# Helical foldamers as selective G-quadruplex ligands

**DOI:** 10.1101/2025.02.25.640101

**Authors:** Alexander König, Vincent Laffilé, Stéphane Thore, Cameron D. Mackereth, Liliya Yatsunyk, Yann Ferrand, Eric Largy, Valérie Gabelica

## Abstract

We investigated the G-quadruplex binding selectivity of short aromatic oligoamide helical foldamers comprising quinoline (Q) and pyridine (P) units. We found that the foldamers prefer parallel G-quadruplex structures, especially when the external G-quartets are sterically accessible. A crystal structure of the tetramer QQPQ with the parallel G-quadruplex formed by dTGGGTTGGGTTGGGTTGGGT shows two quinoline subunits interacting with an external G-quartet through π-stacking, and solution NMR confirms that the foldamer targets the 3’ and 5’ ends of this G-quadruplex. Foldamers can also selectively target sequence variants of the telomeric sequences containing adenine-to-thymine mutation in the loops. The conformational selectivity of foldamers originates from the bulkiness of oligomers with four or more subunits, which imposes steric restrictions on G-quadruplex binding. The flexibility provided by the pyridine subunits was also key to improve affinity. Mixed quinoline-pyridine foldamers are thus a promising class of selective G-quadruplex ligands, and their unique modular scaffold offers new avenues to further improve their affinity and selectivity.

## Introduction

G-quadruplexes (G4s) are non-canonical DNA structures wherein four guanines associate into a quartet through Hoogsten-type H-bonding.^1^ G4 folding topologies are diverse and some sequences can form several energetically close G4 topologies.^2–7^ The structure(s) adopted by a given DNA sequence can depend on the concentration of DNA strand, the type and concentration of cation^2,8–10^ or on the ionic strength and pH of the buffer.^11,12^ Over the past decades, the solution-phase structures of several G4s have been resolved.^13–15^ We will use a previously assembled database of resolved G4 structures featuring circular dichroism (CD), mass spectrometry (MS), UV-melting and nuclear magnetic resonance (NMR) data in both NMR and electrospray ionization mass spectrometry (ESI-MS) compatible conditions.^16^

G4-forming sequences are abundant in promoters and at the telomeres, opening avenues to develop G4-targeting ligands as antiproliferative agents or to regulate gene expression.^17–24^ G4 ligands are generally flat, condensed heteroaromatic systems that allow efficient π-stacking on G-quartets while inhibiting dsDNA intercalation.^25^ Cationic side chains are often added to improve water solubility and favor electrostatic interactions with the nucleic acid backbone.^26^ However, while G4 selectivity over duplexes is often achieved, most ligands lack selectivity among different G4s.^27,28^ Ligands discriminating certain topologies^29–31^ or topology-subclasses^32–34^ have already been reported but rational design efforts have not yet yielded small molecules with significant selectivity for a specific G4 structure.^35^ Molecular scaffolds that deviate from the condensed aromatic structure paradigm may be required to produce ligands with improved topology selectivity.^32^

Inspired by biopolymers, foldamers are synthetic oligomers self-organized into a defined folded structure.^36^ Here we studied quinoline oligoamide foldamers that fold into a helix driven by intramolecular forces including electrostatic repulsions, intramolecular hydrogen bonds, conjugation and extensive aromatic stacking.^37–39^ In the absence of any chirality inducers, helical foldamers exist as a racemic mixture of left- and right-handed helices.^40^ Foldamers possess two characteristic traits of G4 ligands: 1) an aromatic heterocyclic core structure, which is derived from quinoline and 2) extended, flexible side chains. But unlike most G4 ligands, foldamers are not flat. Our foldamers have a positively charged side chain (derived from ornithine) to enhance water solubility (Figure 1). We also introduced pyridine monomers devoid of any side chain to increase the flexibility of the helices.

**Figure 1.**
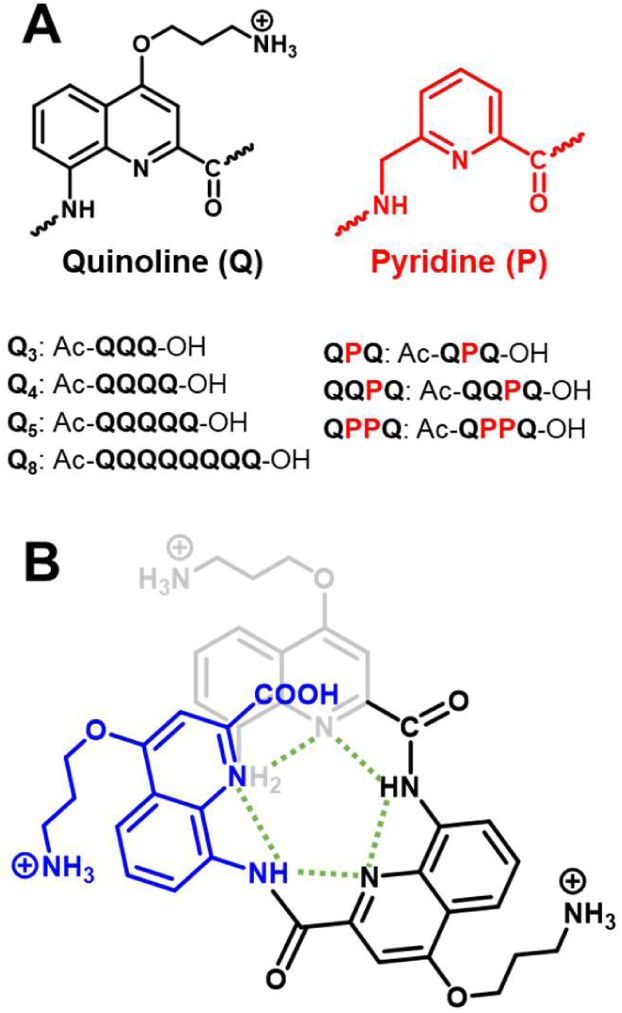
**A)** Oligoquinoline foldamers used in this study, containing quinoline (Q) and pyridine (P) subunits. (panel B shows Q3) **B)** Intramolecular H-bonds (green) sterically prohibit a planar structure, forcing the molecule into a (racemic) helical assembly.

The interaction between oligoquinoline foldamers and G4s has been studied before. They stabilize G4s, but have no effect on DNA duplexes, according to FRET-melting studies.^37,38,41^ Selectivity among different G4 sequences was observed, but not explored further in terms of structure, as the main aim was to demonstrate foldamers targeting G4s over other secondary structures. The takeaway lessons from these previous studies are: 1) Negatively charged foldamers do not interact with G4s, likely due to charge repulsion.^38,42^ Hence, aromatic foldamers designed as G4 ligands should carry positive charges, which are implemented through the side chain. 2) The effect of foldamer helicity on G4 binding is uncertain. It was probed by inducing the handedness quantitatively into a left (*M*) or right (*P*) configuration by covalently attaching either *(R)* or *(S)*-camphanic acid, respectively. However, introducing the camphanyl residue disrupts the foldamer-G4 interaction, indicating that the G4-foldamer interaction site is located at the *N*-terminus of the foldamer.^42^ Helicenes derived from diazaoxatriangulenium (DAOTA) were partially separated using diastereoselective precipitation. The difference in K_D_ value for the *(M)* and *(P)* helices was less than factor 1.5.^43^ 3) Foldamers and antiparallel G4s can co-crystallize without any specific ligand-target interaction.^39^

Here, by native mass spectrometry, we serendipitously rediscovered the high binding affinity of foldamer Q_4_ with the parallel-forming sequence 1XAV (from the *c-myc* promoter region), and large affinity differences with variants of the telomeric sequence. We therefore conducted a systematic investigation of a larger panel of sequences with several foldamers, and detailed biophysical and structural investigations to understand the determinants of sequence or topology preferences.

## Experimental methods

### Foldamer synthesis

Seven foldamer sequences (Figures S1—S2) were prepared according to the synthetic schemes in the supporting information (Figures S3—S6). NMR spectra, HPLC traces and accurate masses for each foldamer are provided in Figures S7—S20.

The aromatic oligoamides were synthesized using a microwave-assisted solid phase synthesis approach. The matrix is a low loading Cl-MPA ProTide® resin. The first monomer unit was attached to the resin using CsI/DIEA. Subsequently, each monomer was coupled to the oligomer chain iteratively through periodic cycles of deprotection. Coupling to the next monomer unit was performed using Appel conditions (PPh_3_/trichloroacetonitrile) with collidine as base. Finally, trifluoroacetic acid was used to simultaneously cleave the oligomer from the resin and deprotect the Boc groups of the side chains to yield the corresponding ammonium salts. After filtration of the remaining solids, the TFA solution was evaporated then the residue was suspended in Et_2_O and centrifuged. The solid was finally dissolved in water and freeze-dried.

All sequences were purified using semi-preparative HPLC on a reverse phase C18 column (mobile phase: H_2_O/ACN/TFA). The purified foldamers were freeze-dried twice more to remove TFA traces. The final product was obtained as a yellow solid with a cotton-like texture, consisting of the foldamer with one trifluoroacetate counterion per ammonium groups. Foldamer stock solutions were prepared by weighing in the purified product on a microbalance (Sartorius ME5) and dissolving them in UPLC/MS grade pure water.

### Oligonucleotides and sample preparation

All oligonucleotide sequences were purchased from Eurogentec (Belgium) and dissolved in UPLC/MS grade pure water (Biosolve Chimie, France) to 1 mM DNA. Stock solutions were annealed for 3 minutes at 85°C, then any residual Na^+^ ions were exchanged with 500 mM NH_4_OAc, which we then flushed out with water using cellulose-matrix 3K centrifugal filter units (MerckMillipore, Ireland). We validated the desalting method by analyzing a DNA stock solution before and after desalting (Figure S21). The concentration of each DNA stock solution was determined with a UV spectrophotometer (Uvikon XL Secomam), utilizing extinction coefficients at 260 nm that were calculated from the nucleotide sequence via nearest-neighbor method.^44^ An overview of the sequences studied is provided in Table 1. To form G4, DNA solutions were doped with either potassium chloride from Sigma Aldrich and Trimethylammonium acetate solution (TMAA) from ChemCruz or ammonium acetate solution (NH_4_OAc) from Sigma Aldrich (Saint-Quentin-Fallavier, France).

**Table 1.**
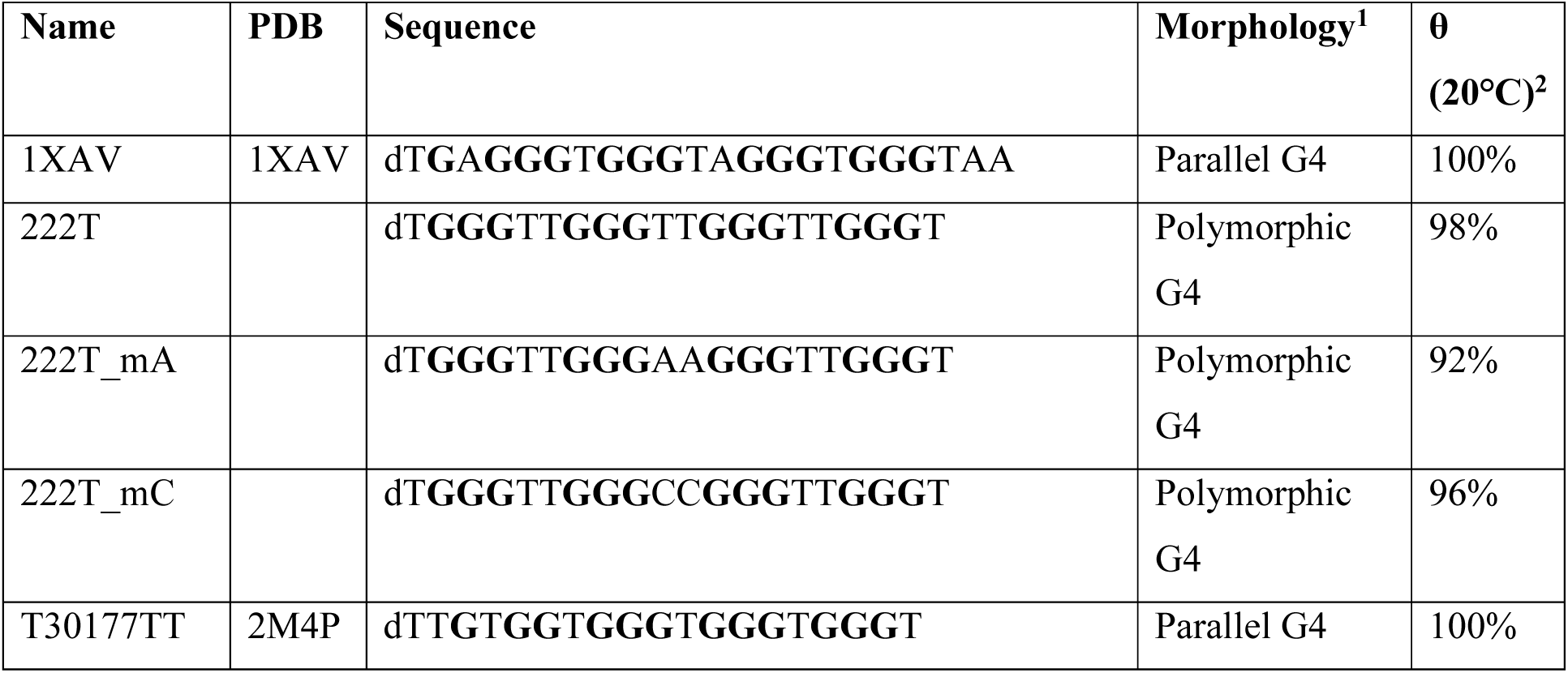
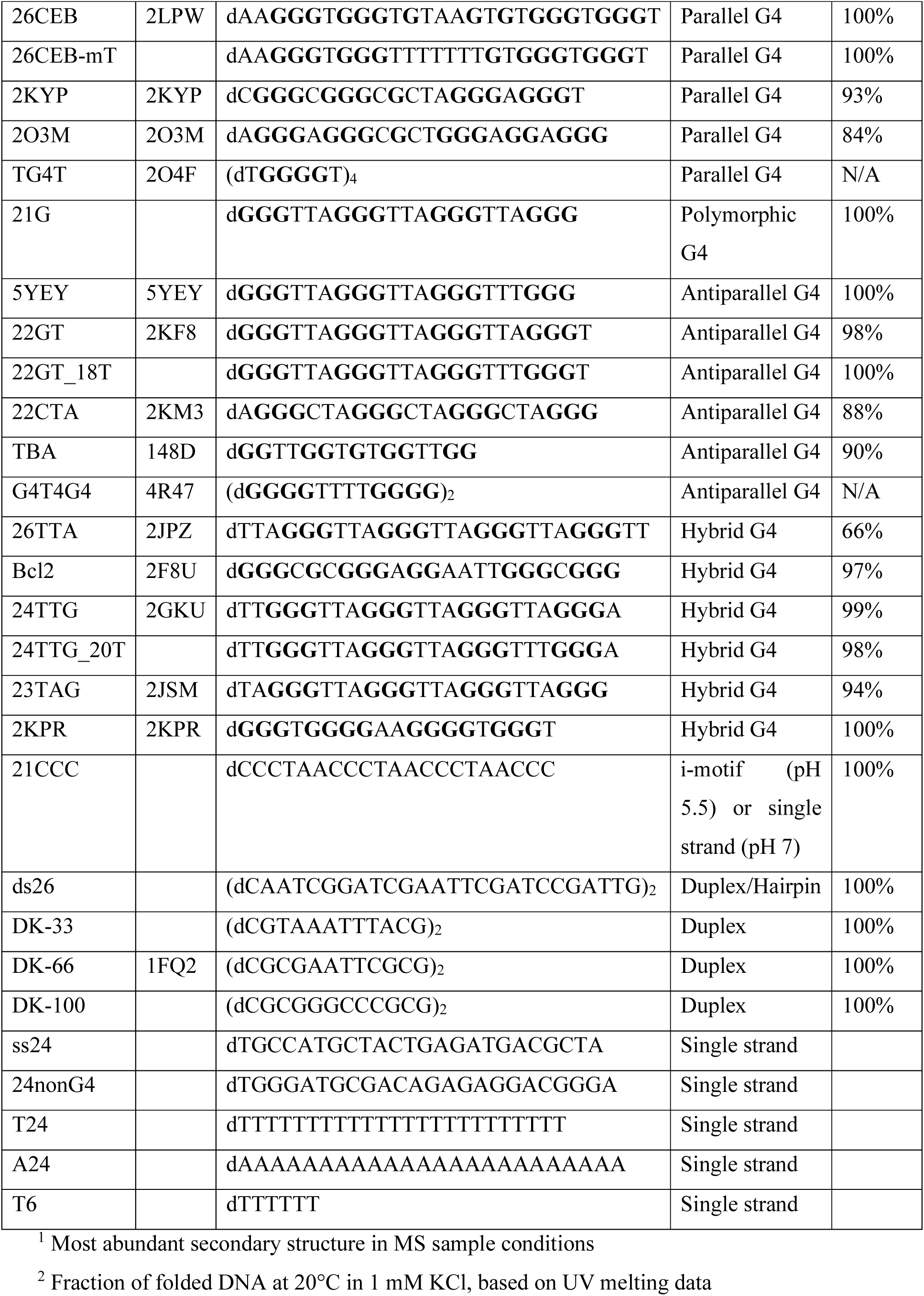
DNA sequences used for the foldamer screening experiments.

### Circular Dichroism spectroscopy

CD measurements were performed on a JASCO J-815 spectrophotometer equipped with a Lauda RE 305 temperature control system with the following parameters: 220-500 nm scan range, 50 nm/min scanning speed, 0.2 nm data pitch, 2 nm bandwidth, 2 s data integration time, 22°C temperature in the sample holder, 3 accumulations. The samples, containing 10 µM of DNA, were placed in Suprasil quartz cuvettes (Hellma) with 2 or 10 mm pathlength.

Raw data was blank subtracted then converted to molar ellipticity with Equation (1), where *θ* is the ellipticity (mdeg), *c* the molar concentration of oligonucleotide (mol/L), and *l* the path length (cm).

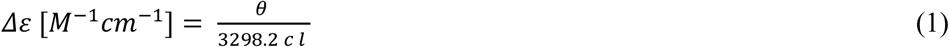

We provide CD spectra in MS conditions for all intramolecular G4 sequences listed in Table 1. They have either been published previously,^16^ or are featured in the supporting information (Figures S22—S29).

### Thermal denaturation

Thermal denaturation was monitored by UV absorption spectroscopy on a SAFAS UVmc2 double-beam spectrophotometer equipped with a high-performance Peltier temperature control unit. The melting ramps were 20→90°C (10 °C/min), 90→4°C (0.2 °C/min), 4→90°C (0.2 °C/min), 90 s data reading interval, 0.5 s averaging time. The samples containing 10 µM of DNA were placed in 10 mm path length Suprasil quartz cuvettes (Hellma). The UV absorption was taken at 260, 295 and 335 nm. The 260 nm wavelength corresponds to the DNA duplex, the 295 nm one to the G4 and the absorption at 335 nm serves as an internal reference to correct for instrumental artifacts. In addition, the buffer alone (100 mM TMAA and 0.5/1 mM KCl) was measured for blank correction.

Raw data, at 260 nm for dsDNA and 295 nm for G4s, was blank subtracted, and corrected with the 335 nm data, then converted to folded fraction *θ* using Equation (2), where *L*0(*T*) and *L*1(*T*) are the baseline values of the unfolded and folded species, respectively.^45,46^

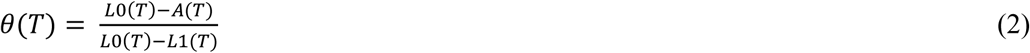

The melting temperature T_M_ is the temperature for which θ = 0.5. We provide *θ*_20°C_ for all intramolecular G4, i-motifs and duplex forming sequences, in MS conditions in Table 1. Melting curves have either been published previously,^16^ or are featured in the supporting information (Figures S22—S27).

### Native mass spectrometry: Instrumental setup

Native MS experiments were conducted on an Agilent 6560 IMS-QTOF. The instrumental settings were optimized to preserve non-covalent DNA structures, albeit making a compromise between signal intensity and softness of the ion transfer (Table S1).^47^ Samples were infused at 3 µL/min for 11 minutes, working foldamer-by-foldamer to avoid cross-contaminations. The instrument is equipped with a drift tube for ion mobility spectrometry. The ions pass through a tube, filled with helium, and we measure the time between ions being released into the drift tube until arriving at the detector. This time is the arrival time t_A_, whose correlation to the collisional cross section (CCS) is described in the Mason-Schamp equation (Equation (3)).

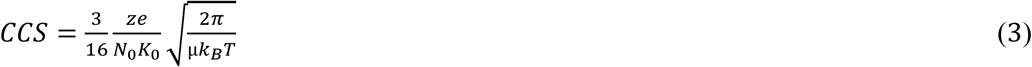

Where *e* is the charge of the electron, *k_B_* is the Boltzmann constant, and *N_0_* is the gas number density (Loschmidt number) at *T*_0_ = 275.15 K and *p*_0_ = 1 atm. The temperature *T* and the reduced mass *µ* (*µ* = (*m*_ion_*m*_He_)/(*m*_ion_+*m*_He_)) remain practically constant within the experimental conditions (details in Table S1). The reduced mobility *K_0_* and charge state *z* differ for every ion species. We obtained the reduced mobility by measuring the arrival time at several drift voltages (Figures S30—S31) and performing a linear fit according to equation (4).

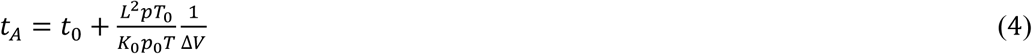

Where *t_0_* is the dead time, *L* is the length of the drift tube (78.1 cm), *p* and *T* are the pressure and temperature inside the drift tube (which are recorded for each experiment) and *K_0_* is the reduced mobility of the ion.

We validated the instrumental settings before each series of measurements using [(dTG_4_T)_4_•(NH_4_)_3_]^z-^ as an external calibrant. We accepted the instrumental settings when the experimental CCS values were within 2% of error to our previously published CCS values (787.5 Å² [5-ion], 735.7 Å² [4-] ion).^48^ The supporting information contains a step-by-step tutorial for how to obtain CCS values and reconstruct CCS distributions (Figures S30—S31 and accompanying text).

### Ligand screening with native mass spectrometry

The panel (Table 1) includes 23 G4-forming sequences featuring different strand stoichiometries (1, 2, 4), numbers of G-quartets (2 to 4), topologies (parallel, antiparallel, hybrid) and origins (*e.g.*, oncogene promoter, telomeres, aptamers). The panel also includes control sequences of alternative secondary DNA structures (duplex, i-motif, single strand). We screened the DNA panel against 4 different foldamer sequences: QPQ, QQPQ, QPPQ and Q_4_. Screening samples contained 10 µM DNA, 20 µM foldamer, 4 µM dT_6_, 0.5 mM KCl and 100 mM TMAA (pH 6.8) in water. 150 mM ammonium acetate (pH 6.8) was used for the sequences T_6_, T_24_, ss24, DK-66 and 21CCC. The multi-stranded G4s TG_4_T and G_4_T_4_G_4_ were screened in both TMAA/KCl and ammonium acetate.

First, each stoichiometry of complex is directly inferred from the mass of the complex. Then, the concentration of DNA species of any stoichiometry (M*) was determined using Equation 5, where *[M]_0_* is the total concentration of DNA in the sample and *I* is the intensity of the different species, obtained by integrating all signal within a specified *m/z* window (monomeric DNA: *M*, ligand: *L*).^49^ The mass spectra in the supporting information show the *m/z* selection windows. The concentration of unbound foldamer is determined by difference (Eq. 6), where *[L]_0_* is the total concentration of foldamer in the sample. The dissociation constants *K*_D1_ and *K*_D2_ values for the 1:1 and 2:1 (*L*:*M*) complexes were then determined from Equations 7–8. The fraction of bound DNA (Eq. 9) is a more concise parameter to present ligand screening results.

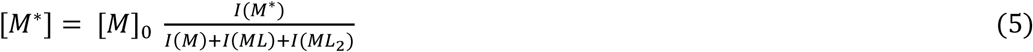

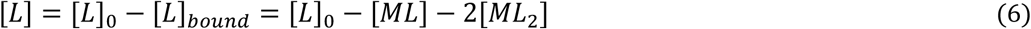

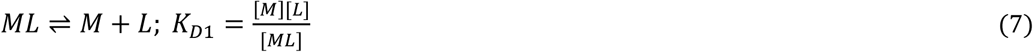

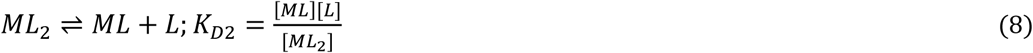

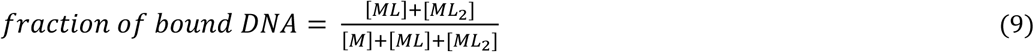

The difference in K_D1_ and K_D2_ highlights ligand cooperativity. We define a positive cooperativity when *K*_D2_ < 4 *K*_D1_ and negative cooperativity when *K*_D2_ > 4 *K*_D1_.^50^

### ESI-MS titrations

The six sequences 1XAV/222T/21G/5YEY/T24/ss24 were titrated with seven foldamers: QPQ, QPPQ, QQPQ, QQQ (Q_3_), QQQQ (Q_4_), QQQQQ (Q_5_) and QQQQQQQQ (Q_8_). The MS titration experiments follow two main objectives: 1) Obtain average *K*_D_ values from a set of measurements alongside with a standard deviation and 2) Study the effect of foldamer length on G4 affinity to aid rational ligand design. We kept the DNA concentration at 10 µM and measured at seven different foldamer concentrations: 0, 5, 10, 15, 20, 30 and 40 µM.

The data processing consists of three steps:

1. Noise correction. Three times the standard deviation of the noise was subtracted from integrated DNA signals. The ligand screening data was noise corrected as well.
2. Response analysis. Response factors depend on the gas-phase transmission yield during electrospray ionization, which can be different for the M, ML and ML_2_ species. We quantify changes in response by adding dT_6_ as an internal calibrant.^51^ The ratio of DNA to dT_6_ intensity as a function of foldamer concentration was monitored and, from changes in that ratio, we derived the effect of foldamer concentration on DNA response (Figure S32). Note that the foldamers do not interact with dT_6_.
3. Dynamic fitting with the DynaFit software (BioKin, MA, USA) which derives a mathematical model from the chemical equations 7 and 8 and matches it to the experimental data with minimal residual error (Figure S33). Response factors of the complexes (ML, ML_2_) were allowed to vary to reflect effects observed at step 2. *K*_D1_ and *K*_D2_ values were extracted from the fitted data.

### Determination of the number of consecutive G-quartets in ligand-free and ligand-bound forms

K^+^ ions bind to G4 forming sequences in two ways: in-between G-quartets (specific) or elsewhere (unspecific). Unspecific K^+^ ions distribution follows a discrete probability, which can be approximated as a Poisson distribution.^52^ Specific K^+^ ions in G4s typically have much higher binding affinities than unspecific ones and have well-defined stoichiometries. An adduct with *n* K^+^ ions is considered specific when it is significantly more populated than its adjacent *n+1* and *n-1* adducts.

For example, in Figure 2, the main adduct of the free DNA is 1 K^+^, which corresponds not only to a 2-quartet G4 (antiparallel),^2^ but also some unfolded DNA stand with non-specific potassium binding. The high relative intensity of the unfolded peak (0 K^+^) is an indicator for an unspecific K^+^ adduct distribution blended within all adduct peaks. The main adduct of the complex is 2 K^+^, which corresponds to a 3-quartet parallel G4. The absence of 0 and 1 K^+^ adducts indicates ligand selectivity towards 3-quartet parallel G4s over 2-quartet antiparallel G4s and unfolded DNA.

**Figure 2.**
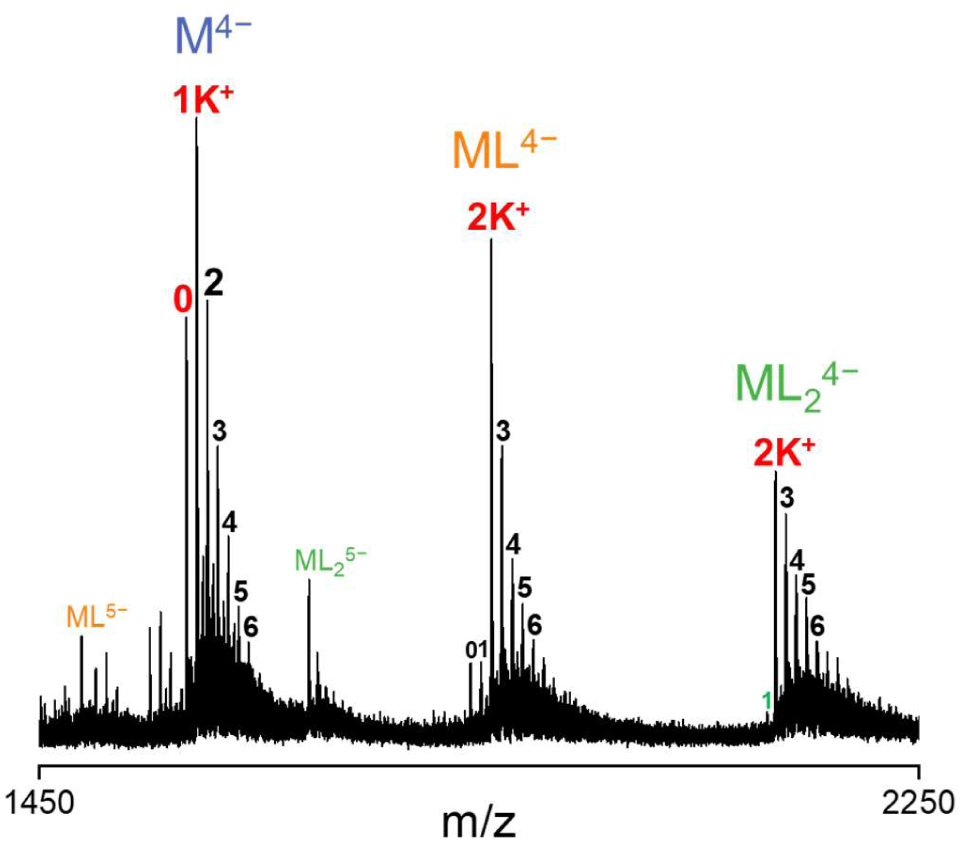
: Mass spectrum of 10 µM 222T_mA (dT**G_3_**TT**G_3_**AA**G_3_**TT**G_3_**T) with 20 µM Q_4_ in 0.5 mM KCl and 100 mM TMAA. Labeled species are: Free DNA (M), 1:1 complex (ML) and 2:1 complex (ML_2_), specific K^+^ adducts (red), non-specific K^+^ adducts (black).

### X-ray crystallography and structure determination

Crystallization was performed in a sitting-drop well plate (SWISSCI MRC 2 Lens Crystallisation Plate). Droplets were obtained by mixing 200 nL of a solution containing 400 µM DNA (222T), 900 µM foldamer (QQPQ) and 20 mM potassium cacodylate (pH 7.0) with an equivalent volume of crystallization screening solutions from various commercial kits. The original condition was the No. 41 of the (Hampton Research) and was composed of 100 mM KCl, 15 mM MgCl_2_, 50 mM Tris (pH 7.5) and 10% PEG-550. We obtained clear, cuboid crystals that were 10-20 µm in size. The crystals were transferred into a cryoprotective solution containing 100 mM KCl, 15 mM MgCl_2_, 50 mM Tris (pH 7.5), 20% PEG-550 and 20% ethylene glycol. Finally, the crystals were flash frozen at -196°C and analyzed at the SOLEIL synchrotron (Saint-Aubin, France).

Several data sets were collected at the French synchrotron Soleil on the beamline Proxima I to a maximum resolution of 2.51 Å. The raw images were processed with XDS.^53^ The structure was solved using the Phaser-Molecular Replacement procedure from the software package Phenix.^54^ The structure was solved using the G4 core from the previously solved structure of the 222T sequence (PDB: 6P45).^55^ We tested various models with and without the connecting nucleotide to help remove model bias. We improved the G4 model by several rounds of manual rebuilding and added the QQPQ foldamer once the electron density from the G4 was filled. QQPQ foldamer restrains were generated using the webserver from GlobalPhasing^56^ and added to the refinement procedure in Phenix. The final model included two potassium ions located between the G-quartets of the same G4 and one potassium ion located between the G-quartets of two dimerizing G4s coordinated by the guanine bases, several magnesium ions and 18 nucleotides of the G4 sequence (missing only the first and the last thymine nucleotide). The summary of data collection and refinement statistics is presented in Table S5. Figures were prepared using ChimeraX.^57^

### NMR spectroscopy

NMR spectra were recorded on a 700 MHz Bruker Avance Neo spectrometer equipped with a triple resonance gradient standard probe, or a 800 MHz Bruker Avance Neo spectrometer equipped with a triple resonance cryoprobe. Topspin version 4.1 (Bruker BioSpin) was used for data collection. Processing and analysis for 1D spectra also used Topspin, whereas the 2D spectra were processed using NMRPipe^58^ followed by analysis with Sparky (T.D. Goddard and D.G. Kneller, University of California). The parallel and monomorphic G4 sequence T95-2T (dT_2_G_3_TG_3_TG_3_TG_3_T, PDB: 2LK7) was used as a substitute for 222T which displayed poor quality NMR spectra.^59^ Identification of key foldamer-binding thymine bases was aided by using three mutated sequences, each of which replaced one thymine with (deoxy)uracil: 2LK7_1U (d**<u>U</u>**TG_3_TG_3_TG_3_TG_3_T), 2LK7_2U (dT**<u>U</u>**G_3_TG_3_TG_3_TG_3_T) and 2LK7_18U (dT_2_G_3_TG_3_TG_3_TG_3_**<u>U</u>**). The 2LK7 and 5YEY NMR samples contained 100 µM DNA in 10 mM potassium phosphate buffer (pH 7, contains 16 mM K^+^) in 90%/10% H_2_O/D_2_O without or with added QQPQ. The 2D ^1^H,^1^H-NOESY used a mixing time of 200 ms, and the ^1^H,^1^H-TOCSY used a mixing time of 60 ms, with both measured at 278 K. Every DNA sequence was measured with and without QQPQ. 1D-NMR spectra of QQPQ alone were measured in d6-DMSO instead of H_2_O/D_2_O and potassium phosphate.

## Results and Discussion

### Foldamer specificity for different nucleic acid secondary structures

To have a first ranking of foldamer binding affinities, we recorded native ESI-MS spectra at one DNA:foldamer concentration ratio. For example, in Figure 3 we can visually compare the mass spectra of foldamers QPPQ and QQPQ with vastly different binding affinities for the same DNA strand 5YEY. The relative abundance of complex (ML) compared to free DNA (M) readily shows that QQPQ binds to the sequence 5YEY with much higher affinity than QPPQ. The calculated *K*_D_ values differ by more than one order of magnitude (Table S2).

**Figure 3.**
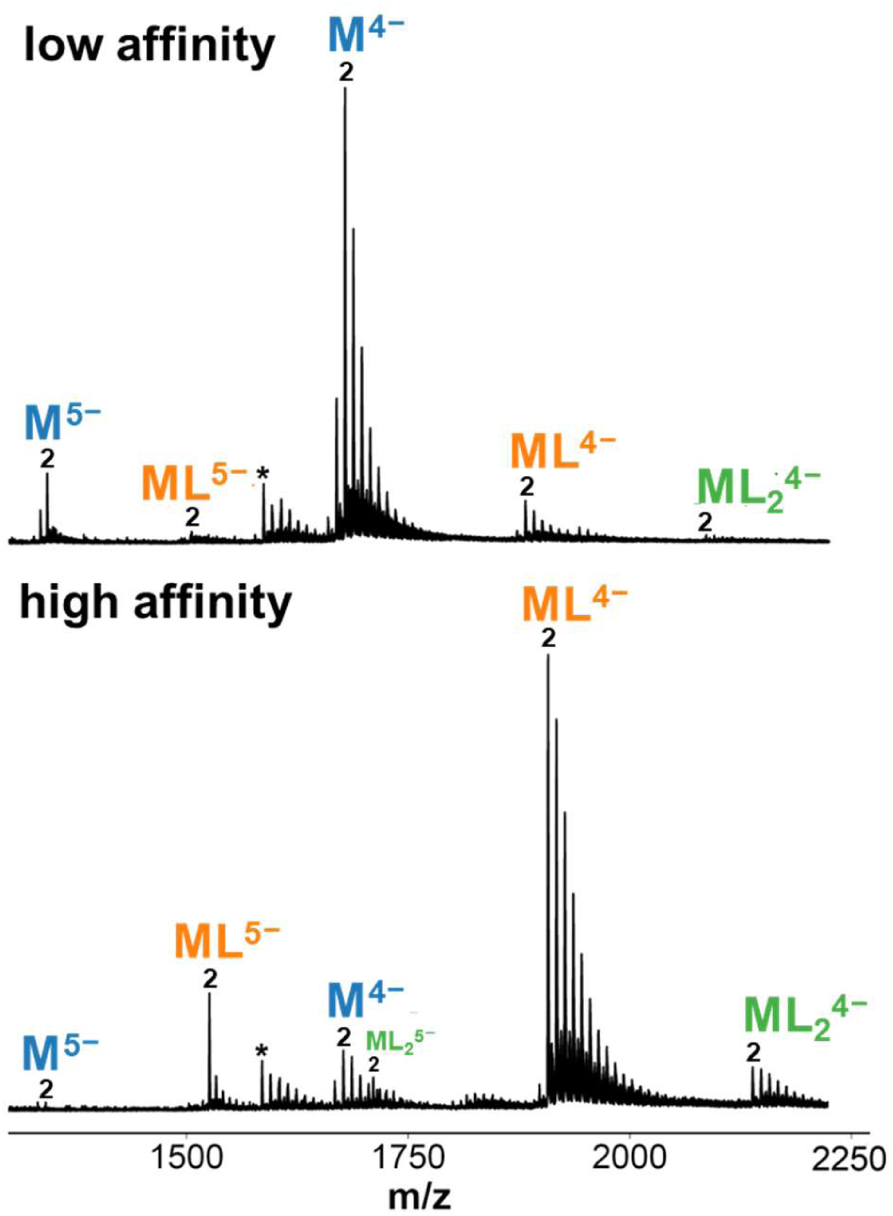
Mass spectra of 10 µM 5YEY (d**G_3_**TTA**G_3_**TTA**G_3_**TTT**G_3_**) with **Top**: 20 µM QPPQ. (K_D1_ = 75 µM, K_D2_ = 40 µM) **Bottom**: 20 µM QQPQ. (K_D1_ = 1.5 µM, K_D2_ = 22 µM). Sample matrix contains: 0.5 mM KCl and 100 mM TMAA (pH 6.8). Labeled species are: Unbound DNA (M), 1:1 complex (ML), 2:1 complex (ML_2_), 2K^+^ adduct (2), depurinated DNA (*).

The low intensity of 2:1 complex compared to the 1:1 complex indicates that the 5YEY G4 only has one accessible binding site. In contrast, foldamer Q_4_ binds to the parallel G4 222T at two binding sites (see Figure 2).

We carried out this general screening with four foldamers (Q_4_, QQPQ, QPPQ and QPQ), and 34 DNA sequences that include well-characterized G4 structures and other DNA secondary structures as controls (i-motif, duplexes, and single-stranded DNA). Figure 4A presents the ligand affinity results as a heatmap. The detailed data, including K_D_ values, are in Table S2 and the mass spectra (showing the foldamer and K^+^ stoichiometries) are shown in Figures S34— S69. Overall, the data indicate that foldamers have a strong preference for parallel G4s and bind very little to DNA controls. Among different foldamers, QQPQ has the most varied affinities and selectivities.

**Figure 4.**
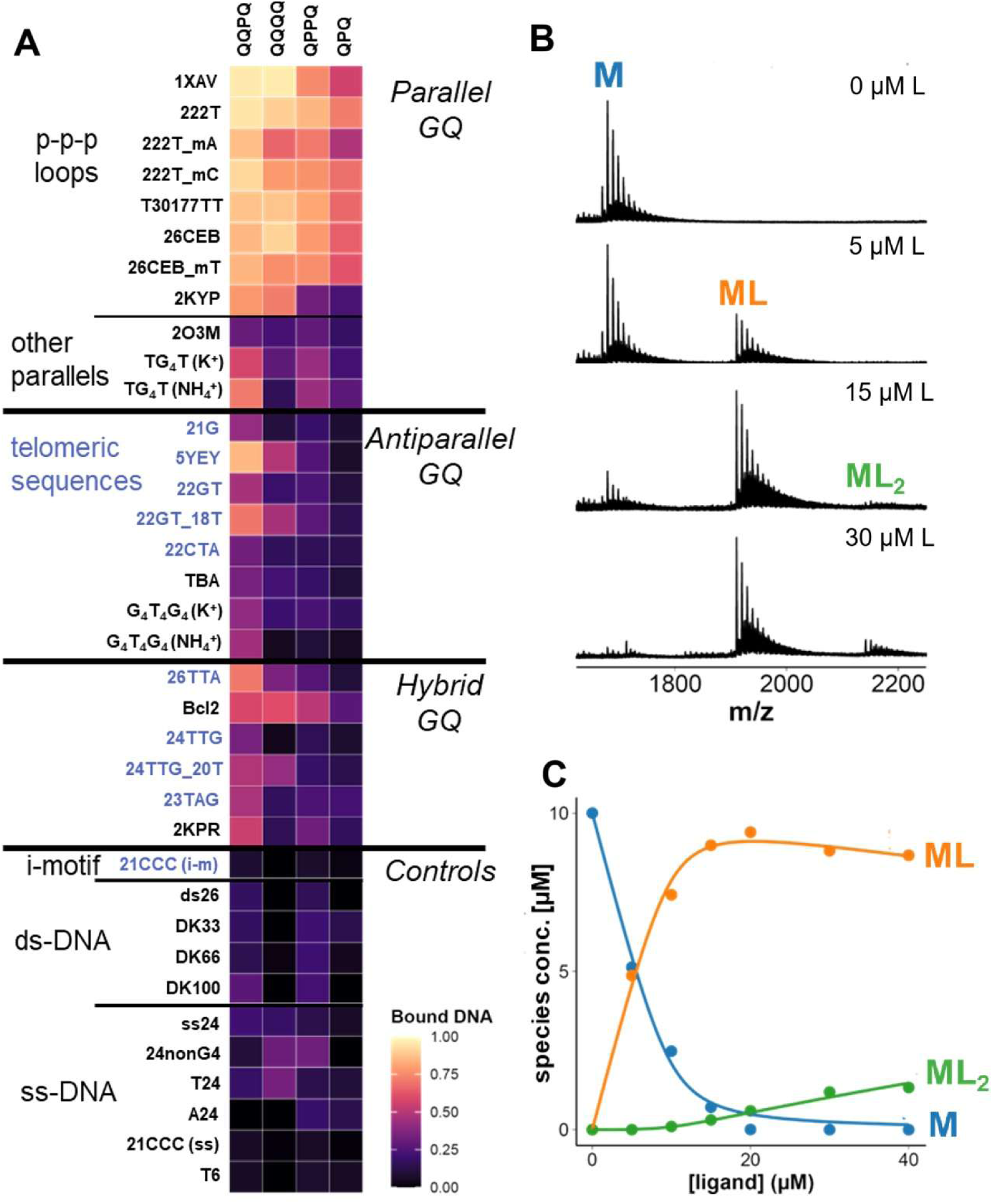
**A)** Heatmap showing the fraction of DNA bound for all DNA/foldamer combinations, calculated from mass spectra. Sample conditions are: 10 µM DNA, 20 µM foldamer, 0.5 mM KCl, 100 mM TMAA (pH 6.8). DNA sequences are listed in Table 1. Telomeric-related sequences are highlighted in blue. **B)** Titration data for 5YEY and QQPQ, showing an extract of the raw mass spectra. **C)** Processed data. The dynamic fitting curves match the experimental datapoints and return K_D1_ = 0.54 ± 0.1 µM, K_D2_ = 200 ± 70 µM.

All four studied foldamers bind parallel G4s with K_D_ values in the low µM range (Table S2). But not all parallel G4s are equivalent. First, 222T_mA and 2KYP are partially unfolded in 0.5 mM K^+^. Therefore, there is a mixture of targetable structures (folded G4) and non-targetable structures (unfolded species). This explains that the fraction of bound DNA is lower compared to the other (fully folded) sequences. Next, the G-quartets of 2O3M and TG4T are less exposed than those of common parallel G4s with three propeller loops. 2O3M has a lateral and a snapback loop can obstruct the two terminal G-quartets. TG4T has eight flanking thymines that can form a sparsely populated T4 tetramer, which also protects the G4 from 5’-5’- dimerization.^60^ We therefore hypothesize that foldamers bind to the external G-quartets, but that loops/bases covering the terminal quartets will impede foldamer binding for steric reasons.

QPQ and QPPQ are highly selective to parallel G4s with three propeller loops but have moderate affinities. These foldamers are more likely crescent-shaped than helical, because the quinoline-oligomer chain is not long enough (it takes 5 Q units for 2 helical turns).^36^ QQPQ and Q_4_ are more stable helices; as indicated by the network of intramolecular H-bonds for QQPQ by NOESY (see NMR section). Their affinity to parallel G4s is 3 to 4 times higher.

Apart from parallel G4s, the QQPQ foldamer tightly binds to 5YEY (K_D1_ = 1.5 µM), but also to TG4T, 22GT_18T and 26TTA (K_D1_ ≈ 5 µM). 5YEY and 22GT_18T are telomeric mutants with a TTA→TTT mutation in the third loop, which induces a switch from a 2-quartet to a 3-quartet antiparallel G4.^61^ QQPQ and Q_4_ are sensitive to this structural switch: The affinities to the mutants is 5-10 times higher than to the ‘wild-type’ sequences (compare: 21G/5YEY, 22GT/22GT_18T, 24TTG/24TTG_20T). 26TTA is a hybrid-2 G4 that is targeted by QQPQ but no other foldamer in this panel. We cannot assess whether QQPQ is selective towards hybrid-2 over hybrid-1 topology.

None of the foldamers interact well with antiparallel G4s, where the G-quartets are usually covered by base triads and/or diagonal loops on both sides (21G, 22GT, 22CTA, TBA, G4T4G4). This finding confirms that foldamers selectively target G4s whose configuration allows exposed G-quartets. The low affinity towards alternative DNA structures (i-motif, duplex, single strand) underlines the G4 specificity of foldamers, except for very specific cases described below. The three-dimensional shape of foldamers prevents duplex intercalation, as they are too bulky to lodge in between base pairs.^43^ Among the control DNA sequences, the foldamers were irresponsive to changes in G-richness, molecularity or strand length.

### Characterization of affinities with ESI-MS titrations

To further characterize binding affinities and stoichiometries, we selected six DNA sequences (two single strands, two parallel G4s, and two derivatives of the telomeric sequences that differ by a single base), and seven foldamers: QPQ, QQPQ, QPPQ, Q_3_, Q_4_, Q_5_ and Q_8_. Figure 4B shows titration data for 5YEY and QQPQ. All K_D_ values are listed in Table S3, and estimated response factors of each DNA:Foldamer complex are in Table S4. The full titration datasets are in Figures S70—S110.

The control sequence ss24 is a single-stranded 24-mer with 25% of each nucleobase. None of the foldamers target this sequence (*K*_D1_ > 100 µM) save for Q_3_, which has significant populations of 1:1 complex, 2:1 complex and 3:1 complex (Figure S98). Q_3_ is thus the least G4-specific ligand in the panel. T24 is another single-stranded control sequence that does not interact with QPQ, QQPQ or QPPQ (K_D_ > 100 µM), but it attracted our attention during the screening with Q_4_. Intriguingly, it forms high-affinity 2:1 complex with Q_3_, Q_4_ and Q_5_ (*K*_D2_ < 1 µM), including a 4:1 complex with Q_3_. Although our foldamers are racemic and display no CD signal on their own, the induced CD in the presence of T24 matches the CD signature of a right-handed foldamer helix (Figure S111). Our hypothesis is that T24 and Q_n_ foldamers associate into a right-handed double helix, wherein the foldamer is dimerized in order to accommodate the full length of the T24 strand. We also tried the RNA counterpart U24 (Figure S112), because U-rich motifs play a role in RNA expression,^62,63^ but there was no induced CD in U24 upon adding Q_n_ foldamers.

The parallel G4s 1XAV and 222T show the highest affinity. All foldamers form high-affinity 1:1 complex with 222T (K_D1_ < 10 µM). The strongest binding partner for 1XAV is Q_4_ (*K*_D1_ = 0.075 ± 0.059 µM), the weakest one is Q_8_ (*K*_D1_ = 28 ± 6.7 µM). All foldamers form 2:1 complex (*K*_D2_ ≈ 1 µM), except for Q_8_ and QPQ. Q_3_ is the only foldamer to form a 3:1 complex, but based on our results with a control sequence ss24, the interaction might not be G4 specific.

Finally, the two telomeric sequences, 21G and 5YEY, are identical, save one nucleotide in the third loop (A in 21G and T in 5YEY). This mutation, however, changes the polymorphic 21G to the antiparallel 5YEY. Q_5_, Q_4_ and QQPQ form a stable 1:1 complex with 5YEY (*K*_D1_ < 10 µM), but not 21G. Q_4_ is the most selective foldamer, with a 20-fold change in *K*_D1_ between 21G and 5YEY. We also compared the sequences 22GT (antiparallel with 2 quartets) and 24TTG (very stable hybrid-1 G4) and the corresponding sequences with the same A→T mutation (22GT_18T and 24TTG_20T), and observed the same increase of affinity for the sequence variant containing the thymine (Figure 4A).

Overall, the ESI-MS titrations indicate that foldamers have the highest affinity (in the low/sub- µM range) towards parallel G4s (1XAV and 222T), alongside a few notable exceptions (5YEY, T24). Foldamer length is important and there seems to be a ‘sweet spot’ at about 4-5 monomers. Having only 3 monomers comes with a decline in affinity (QPQ) or G4 specificity (Q_3_). Having over 5 monomers leads to no affinity increase, but loss of DNA signal that scales with increasing number of Q monomers (Figure S32). The reason is that each Q monomer adds a positive charge to the foldamer, which reduces charge repulsion among DNA polyanions, promoting DNA aggregation. We confirm this hypothesis from 1) seeing precipitation in concentrated DNA solution when an excess of foldamer is added 2) loss of CD signature for a stable (i.e. resistant to disruption) parallel G4 when Q_8_ is added (Figure S113).

### Preferred topologies and foldamer-induced topology switching

Here we focused on a single most promising foldamer, QQPQ and five G4 DNA with differing topologies. We selected QQPQ because 1) it was the foldamer with the highest G4 affinity and 2) we were intrigued by its ability to selectively target some telomeric repeat sequences. We selected telomeric G4 26TTA which adopts hybrid-2 topology and 5YEY, which is antiparallel. We also picked T30177TT as a stable parallel-stranded G4; 222T and 222T_mA which are polymorphic in 0.5 mM K^+^ with a mix of 3-quartet parallel and 2-quartet antiparallel G4.^2^ The equilibrium of 222T in our experimental conditions is shifted towards the parallel form, while the folding of 222T_mA is shifted towards the antiparallel form. We characterized all complexes by mass spectrometry (MS), ion mobility spectrometry (IMS), CD, and thermal stability, and present selected results in Figure 5.

**Figure 5.**
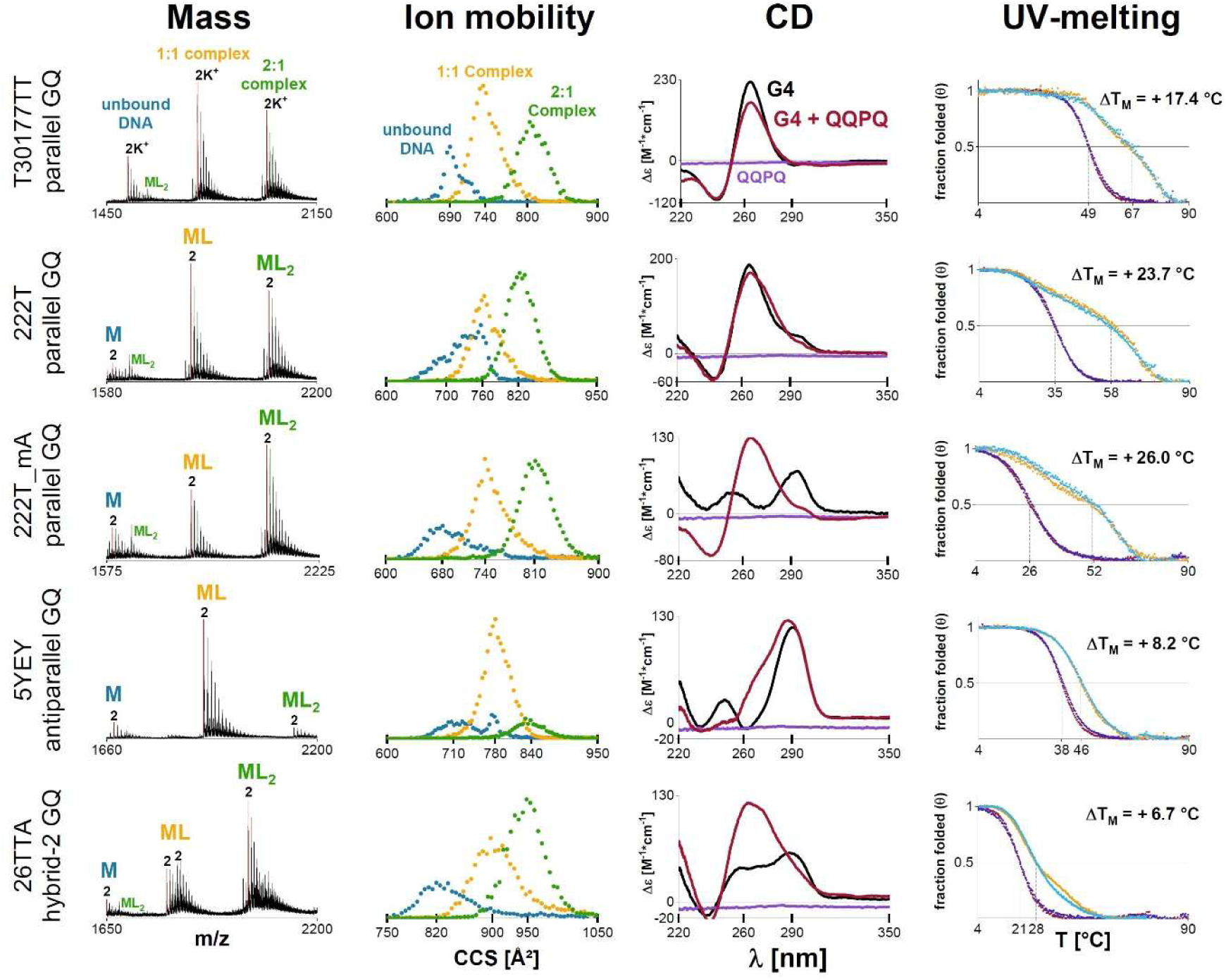
The effect of foldamer QQPQ on five different G4s. Samples contain 10 µM DNA, 20 µM QQPQ, 0.5 mM KCl, 100 mM TMAA (pH 6.8). **1^st^ column**: Mass spectra showing the 4-charge state (5- for 26TTA).**2^nd^ column**: CCS distributions extracted from the 2K^+^ adduct of each species, highlighted with a red line in the corresponding mass spectrum (in column 1). **3^rd^ column**: CD spectra of the complex (red) and control spectra with free DNA (black) and free foldamer (purple). **4^th^ column**: UV-melting curves with foldamer (heating: orange; cooling: cyan) and without foldamer (heating: crimson, cooling: navy).

MS provides a quick readout on ligand stoichiometry, potassium binding stoichiometry and the quantity of each species. The number of specific K^+^ ions correlates to the number of G-quartets. IMS lets us track changes in structure even when there is no change in mass. The ions’ collisional cross section (CCS) depends on the ion size, shape and charge state (within constant experimental settings). Adding ligands is increasing the size of the complex and is expected to lead to a CCS increment. In addition, significant conformational differences for a given size are expected to lead to significant differences in ion mobility, resulting in different CCS distributions. Thus, the shape of the CCS distribution for each ligand stoichiometry indicates how many topologies could be present in solution. CD signatures of DNA alone and with foldamer informs on how the foldamer affects the G4 topology. Finally, the difference in melting temperature with and without foldamer (Δ*T*_m_) provides a quick snapshot of how potent a ligand is at stabilizing the G4 structure. When there are sufficiently populated species with significantly different melting temperatures, multiple melting transitions are visible.

For T30177TT, 2 K^+^ is the dominant adduct in the mass spectra. It has a parallel CD signature and is 100% folded at 25°C. We are thus dealing with a stable, 3-quartet parallel G4. Adding foldamer does not change the K^+^ adduct distribution, the shape of the CCS distribution or the CD signature, meaning that foldamer binding does not significantly change the G4 structure. The slight drop in CD intensity is likely due to foldamer-induced DNA aggregation (Figure S113). The melting temperature of T30177TT increases by 17 °C in the presence of a 2-fold excess of QQPQ, but two distinct melting transitions appear. We know from previous temperature-controlled MS studies on G4 ligands that the 1:1 complex usually melts before the 2:1 complex.^64^ This situation could apply here as well.

222T and 222T_mA are polymorphic in 0.5 mM K^+^. The CCS distributions of free DNA are therefore the sum of multiple Gaussian functions, each representing a different set of conformers. Upon foldamer binding, the CCS distributions are less convoluted (i.e., they look more like a single Gaussian distribution), highlighting that the foldamer selectively targets certain conformers within the conformational ensemble of polymorphic sequences. Figure 5 shows that the complexes (ML/ML_2_) have 2 specific K^+^ ions, therefore the complex must be a 3-quartet G4. QQPQ shifts the CD spectra towards a parallel signature (222T_mA has a remarkable signature switch. The foldamer targets parallel G4 and stabilizes it with Δ*T*_m_ ≈ +26 °C). Similar to T30177TT, the DNA/foldamer system has two separated, broadened melting transitions which likely correspond to unfolding of 1:1 and 2:1 complexes.

26TTA mirrors the behavior of 222T/222T_mA: The K^+^ adduct distribution shifts towards 2K^+^, a parallel CD signature emerges and the UV melting transition broadens in the presence of QQPQ. The foldamer promotes a structural rearrangement of 26TTA, probably from hybrid to mostly parallel topology. As a result, the complex has two binding sites.

The antiparallel 5YEY has only one accessible binding site, as the intensity of 1:1 complex is significantly higher than that of 2:1 complex in the mass spectrum (see also the titration in Figure S87). The foldamer affects G4 folding, causing a CD band at 260 nm to increase. The CD signature indicates that the 1:1 complex could be a hybrid G4, but the CCS distributions give no conclusive evidence whether there is only one hybrid species or multiple species with different topologies. The affinity of the 1:1 complex between 5YEY and QQPQ (K_D1_ = 0.54 ± 0.1 µM) is comparable to that observed for parallel G4s. The melting curve is steeper and shifts less (Δ*T*_m_ = +8.2 °C) compared to those for parallel G4s. That implies that the complex formation with 5YEY is more enthalpy-driven.^46^

### Origin of selectivity for parallel G4s

#### X-ray crystallography

We used X-ray crystallography to test the end-stacking hypothesis based on the CD and native MS results. Molecular crowding induces a bias for the parallel topology,^65,66^ so we selected the parallel G4 sequences 1XAV and 222T to maintain the solution-phase topology and promote crystallization with QQPQ or Q_4_. We also studied the 5YEY/QQPQ and T24/Q_4_ complexes due to their specific binding properties that could provide additional mechanistic insights. We only obtained crystals for the 222T/QQPQ complex, whose structure is shown in Figure 6. Atomic and diffraction data have been deposited in the PDB under the accession code: 8QN2. Crystal and structure refinement information is deposited in Table S5. Electron density maps are shown in Figures S114—S116.

**Figure 6.**
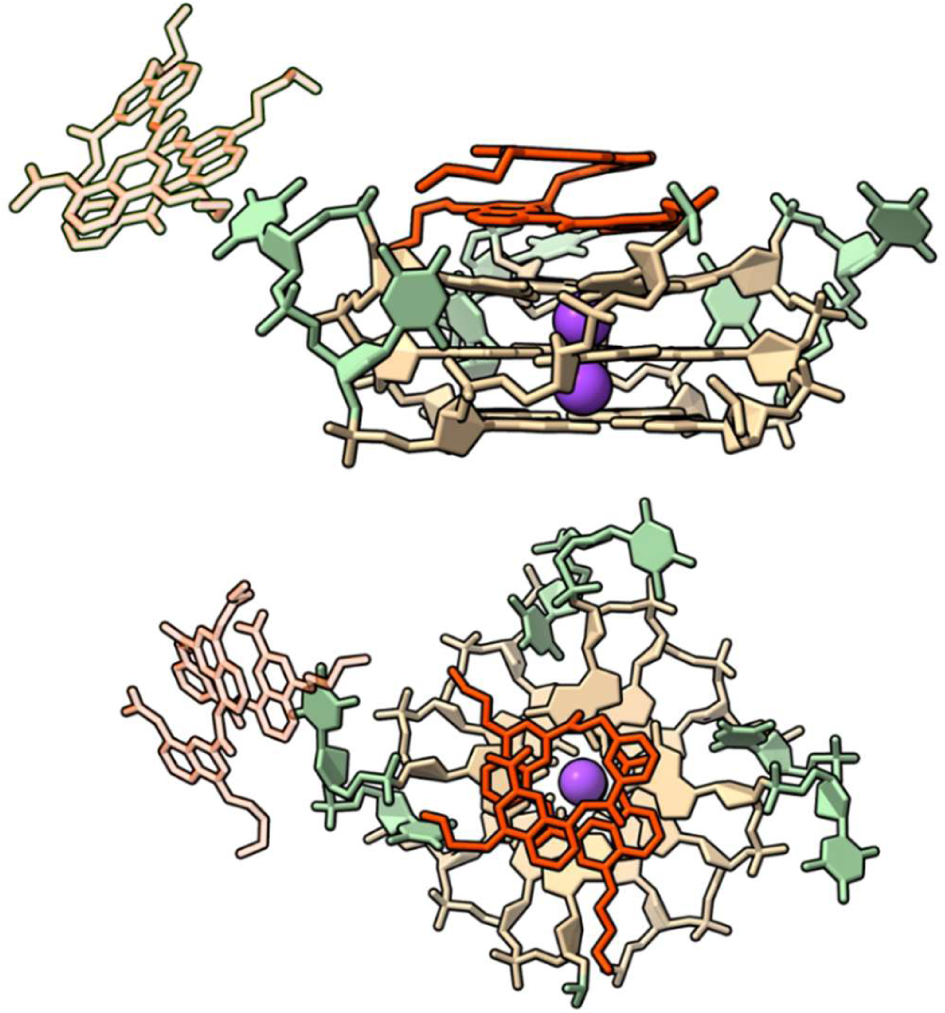
X-ray crystal structure of G4 222T (dT**G_3_**TT**G_3_**TT**G_3_**TT**G_3_**T) and foldamer QQPQ (PDB entry code: 8QN2) from the side (**top**) and from above (**bottom**). Color coding: two QQPQ molecules in red (the binding site not existing in solution phase is set to 50% transparency), guanine in tan, thymine in green, potassium in purple. The crystallization matrix contains 100 mM KCl, 15 mM MgCl_2_, 50 mM Tris (pH 7.5), 10% PEG-550 and a 9:4 ratio of foldamer:DNA.

222T adopts a parallel-stranded topology and forms a 5’-5’ crystallographic dimer. Two QQPQ foldamers interact with one 222T molecule. The first QQPQ has two quinolines π-stacking onto the 3’ G-quartet. The QQ unit is flattened at the binding site to allow more contact area for π-stacking. We suspect that the P unit adds flexibility through its methylene group, which can freely rotate. The flexibility is needed to reduce molecular tension in the foldamer helix caused by the QQ pair flattening. Such molecular architecture explains why QQPQ has the highest G4 affinity: It has two Qs for π-stacking, followed by a P which acts as a step, after which the molecule can resume its helical shape. The top view illustrates how the diameter of the foldamer helix matches the size of a G-quartet. The propeller loops are not limiting the foldamer’s access to the G-quartets and therefore do not interfere with foldamer binding.

According to our ESI-MS titrations, the 222T/QQPQ complex is saturated in a 3-fold excess of foldamer, where it forms a 2:1 complex (Figure S80). We expected to have one ligand on each end of the G4. However, the 5’ interface is utilized for G4 dimer formation and the second foldamer instead stacks onto a thymine base of a TT loop (Figure 6). Parallel G4s tend to dimerize in concentrated solution because of their exposed G-quartets. The preferred interface for dimerization is 5’ to 5’ end.^67,68^ The foldamer sitting atop the thymine establishes a contact with a neighboring unit cell (Figure S117). Such crystal contacts are needed to promote crystal formation and growth, but they can only persist in the solid phase.

The electron density map of the G4 is well-resolved (Figure S114), but the electron density of the foldamer is less defined (high B-factors, see Figure S116), suggesting high dynamics. Several factors can contribute to these dynamics: 1) rotational freedom along the helical axis, 2) exchanges between left- and right-handed helix, since the foldamer is racemic, 3) the foldamer latching onto the G-quartet from its C-terminus or N-terminus, and 4) conformational freedom of the foldamer side chains. We tested models where either the C- or N-terminus of QQPQ interacts with the 3’ G-quartet (Figure S118) and found no significant change in refinement statistics that would suggest any C/N-terminal arrangement being more favorable than the others. So even though the crystal structure depicts a static image of the foldamer-DNA-complex, dynamic exchanges are expected in solution. Alternatively, the foldamer binds to different G4s with different termini and the observed structure is a statistical sum of all arrangements.

In summary, we have validated one binding site being the 3’ G-quartet and the binding mode being π-stacking but need supporting evidence for the second binding site and the foldamer orientation.

### NMR spectroscopy

NMR allows us to investigate foldamer binding to G4 DNA in solution and thus validate (or not) our X-ray structural studies. We could not obtain well-resolved NMR spectra with 222T, but instead used 2LK7 (dTT**G_3_**T**G_3_**T**G_3_**T**G_3_**T), whose structure is very similar to 222T but is more dynamically constrained (1-nt loops) and protected from dimerization (TT motif at 5’).^59^ QQPQ targets 2LK7 with sub-µM affinity (Figure S119, Table S3). All 1D and 2D NMR spectra of 2LK7/QQPQ are gathered in the supporting information (Figures S120—S134).

Our key findings are summarized in Figure 7. Figure 7A shows a ^1^H-^1^H NOESY spectrum of the 2LK7/QQPQ complex. We observe two networks of NOE contacts that correspond to the amide protons of the foldamer (designated *NH1* to *NH4* in Figure 7D). The amide protons are in close contact through the H-bond network that enables the helical foldamer arrangement. The detection of (exchangeable) amide protons in the 1D spectrum (Figure 7C) further indicates that these protons are protected from solvent exchange. These factors support the helical folding of QQPQ in solution phase.

**Figure 7.**
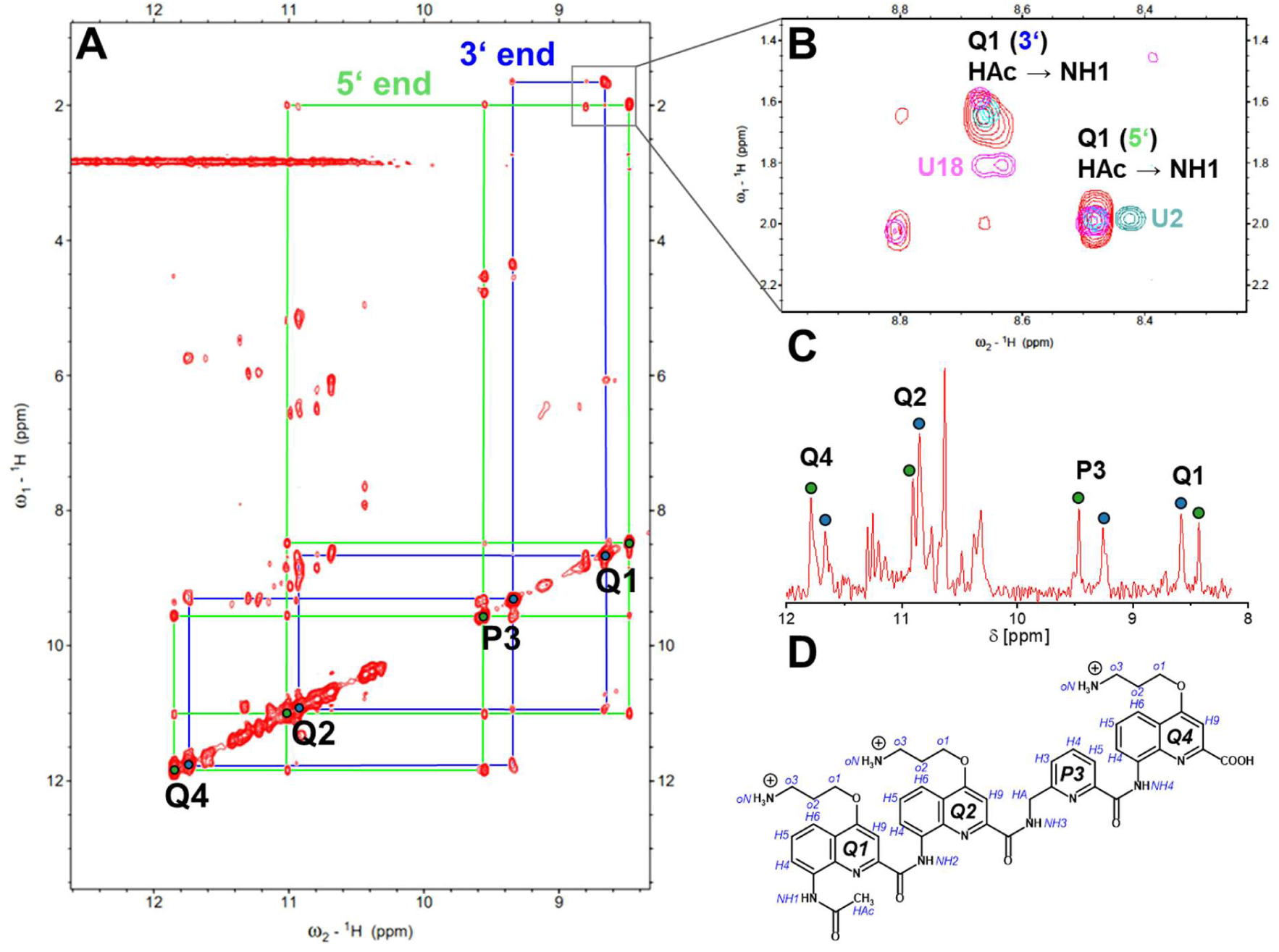
A) ^1^H-NOESY of 2LK7 (dTT**G_3_**T**G_3_**T**G_3_**T**G_3_**T) in excess of QQPQ, showing two NOE networks connecting the foldamer amide protons (NH), indicating that the foldamer is located in two distinct chemical environments. B) Marking the flanking nucleotides through dT→dU mutations induces chemical shifts in the foldamer on the 5’ end (pos. 1,2) and 3’ end (pos. 18) of the G4. C) 1D-NMR of 2LK7 in 3-fold excess of QQPQ, showing the guanine H1-proton region. Foldamer amide protons are highlighted. D) Scheme of QQPQ with proton labels shown in blue. Samples contain 100 µM 2LK7, 300 µM QQPQ, 10 mM KP buffer (pH 7.0) in 90/10 H_2_O/D_2_O.

Figure 7A shows two distinct NOE networks of the same amide protons (lime green and blue). The two significantly different chemical environments for the foldamer must be the first and second binding site, in agreement with the 2:1 complex observed in the mass spectra and our crystal structure. While we do see shifts for the 3’ G-quartet, we do not observe any perturbation of the thymine loop region, indicating that this interaction with loop thymines likely does not take place in solution. A more suitable candidate for the second binding site is the 5’ G-quartet, which is inaccessible in the solid phase because of G4 dimerization.

We then labeled the 5’ and 3’ end of 2LK7 by switching the flanking dT nucleotides with dU (which lacks the characteristic methyl group of thymine). These chemical changes, albeit small, cause visible chemical shifts near each respective thymine base. Figure 7B shows how we assigned a binding site to each NOE network by illustrating the NOE between the amide group (*NH1*) and the methyl group (*HAc*) at the N-terminus of the foldamer. Mutating the 3’ end to uracil (U18) causes a chemical shift in the blue NOE network, while the green NOE network remains unaffected. Likewise, mutating the 5’ end (U2) causes a chemical shift in the green NOE network, while the blue NOE network remains unaffected. Therefore, one foldamer must be close to the 5’ end and another one close to the 3’ end. While the foldamer that interacts with the 3’ G-quartet was clearly seen in the crystal structure, we can now safely deduce that in solution the second foldamer is π-stacking onto the 5’ G-quartet.

### Specificity for variants of the telomeric sequence

In the above characterization of the foldamer-bound G4s based on the telomeric sequence, we observed intriguing specificities for 5YEY, for sequences wherein the third loop is TTT, and also for 26TTA. This latter G4 is not very stable in MS-compatible K^+^ solutions, and therefore the quadruplex can switch its conformation towards a parallel-stranded topology. For 5YEY, our data indicate that QQPQ induces significant base stacking changes in 5YEY (Figure 5). We were thus curious about the structure formed by the 5YEY-QQPQ complex. Unfortunately, we could not crystalize the complex, but we were able to probe the solution states of the free and bound 5YEY by NMR spectroscopy (Figure 8A).

**Figure 8.**
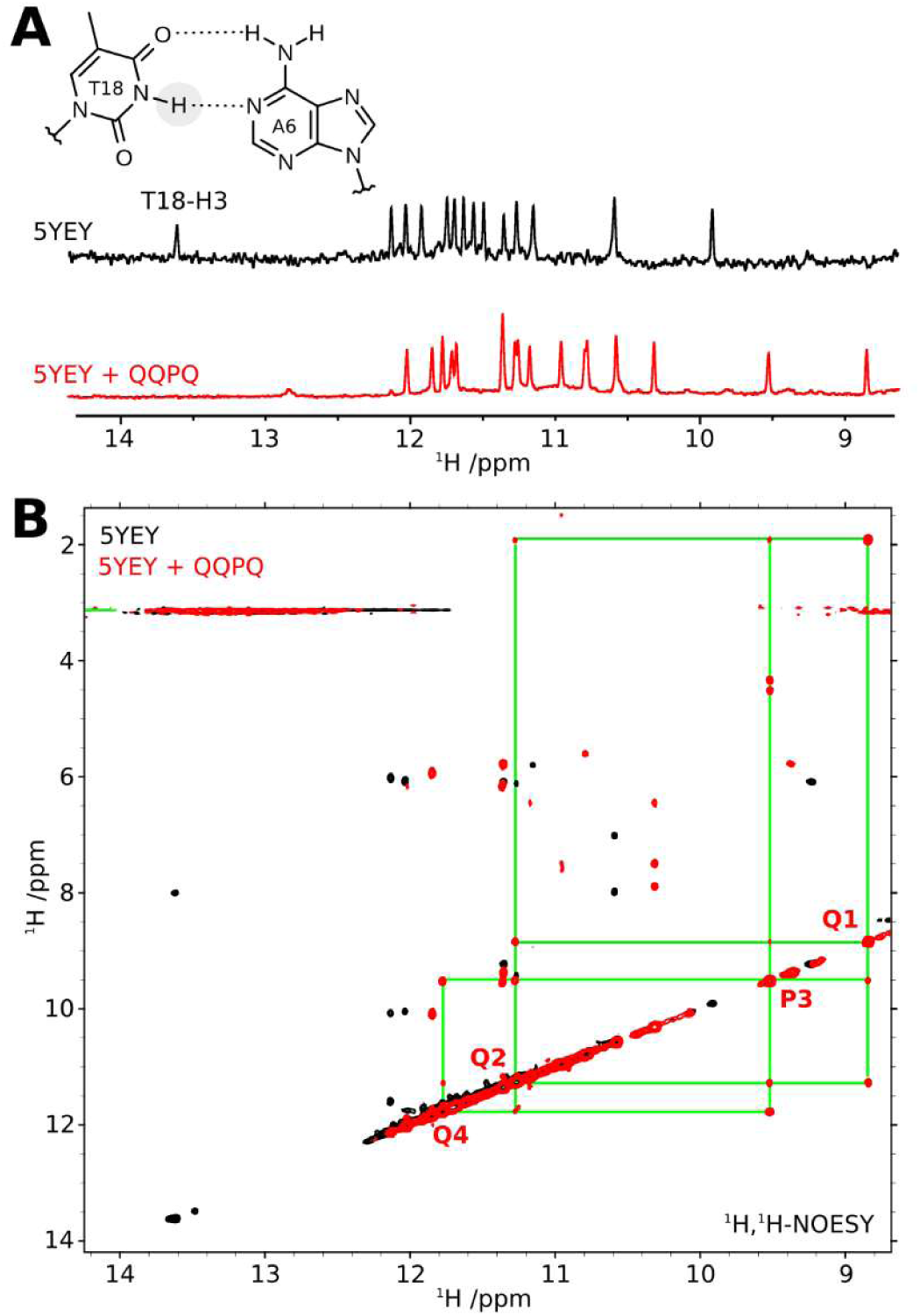
A) In the ligand-free form of 5YEY (black spectrum), the T18-H3 proton is observable and in the region typical of a Watson-Crick base pair (to A6). In the presence of one equivalent of QQPQ (red spectrum), the Watson-Crick peak is no longer present. B) ^1^H,^1^H-NOESY of 5YEY (d**G_3_**TTA**G_3_**TTA**G_3_**TT<u>T</u>**G_3_**) with an excess of QQPQ, showing one NOE network connecting the foldamer amide protons (NH). The single NOE network suggests that the foldamer is located in one distinct chemical environment. 1D spectra were collected at 700 MHz, and 2D spectra at 800 MHz. Samples were measured at 278 K and contained 100 µM 5YEY in 10 mM KP buffer (pH 7.0) in 90/10 H_2_O/D_2_O, without and with 100 µM QQPQ.

NOESY data indicate that only one foldamer NOE network is present (Figure 8B), which reaffirms the 1:1 complex seen in our ESI-MS results (Figure 5). The (lime-colored) NOE network and NMR shifts match well with the lime-colored network in Figure 7, where the foldamer was bound at the 5’ end. Although this observation does not prove that in 5YEY the binding site is also located on the 5’ end, it is consistent with this interpretation.

The 5YEY structure contains an A:T base pair formed between A6 in the first loop and the mutated T18 in the third loop.^69^ A clear indicator of the A:T base pair in the NMR spectrum is the observation of the H-bonded H3 amide proton of T18 within the chemical shift range typical of Watson-Crick base-paired thymine (Figure 8A, top spectrum). In the presence of QQPQ, this characteristic peak disappears from this characteristic region (Figure 8A, bottom spectrum), indicating that the 5YEY/QQPQ complex no longer contains the A:T base pair. Our MS data show 2 specific K^+^ ions for both free 5YEY and the 5YEY/QQPQ complex, therefore both free and bound 5YEY have three G-quartets, but both the circular dichroism spectrum and the ion mobility data suggest that a topology switch could have occurred. Finally, we also recall that the only non-G4 forming sequence to which foldamers significantly bind is the polythymine dT_24_.

We can imagine two scenarios of QQPQ binding to 5YEY. The first scenario is that the G4 topology of 5YEY remains unchanged. The A:T base pair dissociates to make space for the foldamer helix, allowing it to π-stack onto the 5’-opposing G-quartet. However, foldamer binding by stacking on the quartet on the same side of two lateral loops is difficult to reconcile with the observation that, in parallel G4s, steric hindrance on the terminal quartets diminishes the foldamer affinity. We thus favor a second scenario in which the G4 undergoes structural rearrangement. The CD results indicate G-quartet stacking typical of hybrid structures. Hybrid structures typically have a lateral loop on each side of the G-stack core, but recalling the non-negligible affinity of foldamers for polythymines (such as dT_24_), we suggest that the G-quartet on the side of thymine loops (and T or TT overhangs in 22GT_18T and 24TTG_20T) could be the foldamer binding site.

### Molecular dynamics of the complexes

Crystallography experiments indicate that the bound QQPQ remains quite dynamic, showing no clear preference for a specific enantiomer and binding through either its N- or C-terminus. Additionally, these experiments did not allow observation of binding at the 5’-end of the 222T due to its dimerization, nor the peculiar binding to 5YEY. To gain further insights into these aspects, we conducted GPU-enabled one-microsecond molecular dynamics simulations using the OL21 Amber force field (see methods in supporting information). The 1:1 complex from the crystal structure served as a reference, with 222T as a monomer and without the second foldamer interacting with a neighboring unit cell. Based on this reference complex, three other 1:1 complexes were prepared wherein QQPQ binds through its C-terminus (3CM) or as the (P) enantiomer (3NP and 3CP). Additionally, four 2:1 complexes were prepared to study the 5’-end binding: 3NP/5NP, 3NP/5CP, 3NM/5NM, and 3NM/5CM (Figure S135).

MD simulations show that 222T can accommodate QQPQ on both end-quartets without causing steric hindrance or structural modification to either partner (Figures S135—S139). In particular, QQPQ retains its helicity. Both enantiomers bind through their N- and C-termini without clear preference, which aligns with the extensive binding and absence of induced CD. The side chains can bind to the backbone and, to a lower extent sugars (Figures S140—S141). When QQPQ is bound at the 3’-end, the side chains can establish H-bonds with the O4 and O2 acceptors of loop and terminal thymines, but this is almost never the case at the 5’-end (Figure S142).

A notable finding is that QQPQ can rotate along its helical axis (Figures S143—S145), sometimes completing a full rotation within a microsecond of simulation (Figure S146). Additionally, the side chains exhibit significant dynamics and do not form long-lasting interactions with DNA (Figures S140—S142). These observations likely account for the poor electron density of QQPQ in crystallography. Similarly, the 5’ and 3’ dT are highly dynamic, which explains their absence from the crystal data. However, there were a few instances where their N3 H-bound to the amide oxygen of QQPQ, suggesting a particular affinity for thymines (Figure 9 and Figure S147).

**Figure 9:**
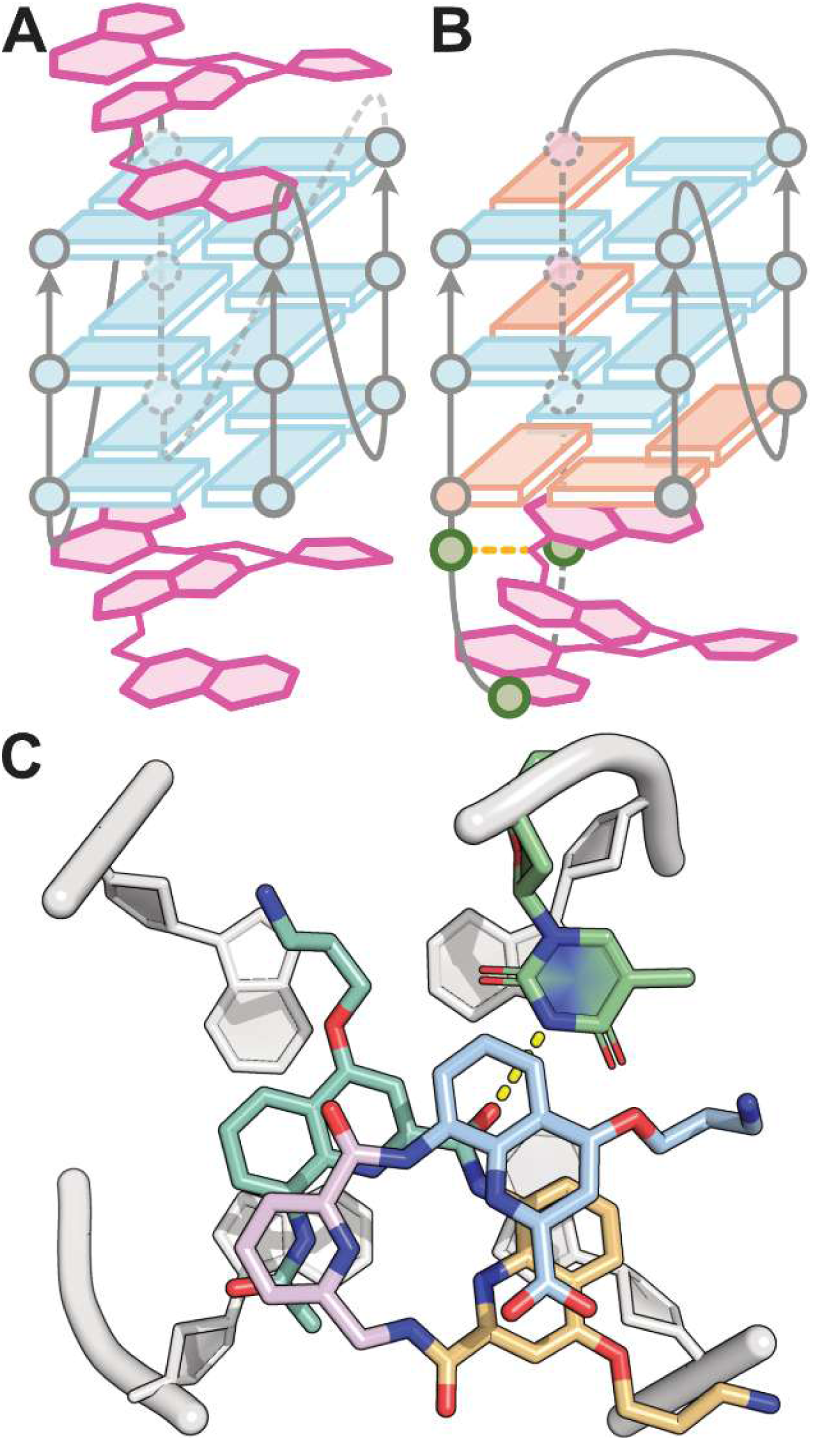
Scheme of the deduced foldamer ligand binding modes. A) Stacking on terminal quartets of a parallel sequence (e.g., 222T), when they are accessible. B) Stacking on a terminal quartet of a non-parallel sequence (e.g., 5YEY), with loop thymines in proximity. C) A snapshot of molecular dynamics started from the crystal structure 8QN2, showing a transient hydrogen bond between an outward pointing amide carbonyl of the foldamer and the N3—H group of a thymine.

Finally, we investigated the binding of QQPQ to 5YEY. A complex was assembled from the hybrid 1 structure of 2JSM, adjusted to match the sequence of 5YEY and capped with QQPQ above the 5’-quartet. The MD trajectory was particularly stable (Figure S148) and provided two key observations supporting the conversion of 5YEY to a hybrid 1 topology with the disruption of the A6:T18 base pair. First, the 5’-quartet of 5YEY can accommodate QQPQ, as it is only occupied by the T16T17T18 lateral loop, which uses minimal space in the absence of the A6:T18 base pair (Figure S149A). Second, the disruption of this base pair allows for the formation of a stable hydrogen bond between the N3 atom of T18 and the amide connecting the N-terminus quinolines of QQPQ (Figure S149A, B). Again, this binding mode is similar to that of the 5’ T1 with 222T described above (Figure S147), and is consistent with the proximity of QQPQ and T2, when the former stacks on the 5’-end of 2LK7 (Figure 8). Finally, T17 stacks over the C-terminus quinoline of QQPQ, further stabilizing the complex (Figure S149C).

The sequence and topology preference of oligoamide quinoline/pyridine containing foldamers can be summarized in Figure 9. The foldamers can stack on terminal G-quartets, preferably when they are sterically accessible. If the two termini are accessible, 2:1 complexes can form (Figure 9A). Exceptionally, binding to some sequences forming non-parallel stranded structures can also occur (e.g., Figure 9B, proposed topology for the 1:1 complex between the 5YEY sequence and QQPQ), provided that there are thymines in the loop, because thymines can form hydrogen bonds with the outward-pointing amide carbonyl groups of the foldamer (Figure 9C, a snapshot extracted from MD simulations).

## Conclusions

Our comprehensive survey of the interaction between G4s and small oligoamide quinoline/pyridine foldamers reveals exciting opportunities to develop sequence- and structure-selective ligands. These foldamers do not target double-stranded DNA, single strands or i-motifs. They selectively target parallel G4s, forming 1:1 and 2:1 complex with sub-μM K_D_ values. The crystal structure reveals two quinoline units in the foldamer helix interact with an exposed G-quartet by π-stacking. The ‘bulkiness’ of the foldamer helix makes it selective for parallel G4 topology with unobstructed 3’ and 5’ G-quartets. The helix clashes with nucleobases surrounding the G-quartet, making it sensitive to loops and flanking nucleotides. We also found that thymines in the loops favor the interaction of foldamers with some specific sequences, offering further opportunities to fine-tune the selectivity.

We believe that through rational design, the structure of the foldamer helix can be adjusted to adapt itself to the terminal G-quartet but also to the surrounding nucleobases. For example, we showed that adding flexibility to the foldamer by incorporating pyridine-based monomers did increase the affinity while preserving G4 specificity. Another intriguing starting point is the particularly high affinity of the QQPQ foldamer to the 5YEY antiparallel variant of the telomeric sequence, which calls for more in-depth structural studies to understand how foldamers could target non-parallel G4s.

As foldamers are made by solid-phase synthesis, other variants can be easily conceived and synthesized. We showed that combinations of four monomer ensured high affinity and some selectivity. Some obvious next steps would be to conceive foldamer compounds with other subunits to modulate the stacking (first two subunits), the flexibility (third subunit), and to explore a variety of sidechains. Overall, our study provides fundamental guidance to those looking to use foldamers as a highly promising ligand scaffold that could be molded to selectively target a G4 target of interest.

## Author Contributions

L.Y. first serendipitously discovered foldamer-G4 interactions and recorded preliminary data. Supervised by Y.F., V.L. synthesized, purified and characterized the foldamer. A.K. prepared and measured the ligand screening, titration, CD and UV melting experiments. A.K. and E.L. carried out MS data processing and visualization. S.T. supervised A.K. during crystallogenesis experiments. S.T. solved and refined the crystal structure. C.M. supervised A.K. for NMR experiments and data processing. A.K. wrote the manuscript with revisions and edits from all authors. V.G. conceived and supervised the entire work and acquired funding.

## Data availability statement

The data underlying this article are available in Zenodo, at https://doi.org/10.5281/zenodo.14964966

## Acknowledgements

We thank Dr. Frédéric Rosu for precious advice on native MS of G4s and useful discussions, Benjamin Lienard and Guilhem Ribera for the technical support during their internship. The work benefited from access to NMR and MS at the “Plateforme de BioPhysico-Chimie Structurale” of the European Institute of Chemistry and Biology (IECB), and the DOREMI CALI v3 cluster of the “Mésocentre de Calcul Intensif Aquitain” (Université de Bordeaux, France). This work was funded by the Agence National de la Recherche (project ANR-18-CE29-0013 POLYnESI to VG and YF) and by the French Ministry of Higher Education and Research (MESRI PhD fellowship to AK). This work was also supported by the Fondation Bettencourt Schueller (Prix Liliane Bettencourt pour les Sciences du Vivant to VG).

## Supporting Information

### Characterization of the foldamers used

#### 1. Materials and methods

##### 1.1. Nuclear Magnetic Resonance

1D NMR spectra of oligomers were recorded on a Bruker Avance NEO NMR spectrometer (Bruker BioSpin) operating at 700,15 MHz for ^1^H observation, equipped with a 5mm TXI probe with a gradient. All NMR experiments were performed at 273 K. Chemical shift values are given in ppm with reference to residual signals of solvent DMSO-d6 (δ=2.50). All coupling constants (J) are given in Hertz and ^1^H NMR splitting patterns with observed first order coupling are designated as singlet (s), broad singlet (brs), doublet (d), triplet (t) or multiplet (m).

##### 1.2. High Performance Liquid Chromatography

HPLC analyses and purification were performed on a reverse phase C8 column on Jasco Extrema analytical and preparative systems. Mobile phases were composed of milli-Q water + 0.1% TFA (solvent A) and Acetonitrile + 0.1% TFA (solvent B). Analyses were done using the following gradient: 0 min: 100% A, 0% B – 2 min: 100% A, 0% B – 12 min: 0% A, 100% B – 15 min: 0% A, 100% B. Purifications were performed using the gradient 0 min: 100% A, 0% B – 2 min: 100% A, 0% B – 32 min: 0% A, 100% B – 35 min: 0% A, 100% B.

##### 1.3. Mass spectrometry analyses

MS characterizations were performed on an Agilent Technologies 6230 TOF LC/MS spectrometer. The instrument is equipped with an ESI source and experiment were recorded in positive mode. The spray voltage was maintained at 3500 V and capillary temperature set at 300 °C. Samples were introduced by injection through a 20 µL sample loop into a 600 µL.min^-1^ flow of acetonitrile from the LC pump.

#### 2. Methods for chemical synthesis

Commercial reagents were purchased from Sigma-Aldrich, Alfa-Aesar or TCI and used without further purification. Low-loading ProTide resin was purchased from CEM. Chloroform (CHCl3), Triethylamine (TEA) and *N*,*N*-diisopropylethylamine (DIEA) were distilled over calcium hydride prior to use. Dry organic solvents: Tetrahydrofuran (THF) and Dichloromethane (DCM), used for solution and solid phase synthesis, were dispensed from a solvent purification system that passes solvents through packed column of dry neutral alumina. Milli-Q water was delivered from a PureLab Prima 7/15/20 system.

##### 2.1. General method for oligomer synthesis

**Figure S1.**
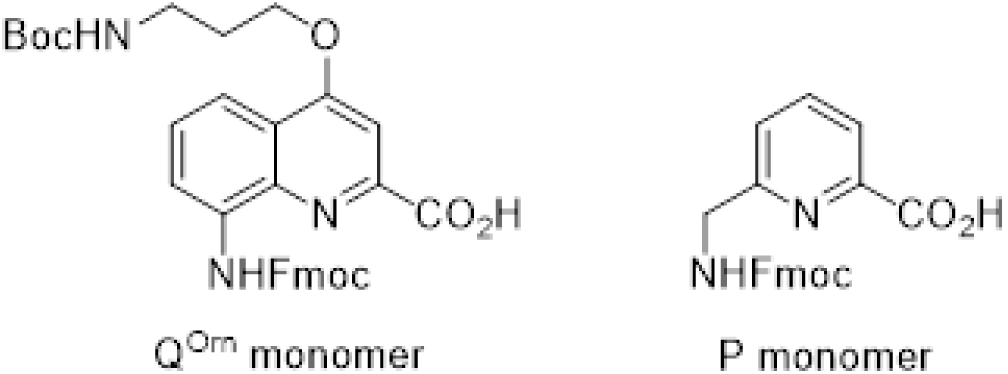
Structures of monomers **Q^Orn^** and **P**. These monomers were synthetized as Fmoc-N-protected and acid free form according to the reported procedure.^1^

**Resin grafting**: On *LL* ProTide resin, the first **Q^Orn^** monomer (3 equiv.) was grafted using CsI (5 equiv.) and DIEA (6 equiv.) in dry DMF. Reaction mixture was vigorously shaking overnight. After reaction, the resin was filtered and washed three times with DMF and dichloromethane.

**Fmoc deprotection**: Grafted resin was washed twice with DMF, suspended in a 20% piperidine in DMF solution (4mL) and slowly stirred for 3 minutes. Resin was then filtered, washed twice with DMF and suspended again in a 20% piperidine in DMF solution and stirred for 7 minutes. The resin was then filtered and washed three times with DMF and dry THF.

***In-situ* coupling procedure**: *For coupling on the aromatic amines of the **Q^Orn^** monomer.* Resin was suspended in dry THF and collidine (9 equiv.) was added. A solution of monomer (3 equiv.), PPh3 (8 equiv.) and trichloroacetonitrile (TCAN, 9 equiv.) in dry CHCl3 was added on the resin. The reaction was assisted by micro-waves (25 W, 50°C) for 15 minutes and repeated once. After reaction, the resin was filtered and washed with dry THF and DMF.

**HBTU coupling**: *For coupling on the aliphatic amines of the **P** monomer*. Resin was suspended in dry DMF. Monomer (3 equiv.) and HBTU (2.9 equiv.) as powder were added followed by DIEA (6 equiv.). The reaction was assisted by micro-waves (50 W, 50°C) for 10 minutes and repeated once. After reaction, the resin was filtered and washed with DMF.

**Resin cleavage**: Resin was washed three times with DMF and dichloromethane and was suspended in a solution of TFA/TIPS/H2O 95:2.5:2.5 (*v/v/v*). The mixture was vigorously stirred for 4 hours. The resin was filtered, and the filtrate was evaporated under reduced pressure. The residual solid was suspended in Et2O and centrifugated at 4°C for 5 minutes. Et2O was removed and the yellow solid was dried under vacuum and then freeze-dried in water.

**Preparative HPLC purifications**: Crude compounds were purified using solvents A and B. The following gradient was used: 0 min: 100% A, 0% B – 2 min: 100% A, 0% B – 22 min: 0% A, 100% B – 27 min: 0% A, 100% B. Collected fractions were analyzed by analytic HPLC and the relevant ones were combined and freeze-dried twice to remove the excess of TFA.

**Figure S2.**
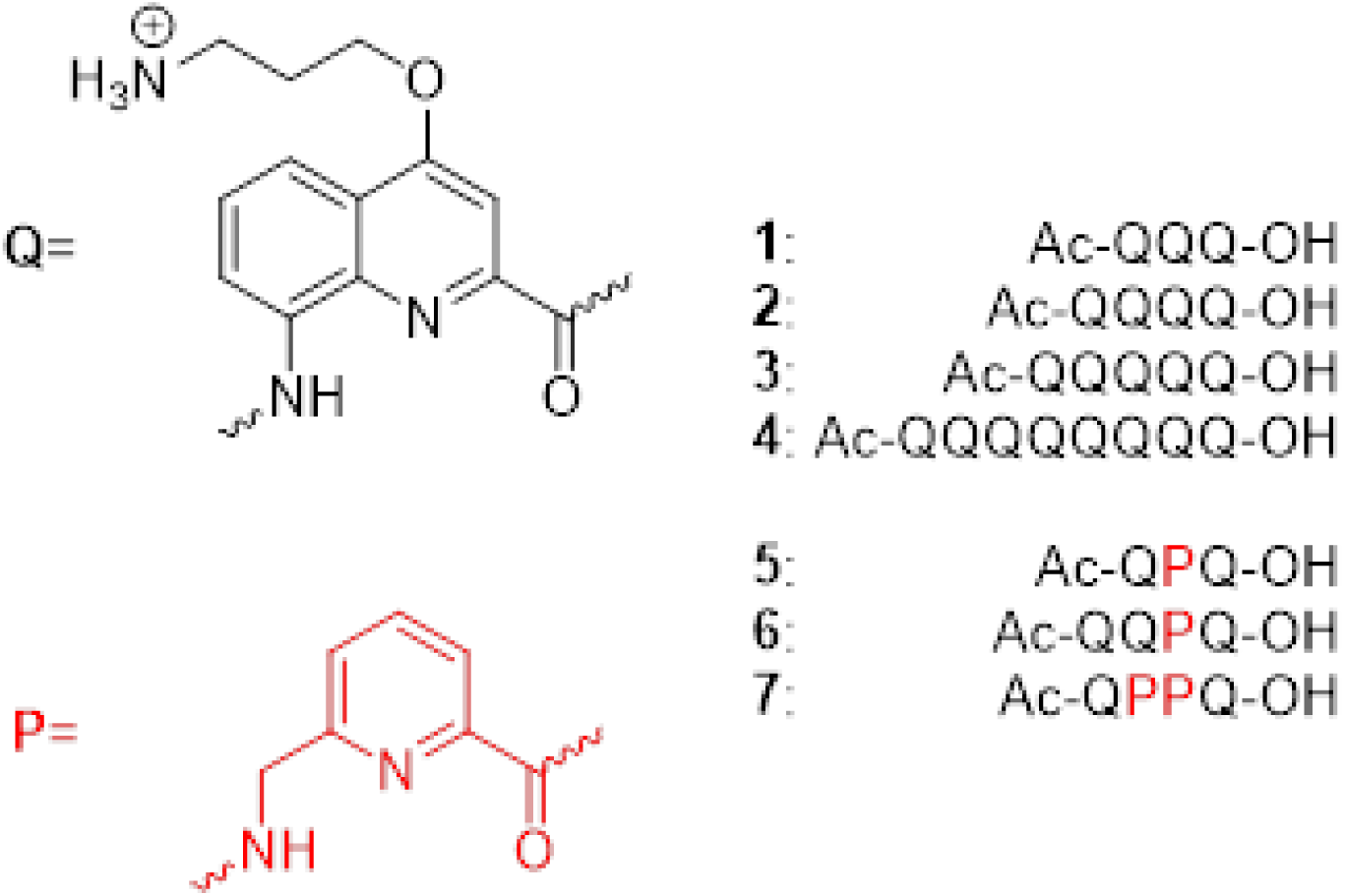
Synthesized oligomers

###### Synthesis of Q3, Q4, Q5 and Q8

**Figure S3.**
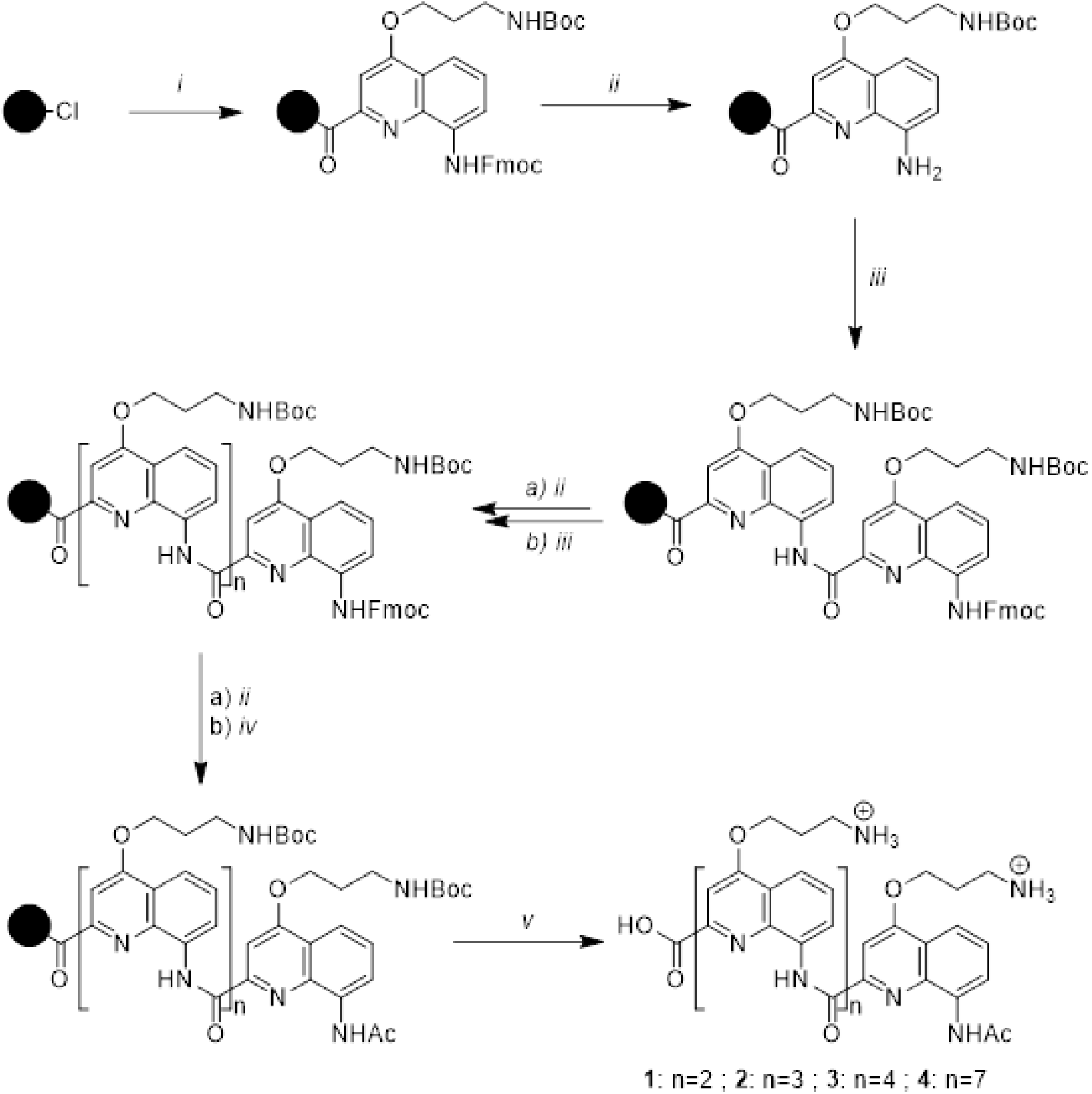
Solid phase synthesis of oligomers **1** to **4.** i) **Q^Orn^** monomer, CsI, DIEA, dry DMF. ii) 2 times, piperidine/DMF 2:8 (v/v), iii) 2 times, **Q^Orn^** monomer, PPh_3_, TCAN, collidine, THF/CHCl_3_. These two last steps are repeated until the desired length is obtained. iv) 2 times, acetyl chloride, DIEA, THF. v) TFA/TIPS/H_2_O 95:2.5:2.5 (v/v/v)

###### Synthesis of QPQ

**Figure S4.**
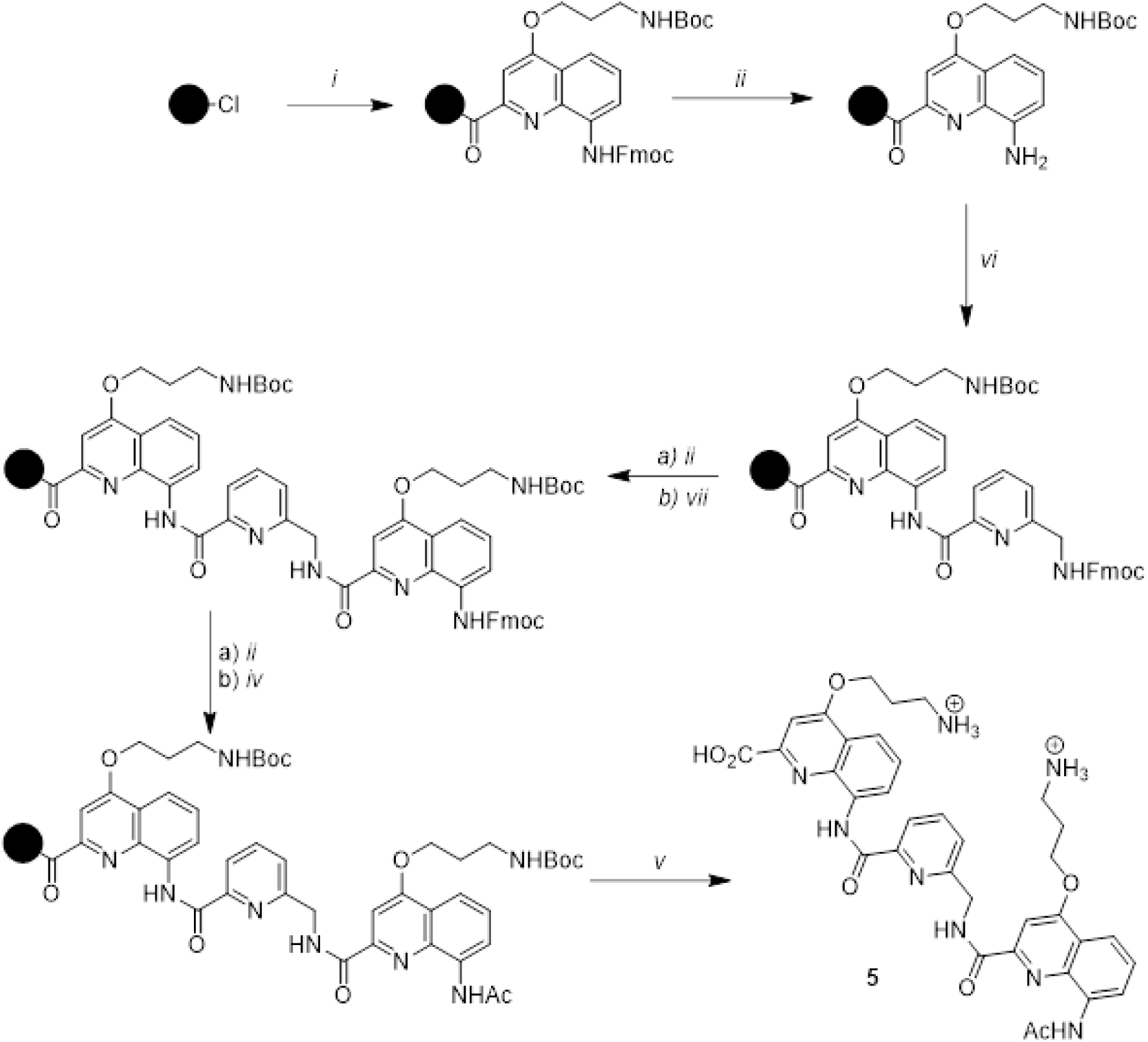
Solid phase synthesis of oligomer **5**. vi) 2 times, **P** monomer, PPh_3_, TCAN, collidine, THF/CHCl_3_. vii) 2 times, **Q^Orn^** monomer, HBTU, DIEA, dry DMF.

###### Synthesis of QQPQ

**Figure S5.**
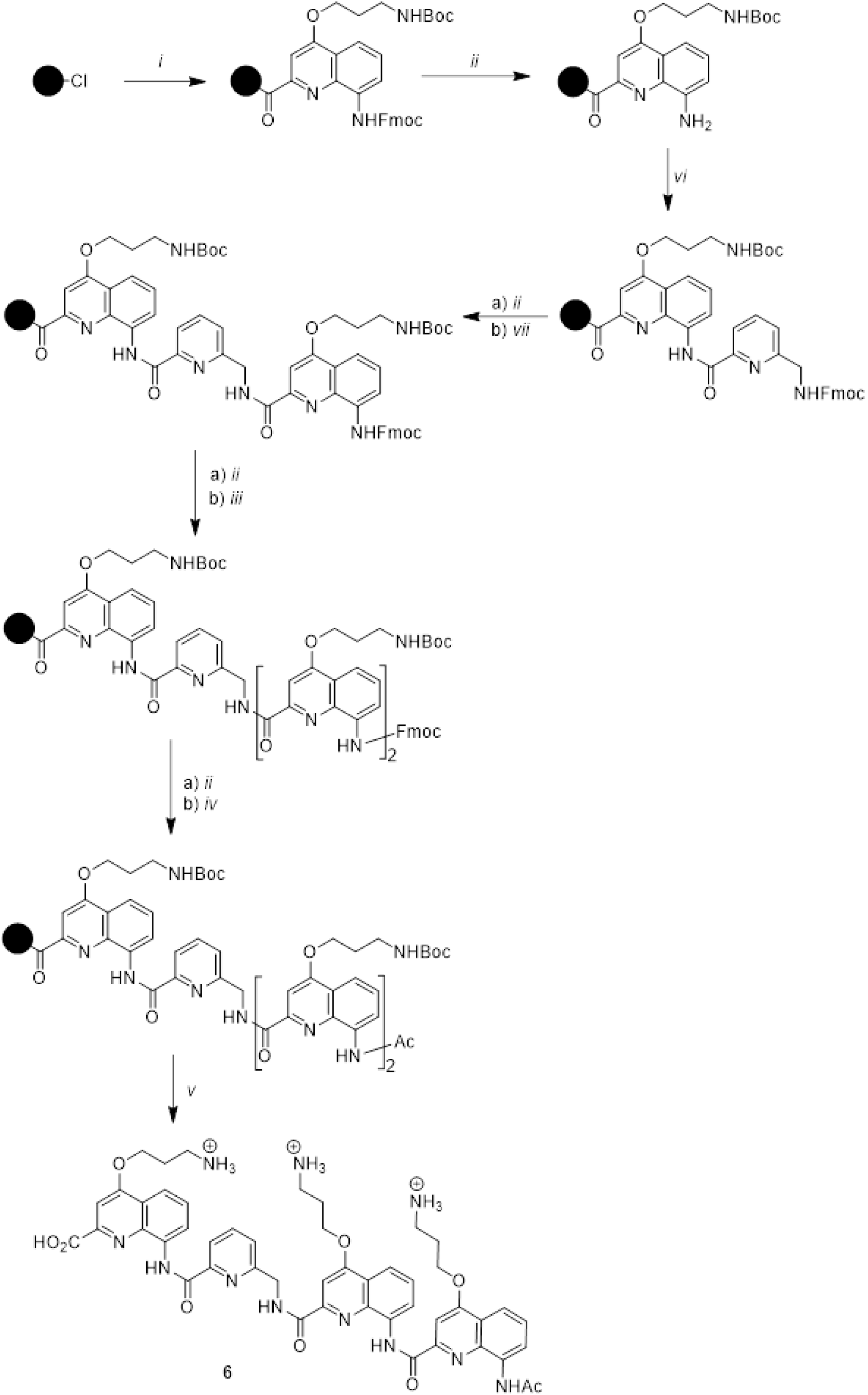
Solid phase synthesis of oligomer **6.**

###### Synthesis of QPPQ

**Figure S6.**
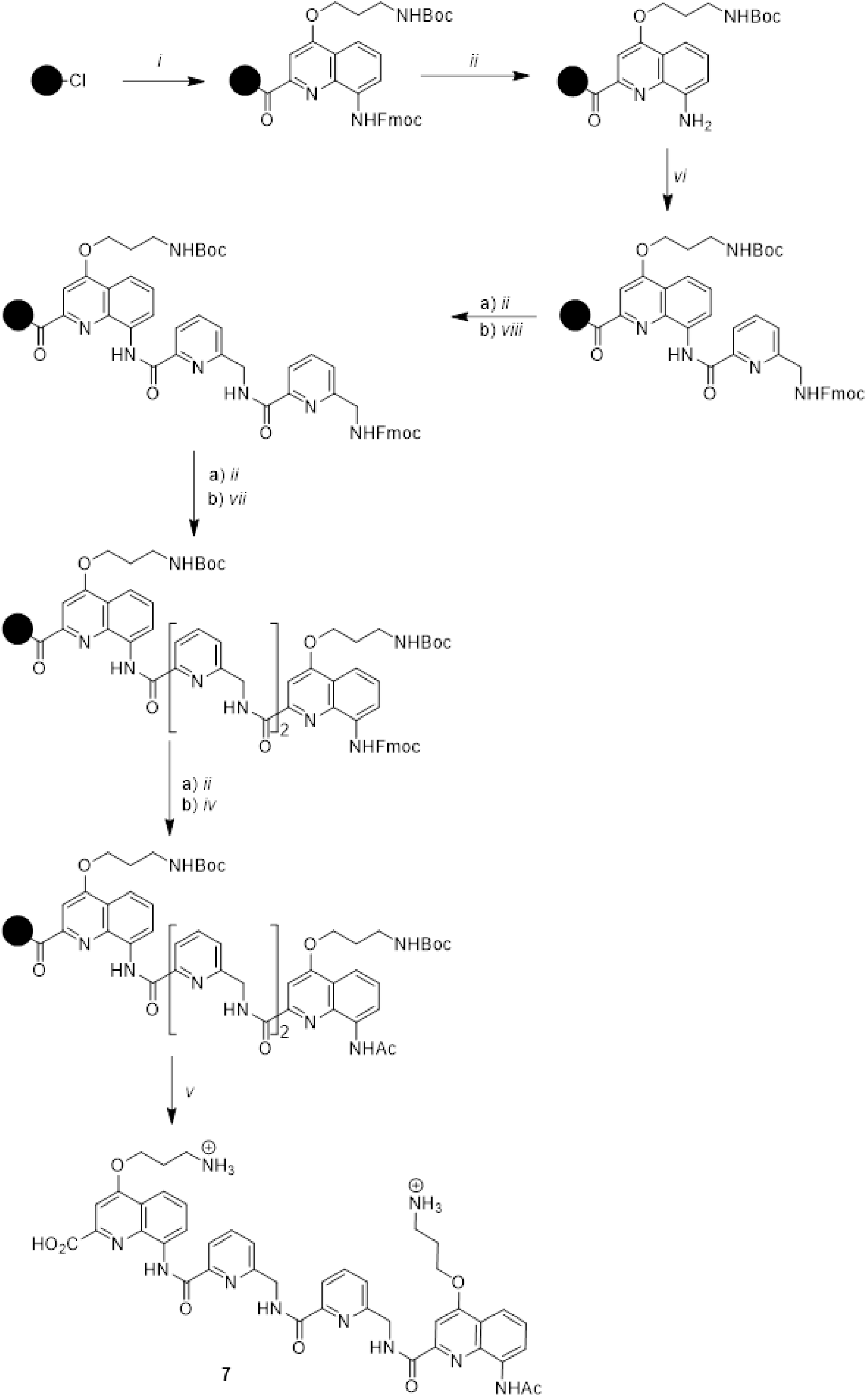
Solid phase synthesis of oligomer **7**. viii) 2 times, **P** monomer, HBTU, DIEA, dry DMF.

###### Characterization of Q3

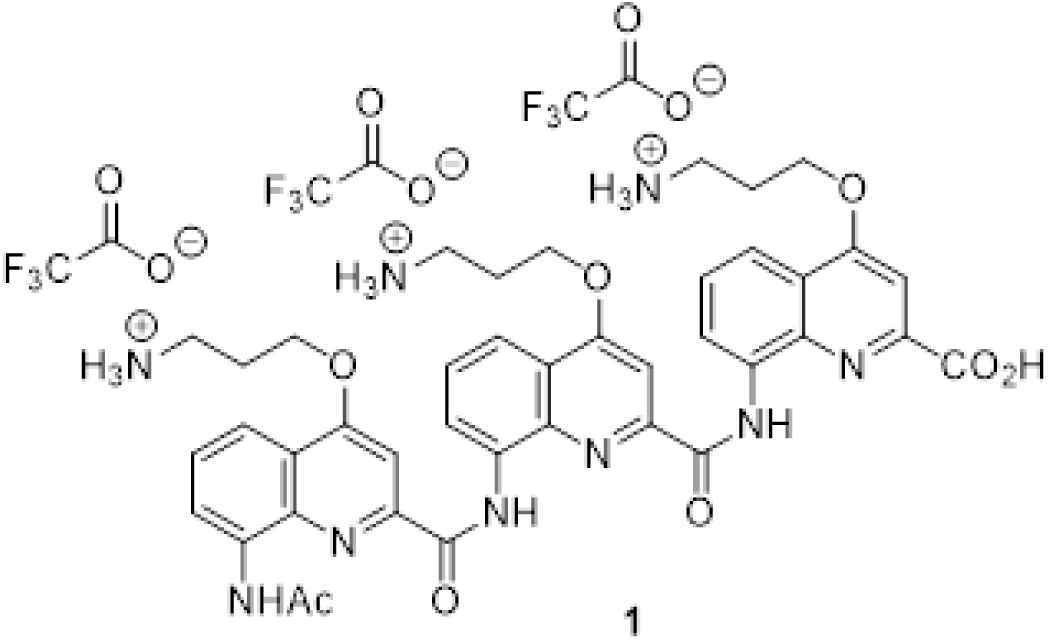

Trimer **1** was synthetized as a trifluoroacetate salt on 8 µmol scale following the *in-situ* activation procedure. The target compound was obtained as a light-yellow solid after purification by preparative HPLC (4 mg, 63% yield). ^1^H NMR (700 MHz, DMSO-d6). δ 12.45 (brs, 1H), 12.27 (s, 1H), 12.16 (s, 1H), 9.23 (s, 1H), 8.91 (d, 1H, ^3^JH-H=7.5 Hz), 8.85 (d, 1H, ^3^JH-H=7.4 Hz), 8.03 (d, 1H, ^3^JH-H=8.2 Hz), 7.76-7.87 (m, 9H), 7.71-7.74 (m, 2H), 7.66-7.70 (m, 2H), 7.28 (t, 1H, ^3^JH-H=8.0 Hz), 6.58 (s, 1H), 4.57 (t, 2H, ^3^JH-H=5.8 Hz), 4.54 (t, 2H, ^3^JH-H=5.5 Hz), 4.12 (t, 2H, ^3^JH-H=5.5 Hz), 3.04-3.16 (m, 6H), 2.21-2.27 (m, 4H), 2.15-2.20 (m, 2H), 1.68 (s, 2H). HRMS (ESI^+^) m/z [M+H]^+^ 790.3303 (calc. 790.3307 for C41H44O8N ^+^).

**Figure S7.**
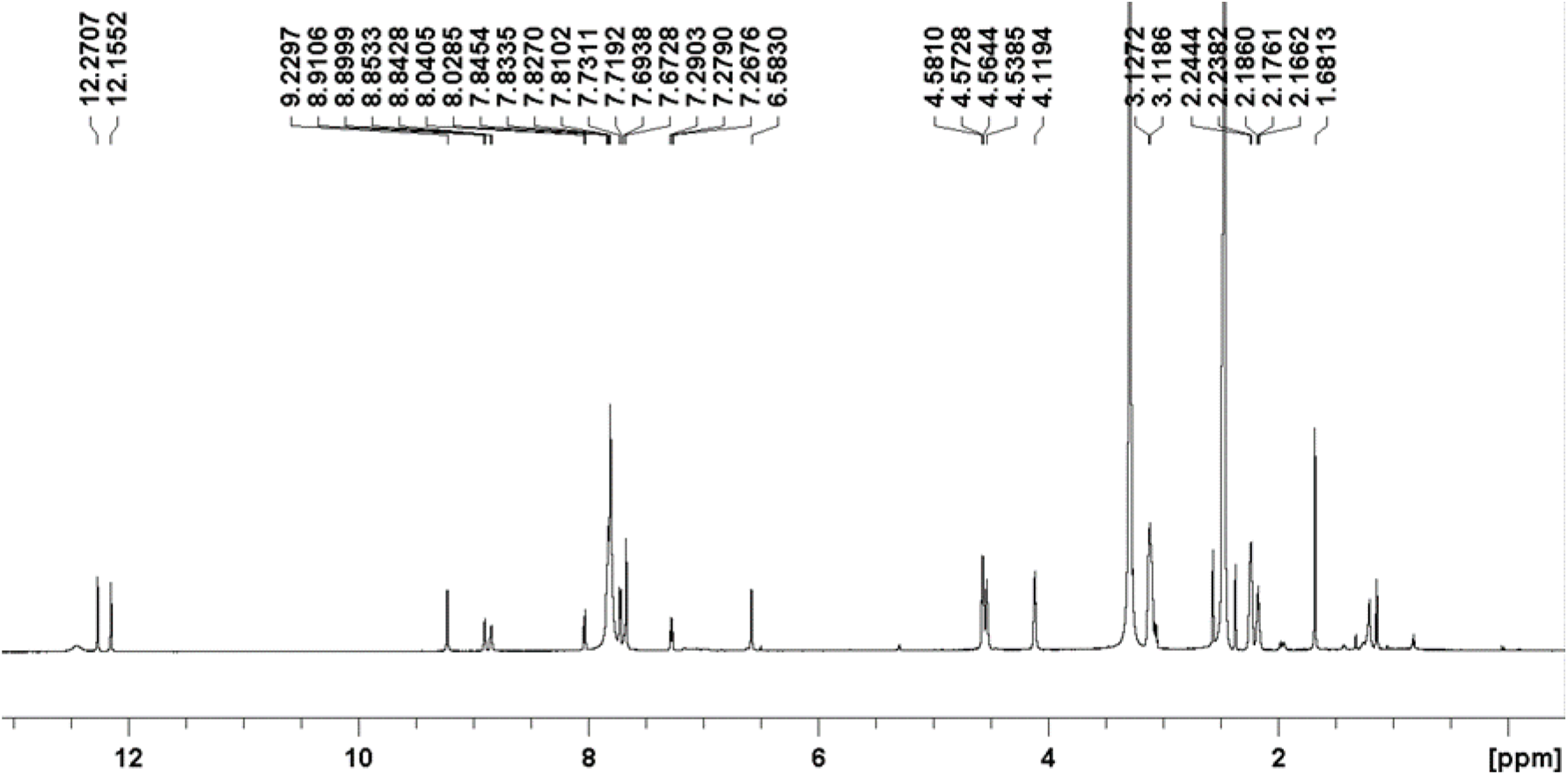
^1^H NMR spectrum of oligomer **1**, measured in DMSO-d6 at 25°C.

**Figure S8.**
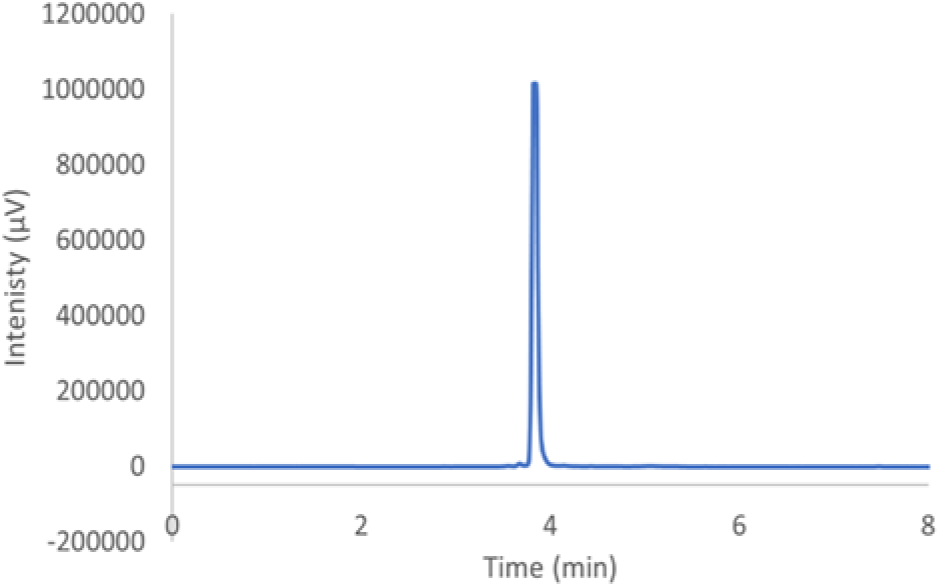
HPLC trace of oligomer **1**. Analysis was performed using solvents A and B with the gradient: 0 min: 100% A, 0% B –2 min: 100% A, 0% B – 12 min: 0% A, 100% B – 15min: 0% A, 100% B.

###### Characterization of Q4

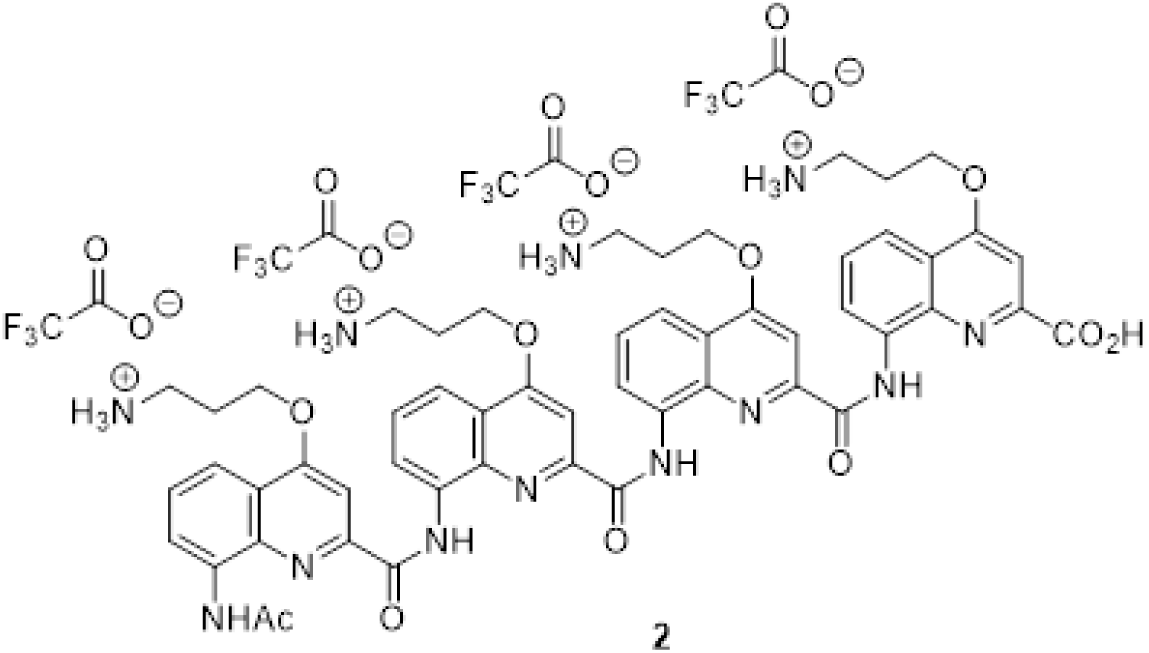

Tetramer **2** was synthetized as a trifluoroacetate salt on 16 µmol scale following the *in-situ* activation procedure. The target compound was obtained as a light-yellow solid after purification by preparative HPLC (16 mg, 67% yield). ^1^H NMR (700 MHz, DMSO-d6). δ 12.53 (brs, 1H), 12.20 (s, 1H), 11.77 (s, 1H), 11.69 (s, 1H), 9.08 (s, 1H), 8.97 (d, 1H, 3JH-H=7.3 Hz), 8.43 (brs, 1H), 7.81-7.99 (m, 12H), 7.75-7.80 (m, 2H), 7.74 (s, 1H), 7.44 (brs, 1H), 7.37 (t, 1H, 3JH-H=7.8 Hz), 7.26 (s, 1H), 6.75 (s, 1H), 6.66 (brs, 1H), 4.54-4.59 (m, 4H), 4.34 (brs, 1H), 4.23 (brs, 1H), 4.16 (brs, 2H), 3.07-3.22 (m, 6H), 2.20-2.46 (m, 8H), 1.69 (s, 3H). HRMS (ESI^+^) m/z 1033,4316 [M+H]^+^ (calc. 1033,4315 for C54H57O10N ^+^).

**Figure S9.**
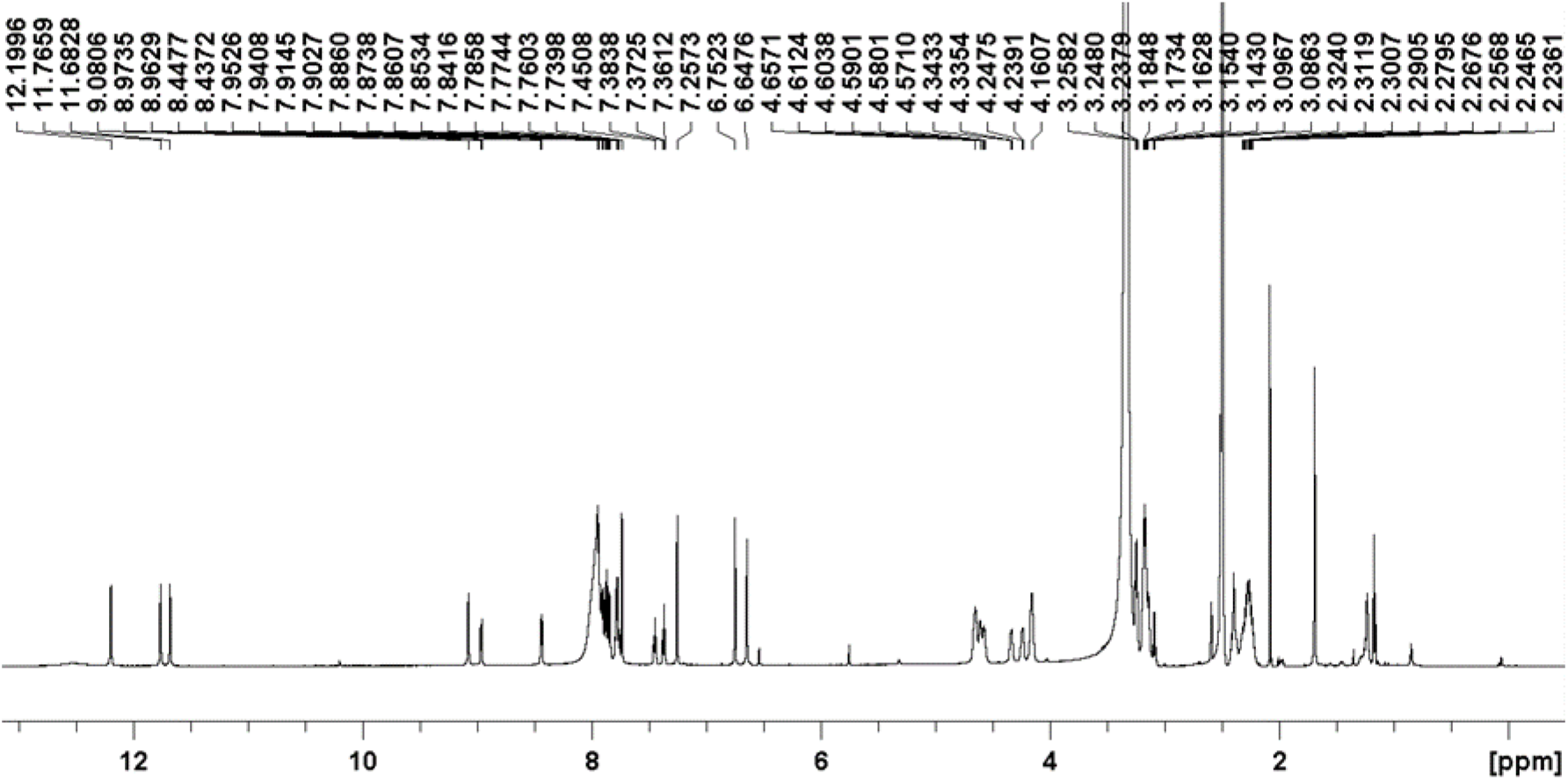
^1^H NMR spectrum of tetramer **2**, measured in DMSO-d6 at 25°C on 700MHz Bruker Avance NEO spectrometer (TXI probe (^1^H, ^13^C, ^15^N, ^2^H), 5mm, z-gradients).

**Figure S10.**
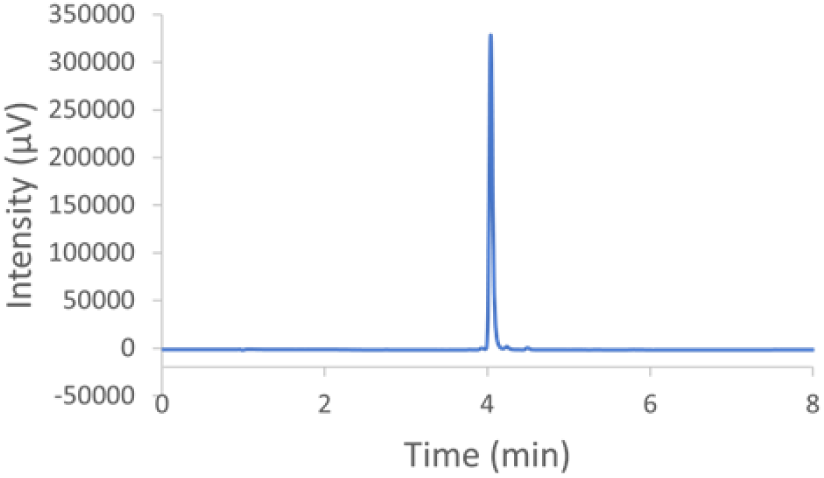
HPLC trace of oligomer **2**. Analysis was performed using solvents A and B with the gradient: 0 min: 100% A, 0% B – 2 min: 100% A, 0% B – 12 min: 0% A, 100% B – 15 min: 0% A, 100% B.

###### Characterization of Q5

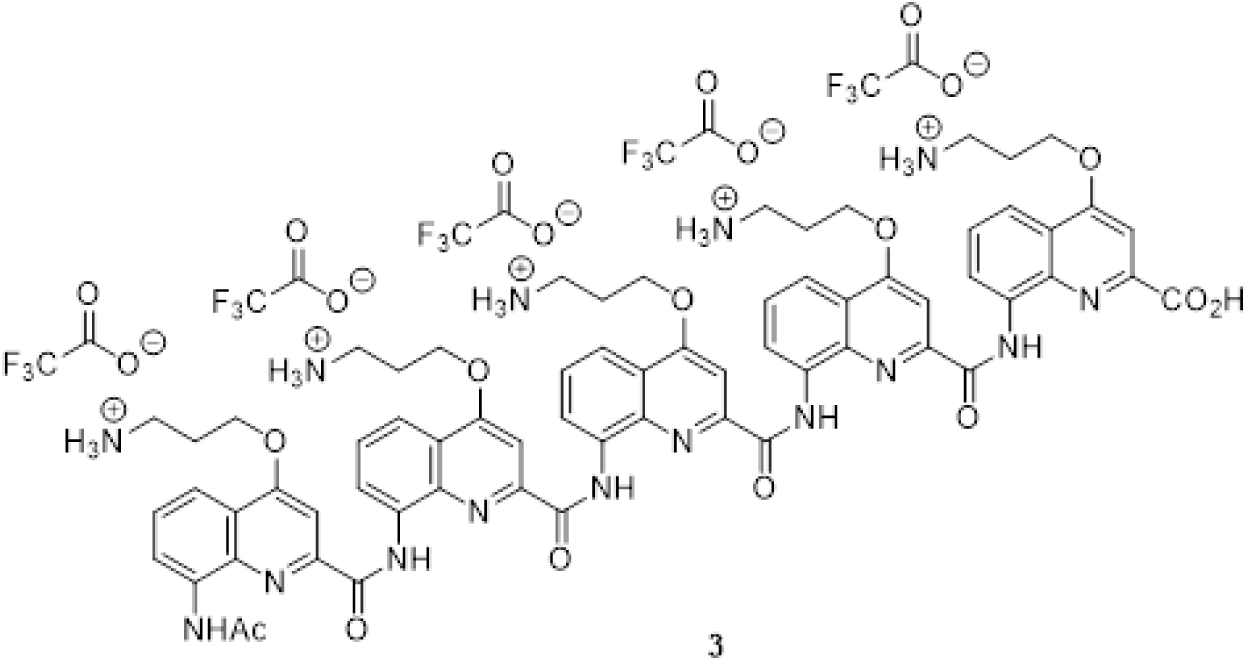

Pentamer **3** was synthetized as a trifluoroacetate salt on 10 µmol scale following the *in-situ* activation procedure. The target compound was obtained as a light-yellow solid after purification by preparative HPLC (7.5 mg, 59% yield). ^1^H NMR (700 MHz, DMSO-d6). δ 12.21 (brs, 1H), 11.74 (s, 1H), 11.68 (s, 1H), 11.64 (s, 1H), 11.52 (s, 1H), 8.87 (s, 1H), 8.50 (d, 2H, ^3^JH-H), 7.4 Hz), 7.88-8.06 (m, 17H), 7.78-7.86 (m, 3H), 7.75 (d, 1H, ^3^JH-H=8.2 Hz), 7.67 (d, 1H, ^3^JH-H=7.3 Hz), 7.51 (t, 1H, ^3^JH-H=7.8 Hz), 7.38 (t, 1H, ^3^JHH=7.8 Hz), 7.33 (t, 1H, ^3^JH-H=7.8 Hz), 7.30 (s, 1H), 7.20 (s, 1H), 6.79 (s, 1H), 6.67 (s, 1H), 6.52 (s, 1H), 4.67-4.73 (m, 2H), 3.08-3.28 (m, 13H), 2.17-2.46 (m, 13H), 1.36 (s, 2H). HRMS (ESI^+^) m/z [M+H]^+^ 1276.5369 (calc. 1276.5323 for C67H70O12N ^+^).

**Figure S11.**
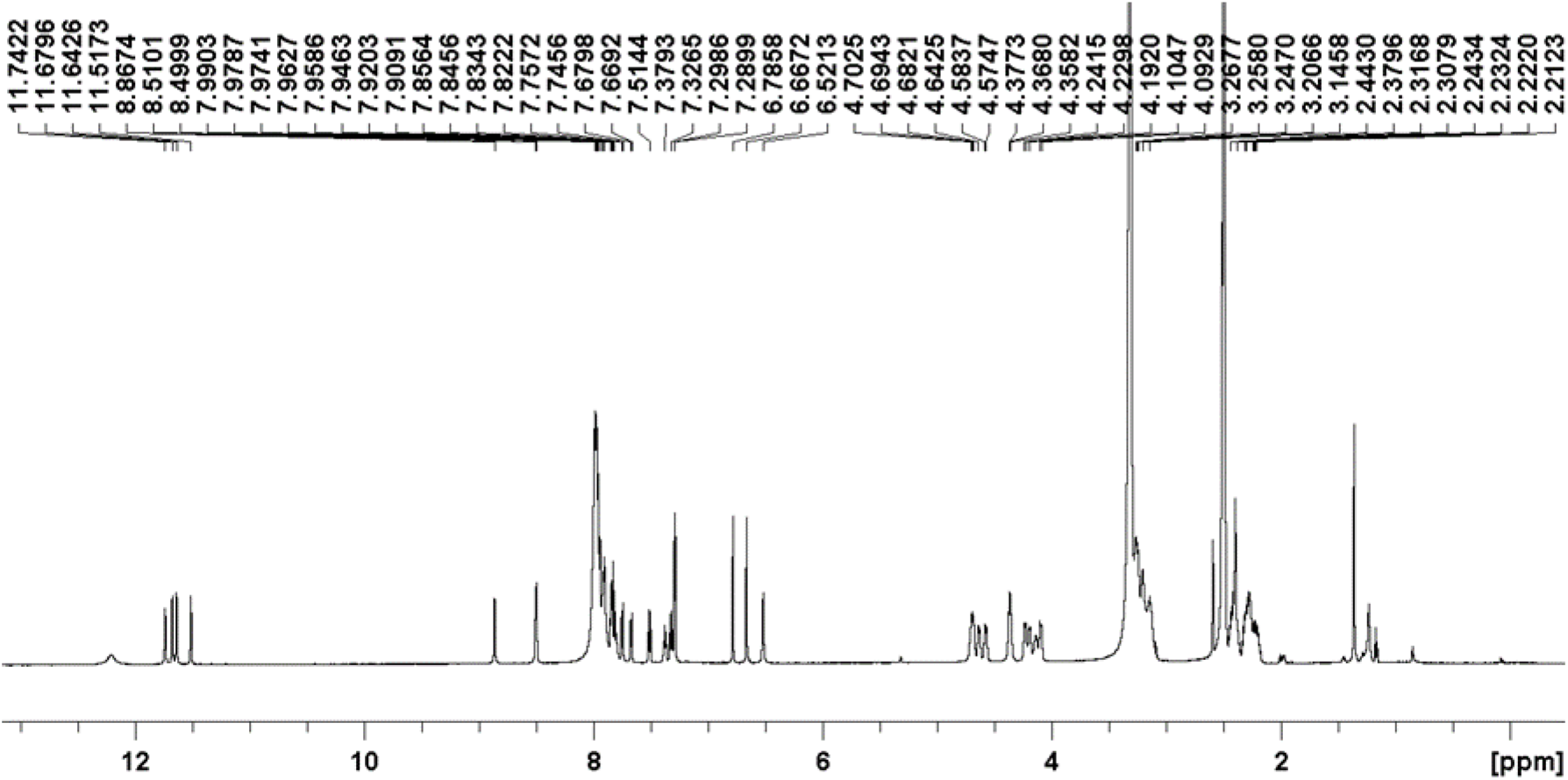
^1^H NMR spectrum of pentamer **3**, measured in DMSO-d6 at 25°C on 700MHz Bruker Avance NEO spectrometer (TXI probe (^1^H, ^13^C, ^15^N, ^2^H), 5mm, z-gradients).

**Figure S12.**
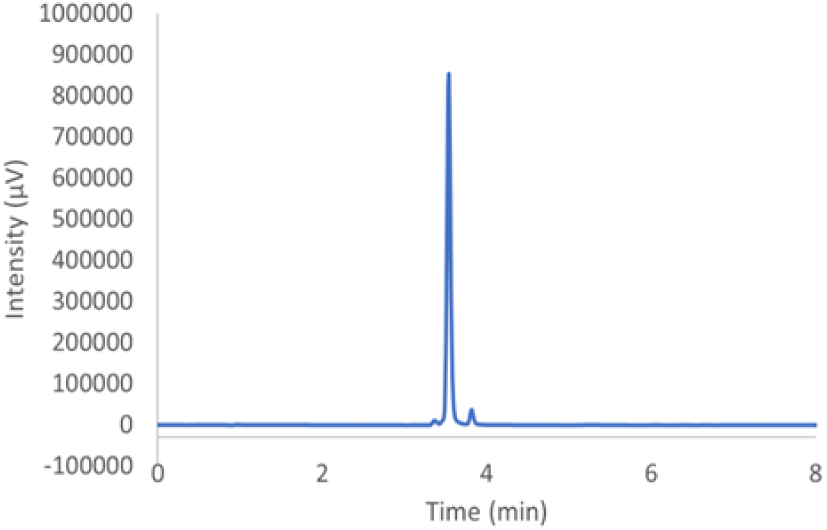
HPLC trace of oligomer **3**. Analysis was performed using solvents A and B with the gradient: 0 min: 100% A, 0% B – 2 min: 100% A, 0% B – 12 min: 0% A, 100% B – 15 min: 0% A, 100% B.

###### Characterization of Q8

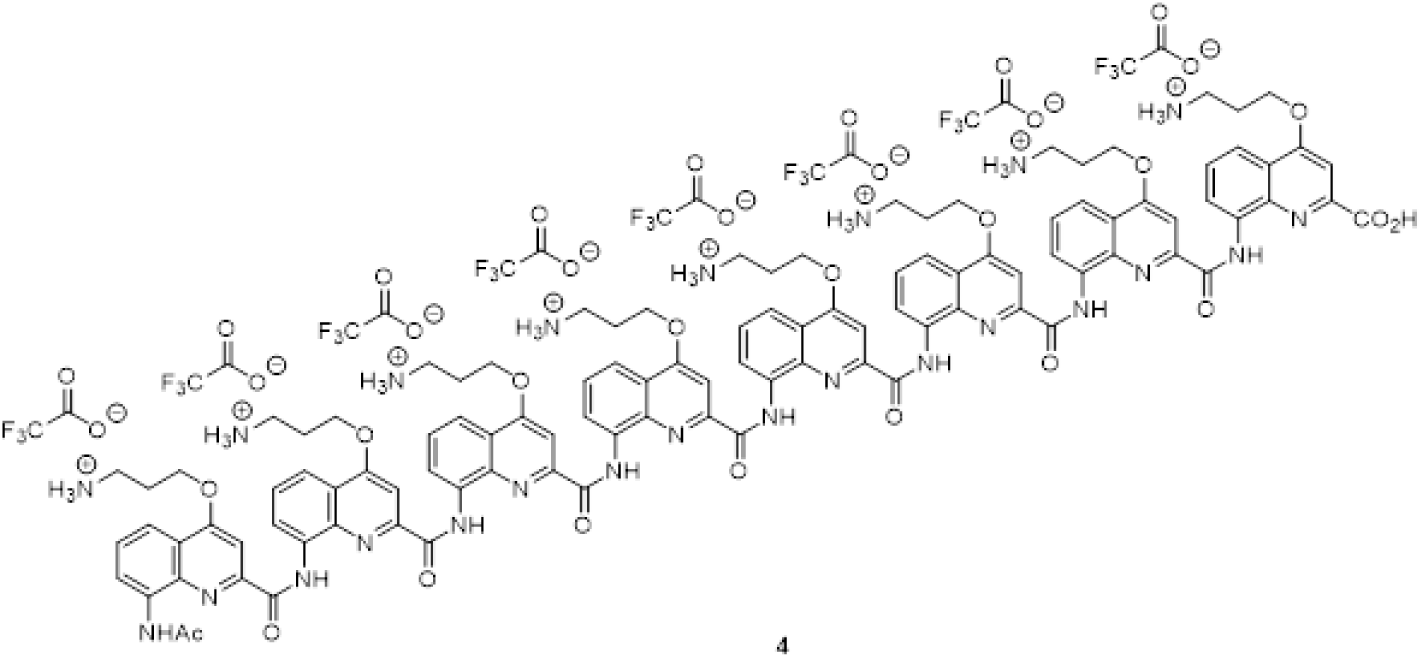

Octamer **4** was synthetized as a trifluoroacetate salt on 10 µmol scale following the *in-situ* activation procedure. The target compound was obtained as a light-yellow solid after purification by preparative HPLC (11.5 mg, 57% yield). ^1^H NMR (700 MHz, DMSO-d6). δ 11.93 (brs, 1H), 11.28 (s, 1H), 11.25 (s, 1H), 11.14 (s, 1H), 11.04 (s, 1H), 11.03 (s, 1H), 10.94 (s, 1H), 10.88 (s, 1H), 8.91 (brs, 1H), 8.44 (s, 1H), 8.07-8.27 (m, 11H), 7.83-8.04 (m, 12H), 7.74-7.81 (m, 3H), 7.67 (d, 2H, ^3^JH-H=7.9 Hz), 7.44-7.51 (m, 3H), 7.31-7.43 (m, 4H), 7.17 (t, 1H, ^3^JH-H=8.3 Hz), 7.12 (t, 1H, ^3^JH-H=8.1 Hz), 7.02 (d, 1H, ^3^JH-H=7.3Hz), 6.96 (s, 1H), 6.85 (s, 1H), 6.58 (s, 1H), 6.53 (s, 1H), 6.36 (s, 1H), 6.30 (s, 1H), 6.17 (s, 1H), 5.91 (s, 1H), 4.36-4.51 (m, 5H), 4.12-4.34 (m, 9H), 3.97-4.07 (m, 3H), 3.85-3.90 (m, 1H), 2.96-3.19 (m, 16H), 1.95-2.35 (m, 15H), 1.24 (s, 3H). HRMS (ESI^+^) m/z [M+H]^+^ 2005.8351 (calc. 2005.8346 for C106H109O18N24

**Figure S13.**
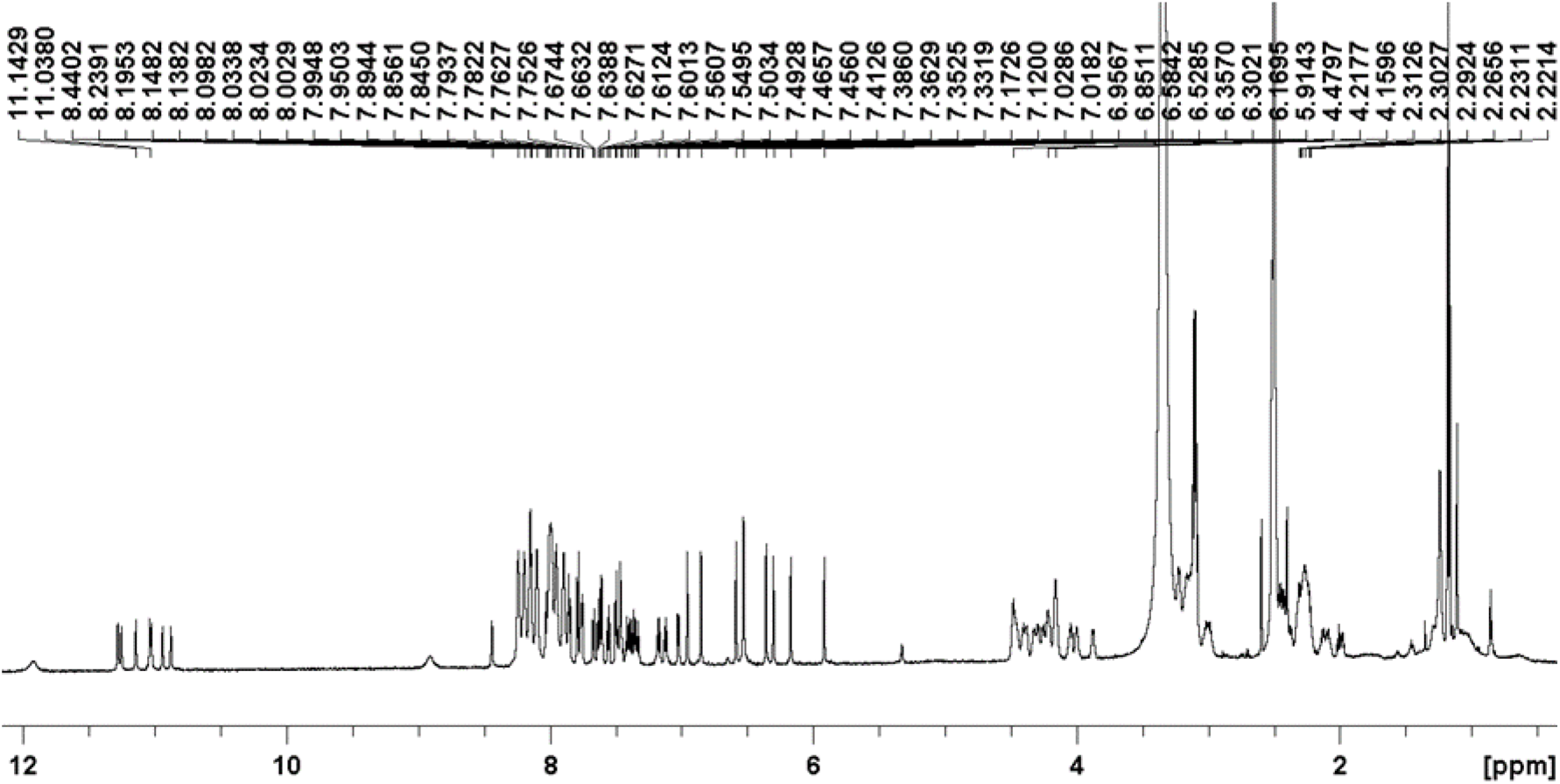
^1^H NMR spectrum of octamer **4**, measured in DMSO-d6 at 25°C on 700MHz Bruker Avance NEO spectrometer (TXI probe (^1^H, ^13^C, ^15^N, ^2^H), 5mm, z-gradients).

**Figure S14.**
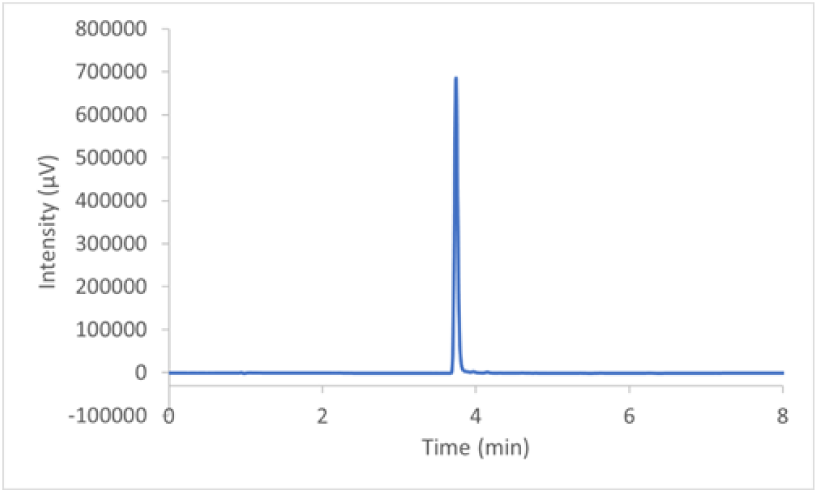
HPLC trace of oligomer **4**. Analysis was performed using solvents A and B with the gradient: 0 min: 100% A, 0% B – 2 min: 100% A, 0% B – 12 min: 0% A, 100% B – 15 min: 0% A, 100% B. Figure S15 ^1^H NMR spectrum of trimer **5**, measured in DMSO-d6 at 25°C on 700MHz Bruker Avance NEO spectrometer (TXI probe (^1^H, ^13^C, ^15^N, ^2^H), 5mm, z-gradients).

###### Characterization of QPQ

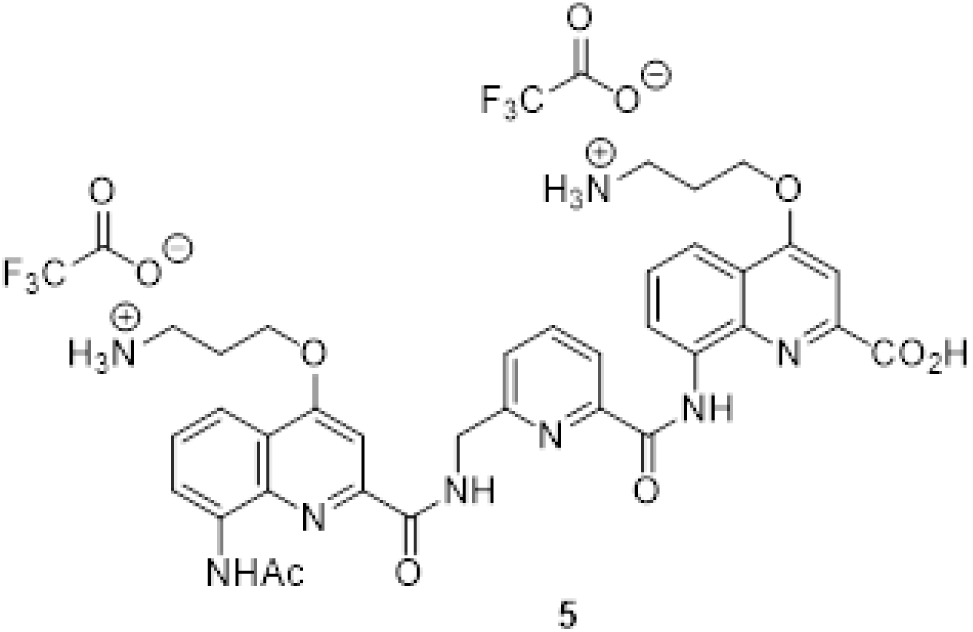

Trimer **5** was synthetized as a trifluoroacetate salt on 16 µmol scale following *in-situ* activation procedure for coupling on **Q^Orn^** monomer and HBTU coupling procedure for coupling on **P** monomer. The target compound was obtained as a white solid after purification by preparative HPLC (7mg, 48% yield). ^1^H NMR (700 MHz, DMSO-d6). δ 13.61 (brs, 1H), 12.60 (s, 1H), 10.35 (t, 1H, 3JH-H=6.2 Hz), 10.25 (s, 1H), 8.91 (d, 1H, 3JH-H=7.2 Hz), 8.74 (d, 1H, 3JH-H=7.7 Hz), 8.08-8.17 (m, 2H), 7.69-7.97 (m, 8H), 7.60-7.67 (m, 3H), 6.55 (s, 1H), 4.99 (d, 2H, 3JH-H=6.1 Hz), 4.43-4.50 (m, 4H), 3.06-3.15 (m, 5H), 2.29 (s, 2H), 2.16-2.25 (m, 4H), 1.24 (s, 3H). HRMS (ESI^+^) m/z [M+H]^+^ 681.2777 (calc. 681.2780 for C35H37O7N ^+^).

**Figure S15.**
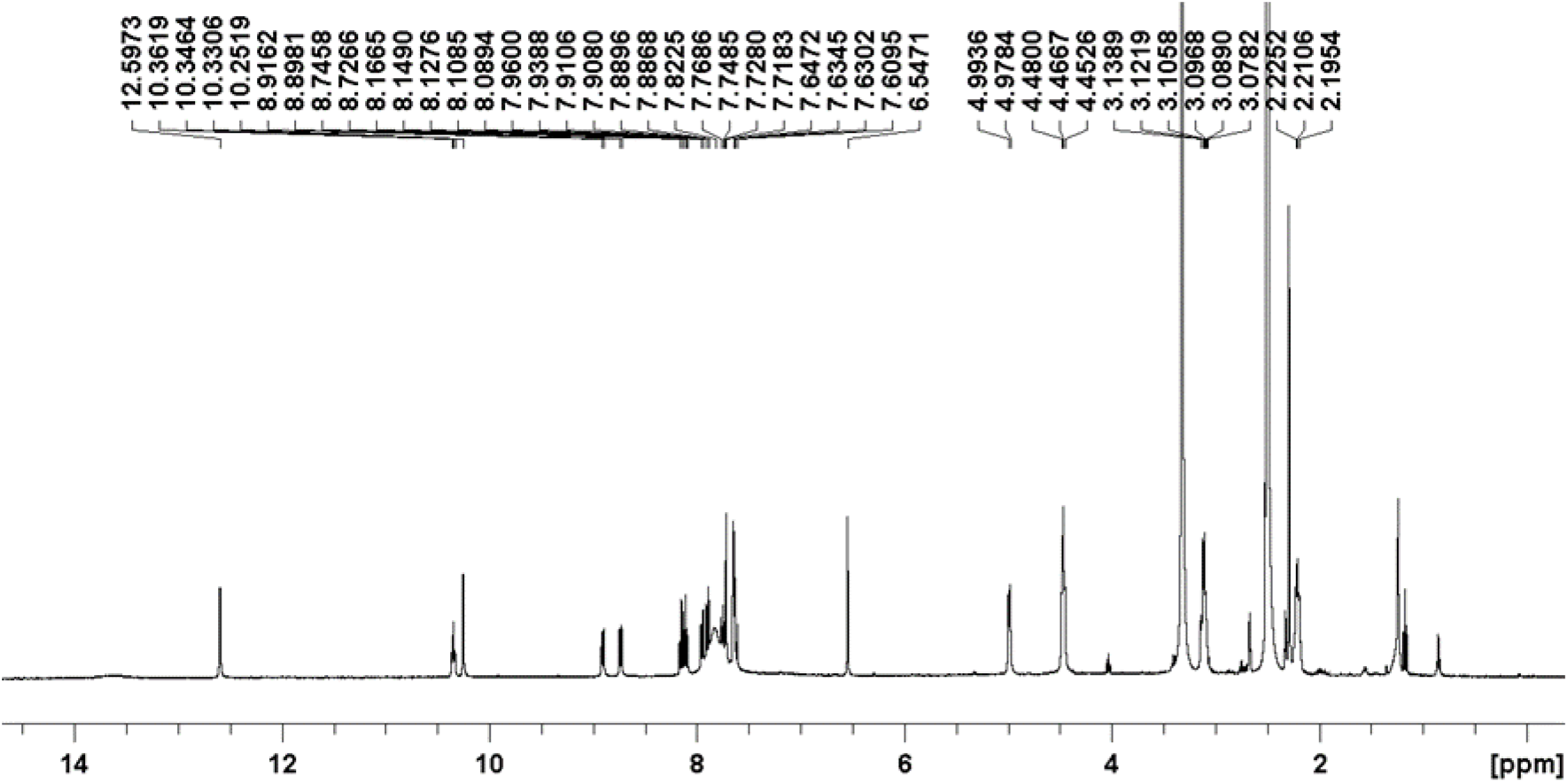
^1^H NMR spectrum of trimer **5**, measured in DMSO-d6 at 25°C on 700MHz Bruker Avance NEO spectrometer (TXI probe (^1^H, ^13^C, ^15^N, ^2^H), 5mm, z-gradients).

**Figure S16.**
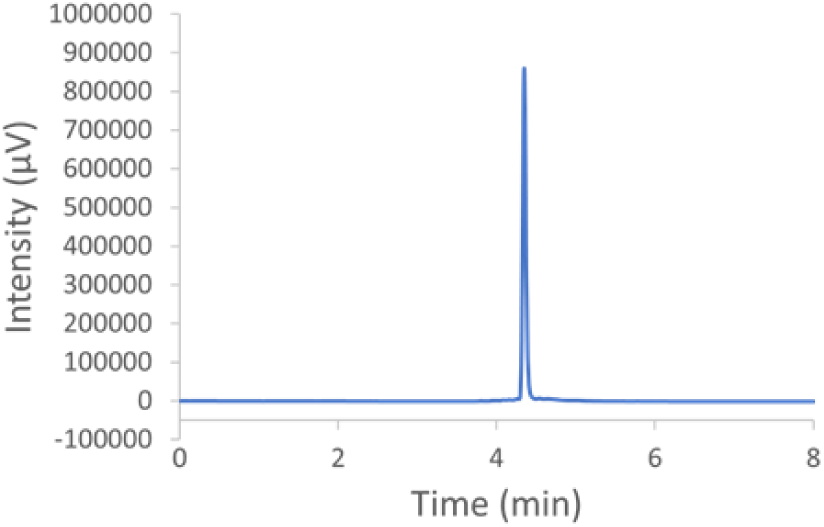
HPLC trace of oligomer **5**. Analysis was performed using solvents A and B with the gradient: 0 min: 100% A, 0% B – 2 min: 100% A, 0% B – 12 min: 0% A, 100% B – 15 min: 0% A, 100% B.

###### Characterization of QQPQ

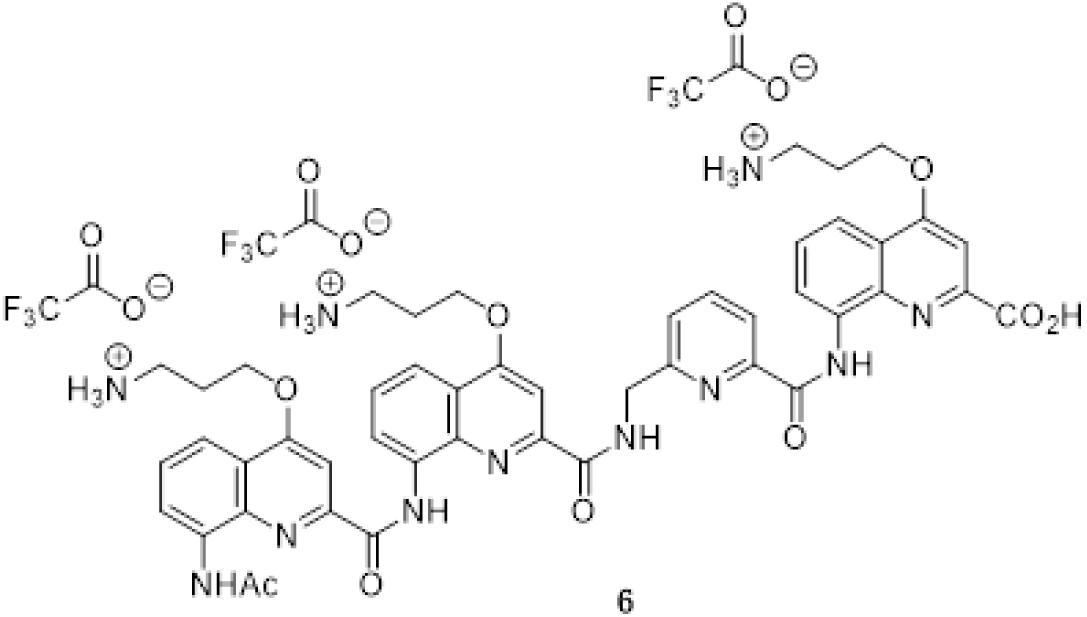

Tetramer **6** was synthetized as a trifluoroacetate salt on 16 µmol scale following *in-situ* activation procedure for coupling on **Q^Orn^** monomer and HBTU coupling procedure for coupling on **P** monomer. The target compound was obtained as a white solid after purification by preparative HPLC (13 mg, 64% yield). ^1^H NMR (700 MHz, DMSO-d6). δ 11.82 (brs, 1H), 11.56 (s, 1H), 9.52 (brs, 1H), 9.30 (s, 1H), 8.31 (d, 1H, ^3^JH-H=7.4 Hz), 7.77-8.14 (m, 10H), 7.61-7.71 (m, 2H), 7.75 (t, 1H, ^3^JH-H=7.9 Hz), 7.43 (t, 1H, 3JH-H=8.0 Hz), 7.31 (d, 1H, ^3^JH-H=7.4 Hz), 7.11 (s, 1H), 6.74 (s, 1H), 6.57 (s, 1H), 5.72 (s, 1H), 4.78 (brs, 2H), 4.53-4.56 (m, 2H), 4.48-4.52 (m, 2H), 4.18-4.22 (m, 2H), 3.06-3.25 (m, 7H), 2.16-2.38 (m, 5H). HRMS (ESI^+^) m/z [M+H]^+^ 924.3048 (calc. 924.3787 for C48H50O9N ^+^).

**Figure S17.**
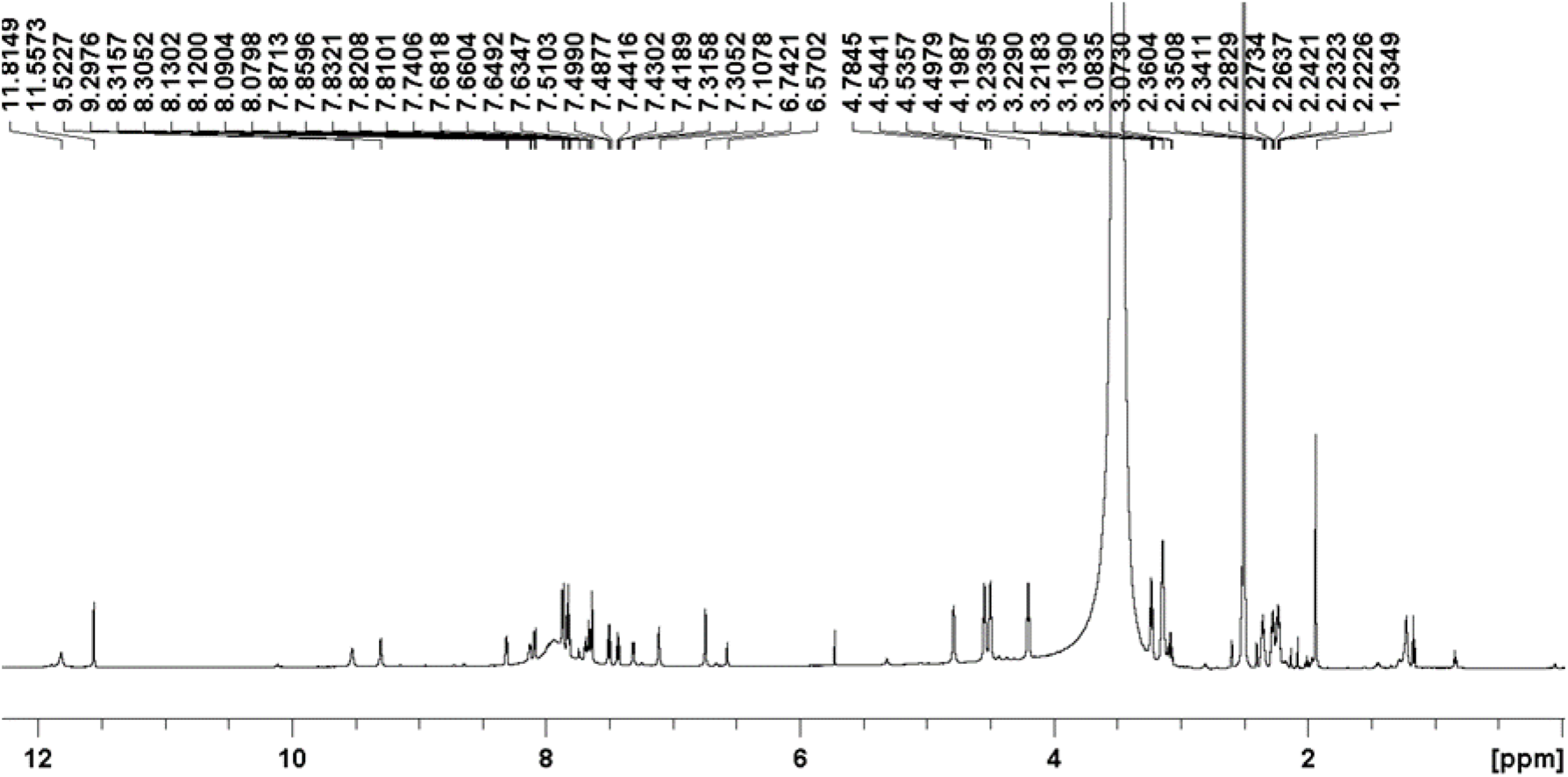
^1^H NMR spectrum of tetramer **6**, measured in DMSO-d6 at 25°C on 700MHz Bruker Avance NEO spectrometer (TXI probe (^1^H, ^13^C, ^15^N, ^2^H), 5mm, z-gradients).

**Figure S18.**
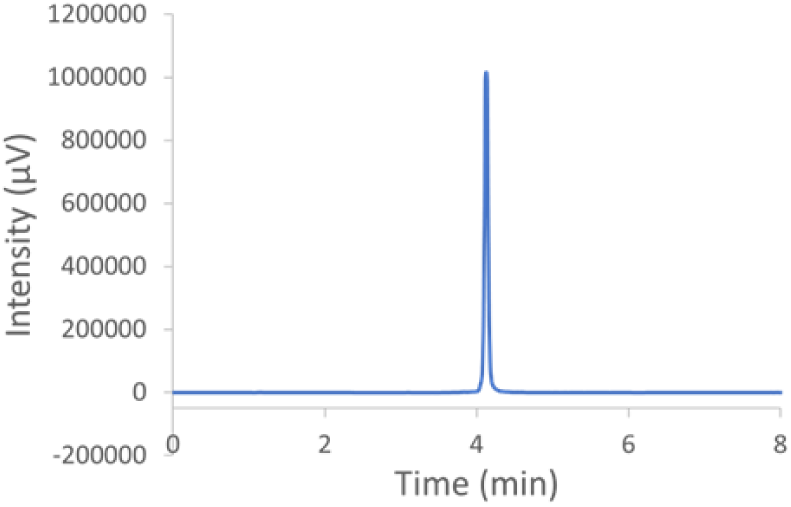
HPLC trace of oligomer **6**. Analysis was performed using solvents A and B with the gradient: 0 min: 100% A, 0% B – 2 min: 100% A, 0% B – 12 min: 0% A, 100% B – 15 min: 0% A, 100% B.

###### Characterization of QPPQ

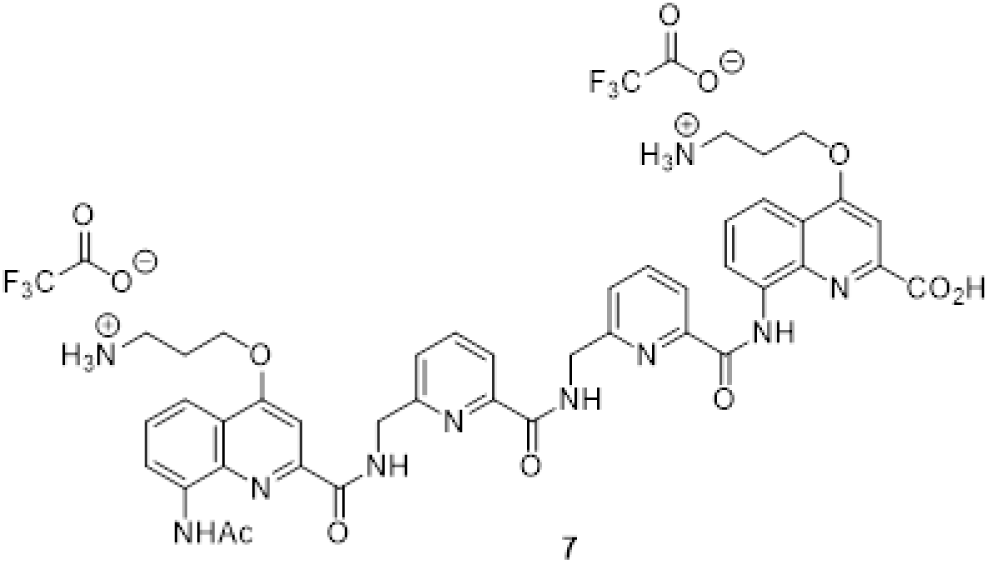

Tetramer **7** was synthetized as a trifluoroacetate salt on 16 µmol scale following *in-situ* activation procedure for coupling on **Q^Orn^** monomer and HBTU coupling procedure for coupling on **P** monomer. The target compound was obtained as a white solid after purification by preparative HPLC (11 mg, 66% yield). ^1^H NMR (300 MHz, DMSO-d6). δ 12.60 (s, 1H), 10.09 (brs, 1H), 9.61 (brs, 1H), 8.91 (d, 1H, ^3^JH-H=7.7 Hz),7.92-8.15 (m, 7H), 7.64-7.82 (m, 9H), 7.48-7.60 (m, 4H), 7.21-7.36 (m, 4H), 6.84 (d, 1H, ^3^JH-H=7.2 Hz), 4.87 (d, 2H, ^3^JH-H=6.2 Hz), 4.73 (d, 2H, ^3^JH-H=6.1 Hz), 4.43-4.52 (m, 2H), 4.34-4.42 (m, 3H), 2.96-3.15 (m, 6H), 1.50-2.69 (m, 8H). HRMS (ESI^+^) m/z [M+H]^+^ 815.3253 (calc. 815.3260 for C42H43O8N ^+^).

**Figure S19.**
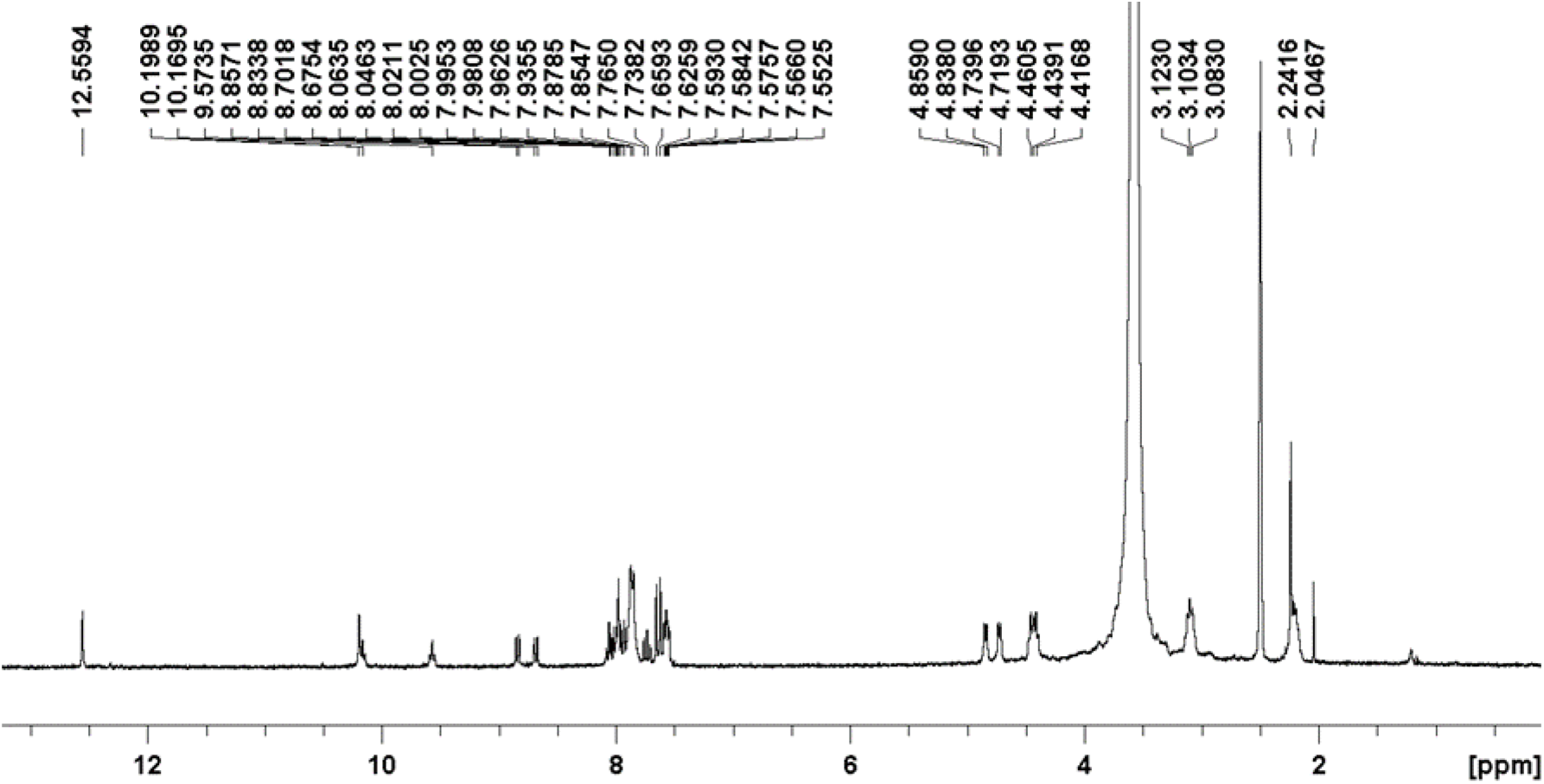
^1^H NMR spectrum of tetramer **7**, measured in DMSO-d6 at 25°C on 300MHz Bruker Avance NEO spectrometer (TXI probe (^1^H, ^13^C, ^15^N, ^2^H), 5mm, z-gradients).

**Figure S20.**
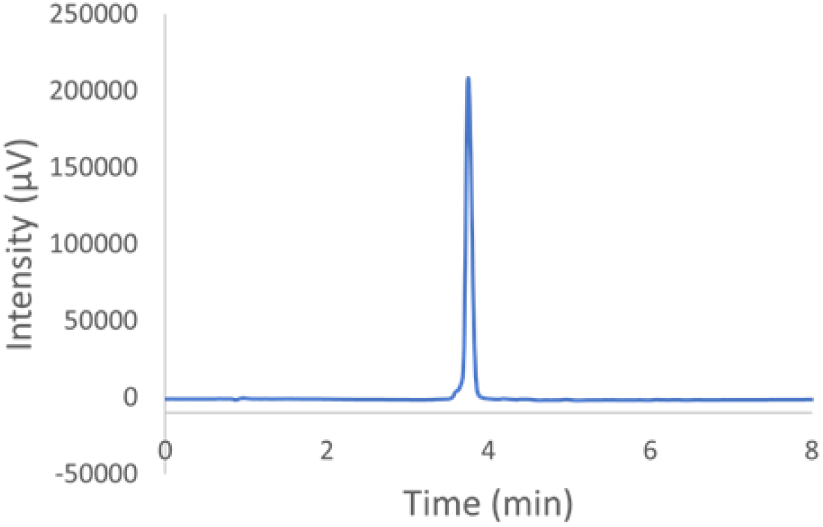
HPLC trace of oligomer **7**. Analysis was performed using solvents A and B with the gradient: 0 min: 100% A, 0% B – 2 min: 100% A, 0% B – 12 min: 0% A, 100% B – 15 min: 0% A, 100% B.

### Desalting procedure

Traces of Na^+^ in the oligonucleotide stock give rise to visible Na^+^ adducts in the mass spectrum, causing loss of resolution and S/N ratio. We therefore strip our stock solutions off of Na^+^ using centrifugal filters (Amicon Ultracel 3K, Millipore). The filters contain a cellulose matrix that will hold back the DNA macromolecule, while allowing the washing solution to pass through via centrifugation.

Prior to desalting, the oligonucleotides are dissolved in water and annealed at 85°C for 2-3 min. Then, the first segment of the desalting process is exchanging the non-volatile Na^+^ ion with volatile NH_4_ ^+^. The filter unit is filled with a washing solution of 500 mM Ammonium acetate and placed in the centrifuge for 15 minutes at 15,000 rpm. This process is repeated four times. The second segment is diluting with pure water to gradually flush out the NH_4_ ^+^ ions. Some NH_4_ ^+^ ions will remain electrostatically bound to the DNA strand, but they will detach during the MS ionization process, given their volatility. This washing process is repeated six times. The third segment is the recovery of the desalted DNA stock solution by placing the filter upside-down in a fresh tube and pushing it out at 1,000 rpm for 3 minutes. A proof of concept for the desalting method is illustrated in Figure S21.

**Figure S21.**
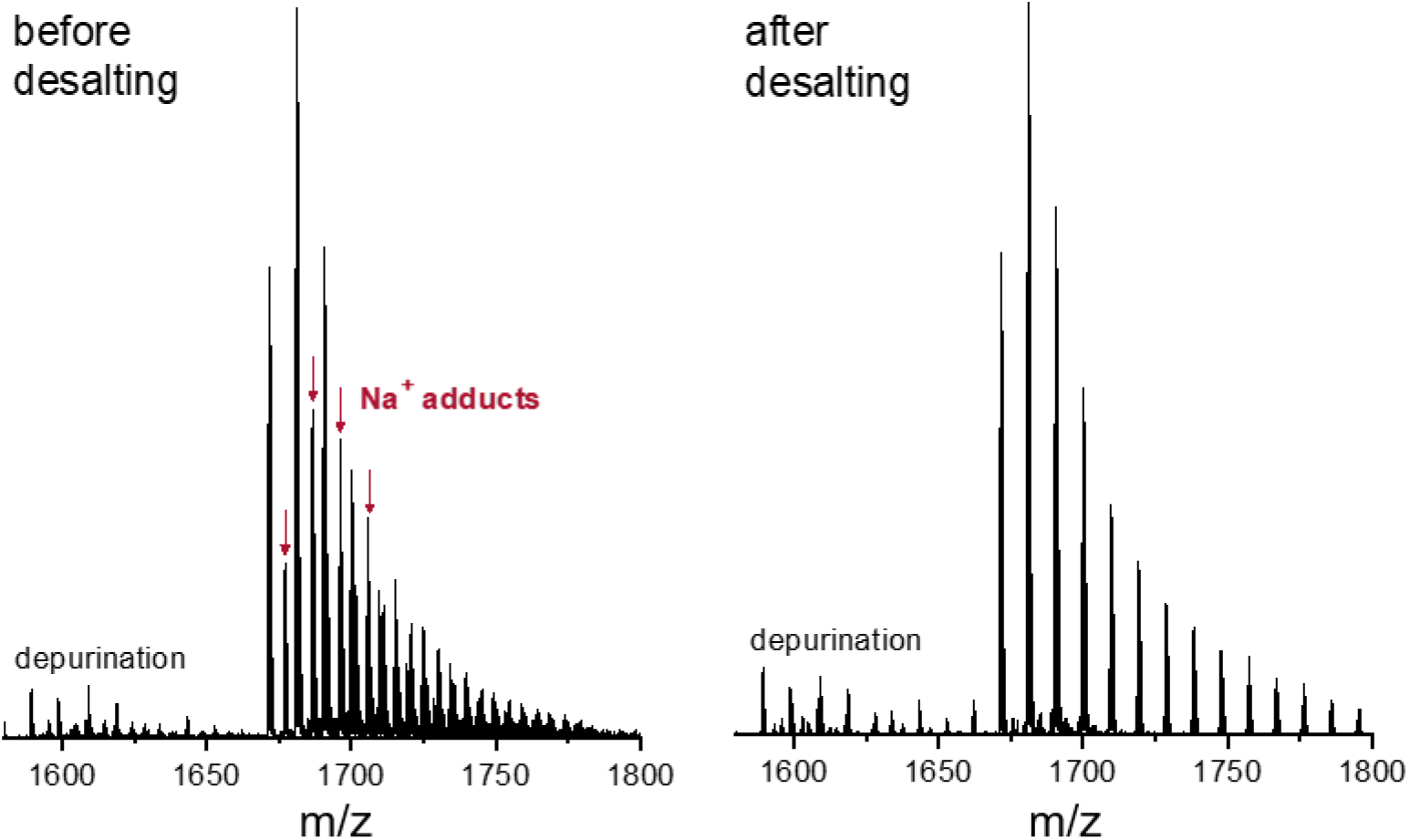
Mass spectrum of DNA sequence 21G (d**G_3_**TTA**G_3_**TTA**G_3_**TTA**G_3_**) before (left) and after (right) desalting, zoomed in on the 4-charge state. Sample conditions: 10 µM DNA, 1 mM KCl, 100 mM TMAA in H_2_O.

### CD/UV-melting curves of mutated G-quadruplex sequences

222T_mA, 222T-mC, 26CEB-mT, 22GT-18T and 24TTG-20T were introduced into the foldamer screening panel to probe the foldamers’ sensitivity to loop-induced G4 topology switches. 22GT-18T and 24TTG-20T are telomeric repeats where the third loop has a TTA to TTT mutation. Removing the adenine disrupts the formation of AGA triads and induces a topology switch.^1^ 222T_mA, 222T-mC and 26CEB-mT are mutated in the middle loop. The effect of mutating the middle loop is unknown for these sequences. We also provide spectra for ‘bcl2’ since we were unsure of its G-quadruplex stability in low K^+^ concentration.

For our six undocumented sequences we provide CD spectra and UV melting curves. One dataset in 1 mM KCl, to be orthogonal with our previously established database.^2^ Another dataset in 0.5 mM KCl, which is the concentration for the ligand screening. In second approach, this dataset gives insight for how much cutting the K^+^ concentration in half affects G-quadruplex stability.

**Figure S22.**
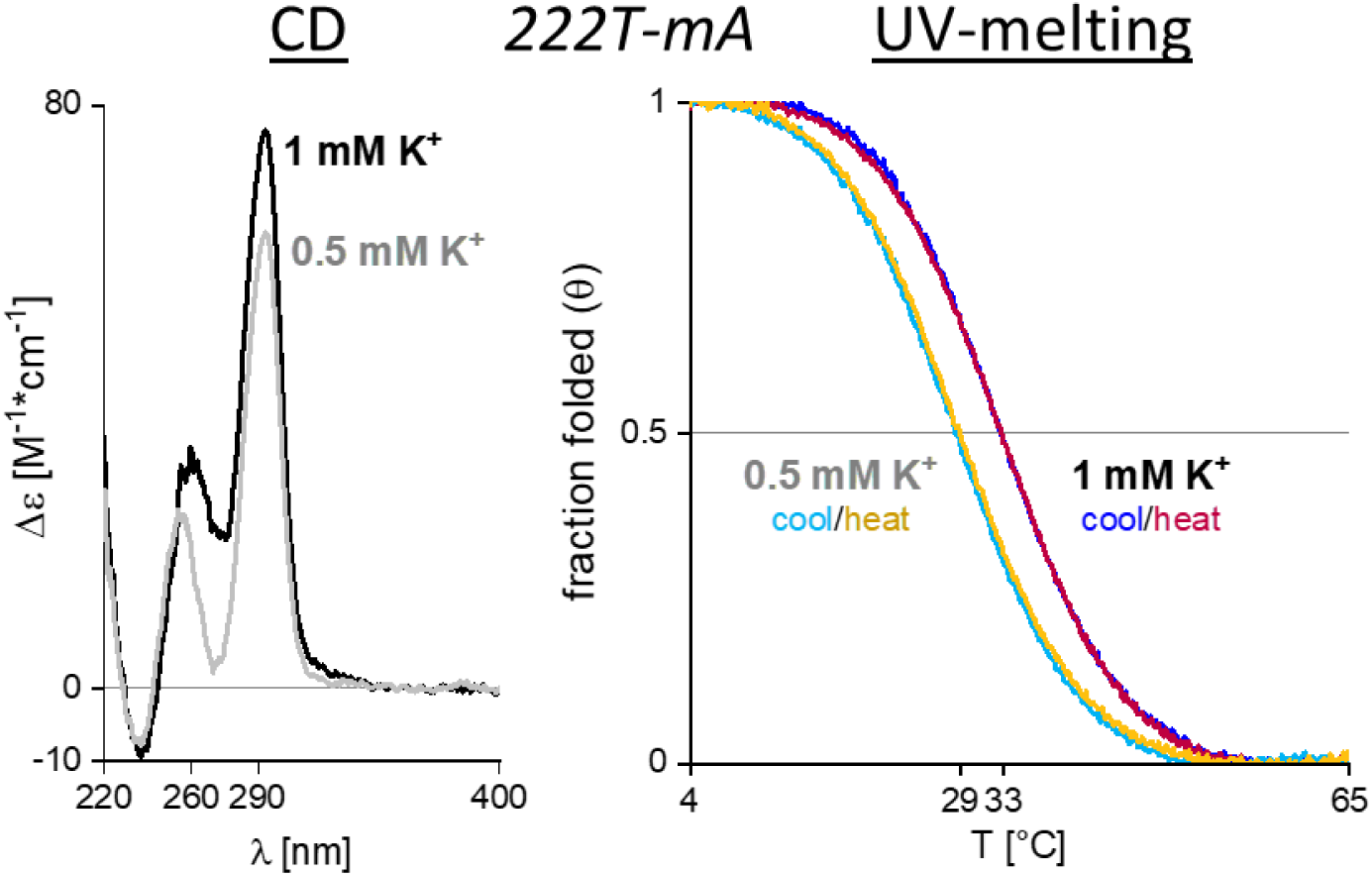
**Left**: CD spectra of 222T_mA (dT**G_3_**TT**G_3_**<u>AA</u>**G_3_**TT**G_3_**T) in 1 mM (black) and 0.5 mM KCl (grey). **Right**: UV-melting curves in 1 mM (blue/red) and 0.5 mM (cyan/orange) KCl. Samples contain 10 µM DNA and 100 mM TMAA (pH 6.8).

**Figure S23.**
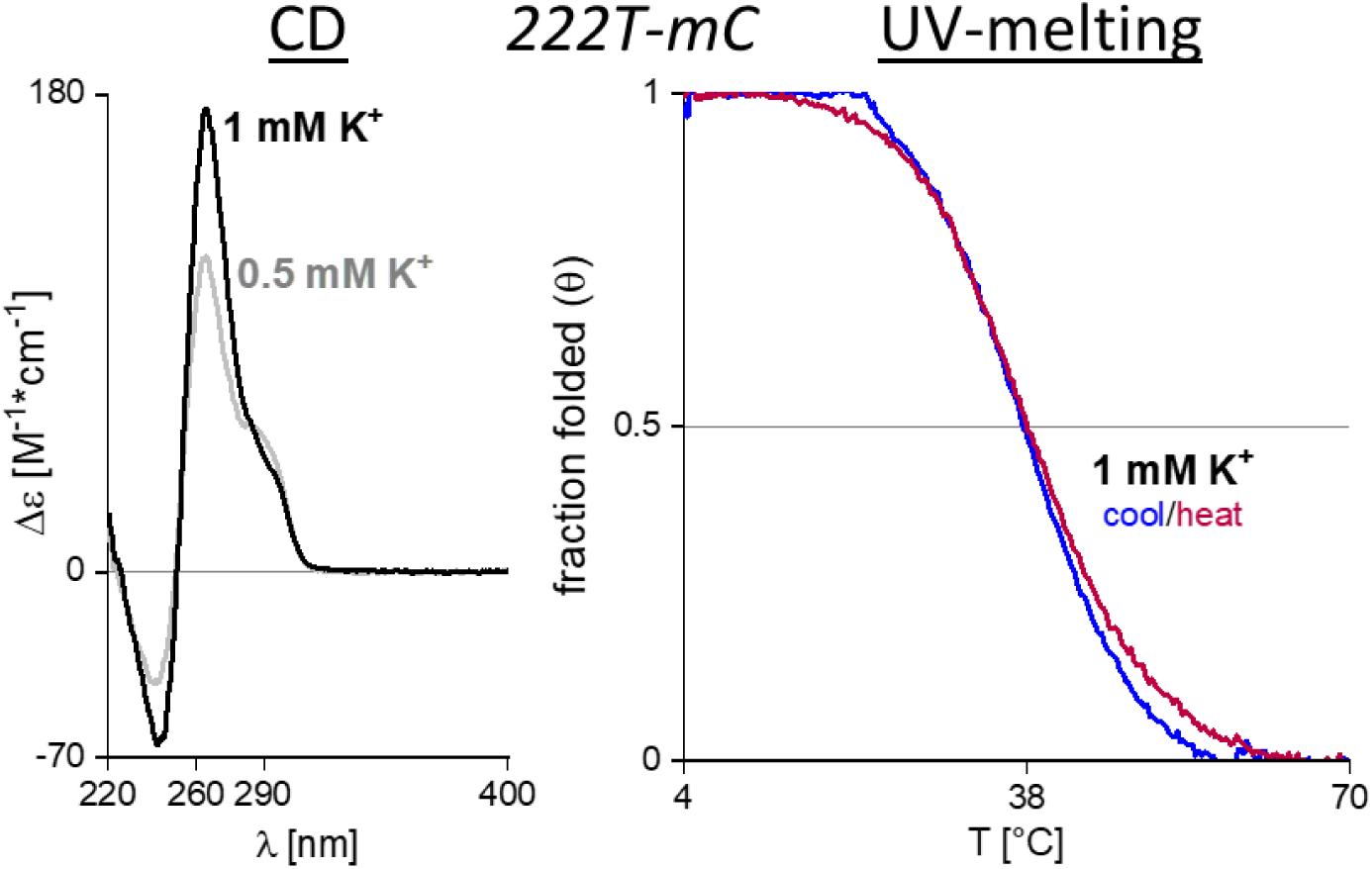
**Left**: CD spectra of 222T-mC (dT**G_3_**TT**G_3_**<u>CC</u>**G_3_**TT**G_3_**T) in 1 mM (black) and 0.5 mM KCl (grey). **Right**: UV-melting curve in 1 mM (blue/red) KCl. No data could be obtained in 0.5 mM KCl. Samples contain 10 µM DNA and 100 mM TMAA (pH 6.8).

**Figure S24.**
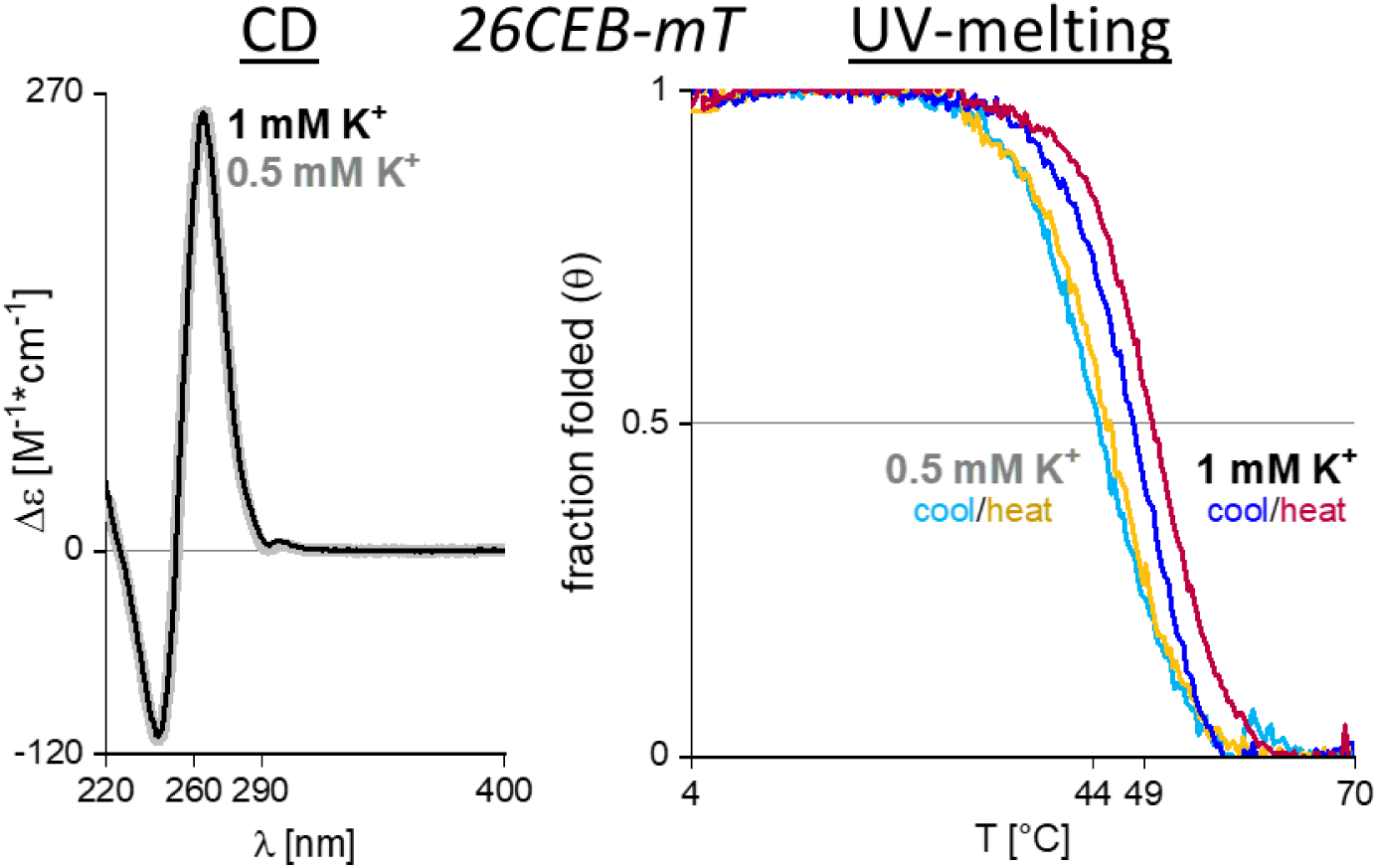
CD spectra of 26CEB-mT (dAA**G_3_**T**G_3_**<u>TTTTTTT</u>**G**T**G_3_**T**G_3_**T) in 1 mM (black) and 0.5 mM KCl (grey). **Right**: UV-melting curve in 1 mM (blue/red) KCl. Samples contain 10 µM DNA and 100 mM TMAA (pH 6.8).

**Figure S25.**
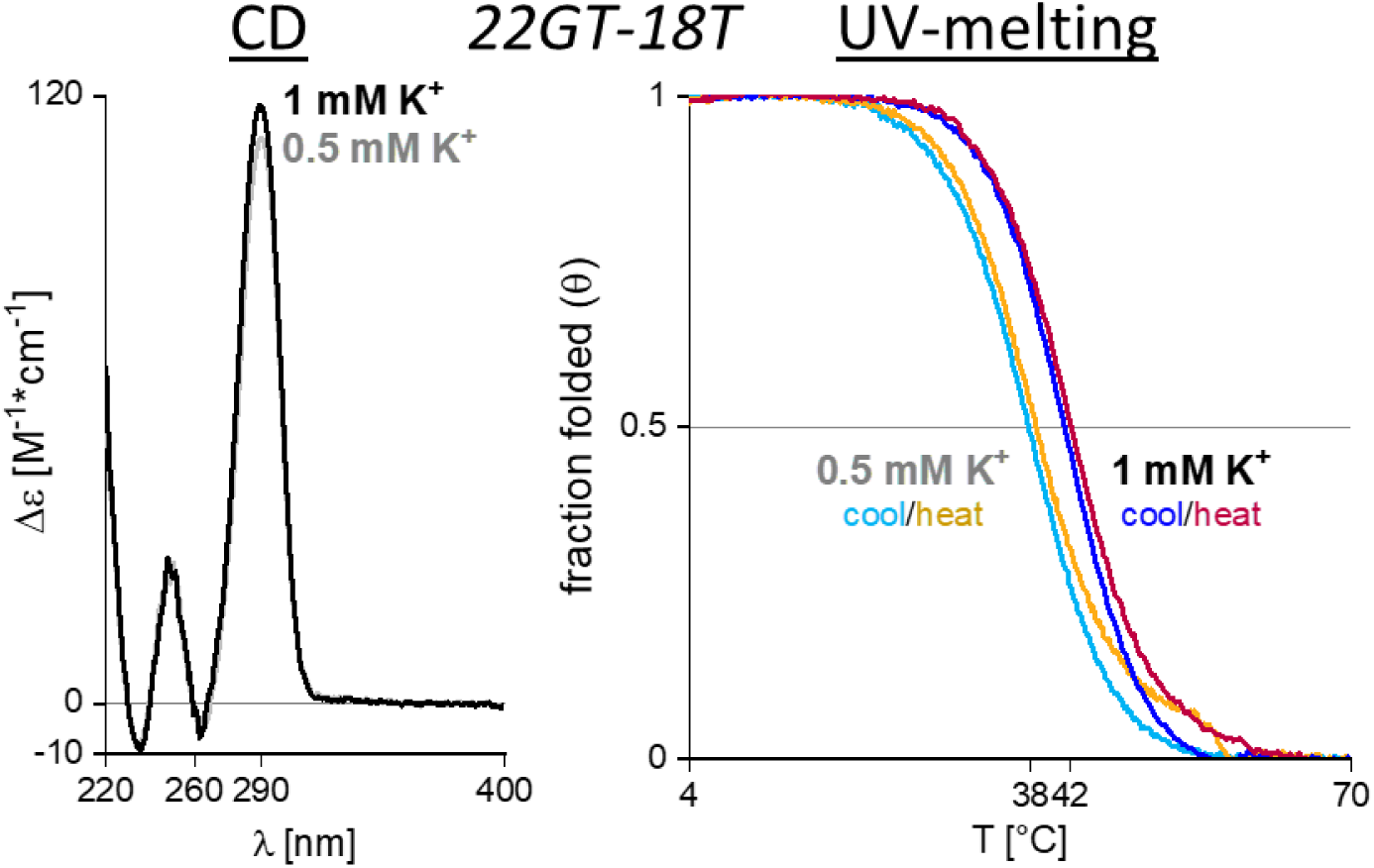
CD spectra of 22GT-18T (d**G_3_**TTA**G_3_**TTA**G_3_**<u>TTT</u>**G_3_**T) in 1 mM (black) and 0.5 mM KCl (grey). **Right**: UV-melting curves in 1 mM (blue/red) and 0.5 mM (cyan/orange) KCl. Samples contain 10 µM DNA and 100 mM TMAA (pH 6.8).

**Figure S26.**
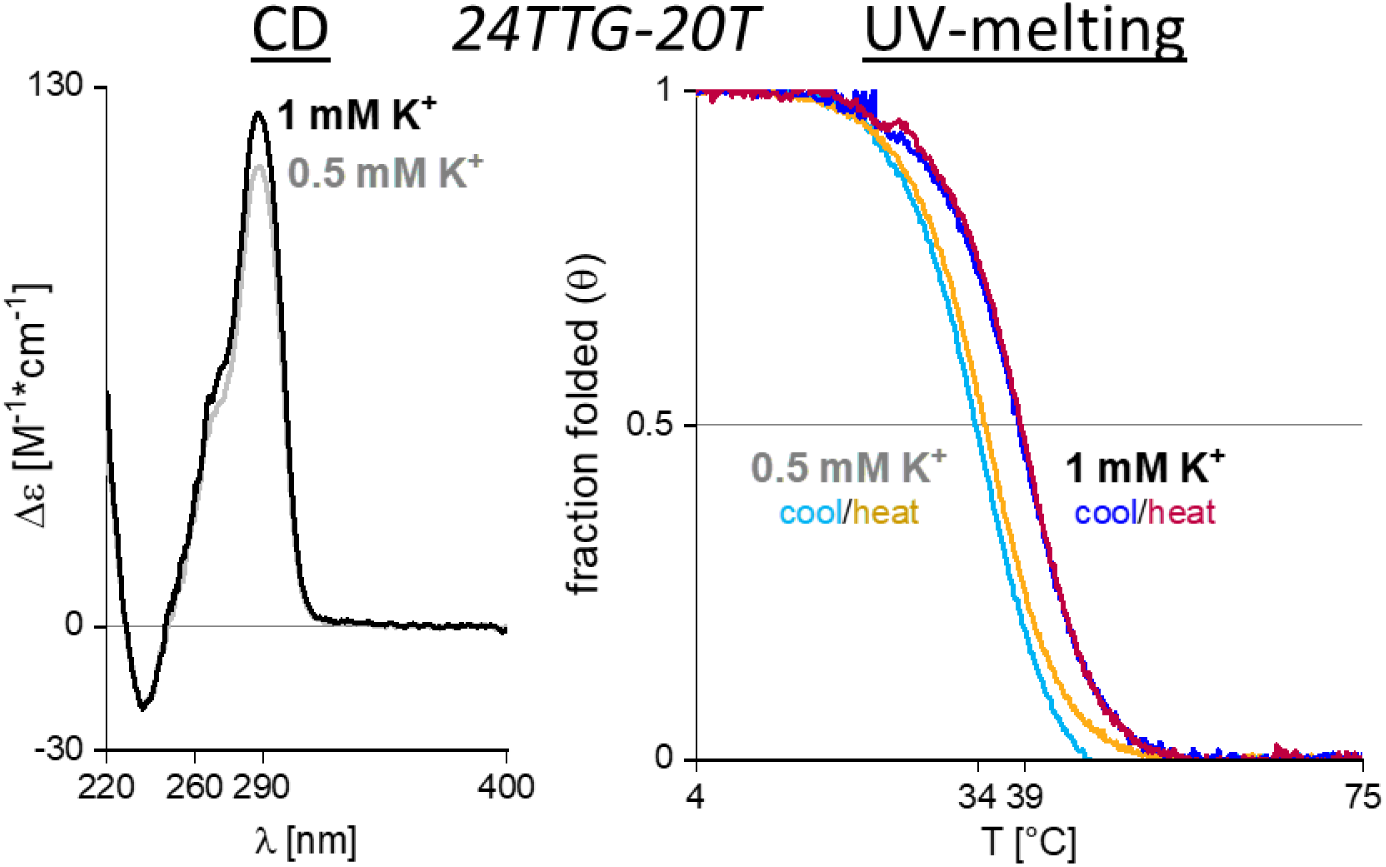
CD spectra of 24TTG-20T (dTT**G_3_**TTA**G_3_**TTA**G_3_**<u>TTT</u>**G_3_**A) in 1 mM (black) and 0.5 mM KCl (grey). **Right**: UV-melting curves in 1 mM (blue/red) and 0.5 mM (cyan/orange) KCl. Samples contain 10 µM DNA and 100 mM TMAA (pH 6.8).

**Figure S27.**
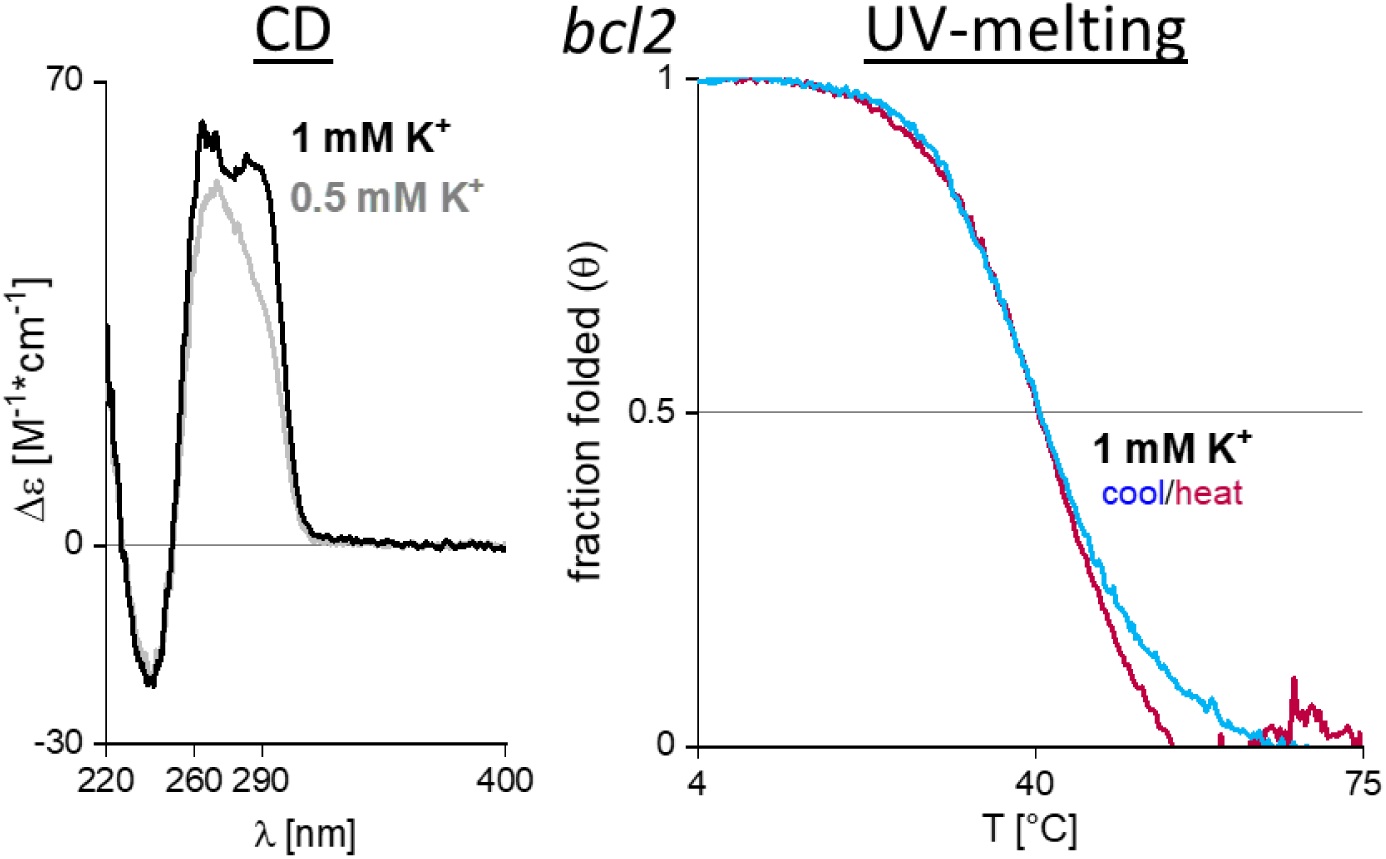
CD spectra of bcl2 (d**G_3_**C**G**C**G_3_**A**GG**AATT**G_3_**C**G_3_**) in 1 mM (black) and 0.5 mM KCl (grey). **Right**: UV-melting curve in 1 mM (blue/red) KCl. No data could be obtained in 0.5 mM KCl due to lack of a low temperature baseline. Samples contain 10 µM DNA and 100 mM TMAA (pH 6.8).

Figure S28 lets us directly compare the CD signatures of the mutated sequences with their ‘wild-type’ versions.

**Figure S28.**
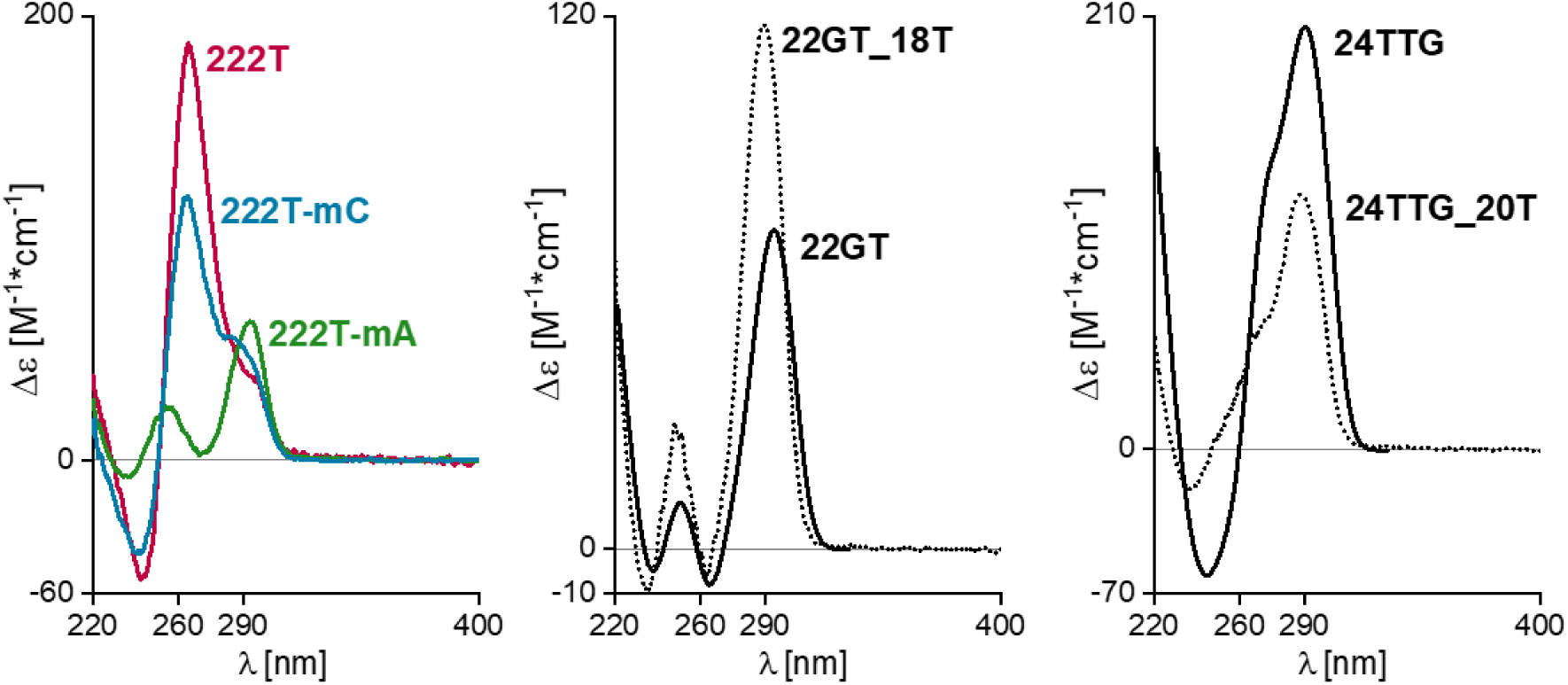
Comparing CD signatures of ‘wild-type’ and mutant sequences. **Left**: The 222T sequence motif (dT**G_3_**TT**G_3_**<u>TT</u>**G_3_**TT**G_3_**T) in 0.5 mM KCl **Middle**: the 22GT sequence motif (d**G_3_**TTA**G_3_**TTA**G_3_**<u>TTA</u>**G_3_**T) in 1 mM KCl **Right**: The 24TTG sequence motif (dTT**G_3_**TTA**G_3_**TTA**G_3_**<u>TTA</u>**G_3_**A) in 1 mM KCl. Samples contain 10 µM DNA, 0.5 or 1 mM KCl and 100 mM TMAA (pH 6.8). ‘wild-type’ signatures of 22GT and 24TTG are taken from previously published data.^2^

### CD curves of multi-stranded G-quadruplexes in NH_4_^+^ and K^+^

The screening panel contains tetramolecular G-quadruplex (dTG4T)4 and bimolecular G-quadruplex (dG4T4G4)2. We know that these quadruplexes take weeks/months to fold completely in solution. We expected the formation to be notoriously slow in 0.5 mM K^+^, which is why we measured another sample in 150 mM NH_4_ ^+^, hoping to speed up the formation kinetics and obtain quantitative amounts of G-quadruplex.

We noticed slight changes in ligand binding when the cation was switched (see **Error! Reference source not found.**). We gathered CD spectra of both sequences in K^+^ and NH_4_ ^+^ (Figure S29) after having let the samples rest for 10/11 months. The CD signatures do not change enough to indicate that switching from K^+^ to NH_4_ ^+^ induces a change in topology. Since the true concentration of folded G-quadruplex is uncertain, we did not convert the ellipticity to Δε.

**Figure S29.**
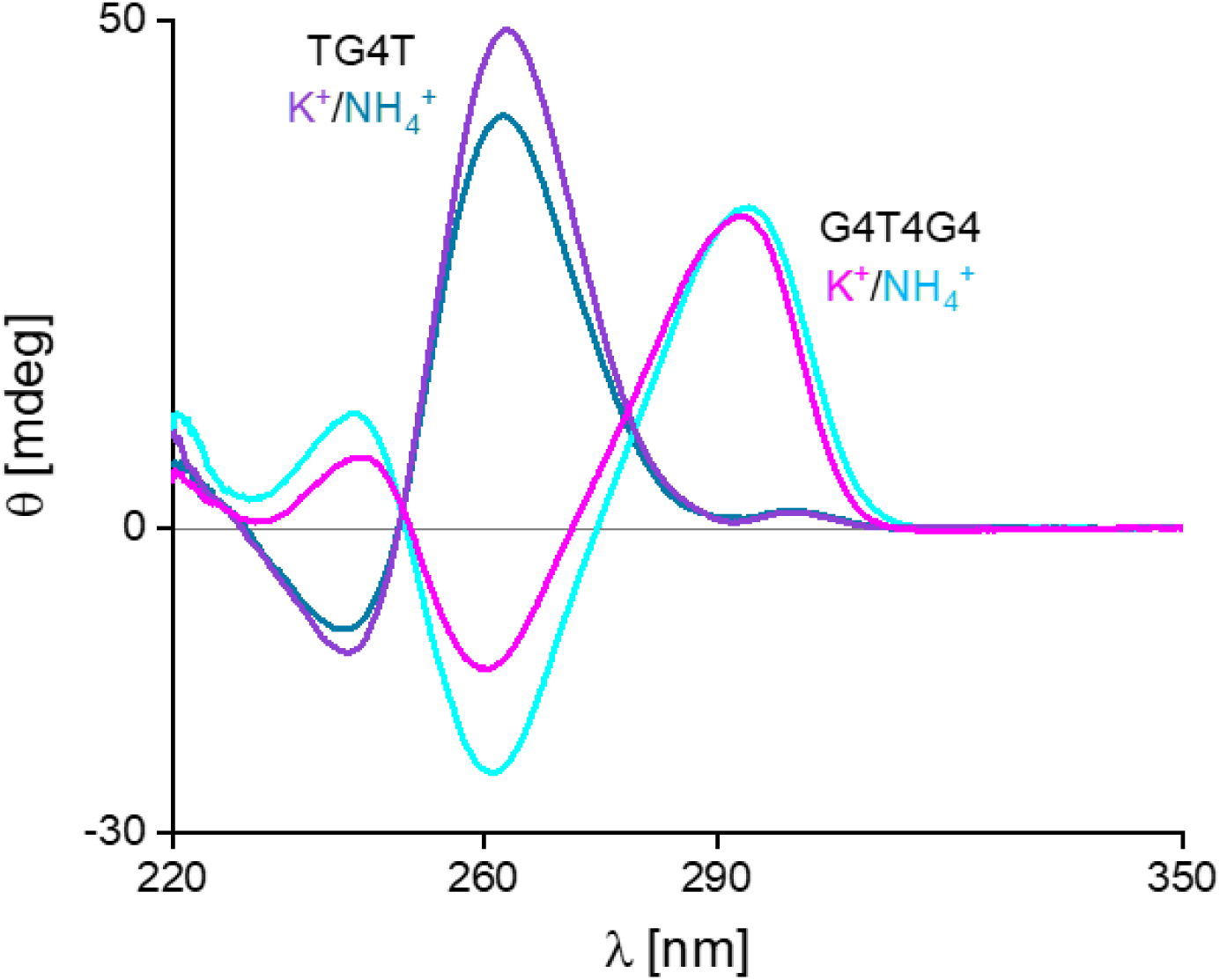
CD spectra of multi-stranded G-quadruplexes TG4T and G4T4G4 in different cation buffers. Samples contain: 10 µM G4T4G4/20 µM TG4T, 150 mM ammonium acetate (pH 6.8) or 0.5 mM KCl and 100 mM TMAA (pH 6.8).

### Parameters of the ESI-IMS-QTOF instrument

**Table S1.**
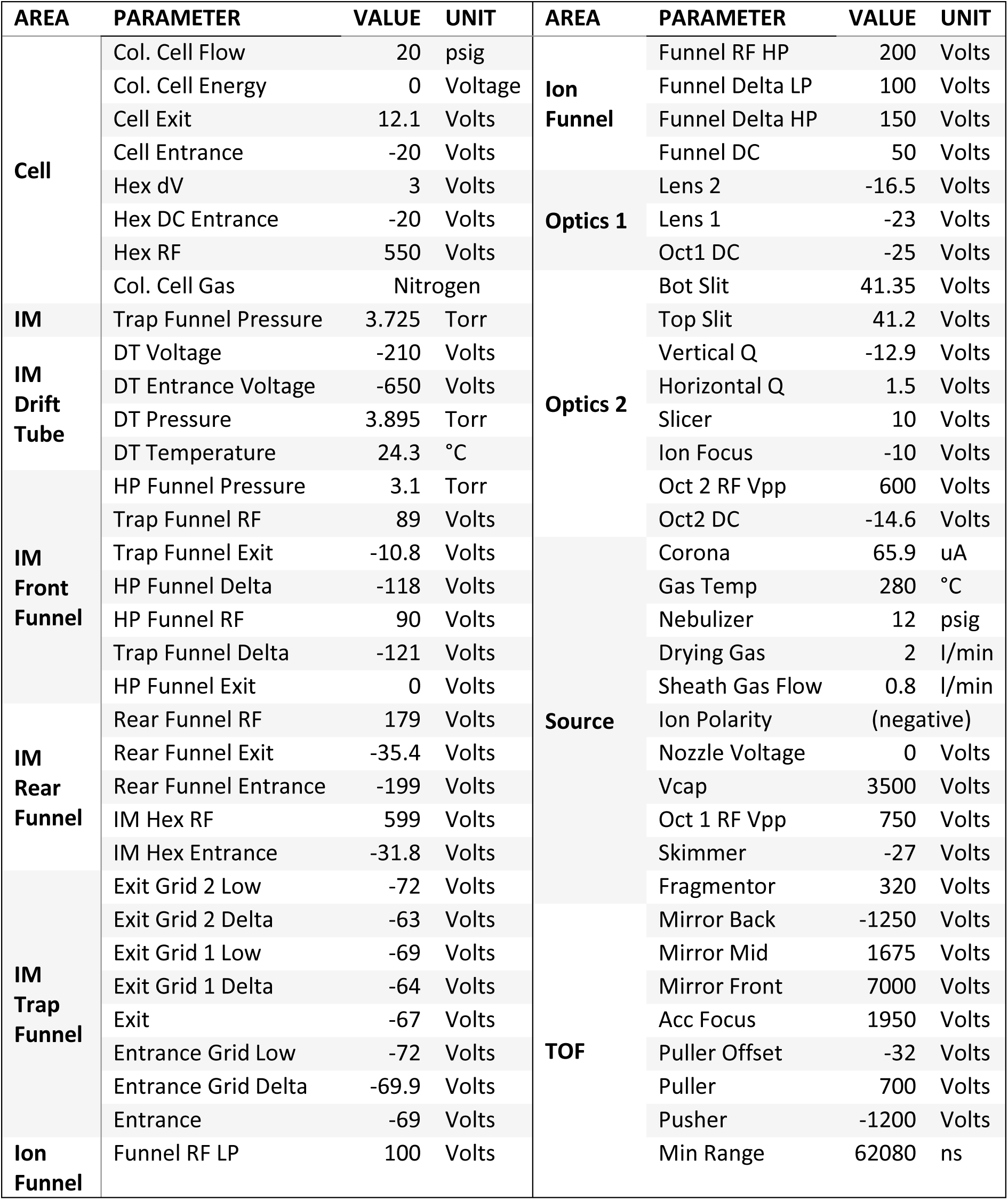
Instrumental settings of the Agilent 6560 IMS-QTOF, optimized for native MS on oligonucleotides.

For the location of all the parts listed consult page 5 of the [overview provided by Agilent](https://www.agilent.com/cs/library/technicaloverviews/public/5991-3244EN.pdf). I cannot reprint the schematic in here because of copyright.

### Generating CCS distributions from ATD distributions

The instrument measures the arrival time (tA) as the time between the release of ions from the trapping funnel until their detection at the TOF mass analyzer. The ion spends a portion of that time in the drift tube (drift time: tD) and the other portion inside the rest of the instrument (dead time: t0). We separate those contributions by operating the drift tube at different drift voltages. Figure S30 shows the shift of arrival time distribution (ATD) at different drift voltages (ΔV) from our external calibrant, which is a solution of 40 mM dTG4T in 150 mM ammonium acetate.

**Figure S30.**
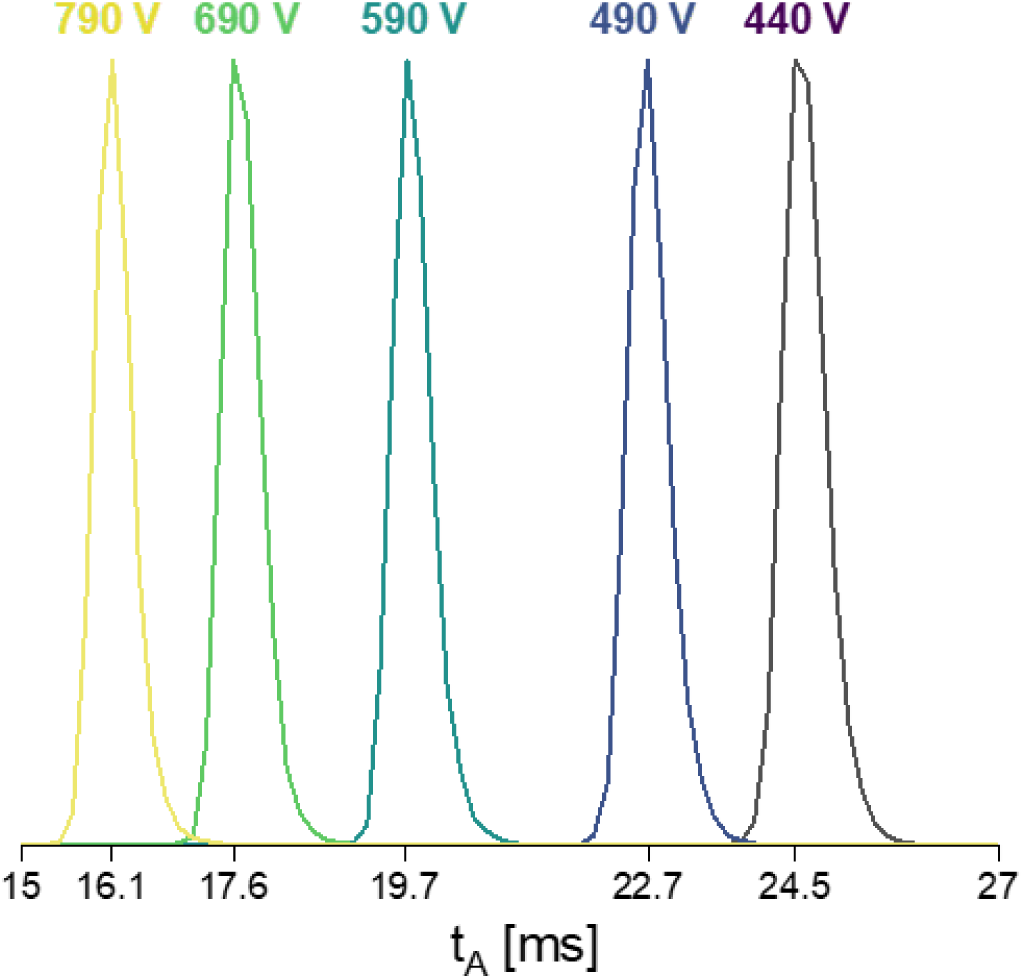
Arrival time distributions of the G-quadruplex species [(dTG_4_T)_4_*3NH ^+^]^5^⁻ at different drift voltages.

The drift time *t_D_* is dependent on the drift voltage *ΔV*, whereas the dead time is not.

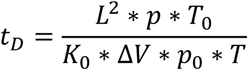

Where *L* is the length of the drift tube (78.1 cm), *p_0_/T_0_* are the standard pressure/temperature (1 bar, 273 K), *p/T* are the pressure/temperature inside the drift tube (which are recorded for each experiment) and *K_0_* is the reduced mobility of the ion species (In this case: [(dTG4T)4*3NH ^+^]^5^⁻). Based on this, we can formulate the correlation between arrival time and drift voltage as a linear function.

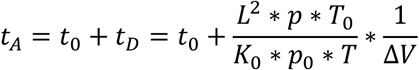

The plot is shown in Figure S31. We extract the dead time from the y-intercept and the reduced ion mobility from the slope. We convert the reduced ion mobility K_0_ to the momentum transfer cross section *Ω* (commonly referred to as the collisional cross section *CCS*) using the Mason-Schamp equation. Note that this equation follows the assumption of a static drift gas and low-field conditions.

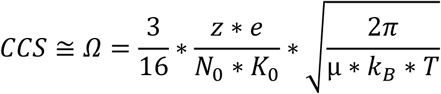

The equation features several constants: *e*, *kB* and *N0* (Loschmidt constant). The temperature *T* and the reduced mass *µ* (µ = (Mion*MHe)/(Mion+MHe)) remain practically constant within the experimental conditions. The reduced mobility *K_0_* and charge state *z* will be different for every ion species.

This method produces a single CCS-value which corresponds to the peak maxima of the arrival time distributions. For the calibrant TG4T, our CCS-values are 788.7 Å² for the 5-ion (ref. Figure S31) and 740.4 Å² for the 4-ion (not shown). These values are in good agreement with previously published experimental results (787.5 Å² [5-], 735.7 Å² [4-]).^3^

**Figure S31.**
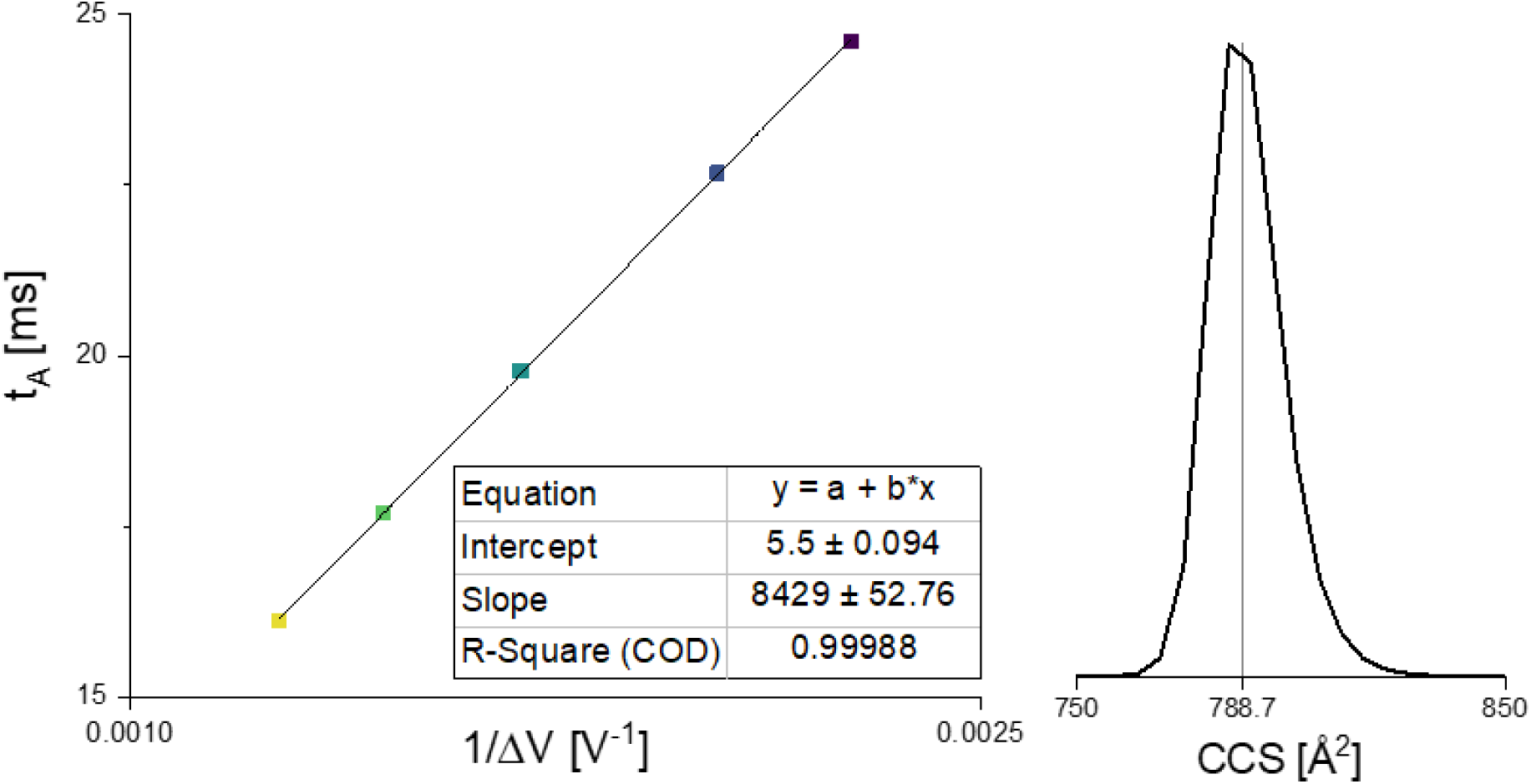
**Left**: Obtaining drift time and reduced ion mobility by linearly fitting the arrival time at different drift voltages. **Right**: CCS distribution of [(dTG_4_T)_4_*3NH ^+^]^5^⁻, calculated from the reduced ion mobility using the Mason-Schamp equation.

We generate CCS distributions by converting the arrival time *t_A_* to the respective CCS value using the ratio of CCS value and arrival time at peak maximum as a proportionality factor.

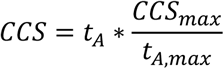

Conceptually, any of the five ATDs shown in Figure S30 can be converted using this method. To be as close as possible to the low-field limit, we consistently choose the ATD at the lowest drift voltage.

### ESI-MS titrations: Data processing

Using dT_6_ as an internal calibrant we quantify the change in DNA response.^4^ Figure S32 shows how the ratio of collective DNA signal vs. dT_6_ signal evolves as a function of ligand concentration. The estimated response factors are listed in table. Based on the estimates we allowed the response factors for the complex signals to fluctuate between 0.8 and 1.1 during dynamic fitting.

**Figure S32.**
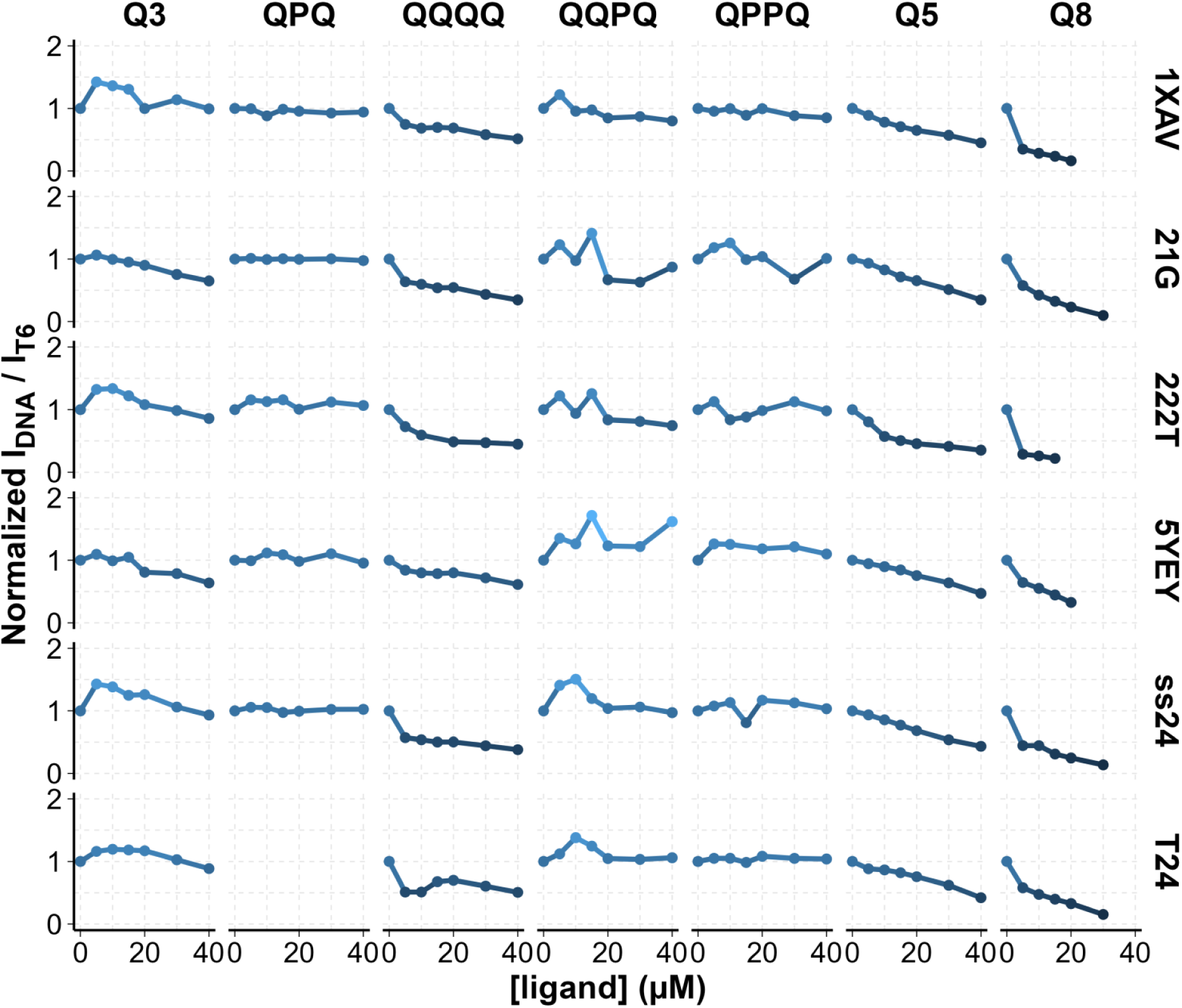
The sum of all DNA signal intensities vs. the signal of calibrant dT_6_. The first datapoint is normalized to 1.

A declining curve means that adding ligand negatively impacts the detection of DNA ions. This leads to an ever-decreasing S/N ratio until the DNA species fall under the limit of detection. The latter happened with Q8, which is why the titration datasets with Q8 are incomplete.

The titration data points, corrected for noise and response, are shown in Figure S33, alongside the dynamic fits from which we obtained refined K_D_ values. All K_D_ values are listed in Table S3.

**Figure S33.**
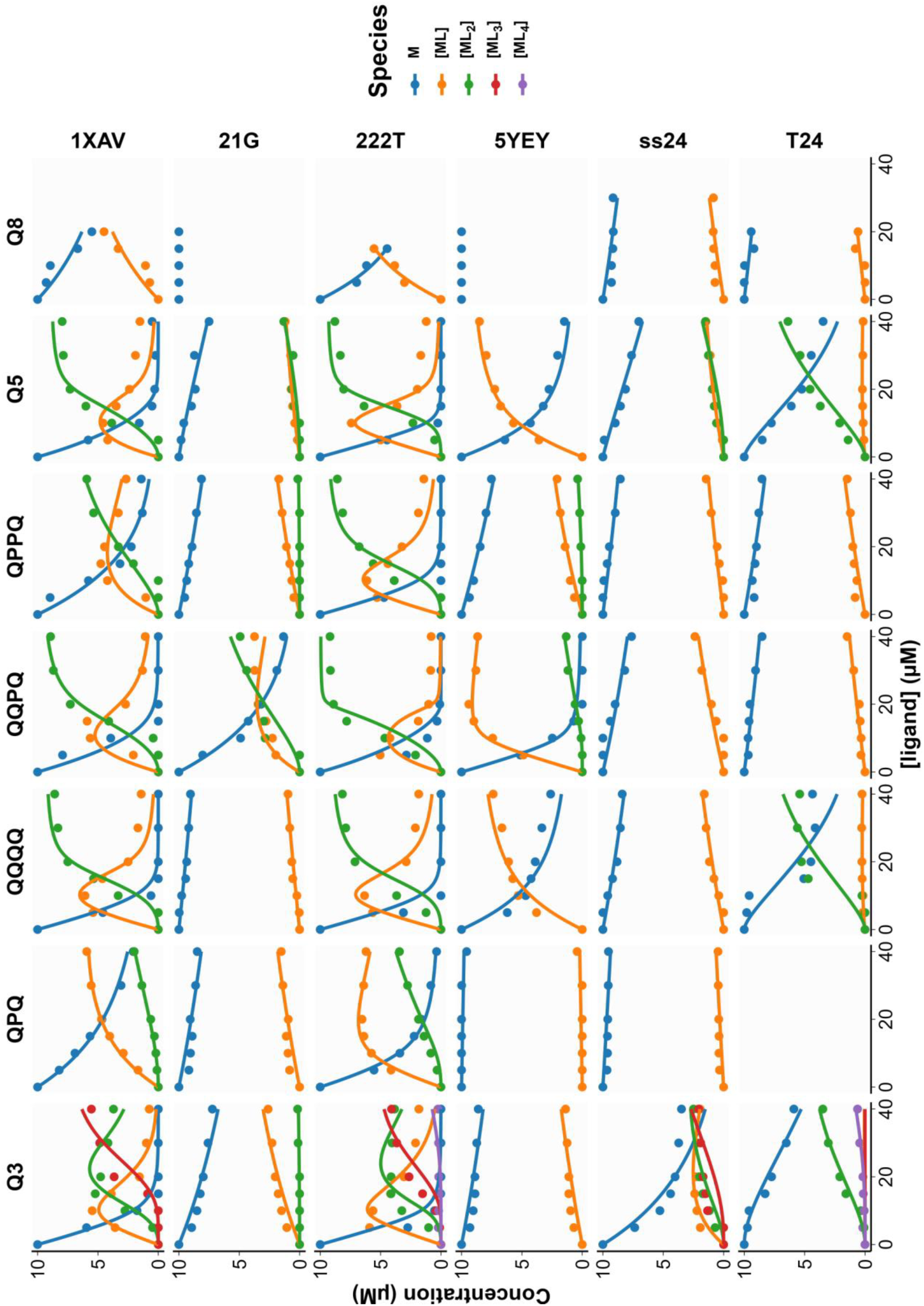
Data points obtained from processing the ESI-MS titration mass spectra. K_D_ values are calculated by dynamically fitting the experimental data based on the complex formation equilibria. Curves show the dynamic fits. Samples contain 10 µM DNA, 0-40 µM ligand, 4 µM dT_6_, 0.5 mM KCl and 100 mM TMAA (pH 6.8).

### Ligand screening: Comprehensive list of DNA/Ligand species concentrations and their K_D_ values

**Table S2.** Data extracted from the ligand screening, featuring: Concentration of unbound DNA (M), 1:1 complex (ML) and 2:1 complex (ML_2_), fraction of DNA bound (fb), complex dissociation constants K_D1_ and K_D2_ as well as ligand cooperativity. Sample conditions were: 10 µM DNA, 20 µM ligand, 0.5 mM KCl, 100 mM TMAA (pH 6.8).

| DNA | Ligand | M<br>[μM] | ML<br>[μM] | ML<sub>2</sub><br>[μM] | fb | $K_{D1}$<br>[μM] | $K_{D2}$<br>[μM] | Coop.<sup>2</sup> |
| --- | --- | --- | --- | --- | --- | --- | --- | --- |
| 1XAV | QQPQ | 0.4 | 3.7 | 5.9 | 0.96 | 0.5 | 1.5 | positive |
| 1XAV | QQQQ | 0.3 | 3.2 | 6.5 | 0.97 | 0.3 | 0.9 | positive |
| 1XAV | QPPQ | 2.5 | 4.6 | 3.0 | 0.75 | 5.2 | 7.4 | positive |
| 1XAV | QPQ | 4.7 | 4.2 | 1.1 | 0.53 | 15 | 27 | positive |
| 222T | QQPQ | 0.4 | 4.3 | 5.3 | 0.96 | 0.5 | 2.1 | negative |
| 222T | QQQQ | 0.8 | 3.9 | 5.4 | 0.92 | 1.1 | 2.0 | positive |
| 222T | QPPQ | 1.4 | 5.0 | 3.6 | 0.86 | 2.3 | 5.4 | positive |
| 222T | QPQ | 2.6 | 5.9 | 1.5 | 0.74 | 4.8 | 22 | negative |
| 222T_mA | QQPQ | 1.2 | 2.6 | 6.3 | 0.88 | 2.2 | 1.0 | positive |
| 222T_mA | QQQQ | 3.6 | 3.0 | 3.4 | 0.64 | 12 | 4.5 | positive |
| 222T_mA | QPPQ | 3.0 | 3.0 | 4.0 | 0.70 | 8.9 | 3.4 | positive |
| 222T_mA | QPQ | 5.5 | 3.0 | 1.5 | 0.45 | 25 | 14 | positive |
| 222T_mC | QQPQ | 0.5 | 2.9 | 6.5 | 0.95 | 0.7 | 0.9 | positive |
| 222T_mC | QQQQ | 1.9 | 4.0 | 4.1 | 0.81 | 3.7 | 3.7 | positive |
| 222T_mC | QPPQ | 2.2 | 3.7 | 4.1 | 0.78 | 4.9 | 3.6 | positive |
| 222T_mC | QPQ | 3.0 | 5.1 | 1.9 | 0.70 | 6.6 | 15 | positive |
| T30177TT | QQPQ | 0.9 | 4.7 | 4.4 | 0.91 | 1.2 | 3.4 | positive |
| T30177TT | QQQQ | 0.7 | 4.5 | 4.7 | 0.93 | 1.0 | 2.9 | positive |
| T30177TT | QPPQ | 1.5 | 6.2 | 2.4 | 0.85 | 2.1 | 12 | negative |
| T30177TT | QPQ | 2.9 | 6.2 | 0.8 | 0.71 | 5.7 | 47 | negative |
| 26CEB | QQPQ | 1.2 | 8.2 | 0.5 | 0.88 | 1.6 | 83 | negative |
| 26CEB | QQQQ | 0.7 | 8.4 | 0.9 | 0.93 | 0.8 | 45 | negative |
| 26CEB | QPPQ | 1.8 | 7.7 | 0.5 | 0.82 | 2.6 | 95 | negative |
| 26CEB | QPQ | 3.3 | 6.4 | 0.3 | 0.67 | 6.6 | 136 | negative |
| 26CEB_mT | QQPQ | 1.2 | 7.9 | 0.9 | 0.88 | 1.5 | 43 | negative |
| 26CEB_mT | QQQQ | 2.2 | 6.2 | 1.6 | 0.78 | 3.8 | 21 | negative |
| 26CEB_mT | QPPQ | 2.1 | 7.3 | 0.7 | 0.79 | 3.3 | 63 | negative |
| 26CEB_mT | QPQ | 3.6 | 6.0 | 0.4 | 0.64 | 8.1 | 103 | negative |
| 2KYP | QQPQ | 2.2 | 2.4 | 5.4 | 0.78 | 6.2 | 1.5 | positive |
| 2KYP | QQQQ | 2.7 | 6.6 | 0.7 | 0.73 | 4.9 | 57 | negative |
| 2KYP | QPPQ | 7.1 | 1.7 | 1.2 | 0.29 | 66 | 11 | positive |
| 2KYP | QPQ | 8.0 | 1.7 | 0.3 | 0.20 | 84 | 47 | positive |
| 2O3M | QQPQ | 7.2 | 1.6 | 1.2 | 0.28 | 71 | 11 | positive |
| 2O3M | QQQQ | 7.8 | 2.2 | 0 | 0.22 | 64 | 0 |  |
| 2O3M | QPPQ | 7.5 | 2.2 | 0.3 | 0.25 | 58 | 58 | positive |
| 2O3M | QPQ | 8.6 | 1.4 | 0 | 0.14 | 119 | 0 |  |

<sup>2</sup> Positive, if $4 \cdot K_{D1} > K_{D2}$ . Negative, if $4 \cdot K_{D1} < K_{D2}$ . Assuming independent and equivalent binding sites.<sup>5</sup>

|  |  |  |  |  |  |  |  |  |
| --- | --- | --- | --- | --- | --- | --- | --- | --- |
| TG4T_K | QQPQ | 4.2 | 5.0 | 0.8 | 0.58 | 11 | 43 | positive |
| TG4T_K | QQQQ | 7.3 | 2.7 | 0 | 0.27 | 46 | 0 |  |
| TG4T_K | QPPQ | 5.7 | 3.8 | 0.5 | 0.43 | 23 | 60 | positive |
| TG4T_K | QPQ | 8.0 | 2.0 | 0 | 0.20 | 70 | 0 |  |
| TG4T_NH4 | QQPQ | 2.8 | 6.2 | 1.0 | 0.72 | 5.3 | 37 | negative |
| TG4T_NH4 | QQQQ | 8.9 | 1.1 | 0 | 0.11 | 149 | 0 |  |
| TG4T_NH4 | QPPQ | 5.8 | 3.9 | 0.3 | 0.42 | 23 | 101 | negative |
| TG4T_NH4 | QPQ | 7.5 | 2.3 | 0.2 | 0.25 | 58 | 114 | positive |
| 21G | QQPQ | 5.9 | 2.6 | 1.5 | 0.41 | 32 | 13 | positive |
| 21G | QQQQ | 9.1 | 0.9 | 0 | 0.09 | 197 | 0 |  |
| 21G | QPPQ | 8.4 | 1.3 | 0.2 | 0.16 | 116 | 53 | positive |
| 21G | QPQ | 9.3 | 0.7 | 0 | 0.07 | 250 | 0 |  |
| 5YEY | QQPQ | 1.2 | 7.3 | 1.6 | 0.88 | 1.5 | 22 | negative |
| 5YEY | QQQQ | 5.0 | 5.0 | 0 | 0.50 | 15 | 0 |  |
| 5YEY | QPPQ | 7.8 | 1.8 | 0.4 | 0.22 | 75 | 40 | positive |
| 5YEY | QPQ | 9.4 | 0.6 | 0 | 0.06 | 309 | 0 |  |
| 22GT | QQPQ | 5.5 | 3.1 | 1.4 | 0.45 | 25 | 16 | positive |
| 22GT | QQQQ | 8.5 | 1.5 | 0 | 0.15 | 104 | 0 |  |
| 22GT | QPPQ | 8.0 | 1.6 | 0.4 | 0.20 | 89 | 36 | positive |
| 22GT | QPQ | 9.1 | 0.9 | 0 | 0.09 | 196 | 0 |  |
| 22GT_18T | QQPQ | 2.6 | 6.5 | 0.9 | 0.74 | 4.6 | 41 | negative |
| 22GT_18T | QQQQ | 5.4 | 4.6 | 0 | 0.46 | 18 | 0 |  |
| 22GT_18T | QPPQ | 7.7 | 1.9 | 0.4 | 0.23 | 69 | 45 | positive |
| 22GT_18T | QPQ | 8.8 | 1.2 | 0.0 | 0.12 | 143 | 0 |  |
| 22CTA | QQPQ | 7.0 | 2.0 | 1.0 | 0.30 | 57 | 16 | positive |
| 22CTA | QQQQ | 8.6 | 1.4 | 0 | 0.14 | 114 | 0 |  |
| 22CTA | QPPQ | 8.6 | 1.4 | 0 | 0.14 | 116 | 0 |  |
| 22CTA | QPQ | 8.8 | 1.2 | 0 | 0.12 | 135 | 0 |  |
| TBA | QQPQ | 6.7 | 3.3 | 0 | 0.33 | 34 | 0 |  |
| TBA | QQQQ | 8.1 | 1.9 | 0 | 0.19 | 78 | 0 |  |
| TBA | QPPQ | 8.6 | 1.4 | 0 | 0.14 | 118 | 0 |  |
| TBA | QPQ | 9.3 | 0.7 | 0 | 0.07 | 252 | 0 |  |
| G4T4G4_K | QQPQ | 6.1 | 3.3 | 0.6 | 0.39 | 28 | 39 | positive |
| G4T4G4_K | QQQQ | 8.2 | 1.8 | 0 | 0.18 | 81 | 0 |  |
| G4T4G4_K | QPPQ | 7.9 | 2.1 | 0 | 0.21 | 67 | 0 |  |
| G4T4G4_K | QPQ | 8.4 | 1.6 | 0 | 0.16 | 99 | 0 |  |
| G4T4G4_NH4 | QQPQ | 5.8 | 3.7 | 0.5 | 0.42 | 24 | 53 | positive |
| G4T4G4_NH4 | QQQQ | 9.7 | 0.3 | 0 | 0.03 | 562 | 0 |  |
| G4T4G4_NH4 | QPPQ | 9.2 | 0.8 | 0 | 0.08 | 225 | 0 |  |
| G4T4G4_NH4 | QPQ | 9.6 | 0.4 | 0 | 0.04 | 470 | 0 |  |
| 26TTA<sup>3</sup> | QQPQ | 2.1 | 3.8 | 3.3 | 0.71 | 4.1 | 4 | positive |
| 26TTA | QQQQ | 6.8 | 2.2 | 1.0 | 0.32 | 48 | 18 | positive |
| 26TTA | QPPQ | 7.7 | 1.9 | 0.4 | 0.23 | 70 | 40 | positive |
| 26TTA | QPQ | 9.1 | 0.9 | 0 | 0.09 | 204 | 0 |  |

<sup>3</sup> This is the only instance where we observed a 3:1 complex. [ML<sub>3</sub>] = 0.8 μM, K<sub>D3</sub> = 22 μM (negative coop.).

|  |  |  |  |  |  |  |  |  |
| --- | --- | --- | --- | --- | --- | --- | --- | --- |
| Bcl2 | QQPQ | 4.4 | 3.4 | 2.1 | 0.56 | 16 | 9.8 | positive |
| Bcl2 | QQQQ | 4.0 | 4.0 | 1.9 | 0.60 | 12 | 13 | positive |
| Bcl2 | QPPQ | 5.3 | 3.6 | 1.1 | 0.47 | 21 | 24 | positive |
| Bcl2 | QPQ | 7.5 | 2.5 | 0 | 0.25 | 52 | 0 |  |
| 24TTG | QQPQ | 7.1 | 1.1 | 1.8 | 0.29 | 102 | 4.6 | positive |
| 24TTG | QQQQ | 9.8 | 0.2 | 0 | 0.02 | 809 | 0 |  |
| 24TTG | QPPQ | 8.8 | 1.2 | 0 | 0.12 | 145 | 0 |  |
| 24TTG | QPQ | 9.4 | 0.6 | 0 | 0.06 | 284 | 0 |  |
| 24TTG_20T | QQPQ | 5.2 | 3.6 | 1.2 | 0.48 | 20 | 22 | positive |
| 24TTG_20T | QQQQ | 6.0 | 4.0 | 0 | 0.40 | 24 | 0 |  |
| 24TTG_20T | QPPQ | 8.3 | 1.7 | 0 | 0.17 | 92 | 0 |  |
| 24TTG_20T | QPQ | 8.9 | 1.1 | 0 | 0.11 | 153 | 0 | positive |
| 23TAG | QQPQ | 5.7 | 3.4 | 1.0 | 0.43 | 25 | 25 | positive |
| 23TAG | QQQQ | 8.9 | 1.1 | 0 | 0.11 | 148 | 0 |  |
| 23TAG | QPPQ | 8.1 | 1.9 | 0 | 0.19 | 75 | 0 |  |
| 23TAG | QPQ | 8.3 | 1.7 | 0 | 0.17 | 93 | 0 |  |
| 2KPR | QQPQ | 4.6 | 1.9 | 3.5 | 0.54 | 26 | 3 | positive |
| 2KPR | QQQQ | 8.7 | 1.3 | 0 | 0.13 | 131 | 0 |  |
| 2KPR | QPPQ | 7.0 | 2.0 | 1.0 | 0.30 | 56 | 17 | positive |
| 2KPR | QPQ | 8.6 | 1.1 | 0.3 | 0.14 | 144 | 37 | positive |
| 21CCC (i-m) | QQPQ | 9.4 | 0.6 | 0 | 0.06 | 294 | 0 |  |
| 21CCC (i-m) | QQQQ | 10 | 0 | 0 | 0.00 | 0 | 0 |  |
| 21CCC (i-m) | QPPQ | 9.4 | 0.6 | 0 | 0.06 | 290 | 0 |  |
| 21CCC (i-m) | QPQ | 9.8 | 0.2 | 0 | 0.02 | 1280 | 0 |  |
| ds26 | QQPQ | 8.7 | 1.3 | 0 | 0.13 | 120 | 0 |  |
| ds26 | QQQQ | 10 | 0 | 0 | 0.00 | 0 | 0 |  |
| ds26 | QPPQ | 8.8 | 0 | 1.2 | 0.12 | 0 | 0 |  |
| ds26 | QPQ | 10 | 0 | 0 | 0.00 | 0 | 0 |  |
| DK33 | QQPQ | 8.5 | 1.5 | 0 | 0.15 | 107 | 0 |  |
| DK33 | QQQQ | 10 | 0 | 0 | 0.00 | 0 | 0 |  |
| DK33 | QPPQ | 8.1 | 1.9 | 0 | 0.19 | 80 | 0 |  |
| DK33 | QPQ | 8.8 | 1.2 | 0 | 0.12 | 141 | 0 |  |
| DK66 | QQPQ | 8.7 | 1.3 | 0 | 0.13 | 123 | 0 |  |
| DK66 | QQQQ | 10 | 0 | 0 | 0.00 | 3981 | 0 |  |
| DK66 | QPPQ | 8.3 | 1.7 | 0 | 0.17 | 88 | 0 |  |
| DK66 | QPQ | 9.7 | 0.3 | 0 | 0.03 | 586 | 0 |  |
| DK100 | QQPQ | 7.7 | 2.3 | 0 | 0.23 | 58 | 0 |  |
| DK100 | QQQQ | 10 | 0.0 | 0 | 0.00 | 0 | 0 |  |
| DK100 | QPPQ | 8.1 | 1.9 | 0 | 0.19 | 77 | 0 |  |
| DK100 | QPQ | 10 | 0 | 0 | 0.00 | 0 | 0 |  |
| ss24 | QQPQ | 8.2 | 1.8 | 0 | 0.18 | 83 | 0 |  |
| ss24 | QQQQ | 8.7 | 1.3 | 0 | 0.13 | 126 | 0 |  |
| ss24 | QPPQ | 9.1 | 0.9 | 0 | 0.09 | 182 | 0 |  |
| ss24 | QPQ | 9.7 | 0.3 | 0 | 0.03 | 626 | 0 |  |
| 24nonG4 | QQPQ | 9.2 | 0.8 | 0 | 0.08 | 208 | 0 |  |
| 24nonG4 | QQQQ | 7.2 | 2.8 | 0 | 0.28 | 44 | 0 |  |
| 24nonG4 | QPPQ | 7.5 | 2.5 | 0 | 0.25 | **51** | **0** |  |
| 24nonG4 | QPQ | 10 | 0 | 0 | 0.00 | **0** | **0** |  |
| T24 | QQPQ | 8.3 | 1.5 | 0.2 | 0.17 | **102** | **65** | positive |
| T24 | QQQQ | 6.8 | 0.4 | 2.8 | 0.32 | **235** | **1.0** | positive |
| T24 | QPPQ | 8.8 | 1.1 | 0.1 | 0.12 | **149** | **195** | positive |
| T24 | QPQ | 9.2 | 0.8 | 0 | 0.08 | **210** | **0** |  |
| A24 | QQPQ | 10 | 0 | 0 | 0.00 | **0** | **0** |  |
| A24 | QQQQ | 10 | 0 | 0 | 0.00 | **0** | **0** |  |
| A24 | QPPQ | 8.3 | 1.7 | 0 | 0.17 | **92** | **0** |  |
| A24 | QPQ | 8.8 | 1.2 | 0 | 0.12 | **138** | **0** |  |
| 21CCC (ss) | QQPQ | 9.5 | 0.5 | 0 | 0.05 | **354** | **0** |  |
| 21CCC (ss) | QQQQ | 10 | 0 | 0 | 0.00 | **4064** | **0** |  |
| 21CCC (ss) | QPPQ | 9.5 | 0.5 | 0 | 0.05 | **359** | **0** |  |
| 21CCC (ss) | QPQ | 9.9 | 0.1 | 0 | 0.01 | **3365** | **0** |  |
| T6 | QQPQ | 9.7 | 0.3 | 0 | 0.03 | **722** | **0** |  |
| T6 | QQQQ | 10 | 0 | 0 | 0.00 | **0** | **0** |  |
| T6 | QPPQ | 9.7 | 0.3 | 0 | 0.03 | **725** | **0** |  |
| T6 | QPQ | 9.8 | 0.2 | 0 | 0.02 | **1150** | **0** |  |

### Ligand screening – Mass spectra

**Figure S34.**
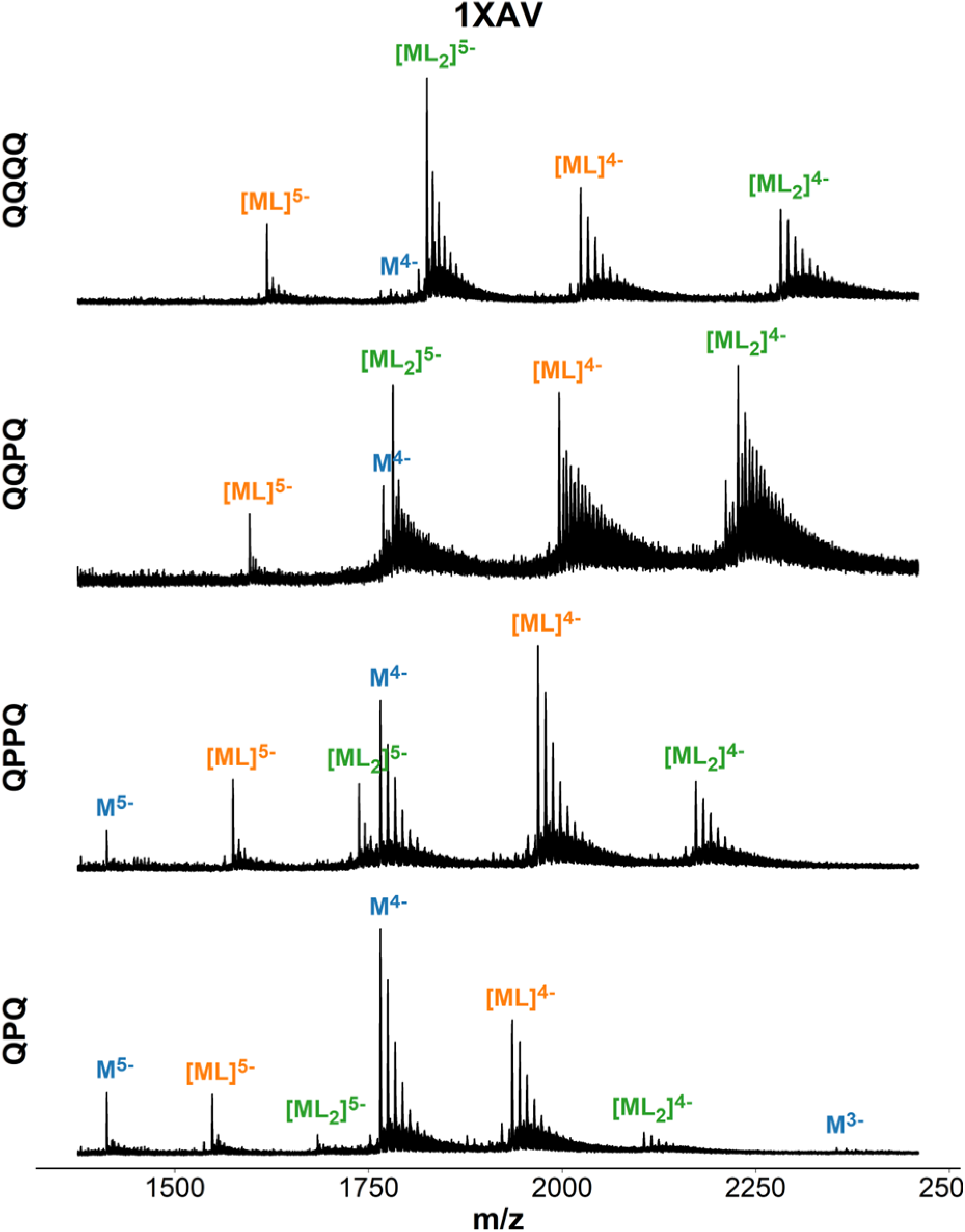
Mass spectra of 1XAV (dT**G**A**GGG**T**GGG**TA**GGG**T**GGG**TAA) in presence of ligand. Samples contain 10 µM DNA, 20 µM ligand, 0.5 mM KCl, 100 mM TMAA (pH 6.8).

**Figure S35.**
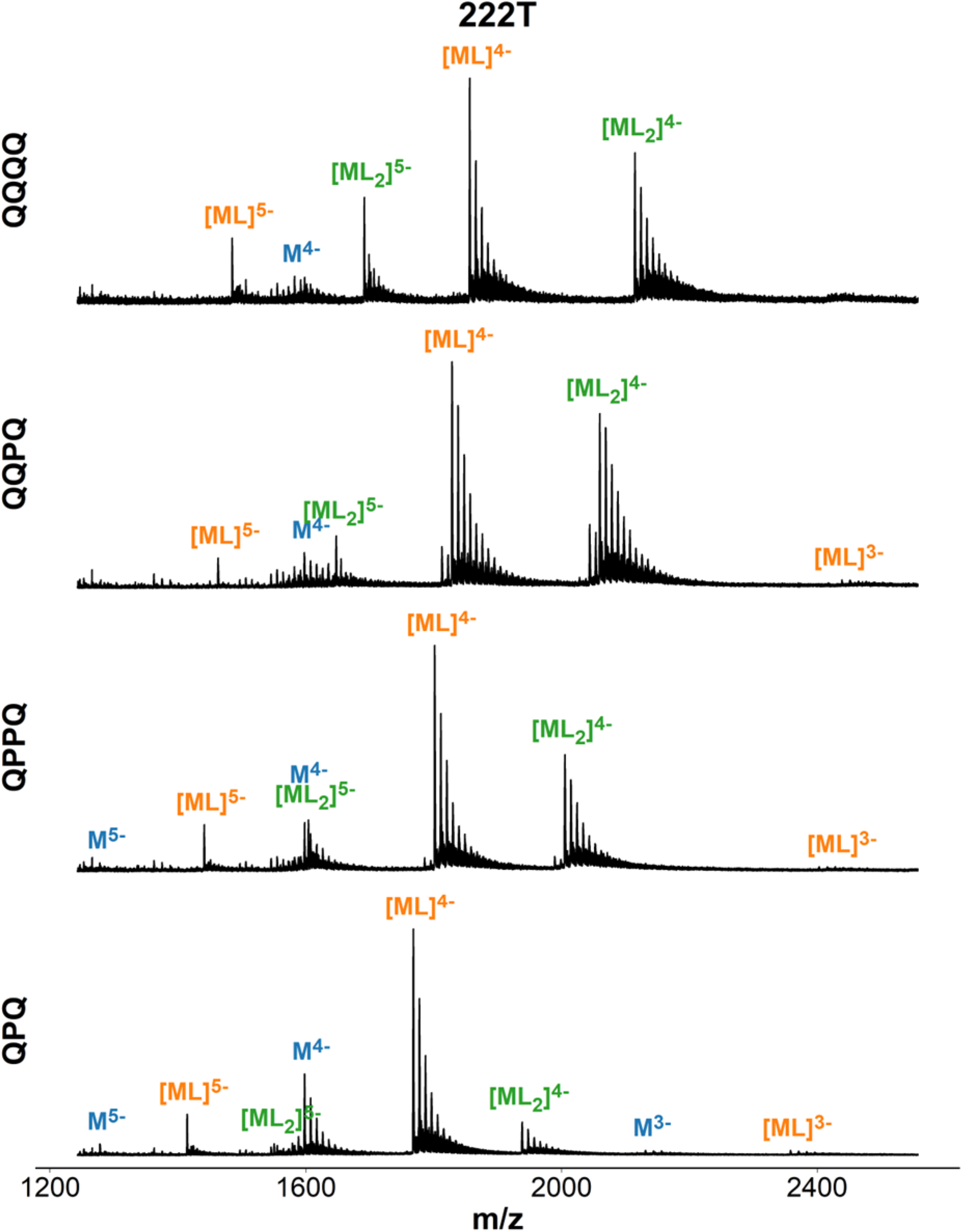
Mass spectra of 222T (dT**GGG**TT**GGG**TT**GGG**TT**GGG**T) in presence of ligand. Samples contain 10 µM DNA, 20 µM ligand, 0.5 mM KCl, 100 mM TMAA (pH 6.8).

**Figure S36.**
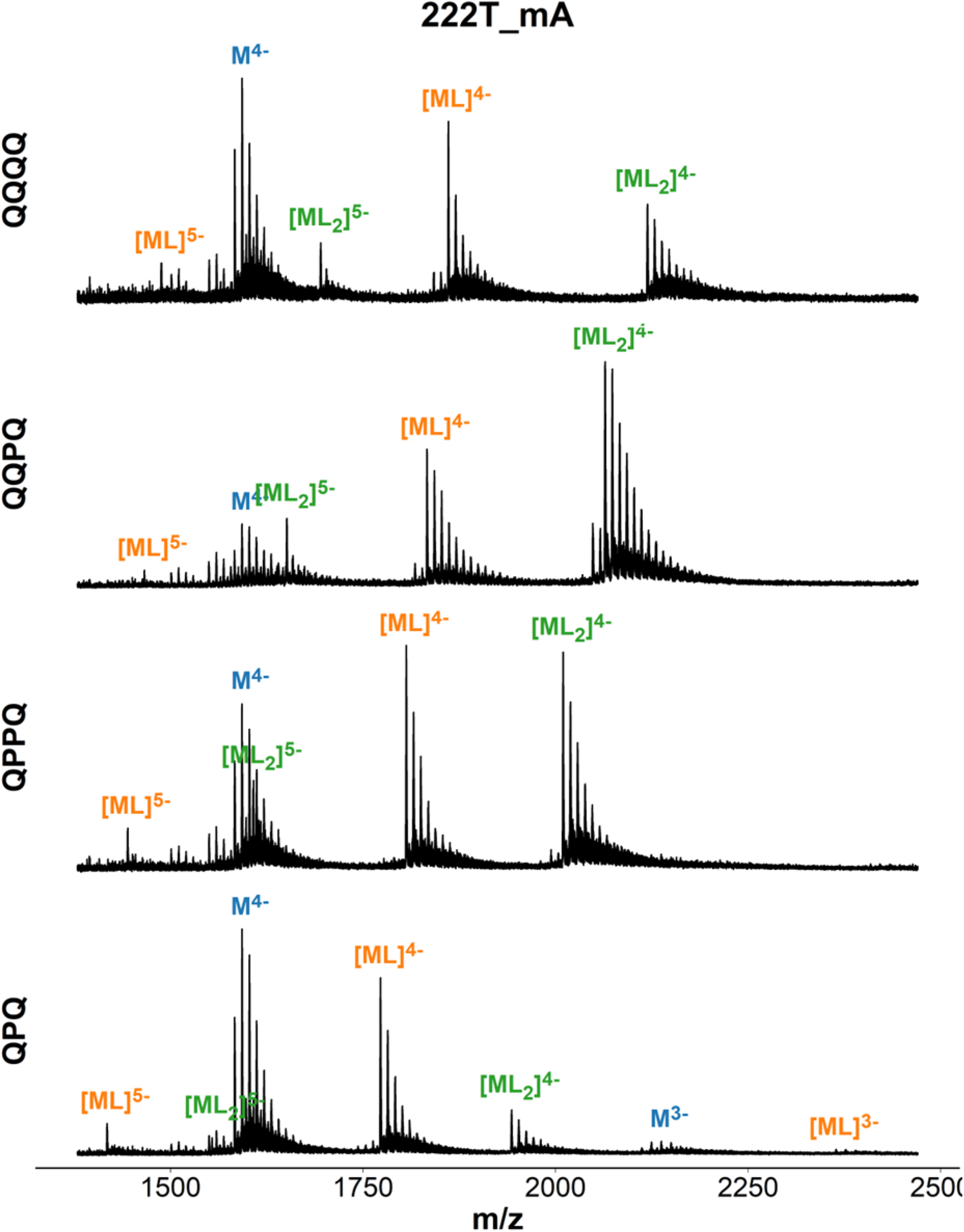
Mass spectra of 222T_mA (dT**GGG**TT**GGG**AA**GGG**TT**GGG**T) in presence of ligand. Samples contain 10 µM DNA, 20 µM ligand, 0.5 mM KCl, 100 mM TMAA (pH 6.8).

**Figure S37.**
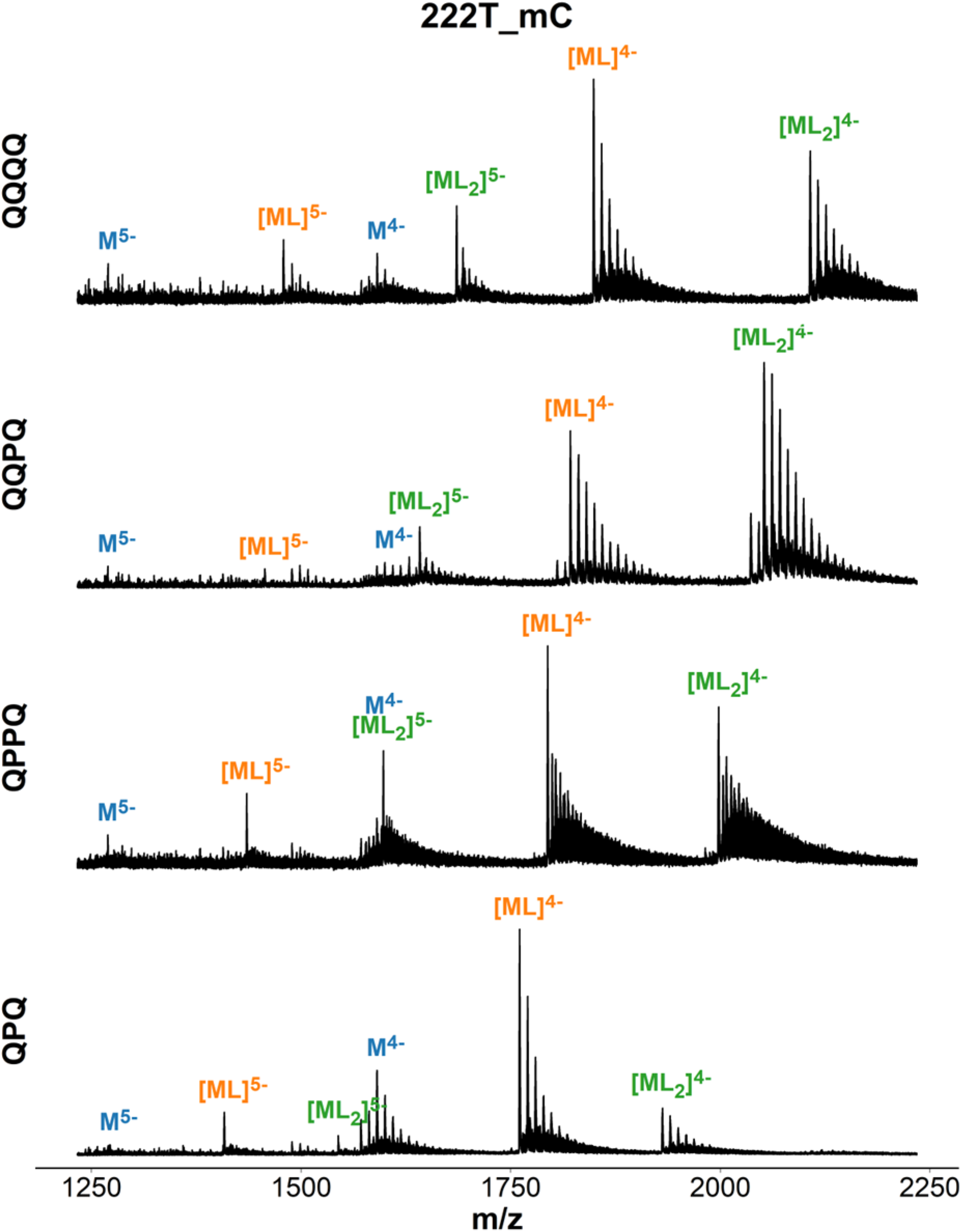
Mass spectra of 222T-mC (dT**GGG**TT**GGG**CC**GGG**TT**GGG**T) in presence of ligand. Samples contain 10 µM DNA, 20 µM ligand, 0.5 mM KCl, 100 mM TMAA (pH 6.8).

**Figure S38.**
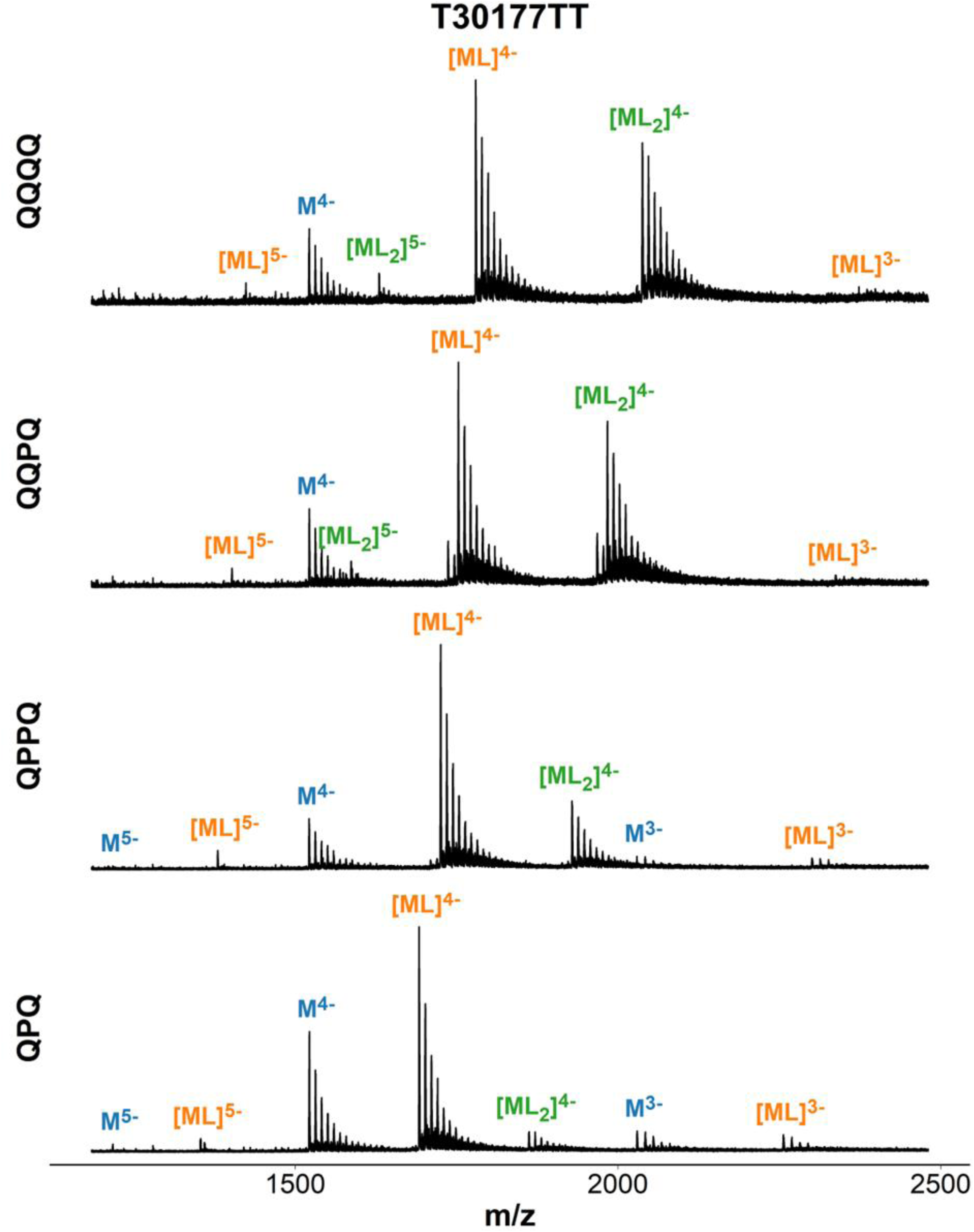
Mass spectra of T30177TT (dTT**G**T**GG**T**GGG**T**GGG**T**GGG**T) in presence of ligand. Samples contain 10 µM DNA, 20 µM ligand, 0.5 mM KCl, 100 mM TMAA (pH 6.8).

**Figure S39.**
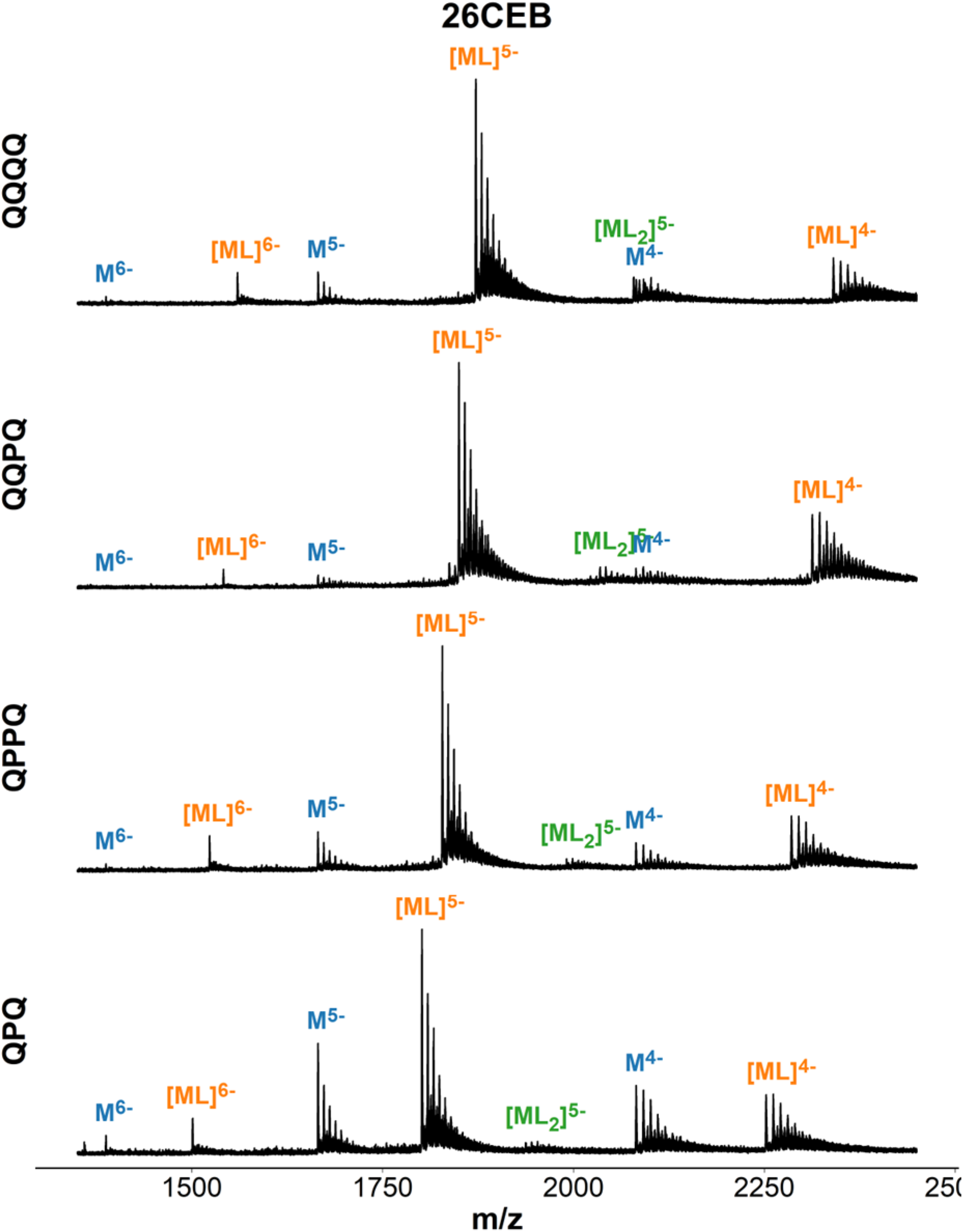
Mass spectra of 26CEB (dAA**GGG**T**GGG**T**G**TAA**G**T**G**T**GGG**T**GGG**T) in presence of ligand. Samples contain 10 µM DNA, 20 µM ligand, 0.5 mM KCl, 100 mM TMAA (pH 6.8).

**Figure S40.**
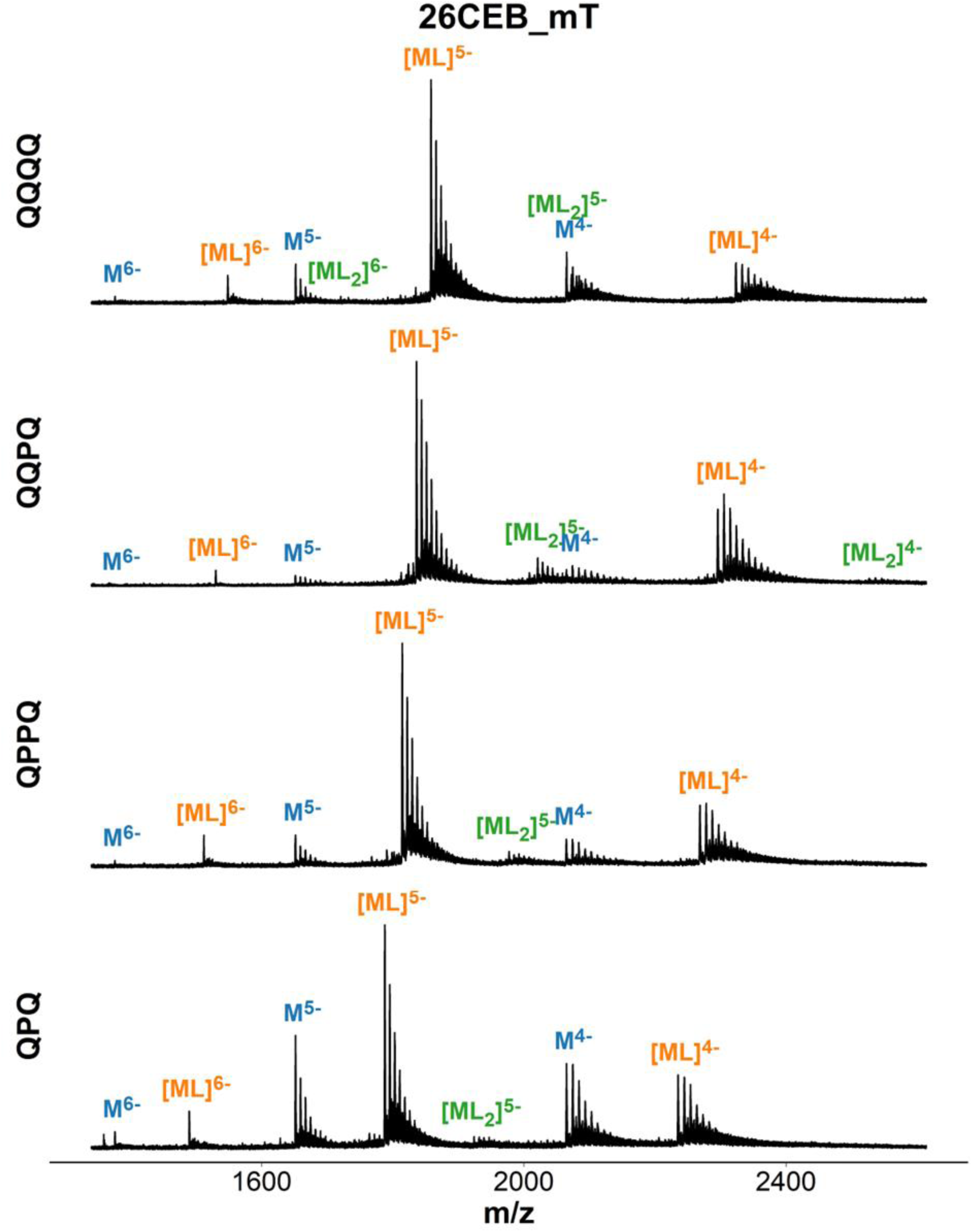
Mass spectra of 26CEB-mT (dAA**GGG**T**GGG**TTTTTTT**G**T**GGG**T**GGG**T) in presence of ligand. Samples contain 10 µM DNA, 20 µM ligand, 0.5 mM KCl, 100 mM TMAA (pH 6.8).

**Figure S41.**
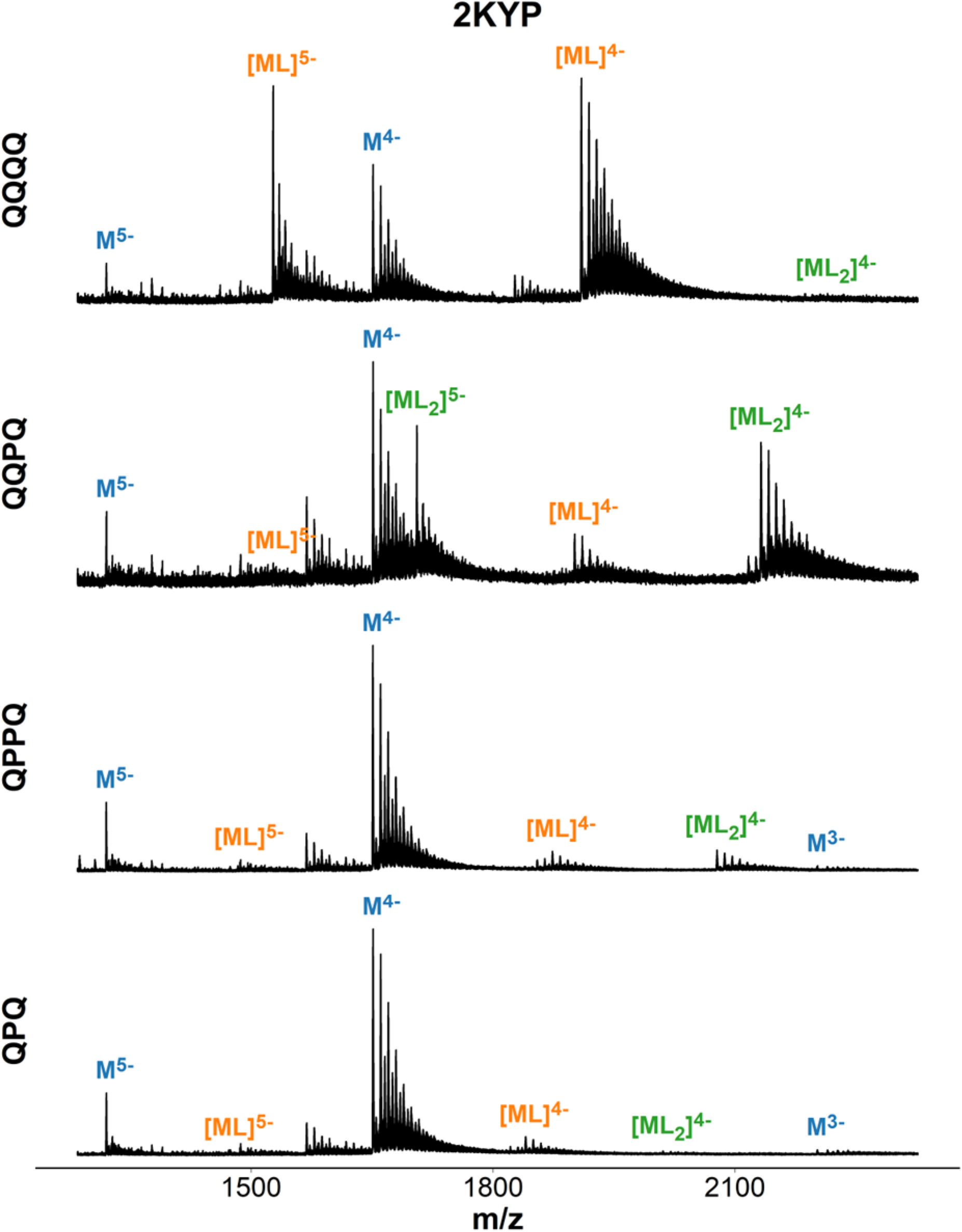
Mass spectra of 2KYP (dC**GGG**C**GGG**C**G**CTA**GGG**A**GGG**T) in presence of ligand. Samples contain 10 µM DNA, 20 µM ligand, 0.5 mM KCl, 100 mM TMAA (pH 6.8).

**Figure S42.**
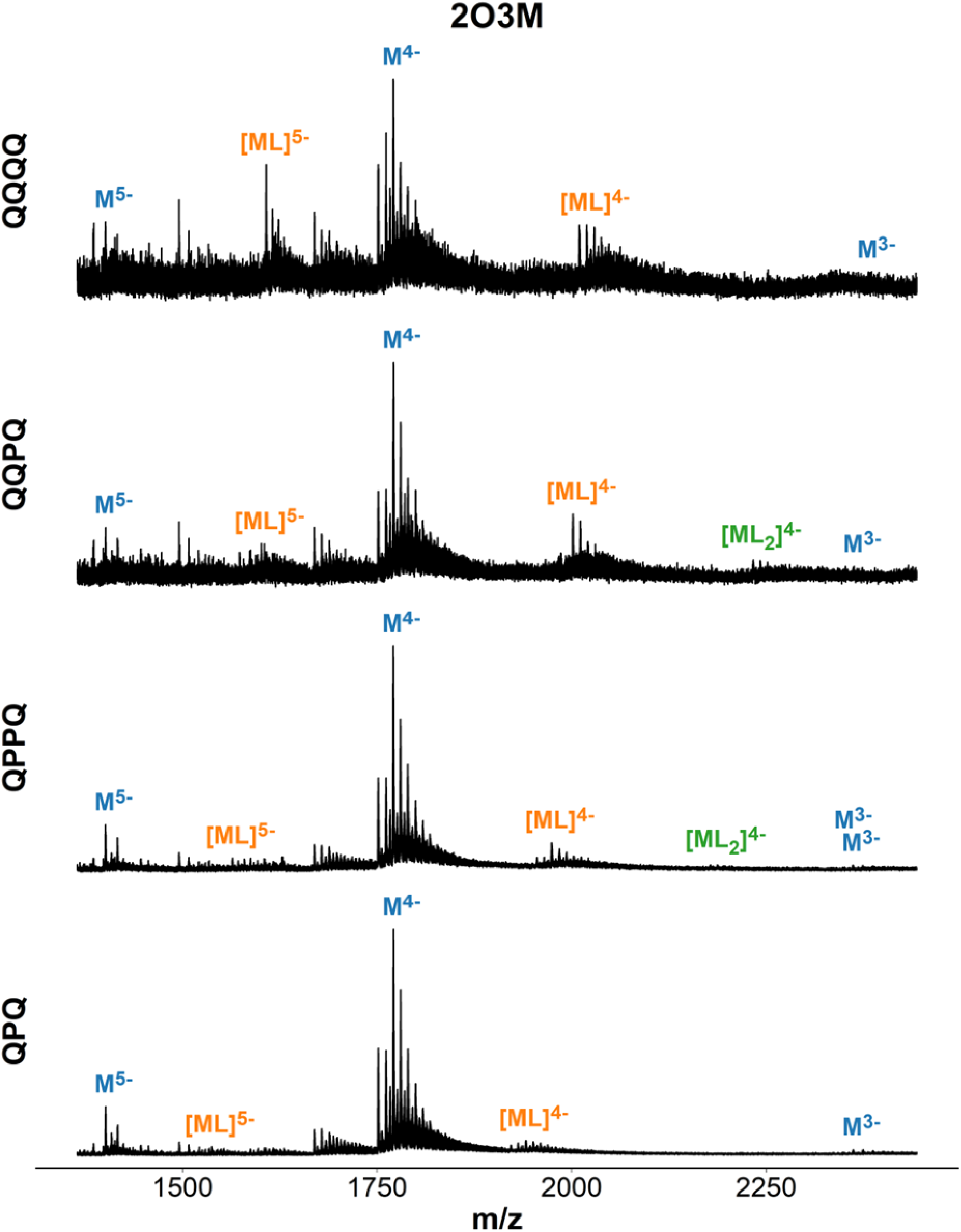
Mass spectra of 2O3M (dA**GGG**A**GGG**C**G**CT**GGG**A**GG**A**GGG**) in presence of ligand. Samples contain 10 µM DNA, 20 µM ligand, 0.5 mM KCl, 100 mM TMAA (pH 6.8).

**Figure S43.**
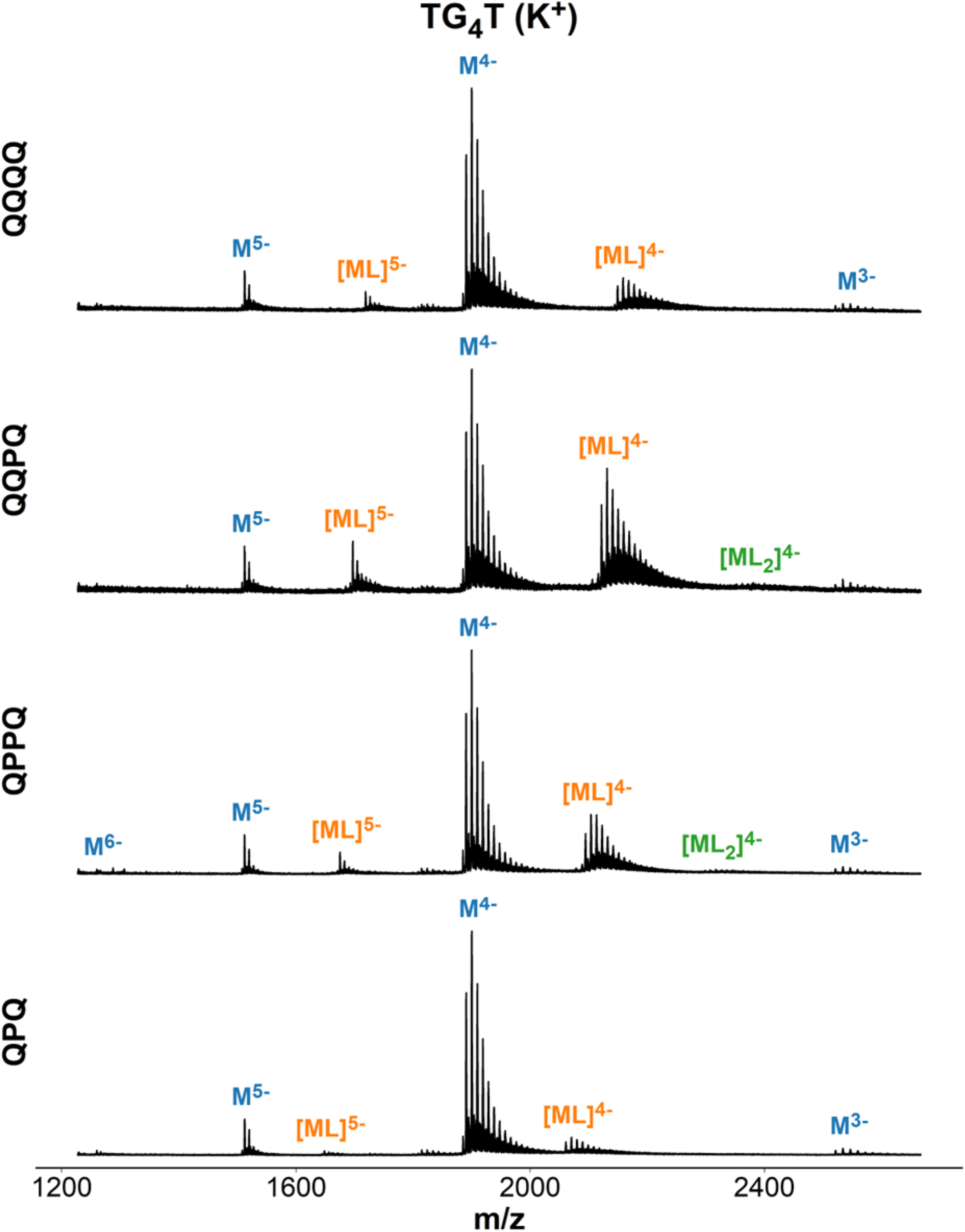
Mass spectra of TG4T ([dT**GGGG**T)_4_) in presence of ligand. Samples contain 40 µM DNA, 20 µM ligand, 0.5 mM KCl, 100 mM TMAA (pH 6.8).

**Figure S44.**
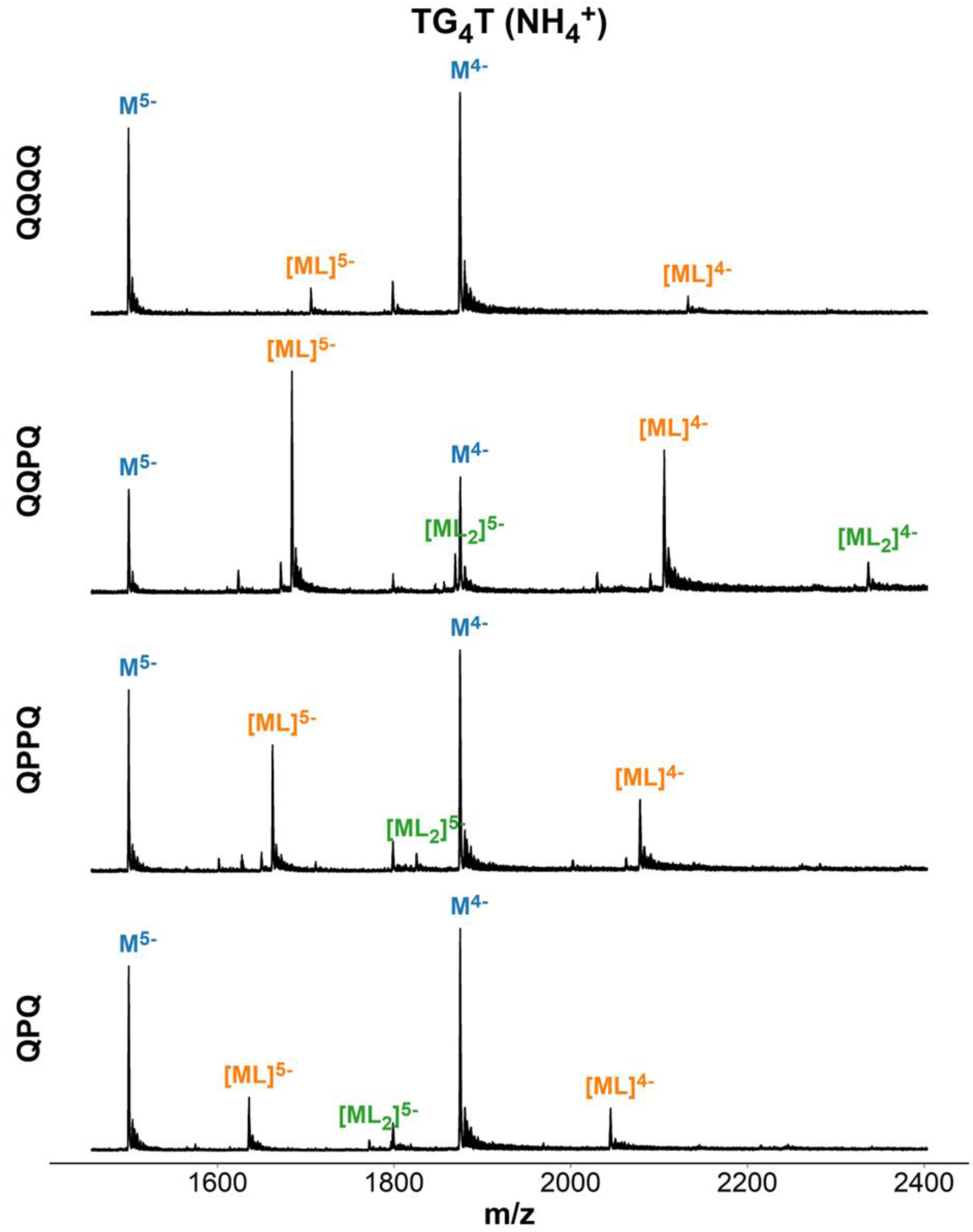
Mass spectra of TG4T ([dT**GGGG**T)_4_) in presence of ligand. Samples contain 40 µM DNA, 20 µM ligand, 150 mM ammonium acetate (pH 6.8).

**Figure S45.**
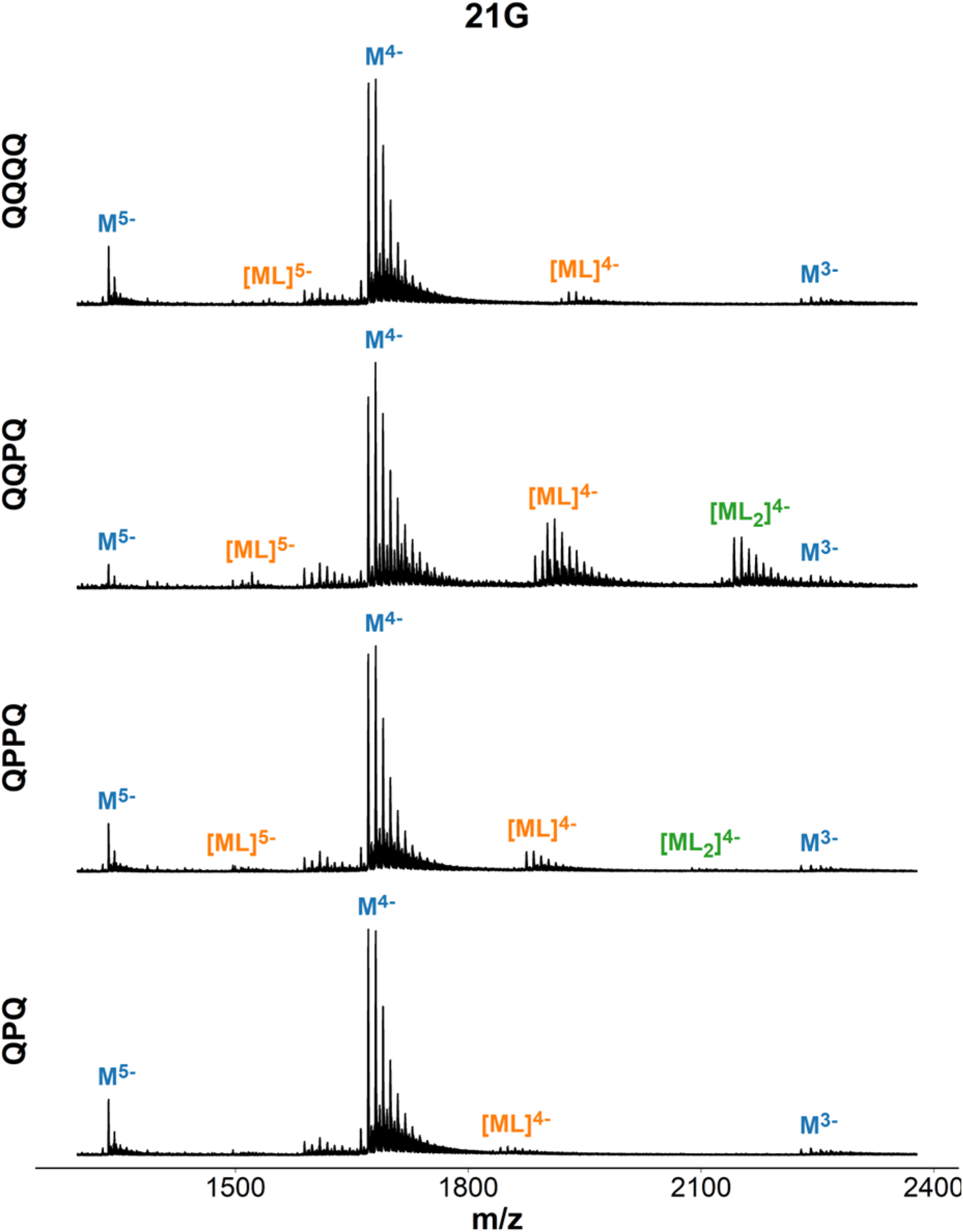
Mass spectra of 21G (d**GGG**TTA**GGG**TTA**GGG**TTA**GGG**) in presence of ligand. Samples contain 10 µM DNA, 20 µM ligand, 0.5 mM KCl, 100 mM TMAA (pH 6.8).

**Figure S46.**
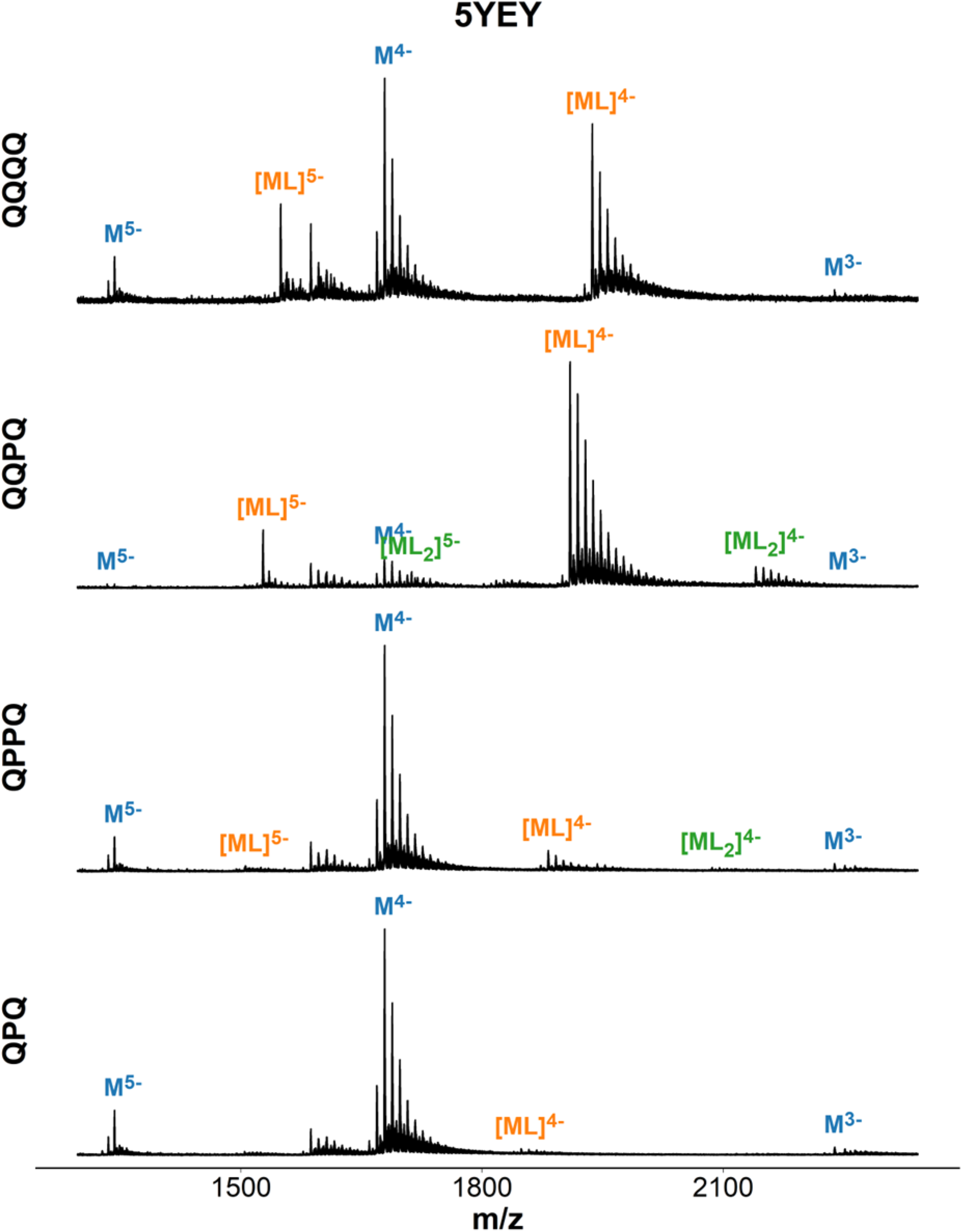
Mass spectra of 5YEY (d**GGG**TTA**GGG**TTA**GGG**TTT**GGG**) in presence of ligand. Samples contain 10 µM DNA, 20 µM ligand, 0.5 mM KCl, 100 mM TMAA (pH 6.8).

**Figure S47.**
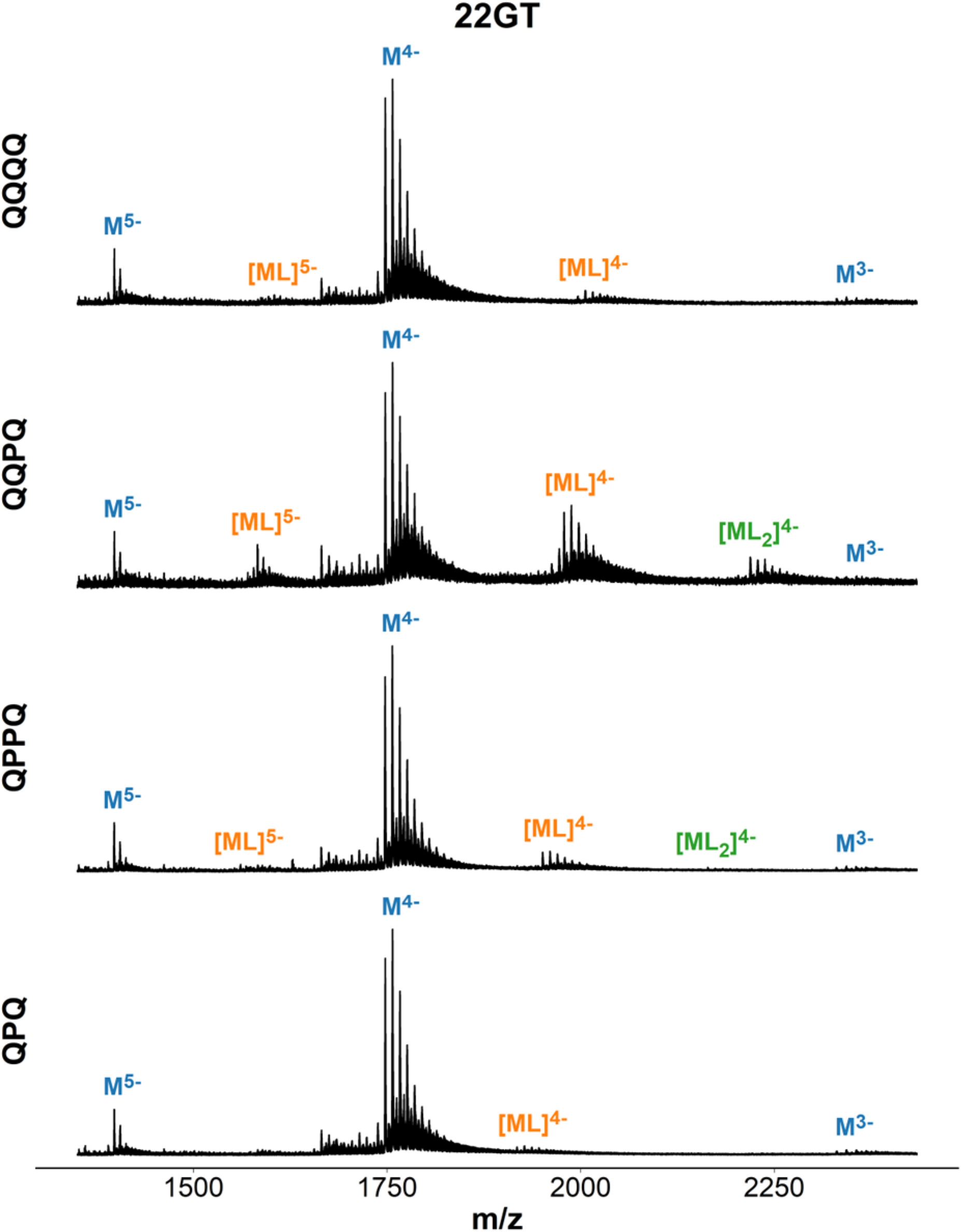
Mass spectra of 22GT (d**GGG**TTA**GGG**TTA**GGG**TTA**GGG**T) in presence of ligand. Samples contain 10 µM DNA, 20 µM ligand, 0.5 mM KCl, 100 mM TMAA (pH 6.8).

**Figure S48.**
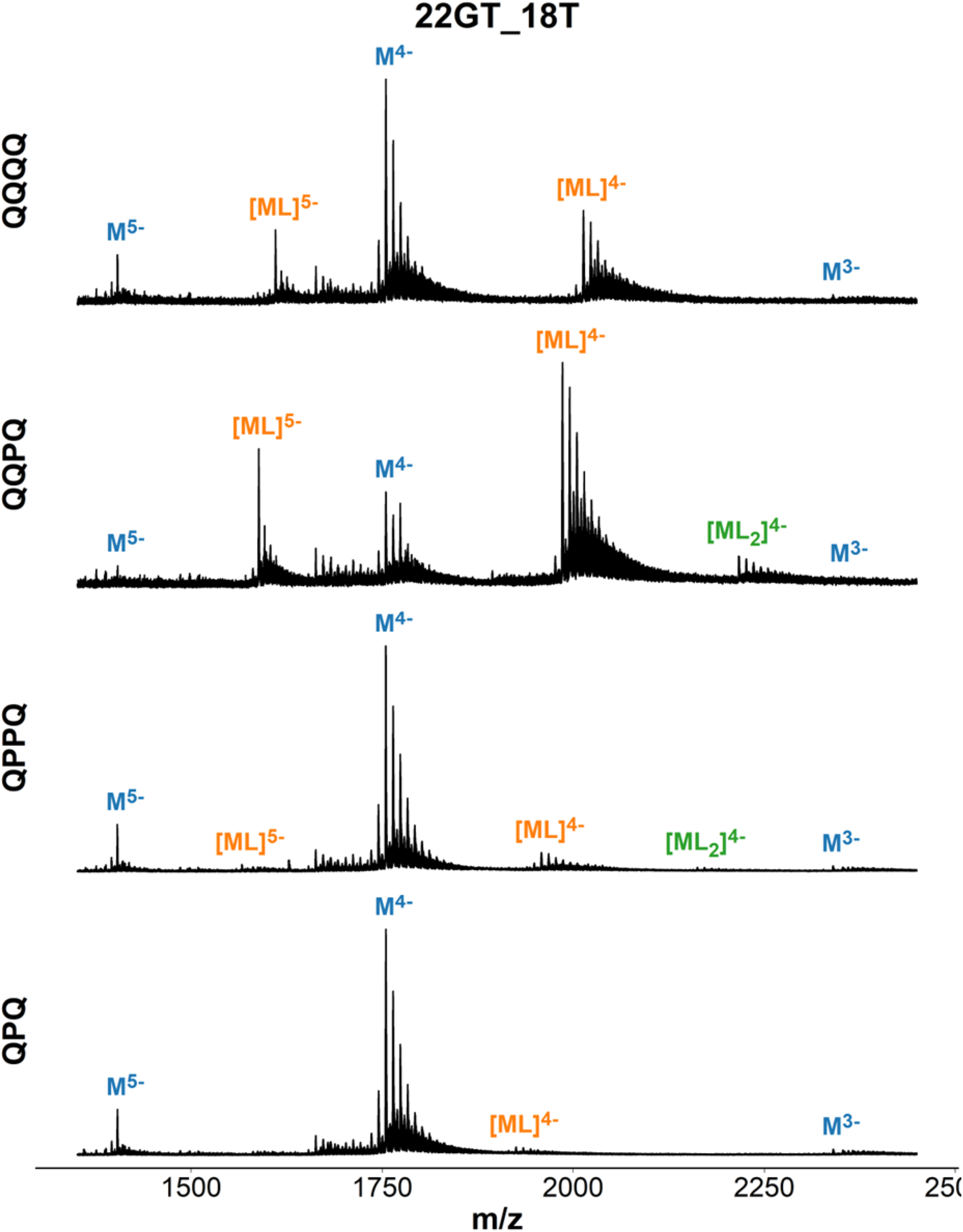
Mass spectra of 22GT-18T (d**GGG**TTA**GGG**TTA**GGG**TTT**GGG**T) in presence of ligand. Samples contain 10 µM DNA, 20 µM ligand, 0.5 mM KCl, 100 mM TMAA (pH 6.8).

**Figure S49.**
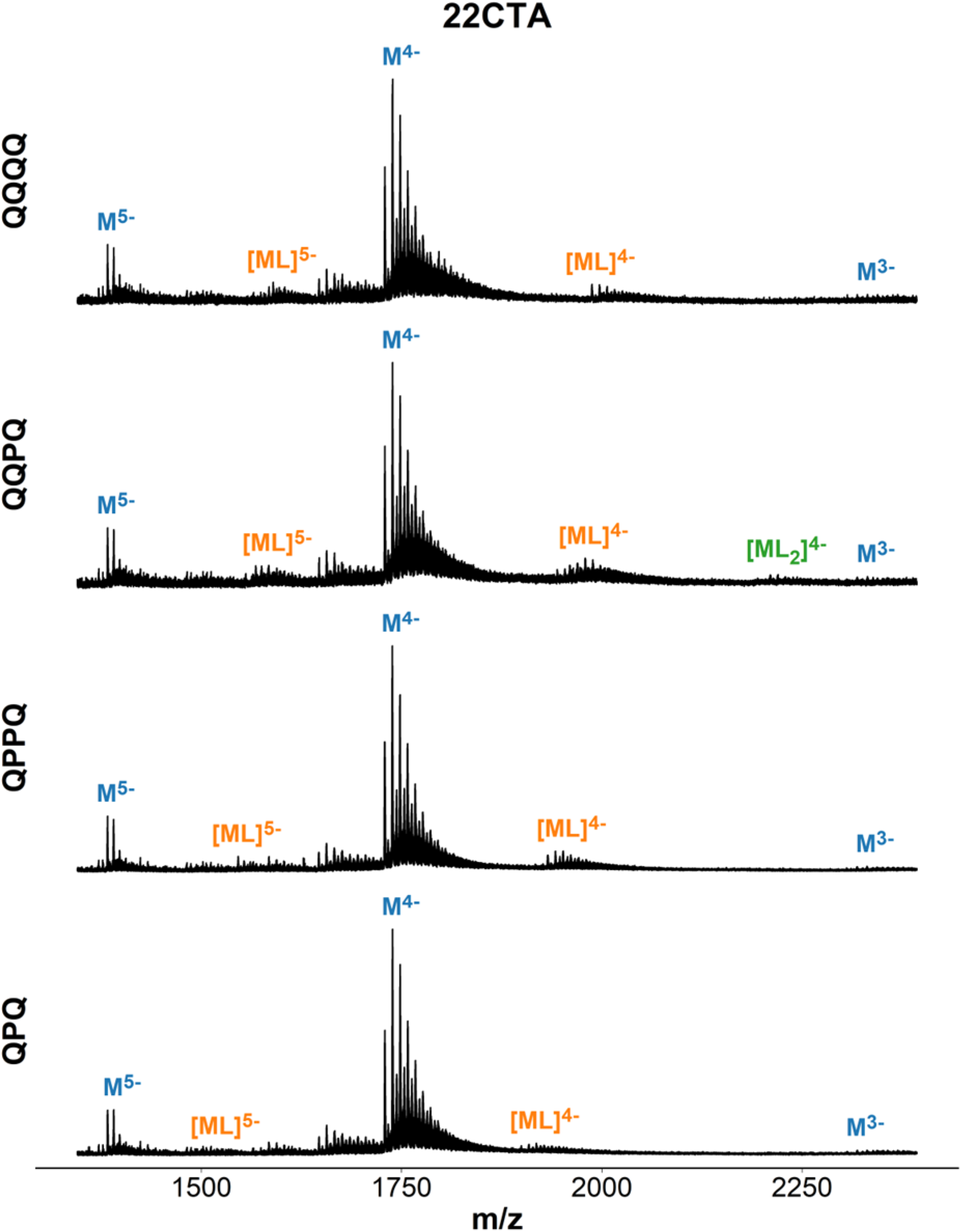
Mass spectra of 22CTA (dA**GGG**CTA**GGG**CTA**GGG**CTA**GGG**) in presence of ligand. Samples contain 10 µM DNA, 20 µM ligand, 0.5 mM KCl, 100 mM TMAA (pH 6.8).

**Figure S50.**
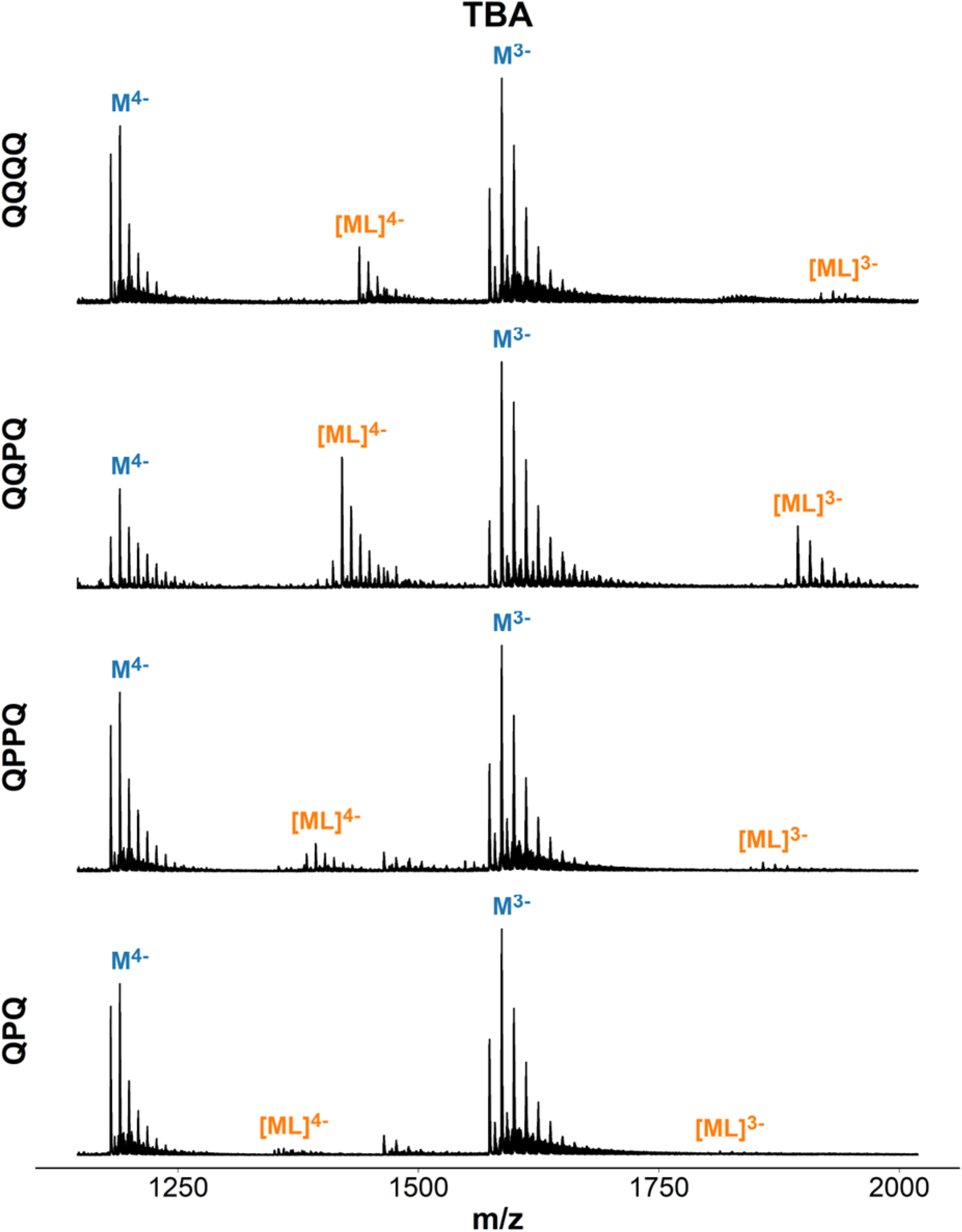
Mass spectra of TBA (d**GG**TT**GG**T**G**T**GG**TT**GG**) in presence of ligand. Samples contain 10 µM DNA, 20 µM ligand, 0.5 mM KCl, 100 mM TMAA (pH 6.8).

**Figure S51.**
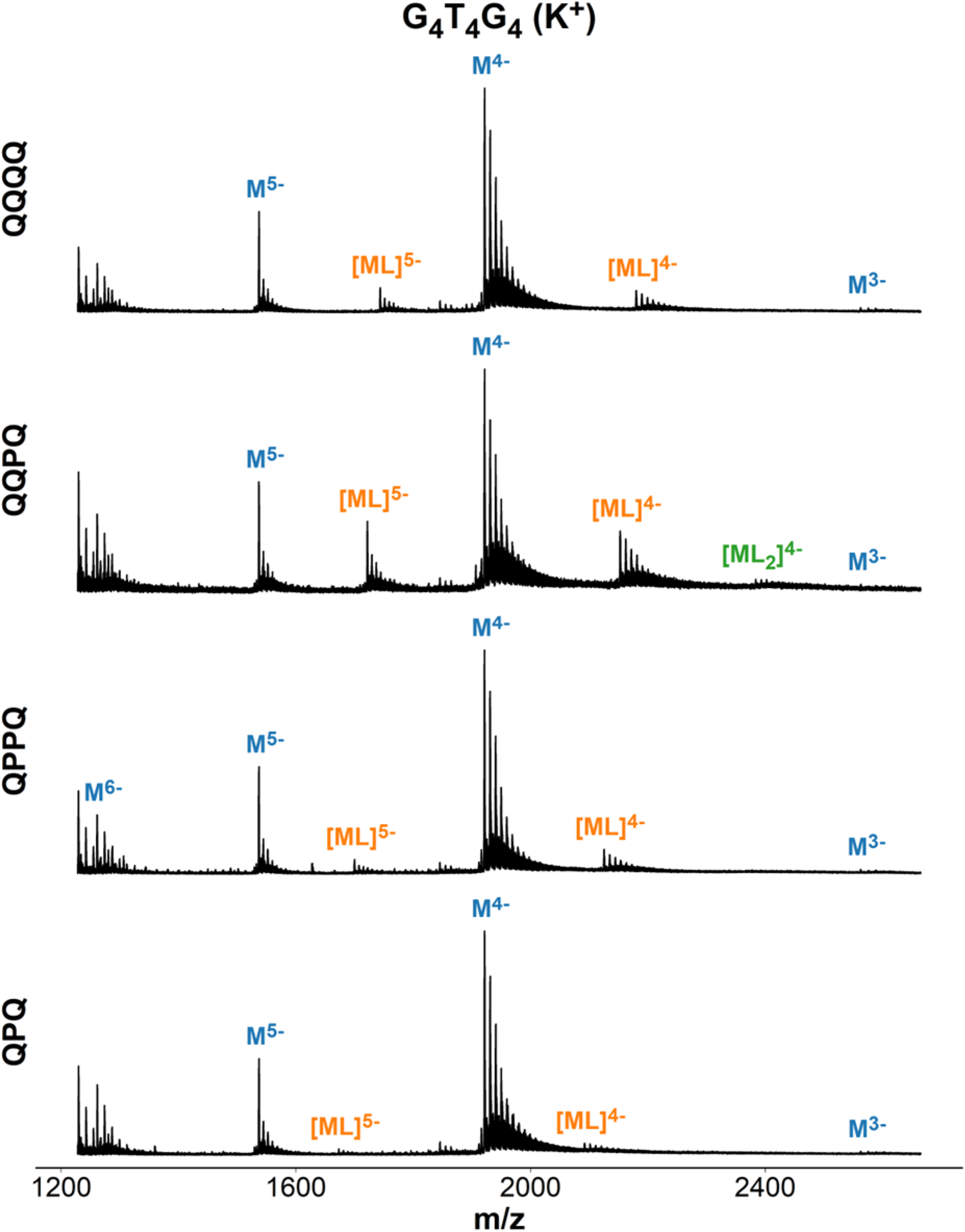
Mass spectra of G4T4G4 ([d**GGGG**TTTT**GGGG**]_2_) in presence of ligand. Samples contain 20 µM DNA, 20 µM ligand, 0.5 mM KCl, 100 mM TMAA (pH 6.8).

**Figure S52.**
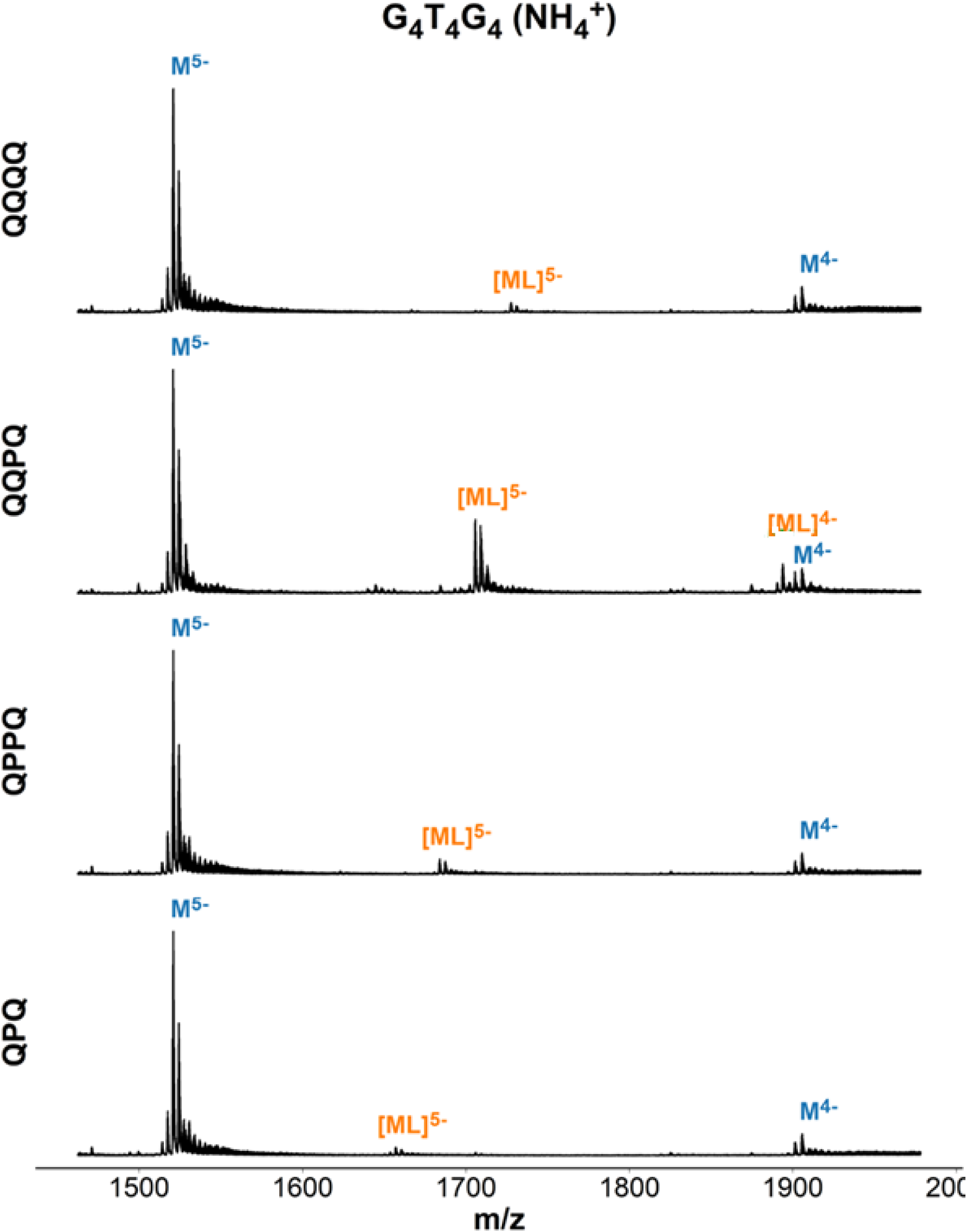
Mass spectra of G4T4G4 ([d**GGGG**TTTT**GGGG**]_2_) in presence of ligand. Samples contain 20 µM DNA, 20 µM ligand, 150 mM ammonium acetate (pH 6.8).

**Figure S53.**
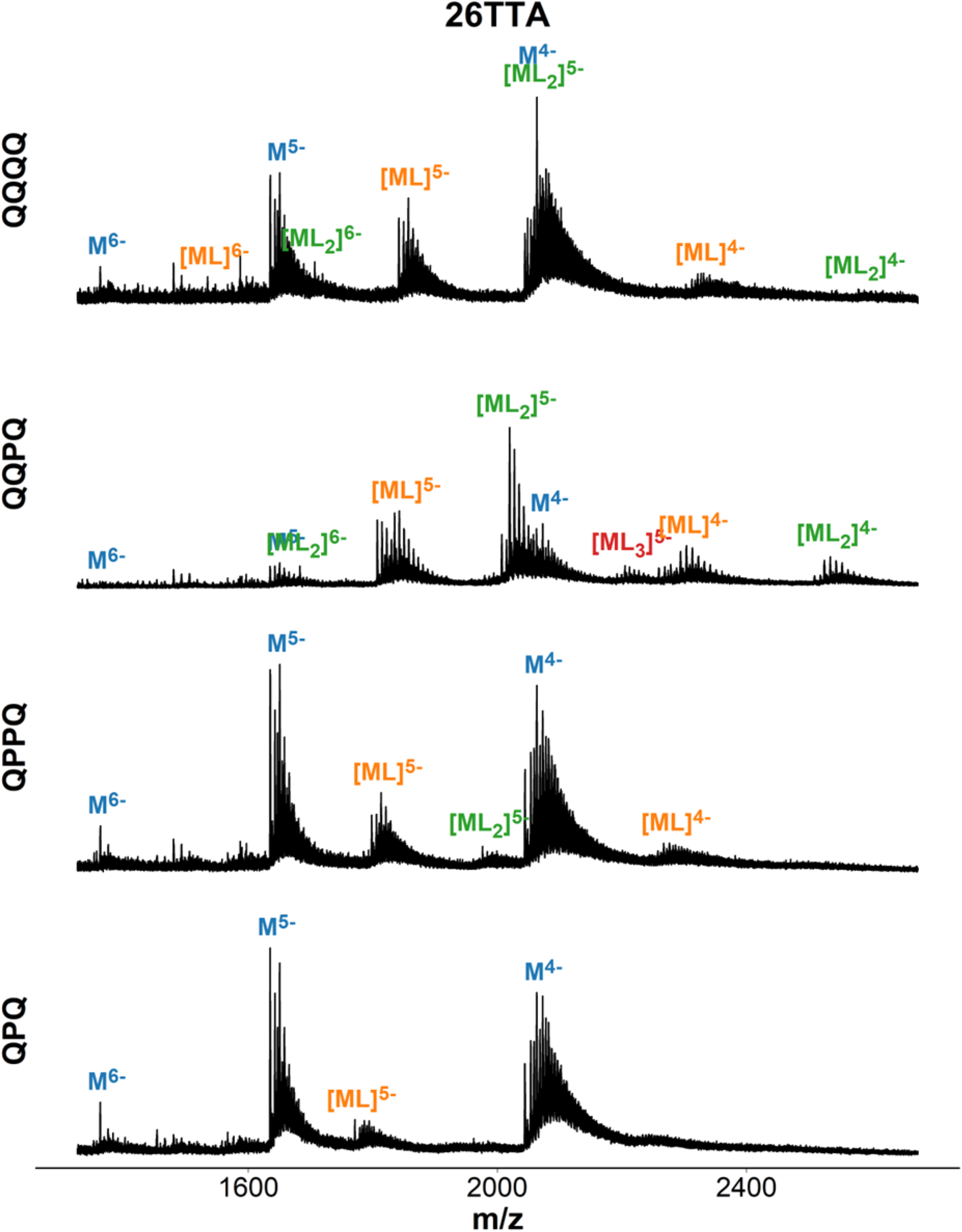
Mass spectra of 26TTA (dTTA**GGG**TTA**GGG**TTA**GGG**TTA**GGG**TT) in presence of ligand. Samples contain 10 µM DNA, 20 µM ligand, 0.5 mM KCl, 100 mM TMAA (pH 6.8).

**Figure S54.**
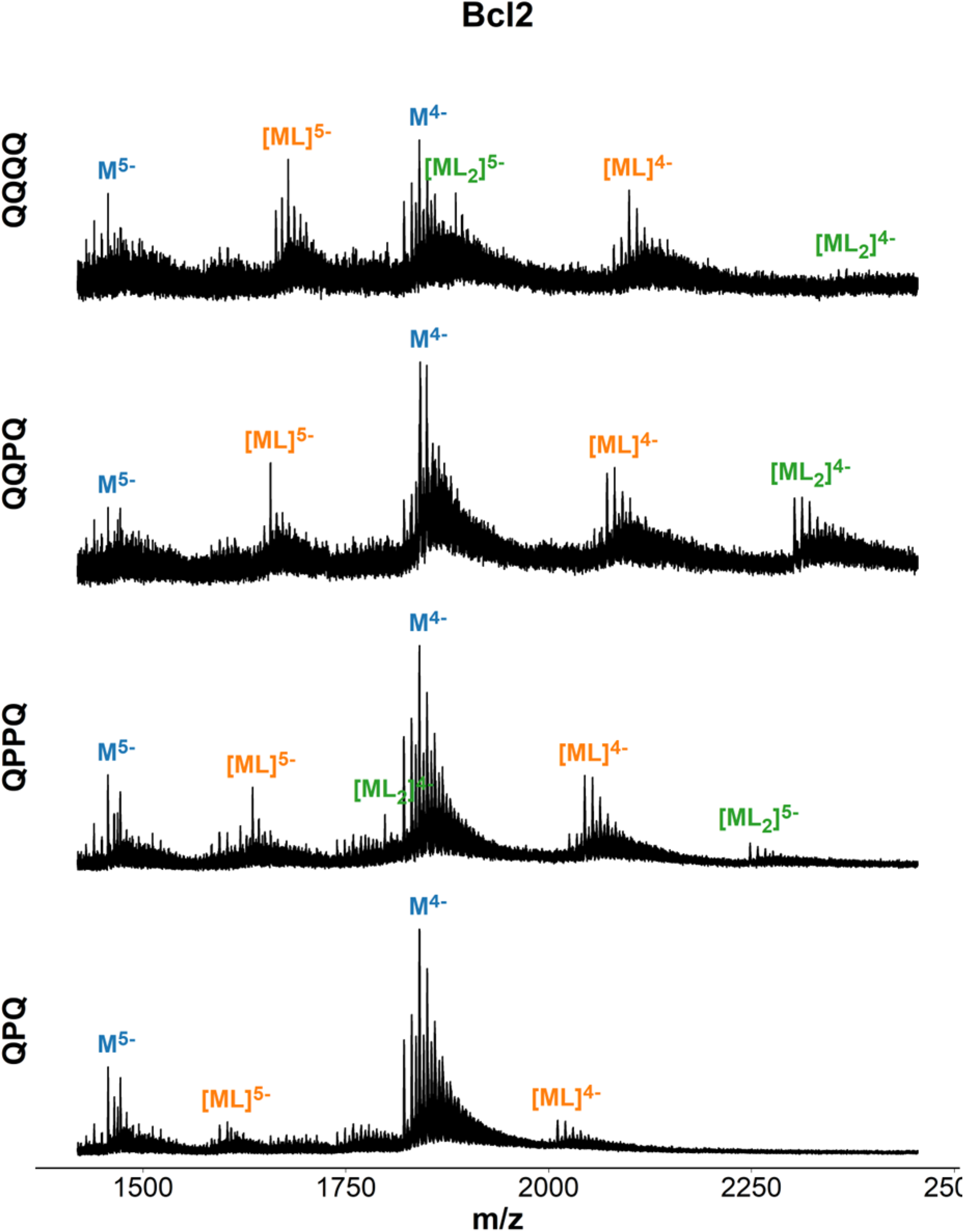
Mass spectra of bcl2 (d**GGG**C**G**C**GGG**A**GG**AATT**GGG**C**GGG**) in presence of ligand. Samples contain 10 µM DNA, 20 µM ligand, 0.5 mM KCl, 100 mM TMAA (pH 6.8).

**Figure S55.**
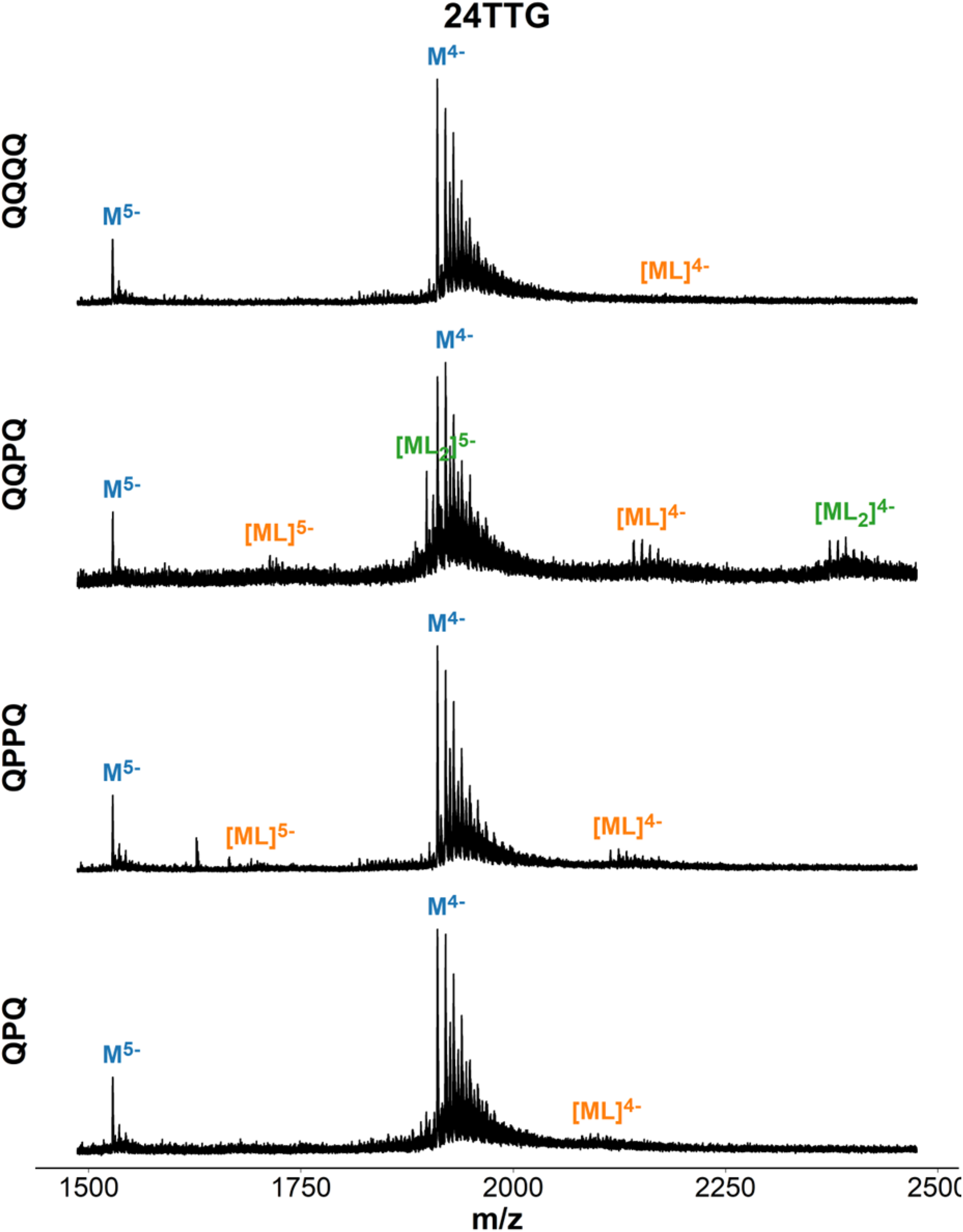
Mass spectra of 24TTG (dTT**GGG**TTA**GGG**TTA**GGG**TTA**GGG**A) in presence of ligand. Samples contain 10 µM DNA, 20 µM ligand, 0.5 mM KCl, 100 mM TMAA (pH 6.8).

**Figure S56.**
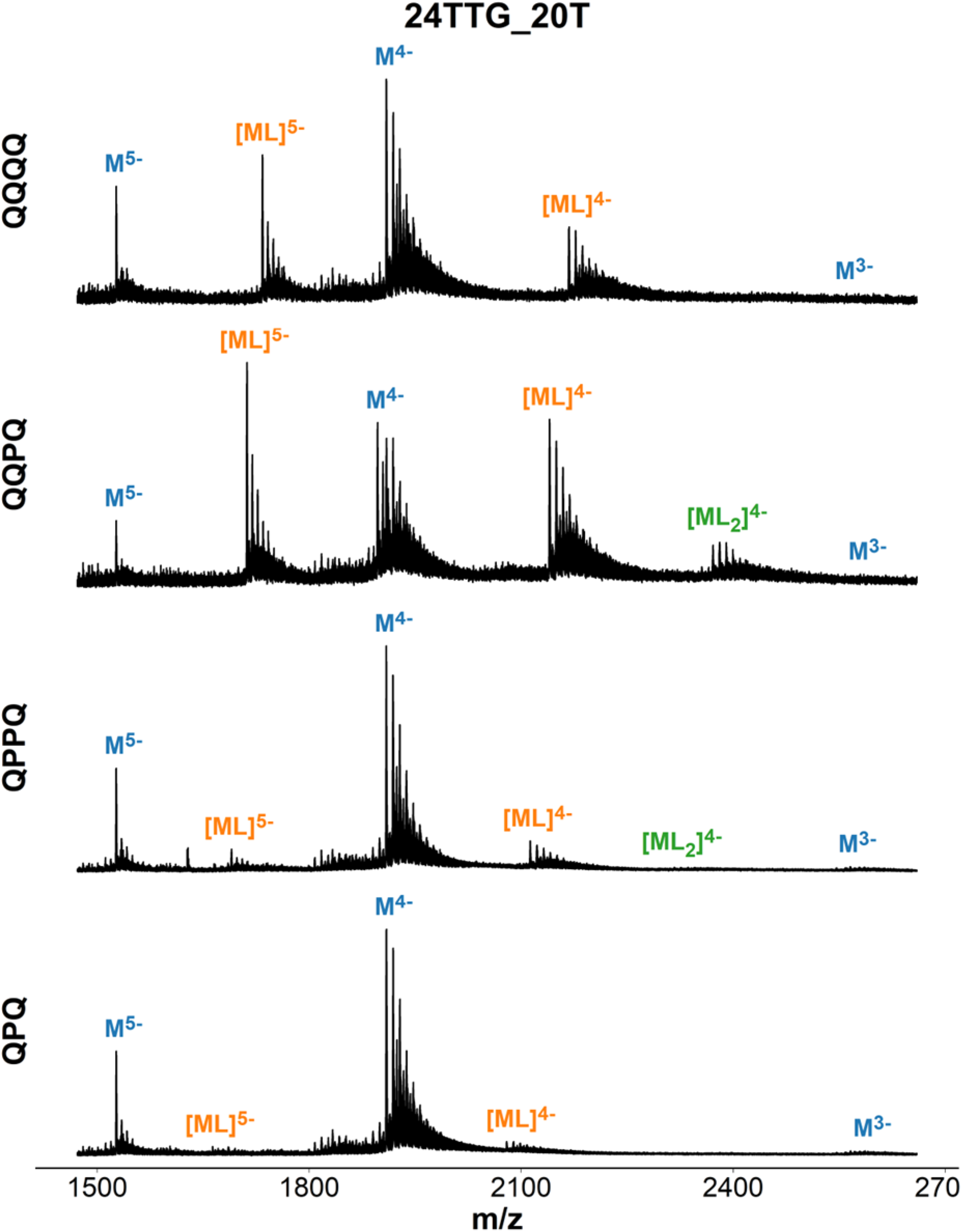
Mass spectra of 24TTG-20T (dTT**GGG**TTA**GGG**TTA**GGG**TTT**GGG**A) in presence of ligand. Samples contain 10 µM DNA, 20 µM ligand, 0.5 mM KCl, 100 mM TMAA (pH 6.8).

**Figure S57.**
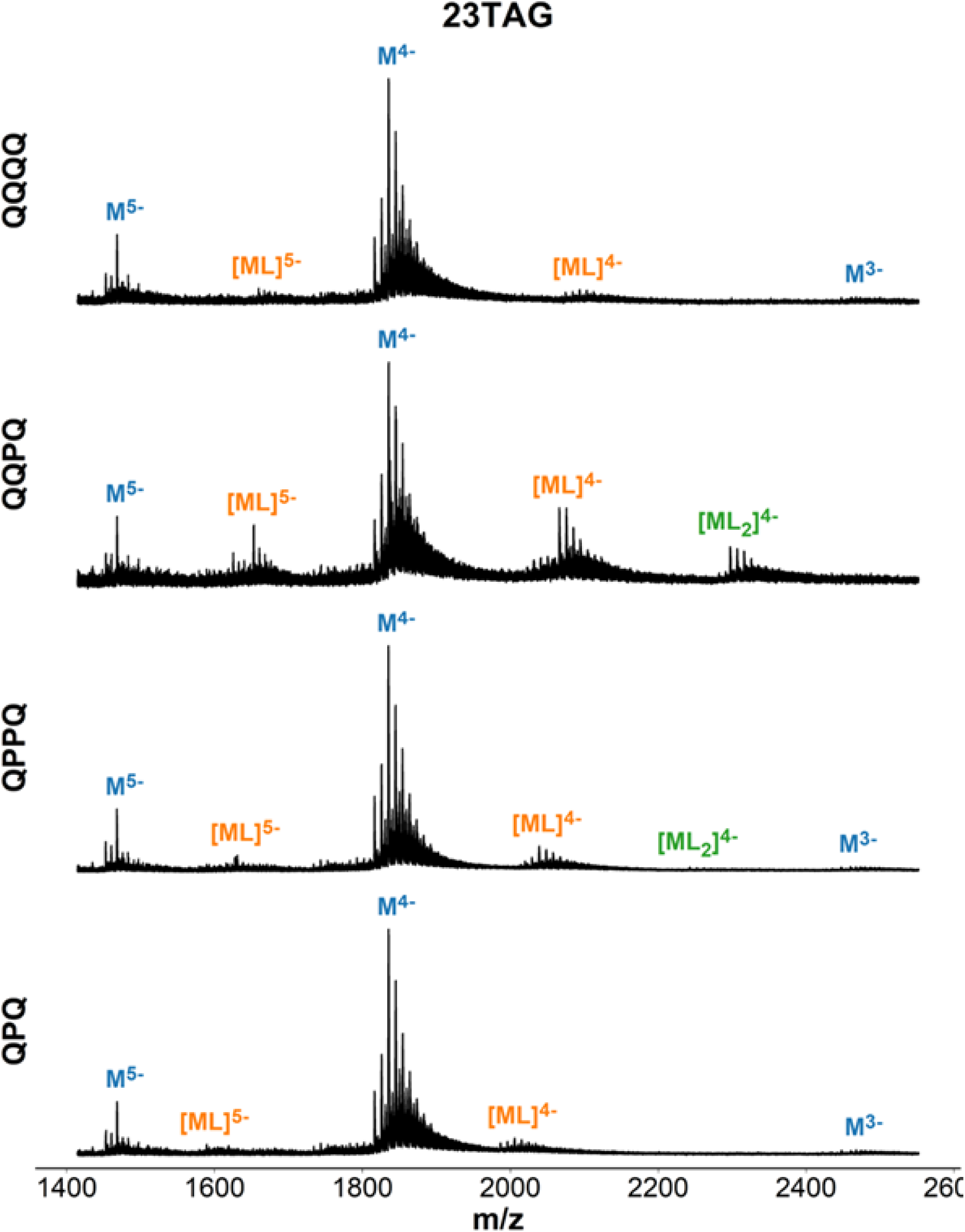
Mass spectra of 23TAG (dTA**GGG**TTA**GGG**TTA**GGG**TTA**GGG**) in presence of ligand. Samples contain 10 µM DNA, 20 µM ligand, 0.5 mM KCl, 100 mM TMAA (pH 6.8).

**Figure S58.**
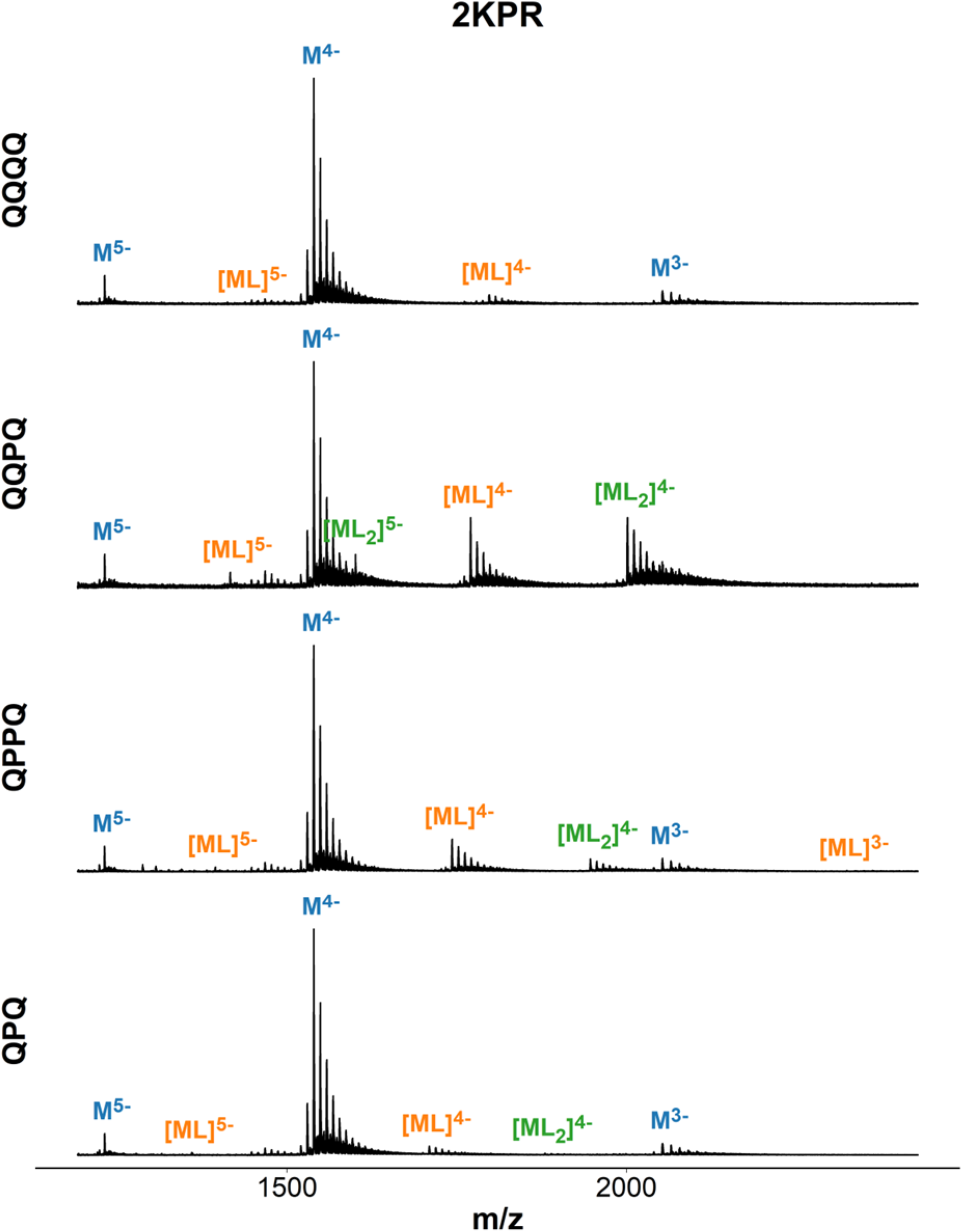
Mass spectra of 2KPR (d**GGG**T**GGGG**AA**GGGG**T**GGG**T) in presence of ligand. Samples contain 10 µM DNA, 20 µM ligand, 0.5 mM KCl, 100 mM TMAA (pH 6.8).

**Figure S59.**
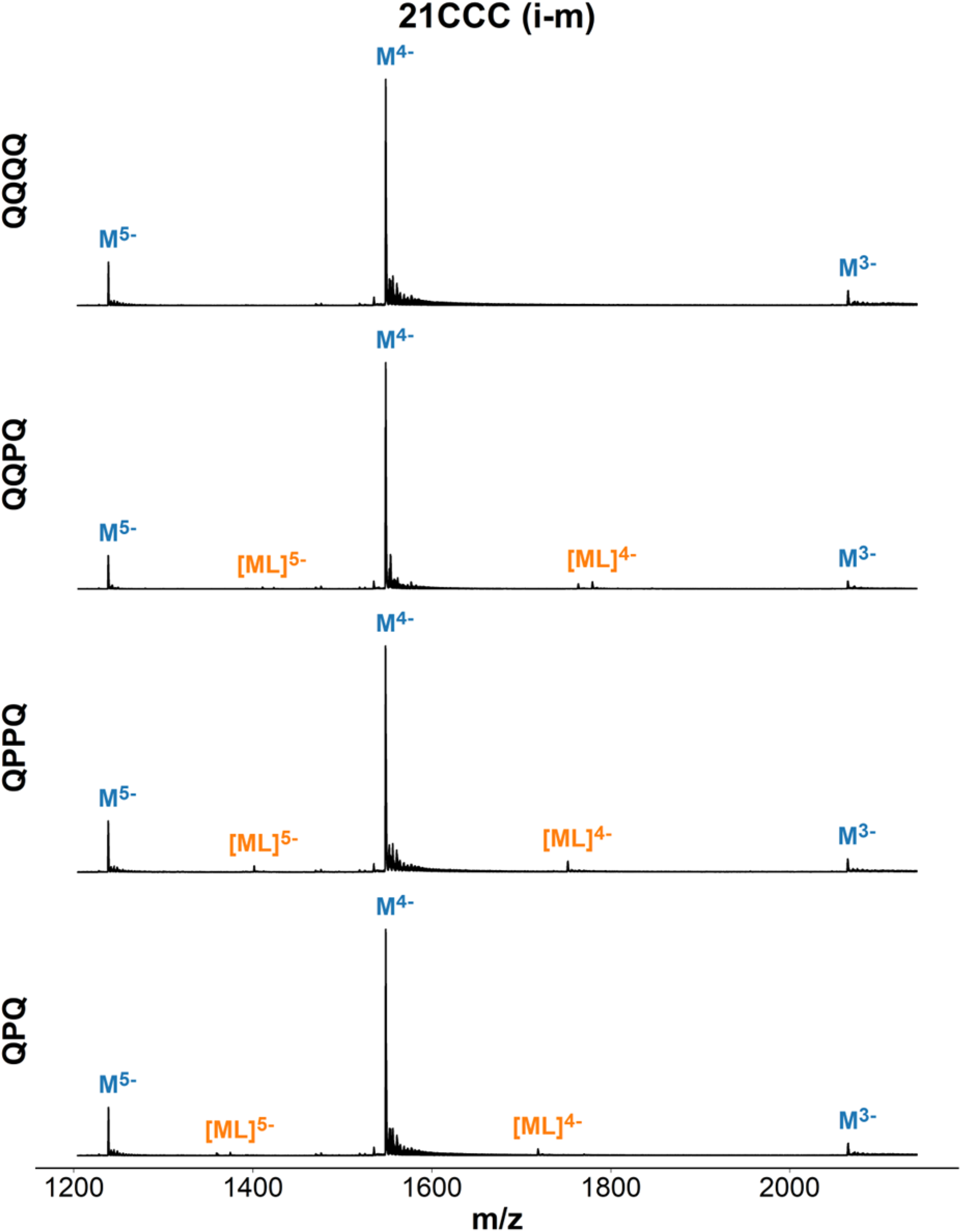
Mass spectra of 21CCC (d**CCC**TAA**CCC**TAA**CCC**TAA**CCC**) in presence of ligand. Samples contain 10 µM DNA, 20 µM ligand, 150 mM ammonium acetate (<u>pH 5.5</u>).

**Figure S60.**
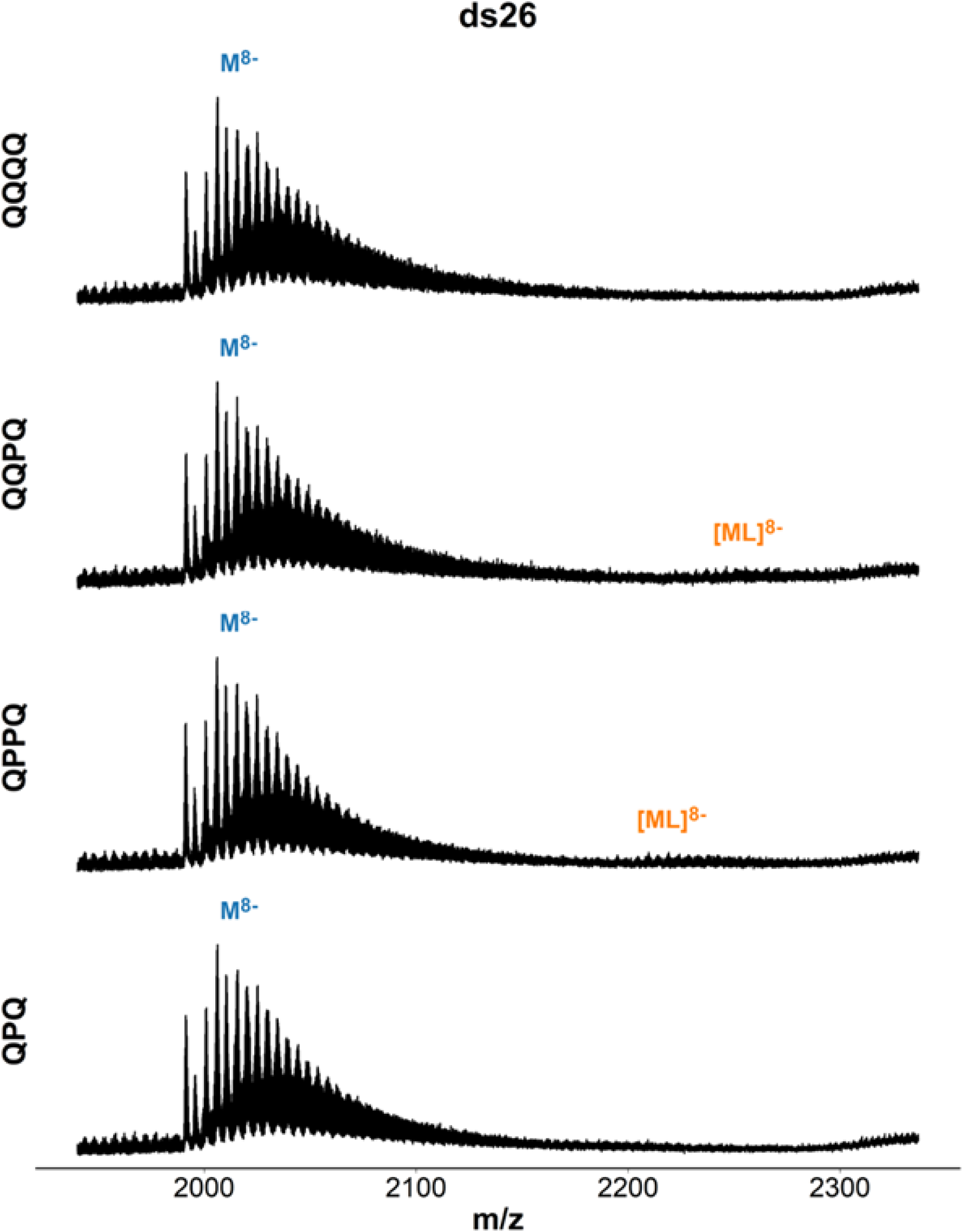
Mass spectra of ds26 ([dCAATCGGATCGAATTCGATCCGATTG]_2_) in presence of ligand. Samples contain 20 µM DNA, 20 µM ligand, 0.5 mM KCl, 100 mM TMAA (pH 6.8).

**Figure S61.**
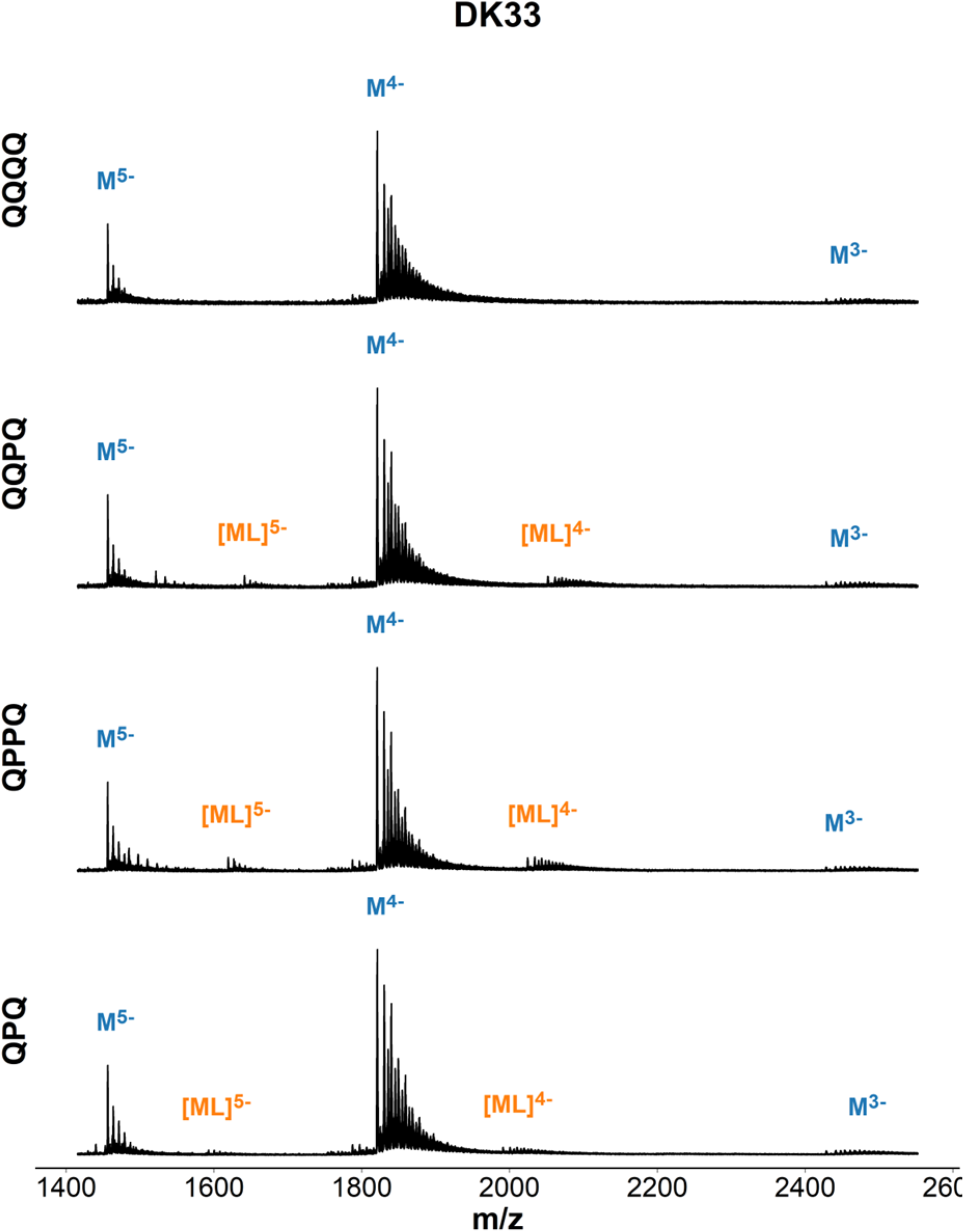
Mass spectra of DK-33 ([dCGTAAATTTACG]_2_) in presence of ligand. Samples contain 20 µM DNA, 20 µM ligand, 0.5 mM KCl, 100 mM TMAA (pH 6.8).

**Figure S62.**
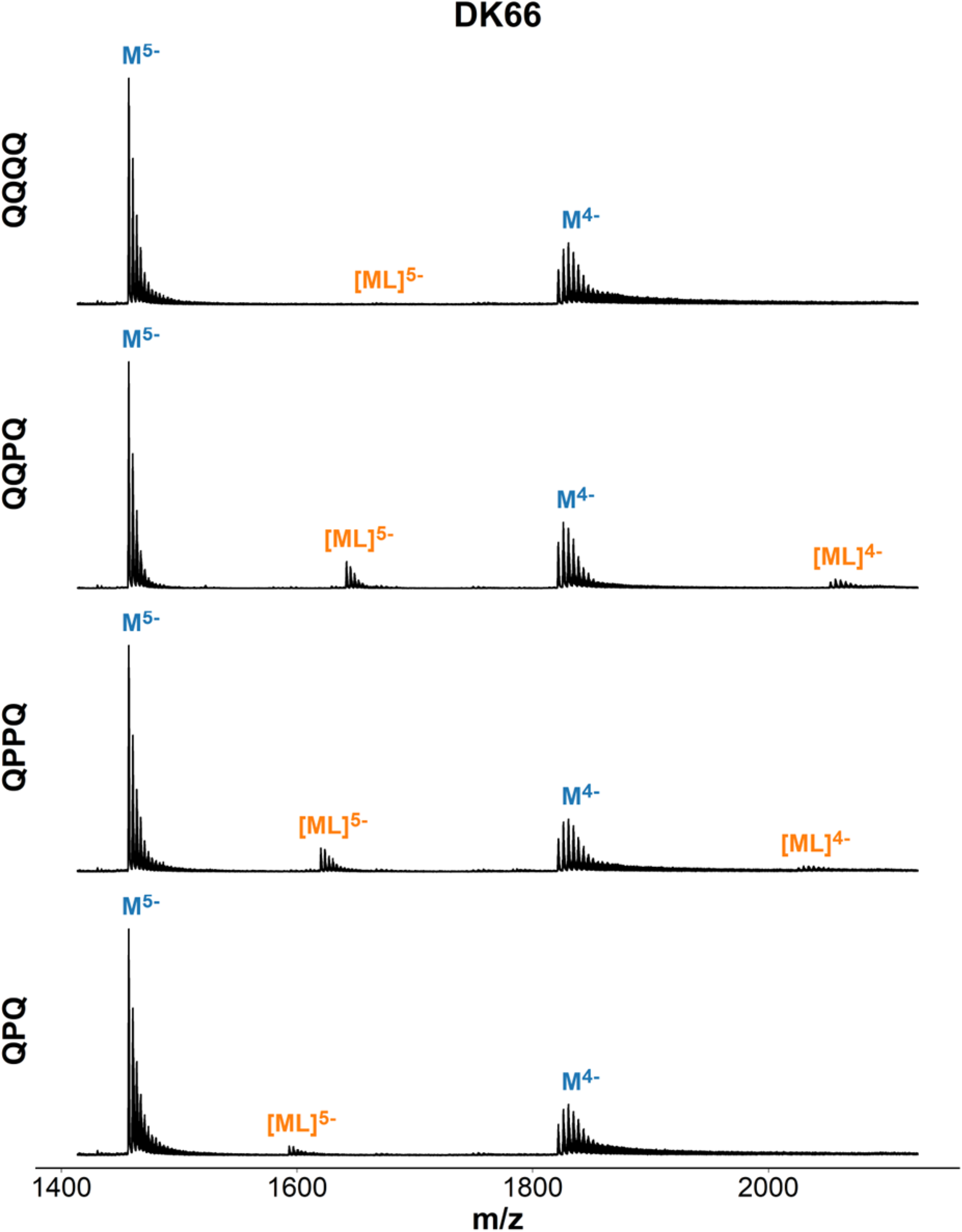
Mass spectra of DK-66 ([dCGCGAATTCGCG]_2_) in presence of ligand. Samples contain 20 µM DNA, 20 µM ligand, 150 mM ammonium acetate (pH 6.8).

**Figure S63.**
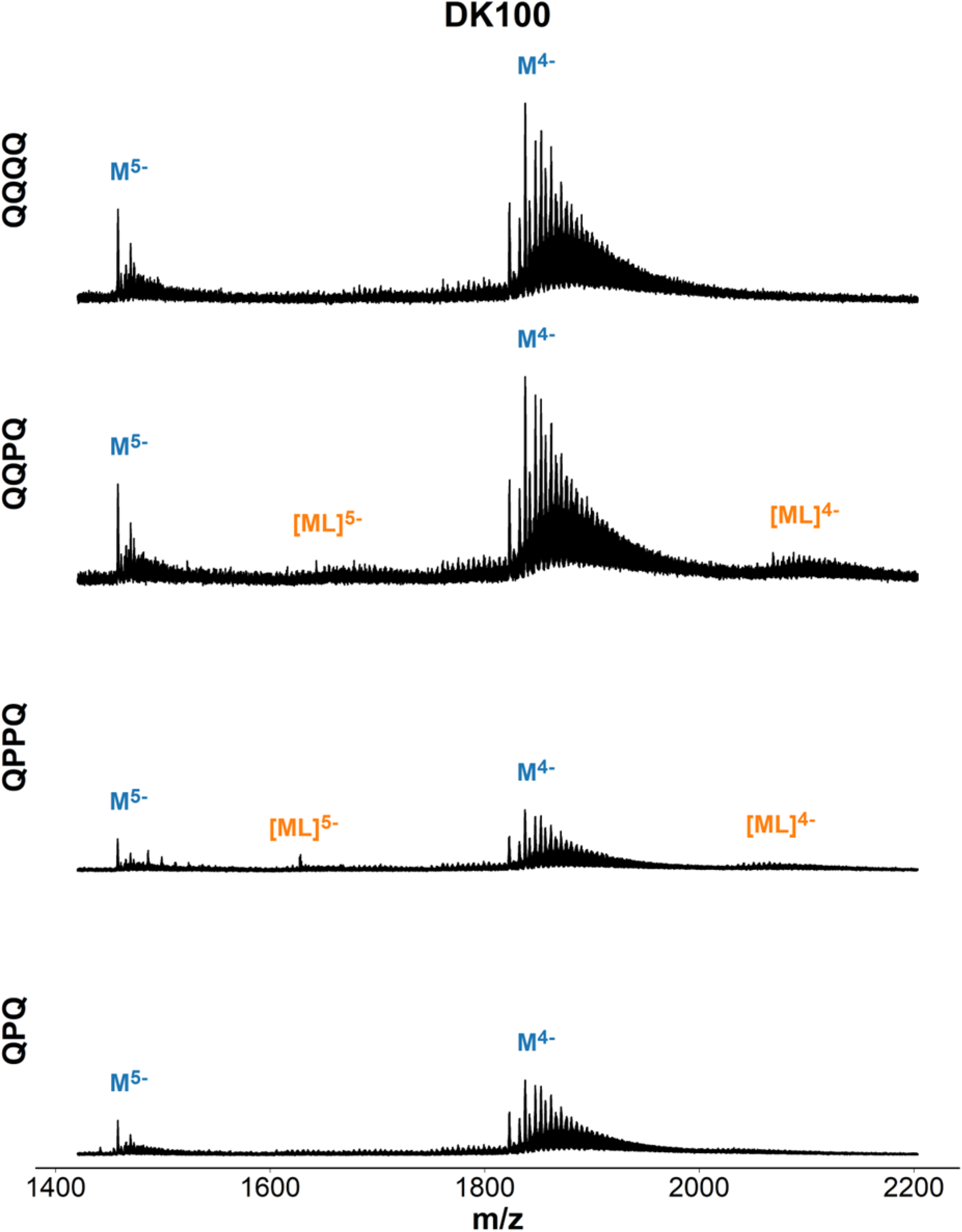
Mass spectra of DK-100 ([dCGCGGGCCCGCG]_2_) in presence of ligand. Samples contain 20 µM DNA, 20 µM ligand, 0.5 mM KCl, 100 mM TMAA (pH 6.8).

**Figure S64.**
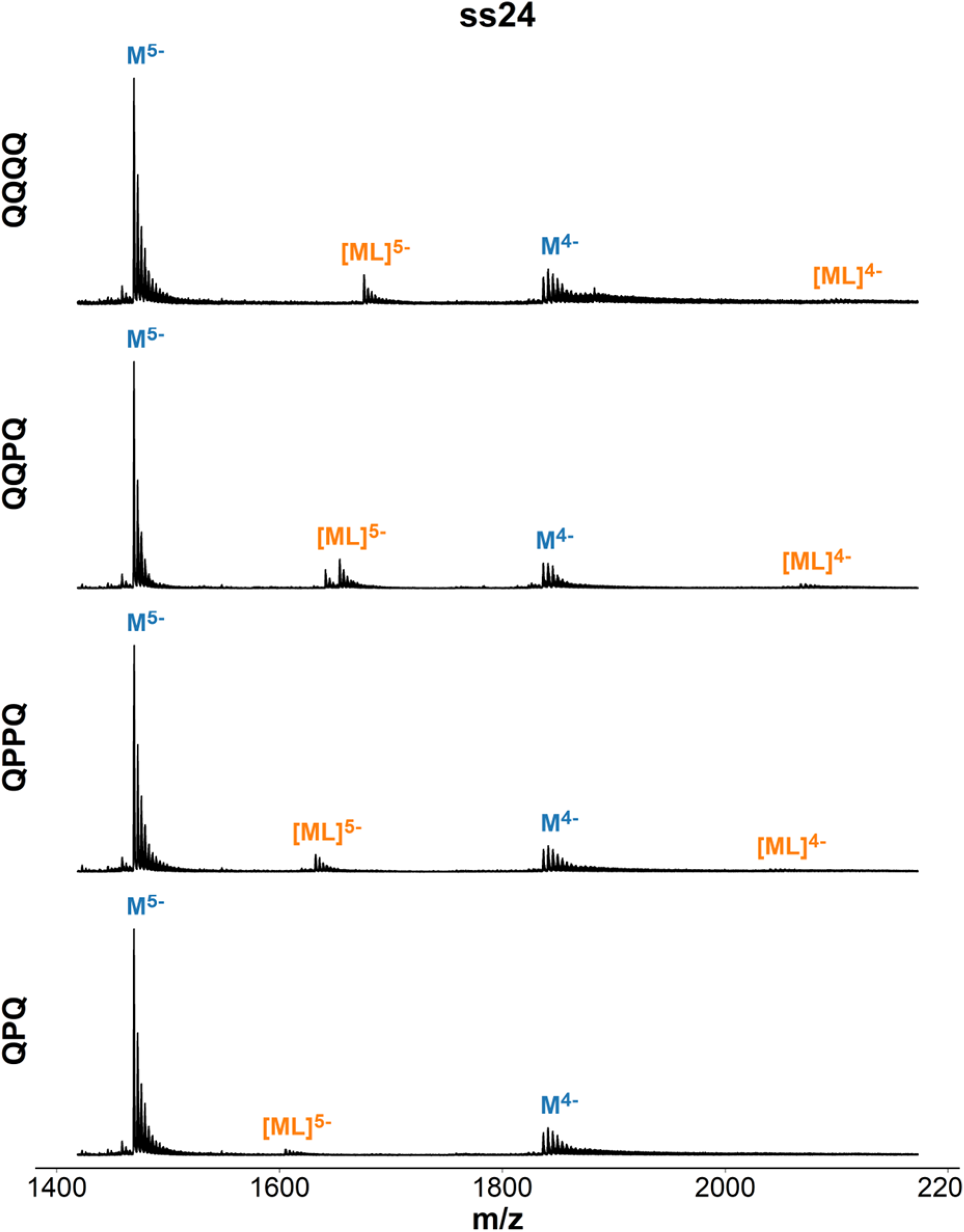
Mass spectra of ss24 (dTGCCATGCTACTGAGATGACGCTA) in presence of ligand. Samples contain 10 µM DNA, 20 µM ligand, 150 mM ammonium acetate (pH 6.8).

**Figure S65.**
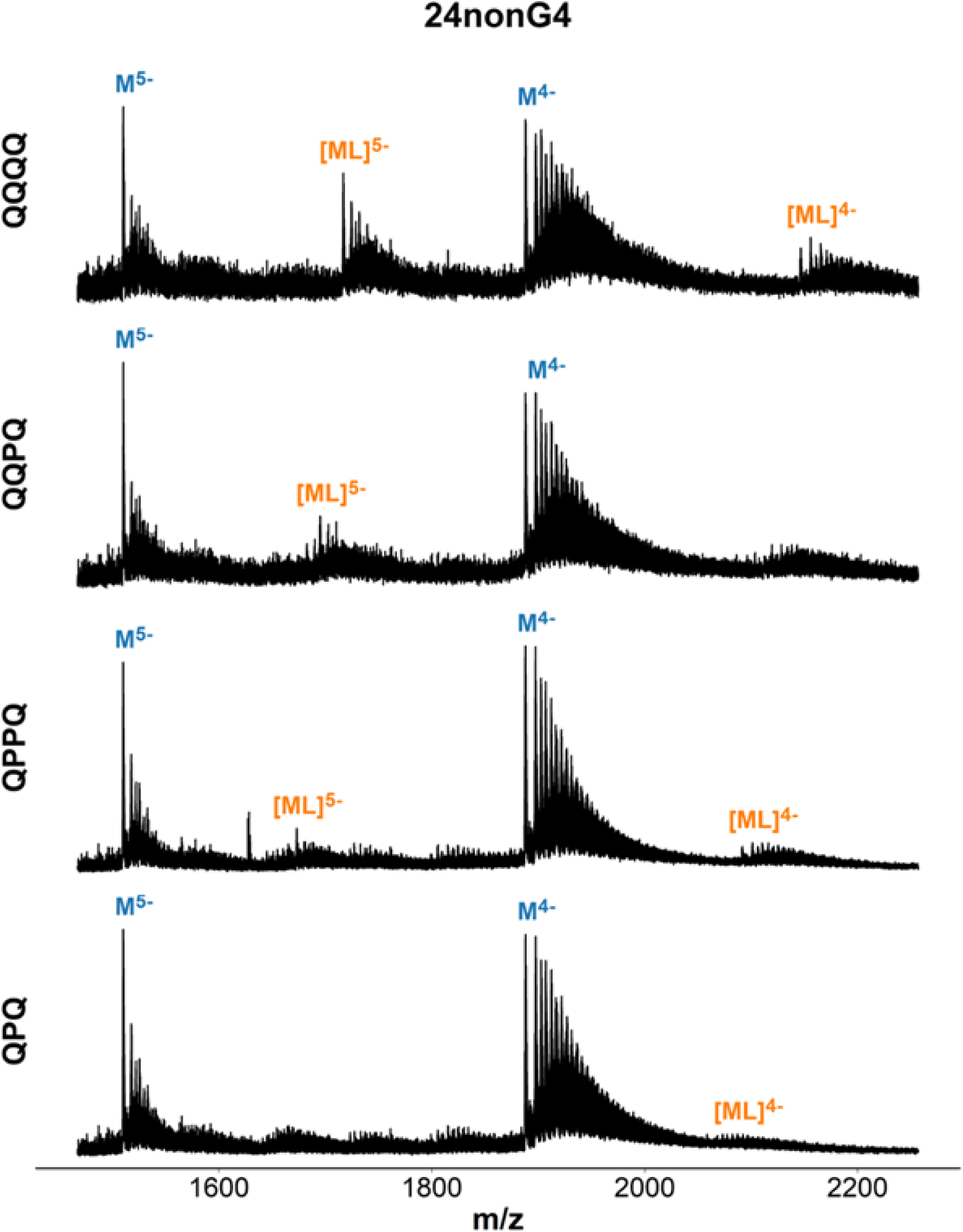
Mass spectra of 24nonG4 (dTGGGATGCGACAGAGAGGACGGGA) in presence of ligand. Samples contain 10 µM DNA, 20 µM ligand, 0.5 mM KCl, 100 mM TMAA (pH 6.8).

**Figure S66.**
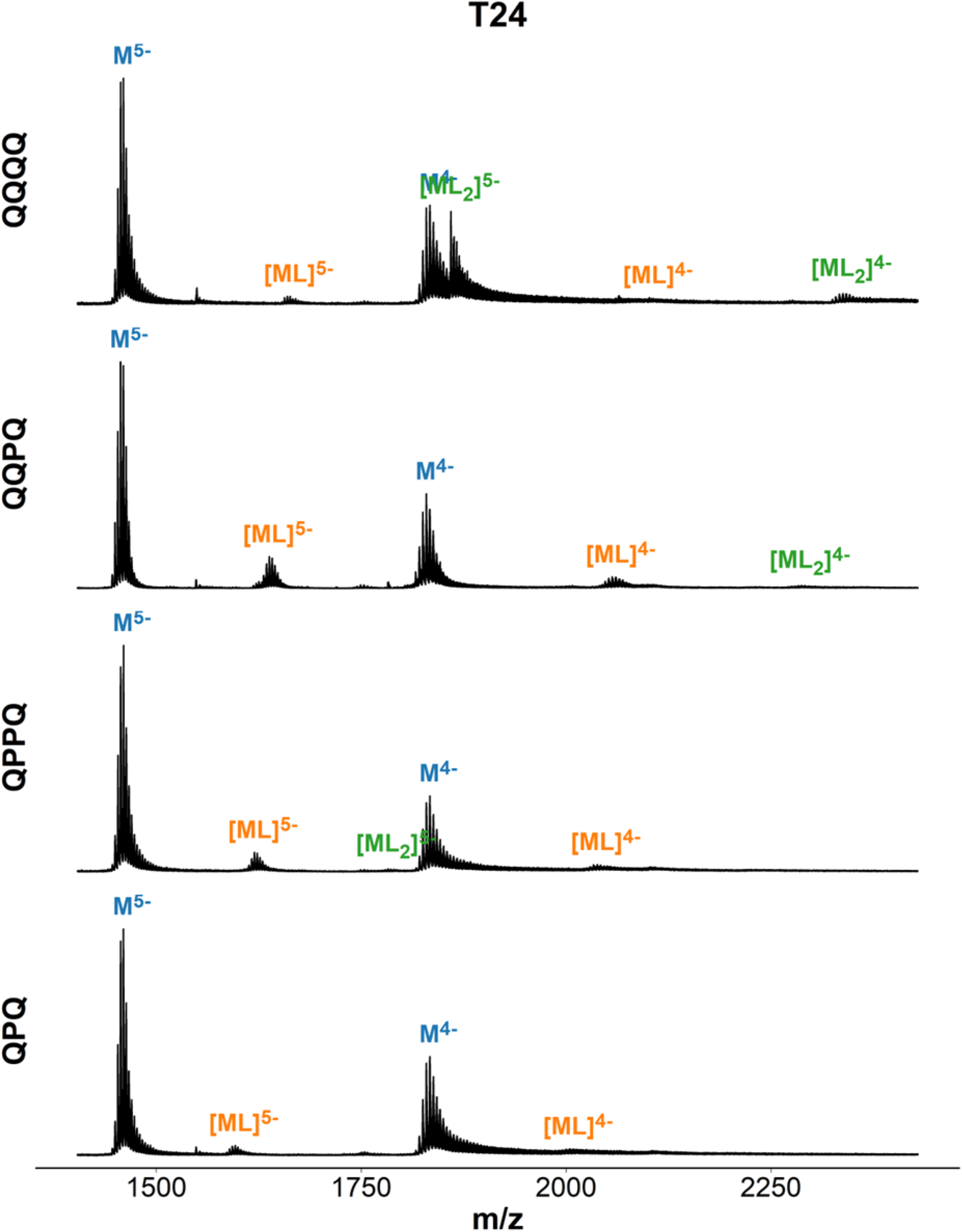
Mass spectra of T24 (dTTTTTTTTTTTTTTTTTTTTTTTT) in presence of ligand. Samples contain 10 µM DNA, 20 µM ligand, 150 mM ammonium acetate (pH 6.8).

**Figure S67.**
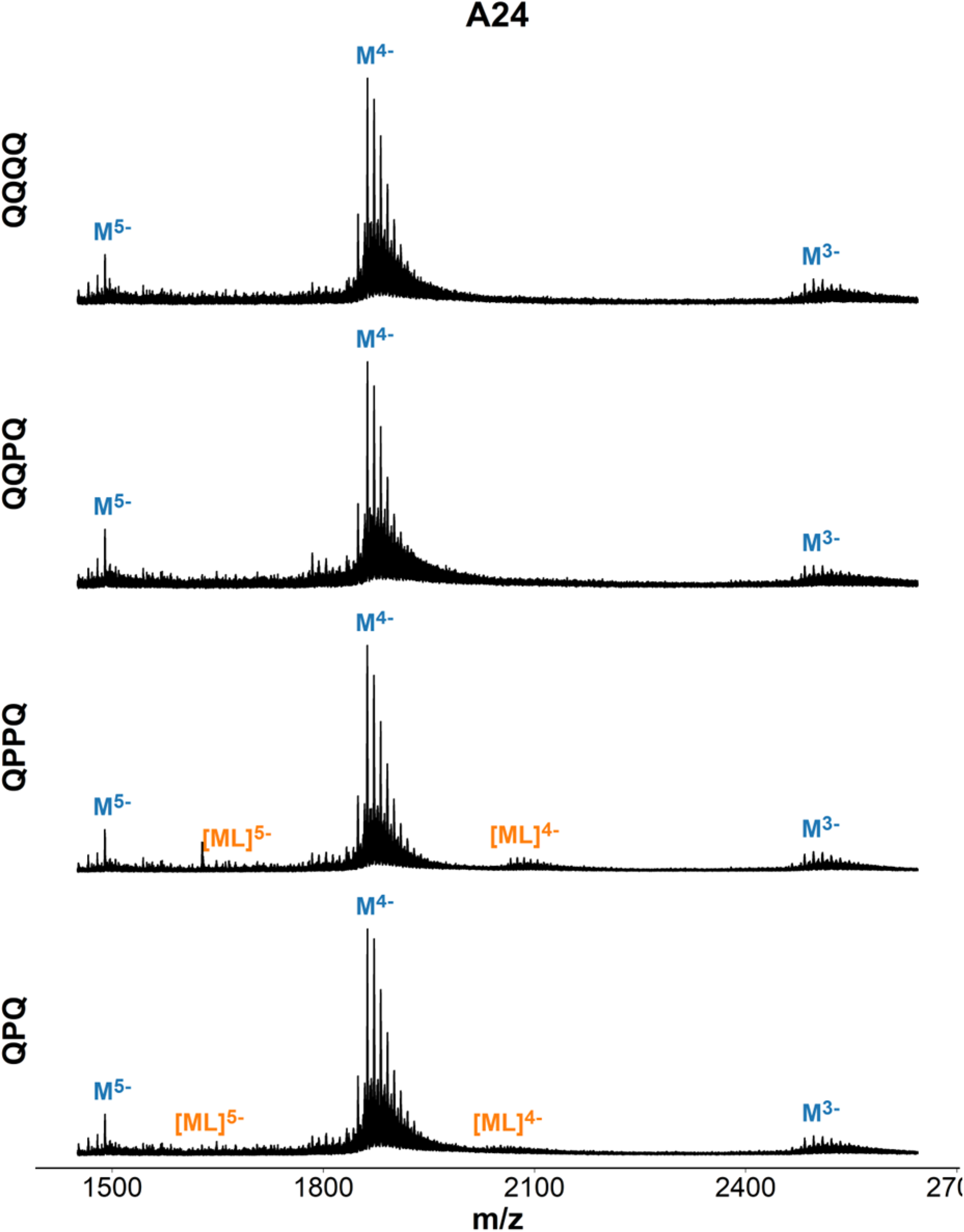
Mass spectra of A24(dAAAAAAAAAAAAAAAAAAAAAAAA) in presence of ligand. Samples contain 10 µM DNA, 20 µM ligand, 0.5 mM KCl, 100 mM TMAA (pH 6.8).

**Figure S68.**
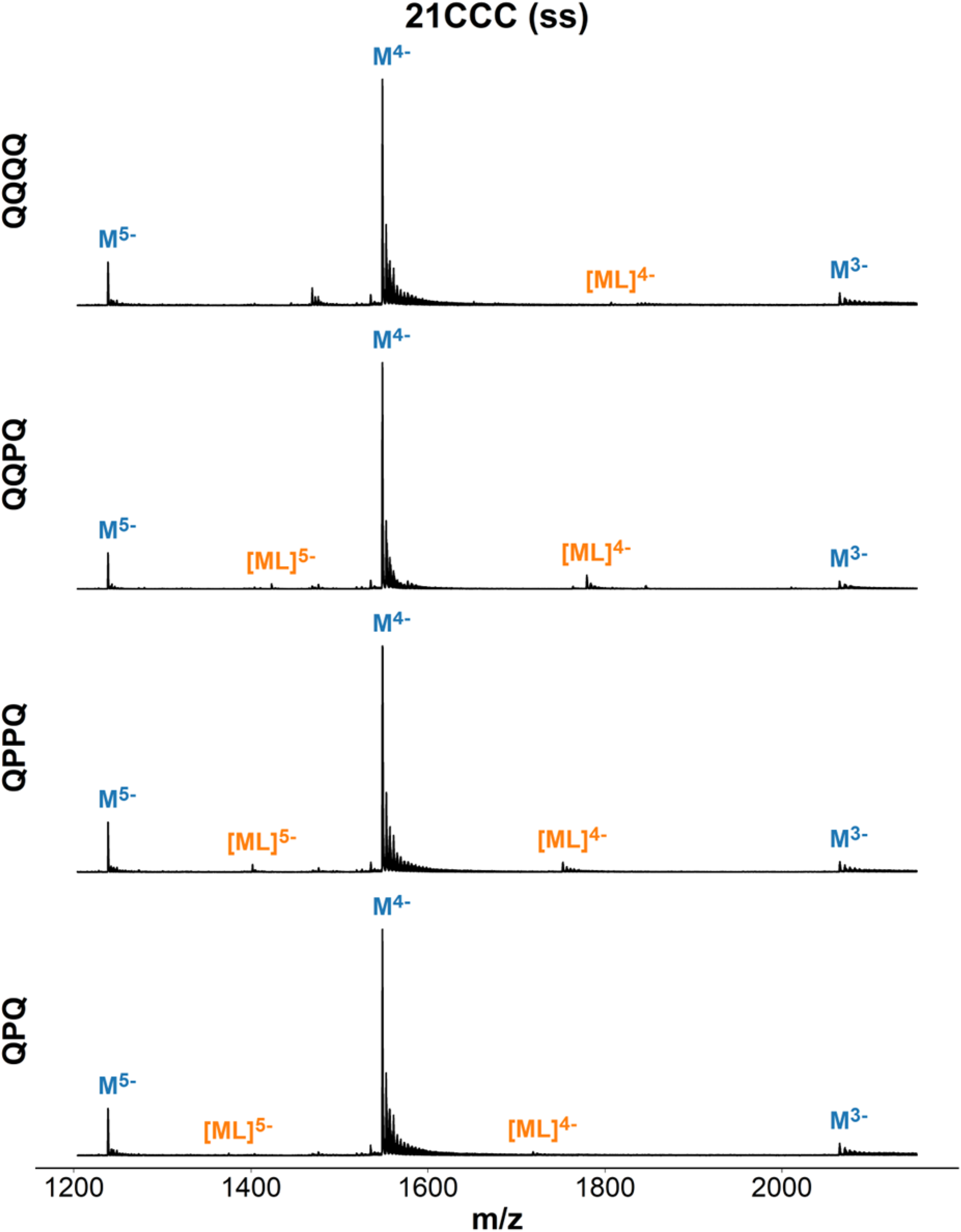
Mass spectra of 21CCC (dCCCTAACCCTAACCCTAACCC) in presence of ligand. Samples contain 10 µM DNA, 20 µM ligand, 150 mM ammonium acetate (pH 6.8).

**Figure S69.**
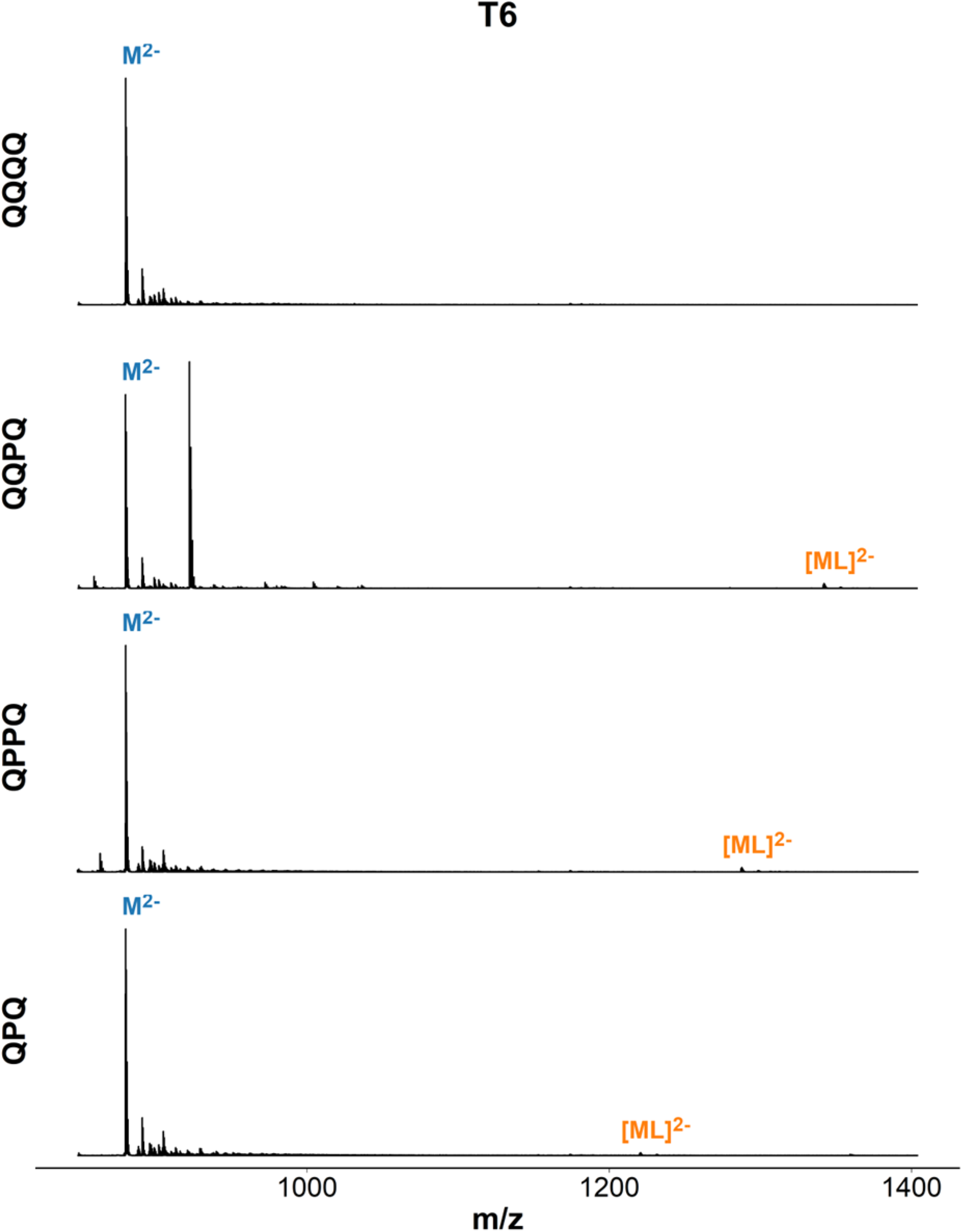
Mass spectra of T6 (dTTTTTT) in presence of ligand. Samples contain 10 µM DNA, 20 µM ligand, 150 mM ammonium acetate (pH 6.8).

#### ESI-MS titrations: K_D_ values and response factor estimates

**Table S3.** All K_D_ values obtained from ESI-MS titration of 6 DNA sequences with 7 foldamer ligands. K_D_ < 5 µM are highlighted in blue, K_D_ > 100 µM are faded out.

| Dissociation constant estimates ($\mu\text{M}$) | | | | | | | | | |
| --- | --- | --- | --- | --- | --- | --- | --- | --- | --- |
| | $K_{D1}$ | | $K_{D2}$ | | $K_{D3}$ | | $K_{D4}$ | | |
|  | ESTIMATE | STD.DEV. | ESTIMATE | STD.DEV. | ESTIMATE | STD.DEV. | ESTIMATE | STD.DEV. |  |
| **2LK7** |  |  |  |  |  |  |  |  |  |
| QQPQ | 0.18 | 0.079 | 2.1 | 0.69 |  |  |  |  |  |
| **1XAV** |  |  |  |  |  |  |  |  |  |
| Q3 | 0.11 | 0.074 | 1.2 | 0.94 | 4.1 | 2.5 |  |  |  |
| QPQ | 13 | 0.75 | 110 | 43 |  |  |  |  |  |
| QQQQ | 0.075 | 0.059 | 0.83 | 0.49 |  |  |  |  |  |
| QQPQ | 0.41 | 0.29 | 1.9 | 1.2 |  |  |  |  |  |
| QPPQ | 6.9 | 2 | 9.7 | 5.7 |  |  |  |  |  |
| Q5 | 0.25 | 0.2 | 0.7 | 0.53 |  |  |  |  |  |
| Q8 | 28 | 6.7 |  |  |  |  |  |  |  |
| **222T** |  |  |  |  |  |  |  |  |  |
| Q3 | 0.1 | 0.1 | 2.8 | 3.1 | 6.1 | 11 | 64 | 440 |  |
| QPQ | 1.8 | 0.25 | 50 | 13 |  |  |  |  |  |
| QQQQ | 0.088 | 0.11 | 1.6 | 1.1 |  |  |  |  |  |
| QQPQ | 0.027 | 0.081 | 0.064 | 0.19 |  |  |  |  |  |
| QPPQ | 0.1 | 0.093 | 1.4 | 0.85 |  |  |  |  |  |
| Q5 | 0.011 | 0.014 | 0.4 | 0.35 |  |  |  |  |  |
| Q8 | 7.8 | 1.1 |  |  |  |  |  |  |  |
| **5YEY** |  |  |  |  |  |  |  |  |  |
| Q3 | 180 | 28 |  |  |  |  |  |  |  |
| QPQ | 1900 | 780 |  |  |  |  |  |  |  |
| QQQQ | 6.6 | 1.3 |  |  |  |  |  |  |  |
| QQPQ | 0.54 | 0.098 | 200 | 70 |  |  |  |  |  |
| QPPQ | 110 | 23 | 310 | 760 |  |  |  |  |  |
| Q5 | 3.9 | 0.37 |  |  |  |  |  |  |  |
| **21G** |  |  |  |  |  |  |  |  |  |
| Q3 | 78 | 19 | 1500 | 21000 |  |  |  |  |  |
| QPQ | 170 | 26 |  |  |  |  |  |  |  |
| QQQQ | 330 | 23 |  |  |  |  |  |  |  |
| QQPQ | 8.6 | 2.3 | 14 | 9.2 |  |  |  |  |  |
| QPPQ | 160 | 20 | 1000 | 4500 |  |  |  |  |  |
| Q5 | 210 | 74 | 47 | 45 |  |  |  |  |  |
| **SS24** |  |  |  |  |  |  |  |  |  |
| Q3 | 12 | 6.2 | 28 | 87 | 15 | 63 |  |  |  |
| QPQ | 580 | 110 |  |  |  |  |  |  |  |
| QQQQ | 180 | 14 |  |  |  |  |  |  |  |
| QQPQ | 150 | 21 |  |  |  |  |  |  |  |
| QPPQ | 270 | 30 |  |  |  |  |  |  |  |
| Q5 | 130 | 42 | 44 | 41 |  |  |  |  |  |
| Q8 | 210 | 45 |  |  |  |  |  |  |  |
| **T24** |  |  |  |  |  |  |  |  |  |
| Q3 | 3400 | 8200 | 0.36 | 0.91 | 260000 | 1.2e+08 | 0.0092 | 4.2 |  |
| QQQQ | 350 | 3400 | 0.53 | 5.5 |  |  |  |  |  |
| QQPQ | 270 | 24 |  |  |  |  |  |  |  |
| QPPQ | 190 | 25 |  |  |  |  |  |  |  |
| Q5 | 180 | 490 | 1 | 3 |  |  |  |  |  |
| Q8 | 290 | 87 |  |  |  |  |  |  |  |

**Table S4.**
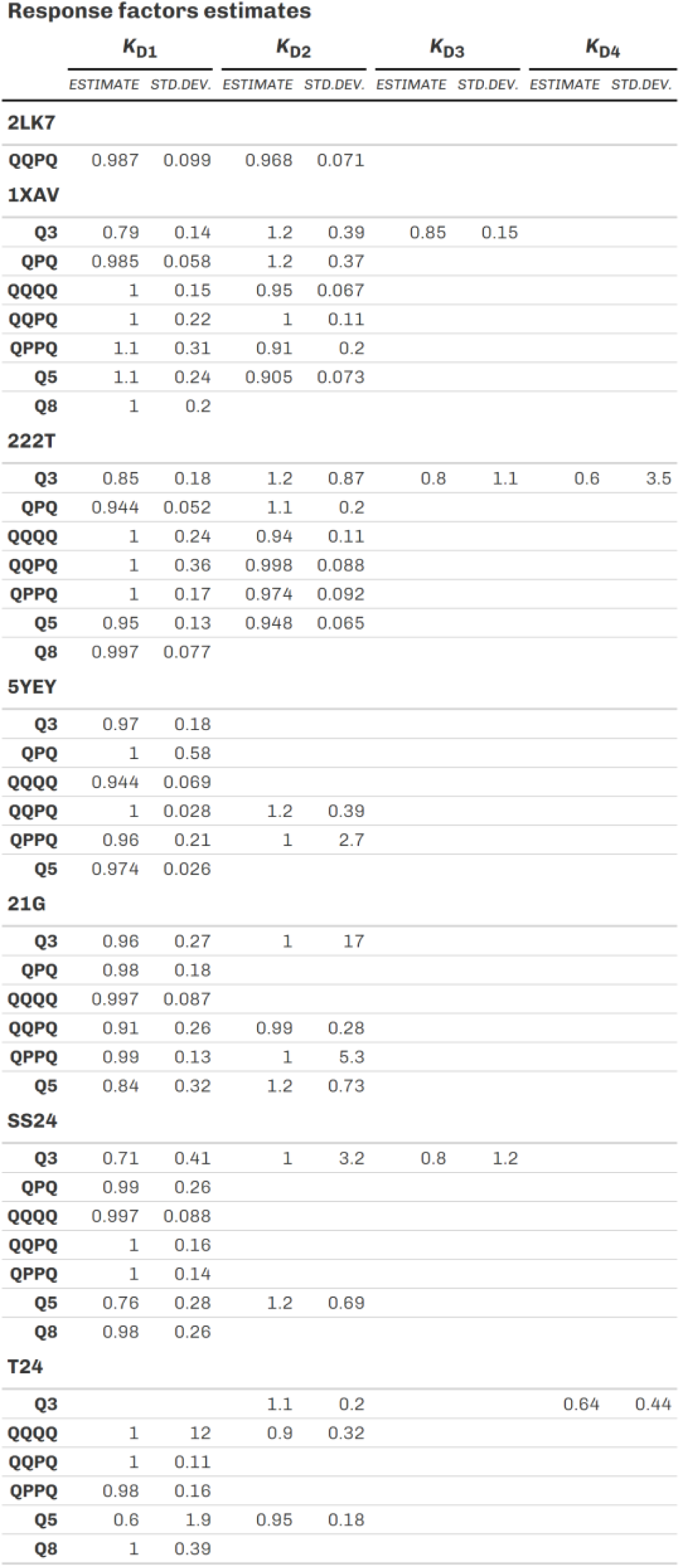
Estimated response factors of complex species ML, ML_2_, ML_3_, ML_4_ relative to the unbound DNA (M with R = 1) using dT_6_ as an internal calibrant. The calculation method was previously described in [^4^].

#### ESI-MS titrations: Mass spectra

**Figure S70.**
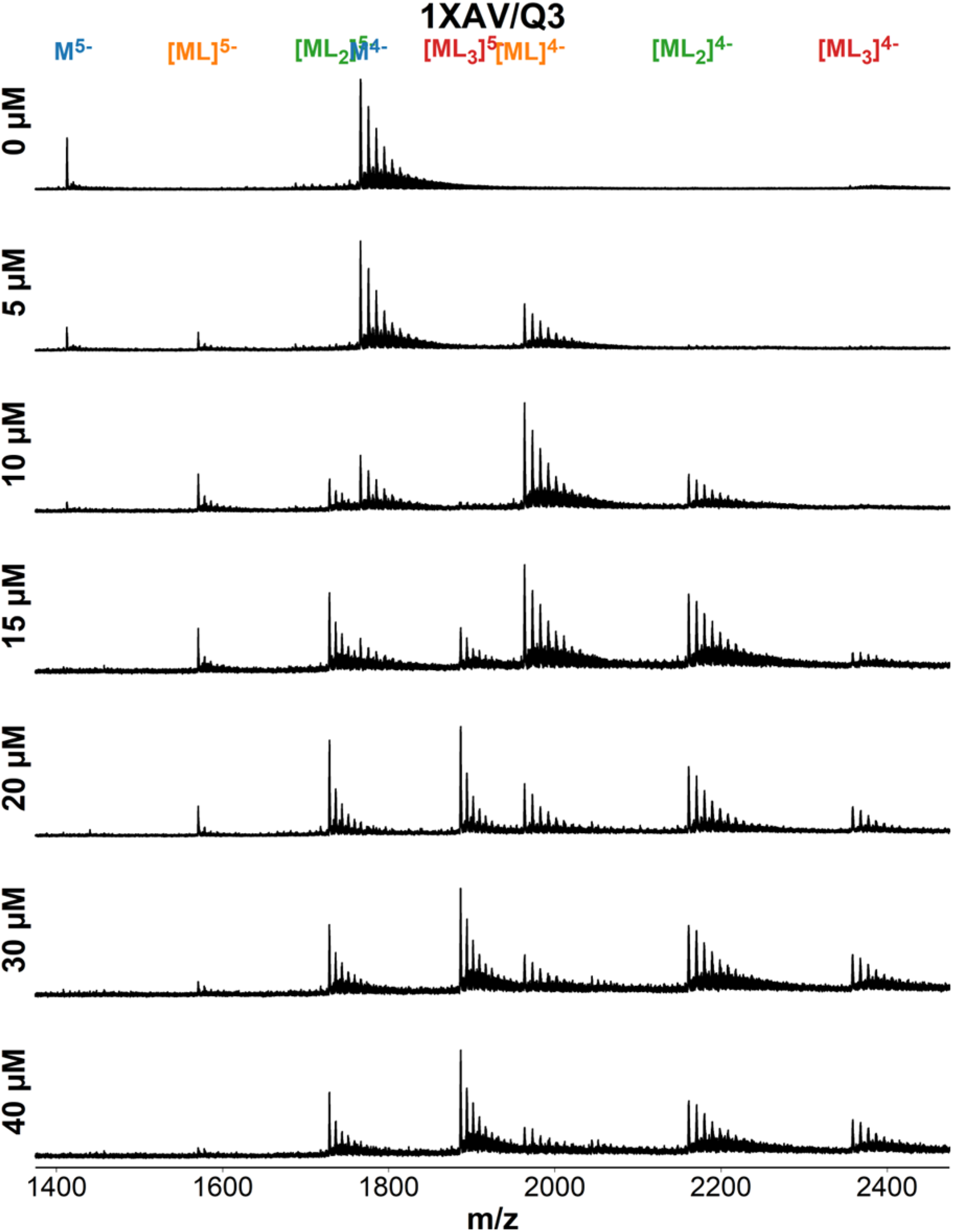
ESI-MS titration of 1XAV (dT**G**A**GGG**T**GGG**TA**GGG**T**GGG**TAA) with foldamer QQQ. Samples contain 10 µM DNA, 0-40 µM ligand, 0.5 mM KCl, 100 mM TMAA (pH 6.8).

**Figure S71.**
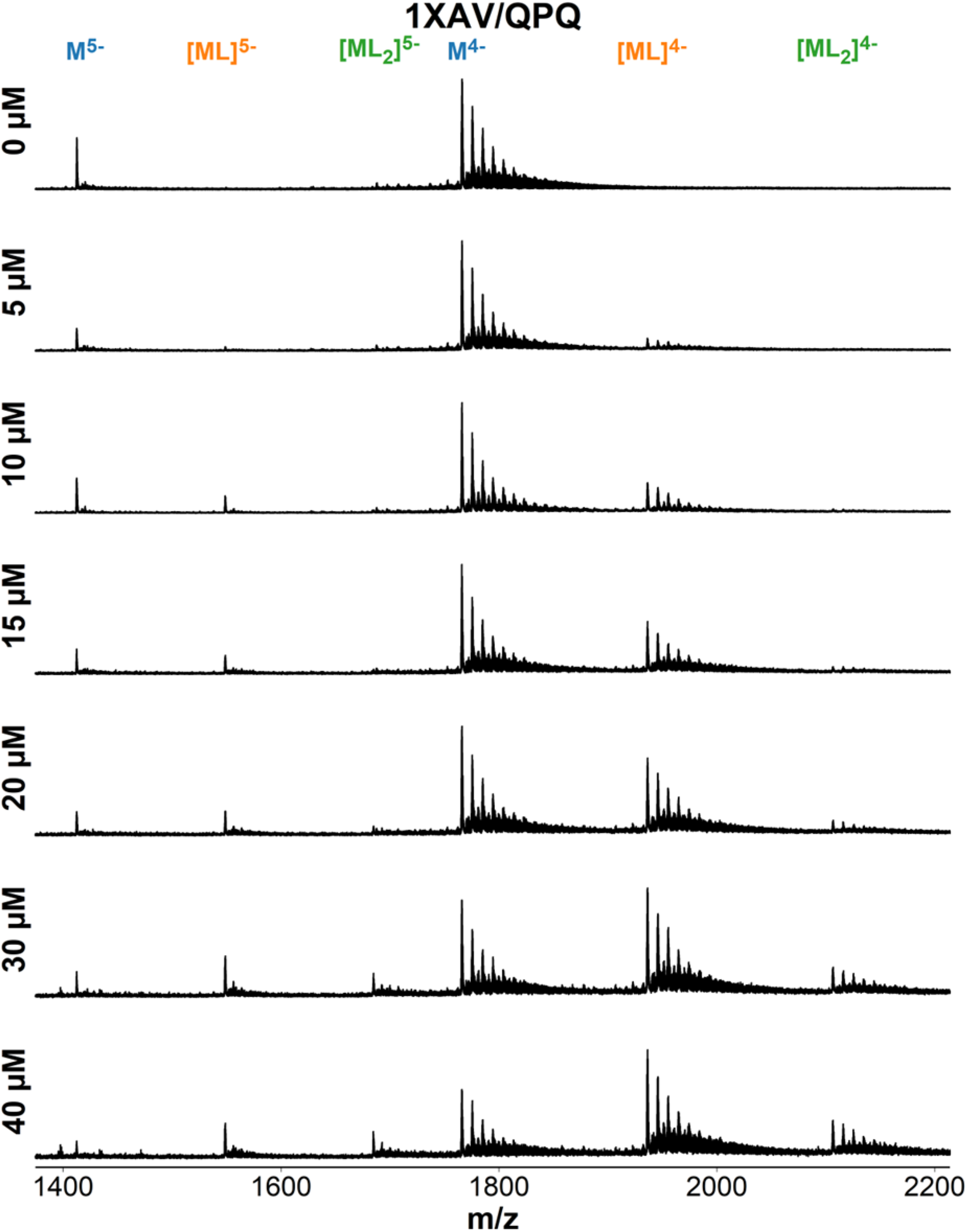
ESI-MS titration of 1XAV (dT**G**A**GGG**T**GGG**TA**GGG**T**GGG**TAA) with foldamer QPQ. Samples contain 10 µM DNA, 0-40 µM ligand, 0.5 mM KCl, 100 mM TMAA (pH 6.8).

**Figure S72.**
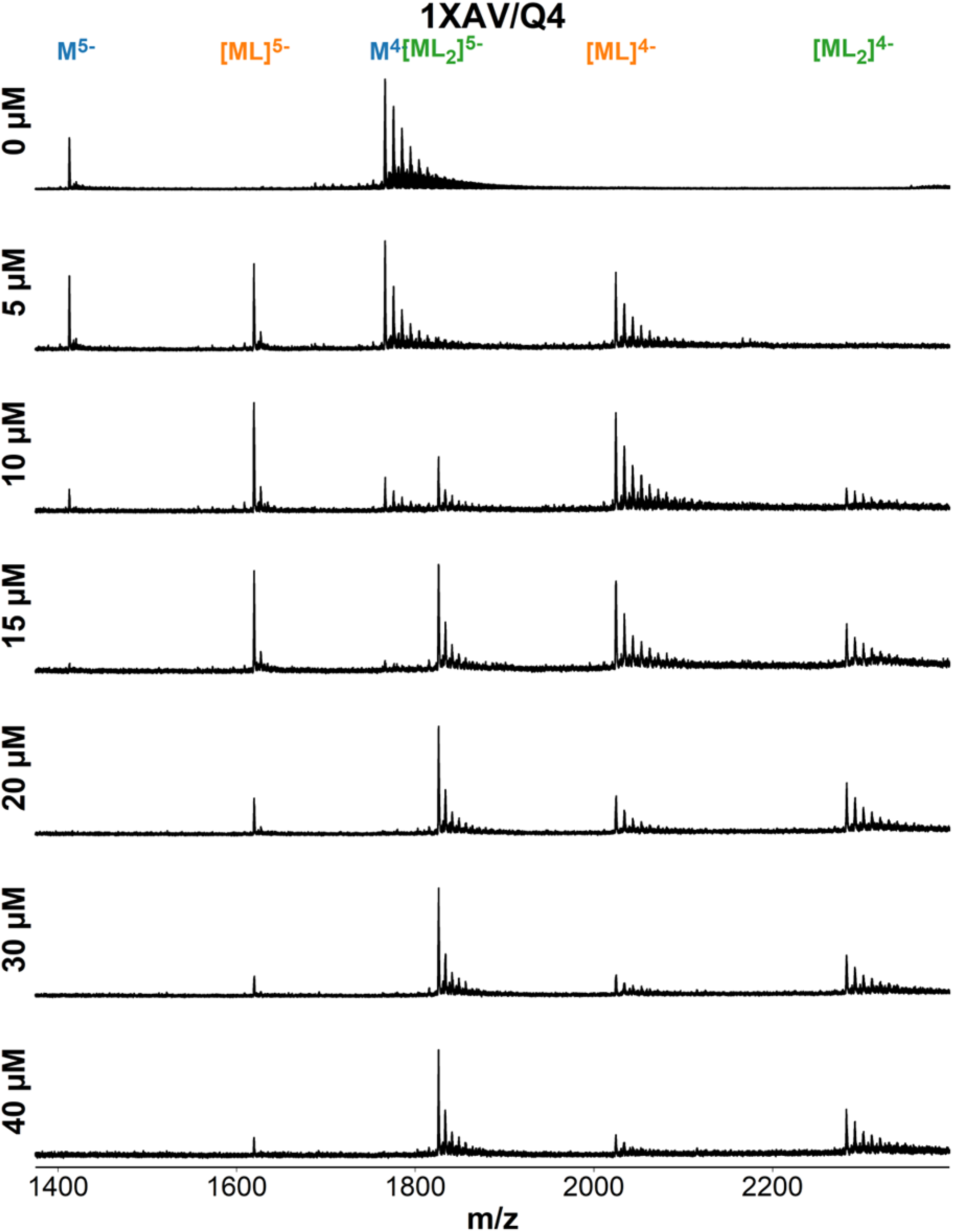
ESI-MS titration of 1XAV (dT**G**A**GGG**T**GGG**TA**GGG**T**GGG**TAA) with foldamer QQQQ. Samples contain 10 µM DNA, 0-40 µM ligand, 0.5 mM KCl, 100 mM TMAA (pH 6.8).

**Figure S73.**
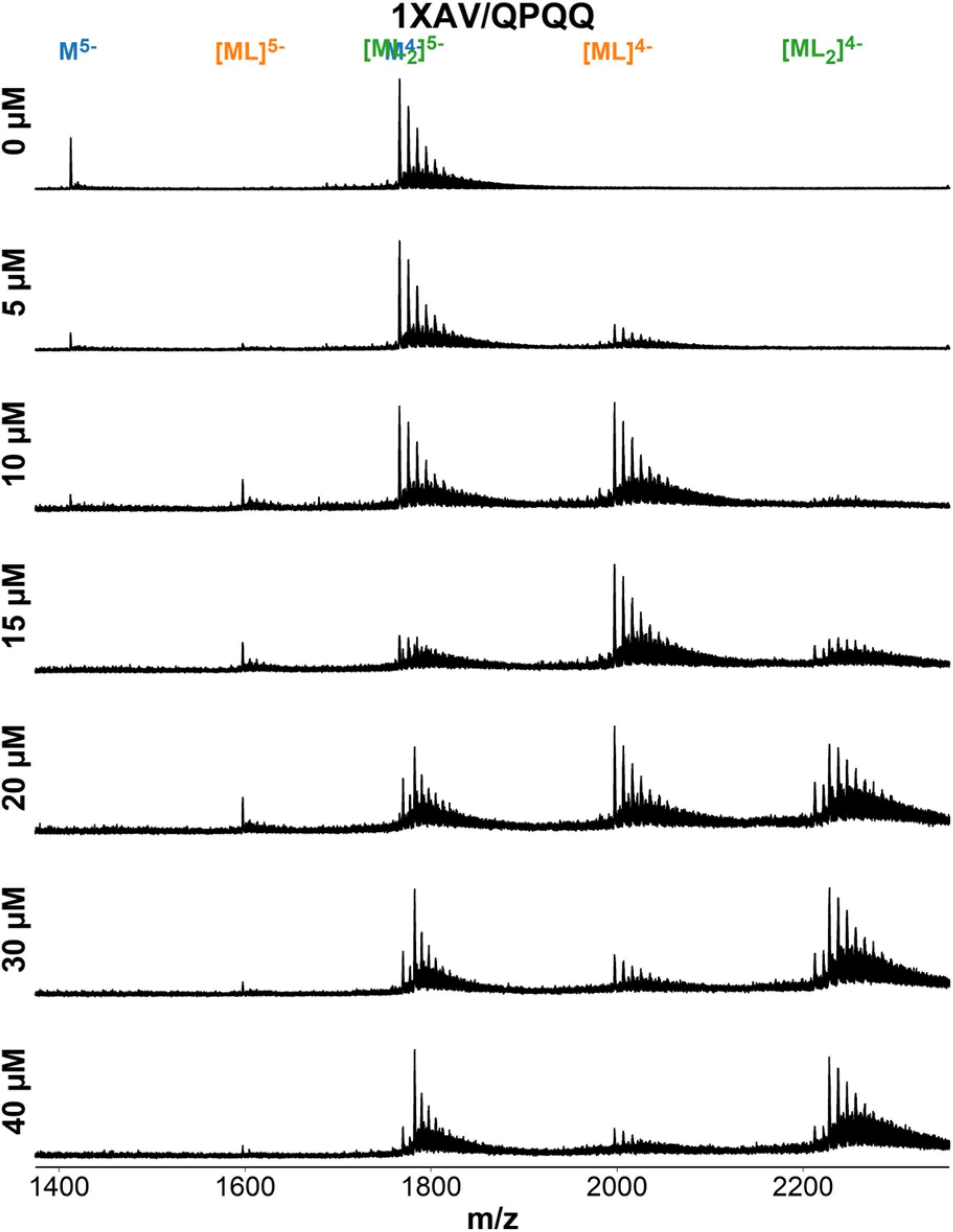
ESI-MS titration of 1XAV (dT**G**A**GGG**T**GGG**TA**GGG**T**GGG**TAA) with foldamer QQPQ. Samples contain 10 µM DNA, 0-40 µM ligand, 0.5 mM KCl, 100 mM TMAA (pH 6.8).

**Figure S74.**
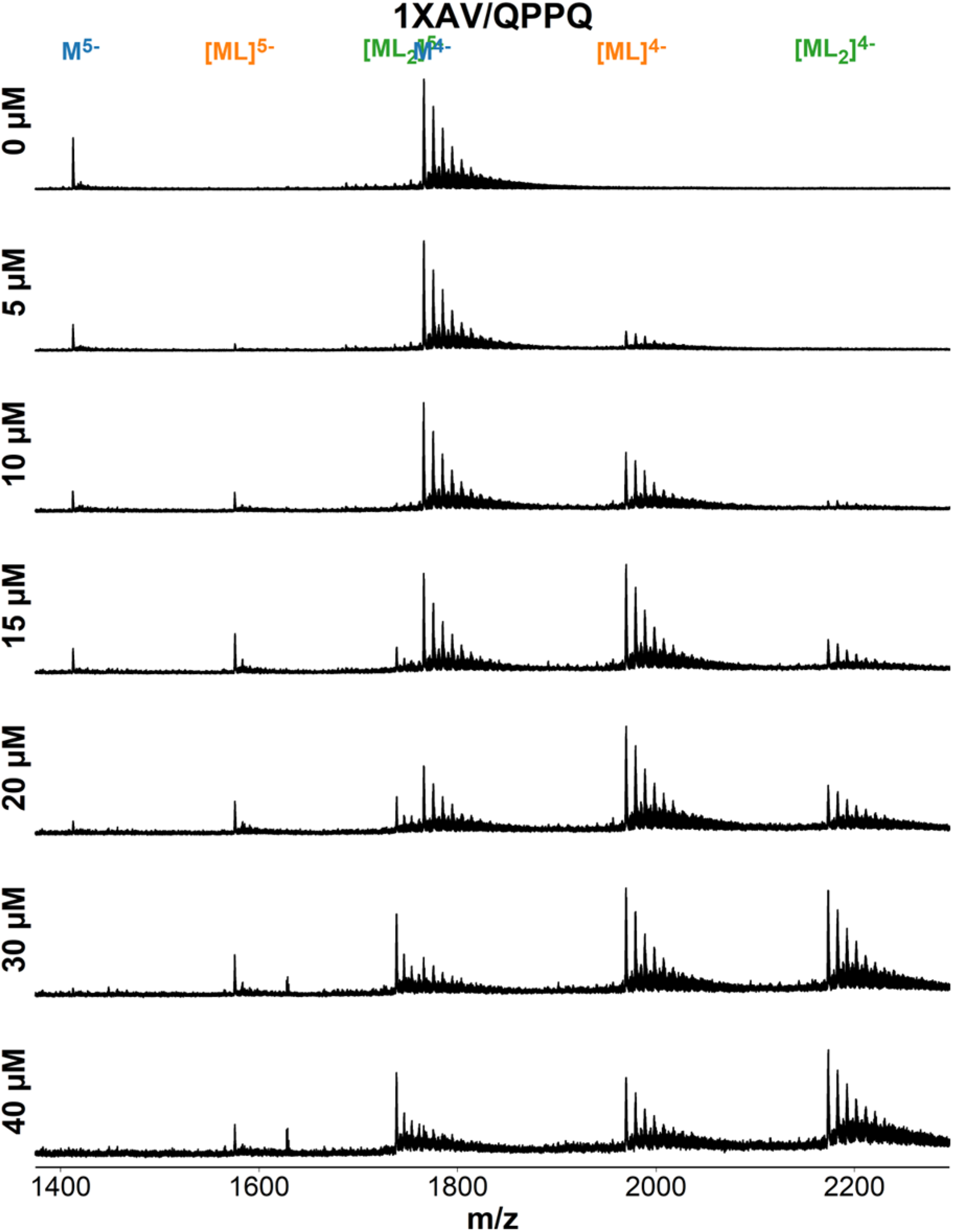
ESI-MS titration of 1XAV (dT**G**A**GGG**T**GGG**TA**GGG**T**GGG**TAA) with foldamer QPPQ. Samples contain 10 µM DNA, 0-40 µM ligand, 0.5 mM KCl, 100 mM TMAA (pH 6.8).

**Figure S75.**
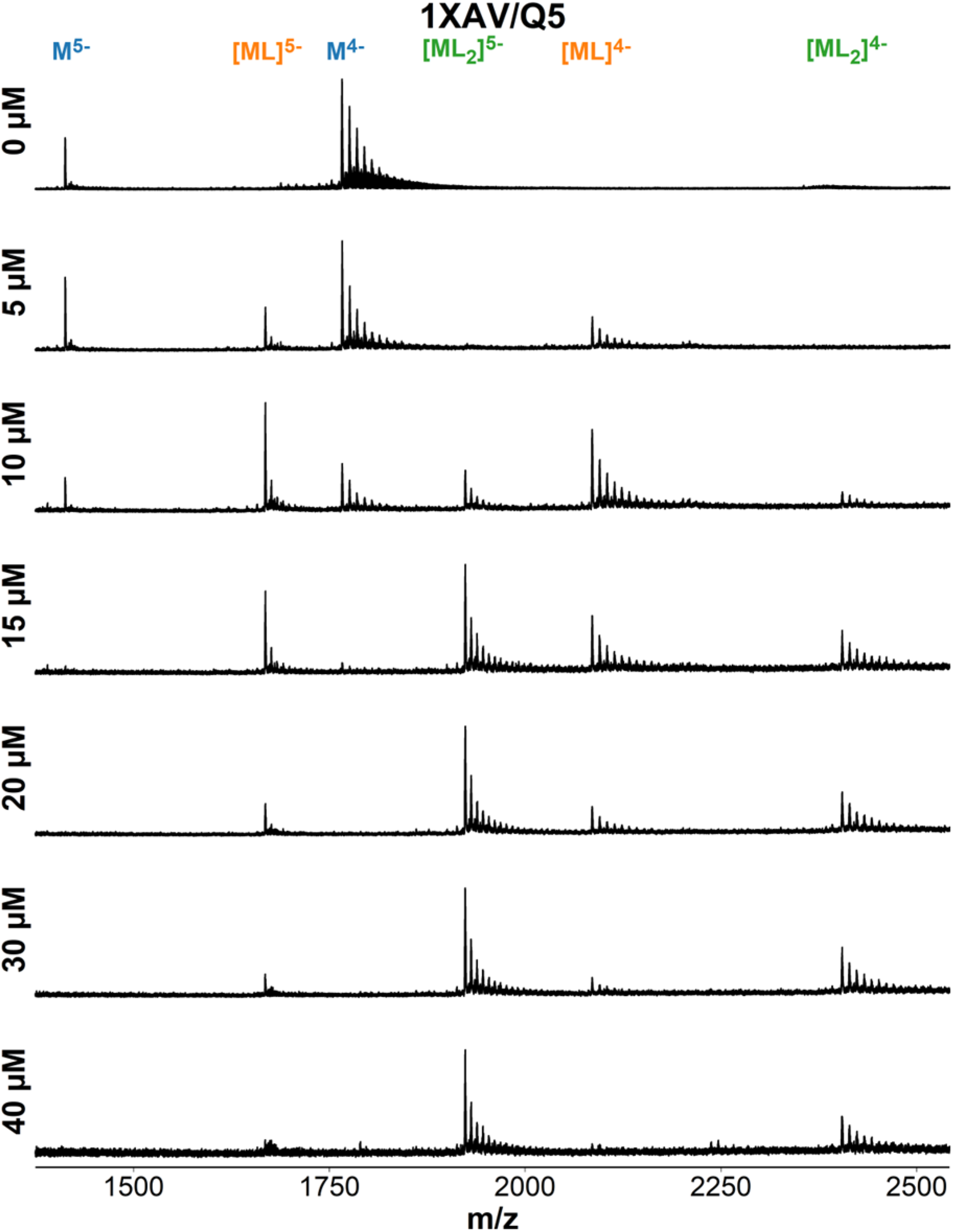
ESI-MS titration of 1XAV (dT**G**A**GGG**T**GGG**TA**GGG**T**GGG**TAA) with foldamer QQQQQ. Samples contain 10 µM DNA, 0-40 µM ligand, 0.5 mM KCl, 100 mM TMAA (pH 6.8).

**Figure S76.**
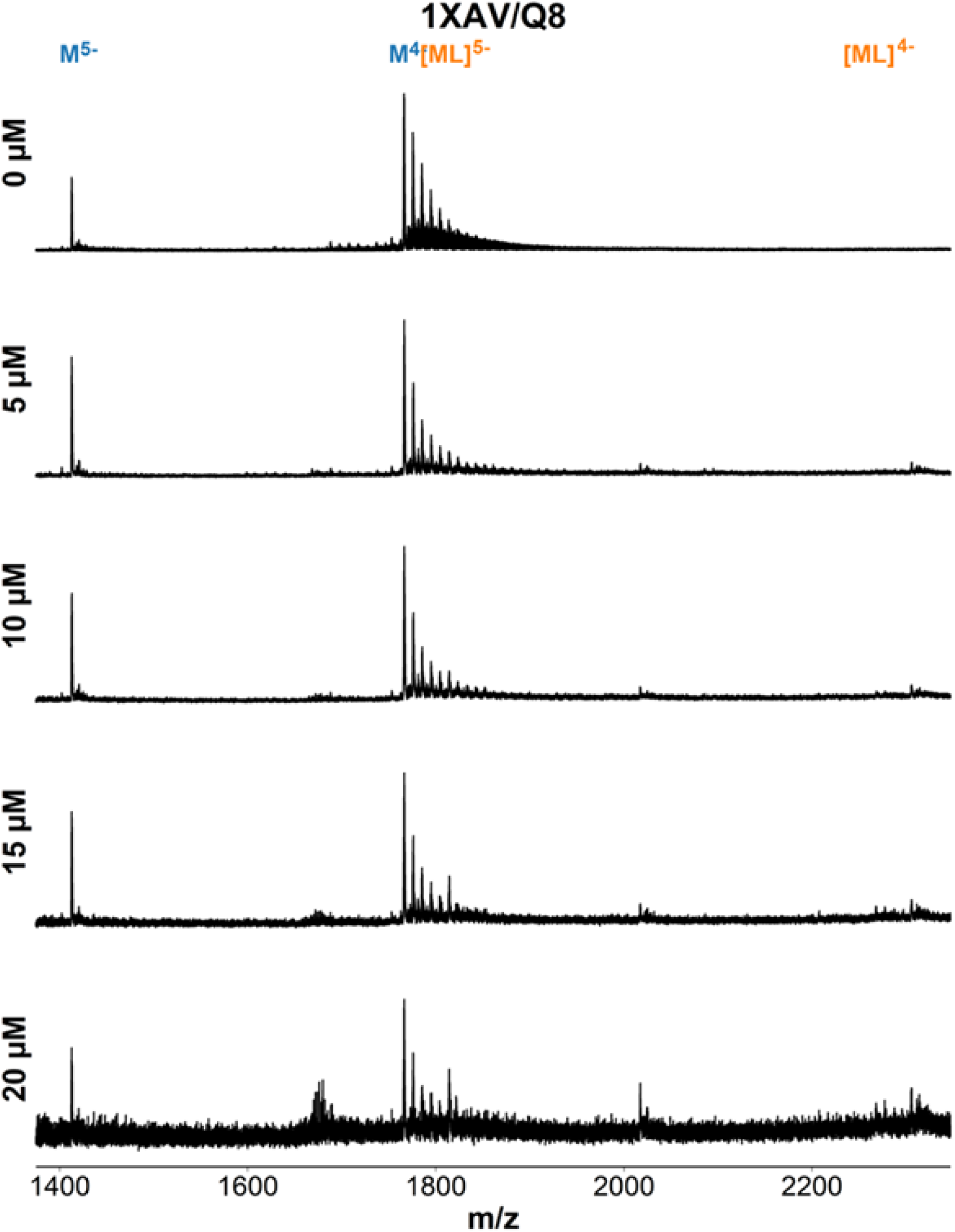
ESI-MS titration of 1XAV (dT**G**A**GGG**T**GGG**TA**GGG**T**GGG**TAA) with foldamer QQQQQQQQ. Samples contain 10 µM DNA, 0-20 µM ligand, 0.5 mM KCl, 100 mM TMAA (pH 6.8).

**Figure S77.**
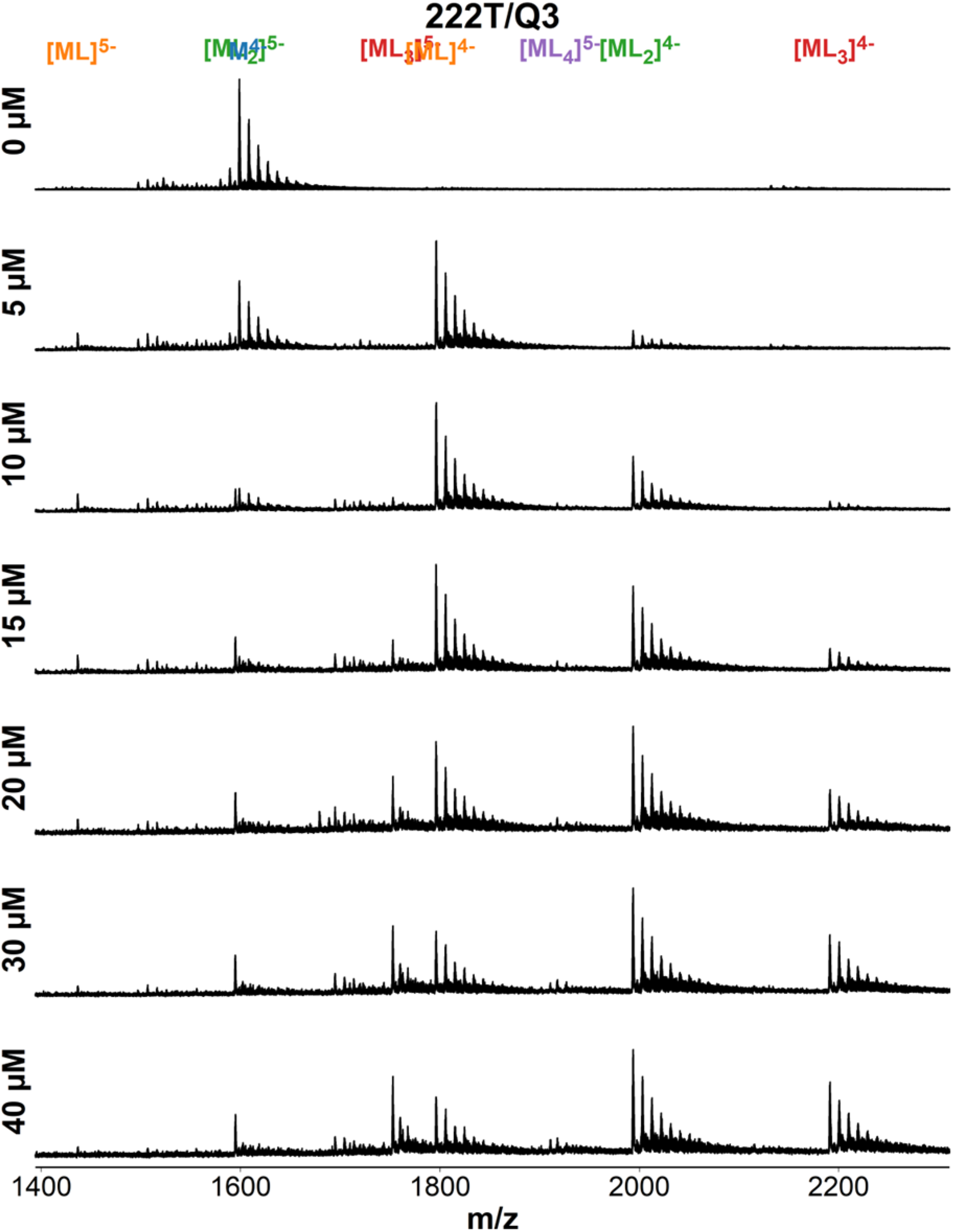
ESI-MS titration of 222T (dT**GGG**TT**GGG**TT**GGG**TT**GGG**T) with foldamer QQQ. Samples contain 10 µM DNA, 0-40 µM ligand, 0.5 mM KCl, 100 mM TMAA (pH 6.8).

**Figure S78.**
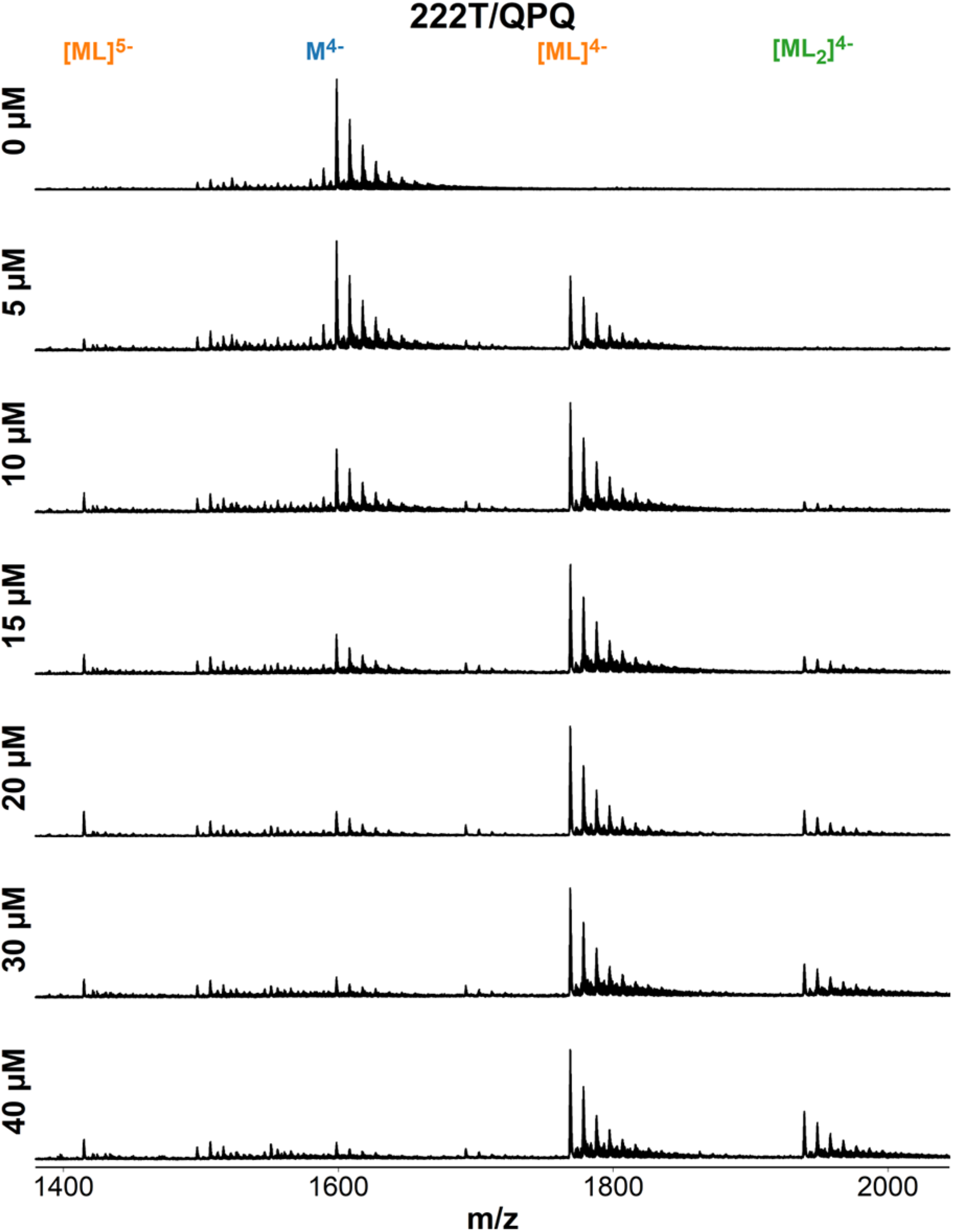
ESI-MS titration of 222T (dT**GGG**TT**GGG**TT**GGG**TT**GGG**T) with foldamer QPQ. Samples contain 10 µM DNA, 0-40 µM ligand, 0.5 mM KCl, 100 mM TMAA (pH 6.8).

**Figure S79.**
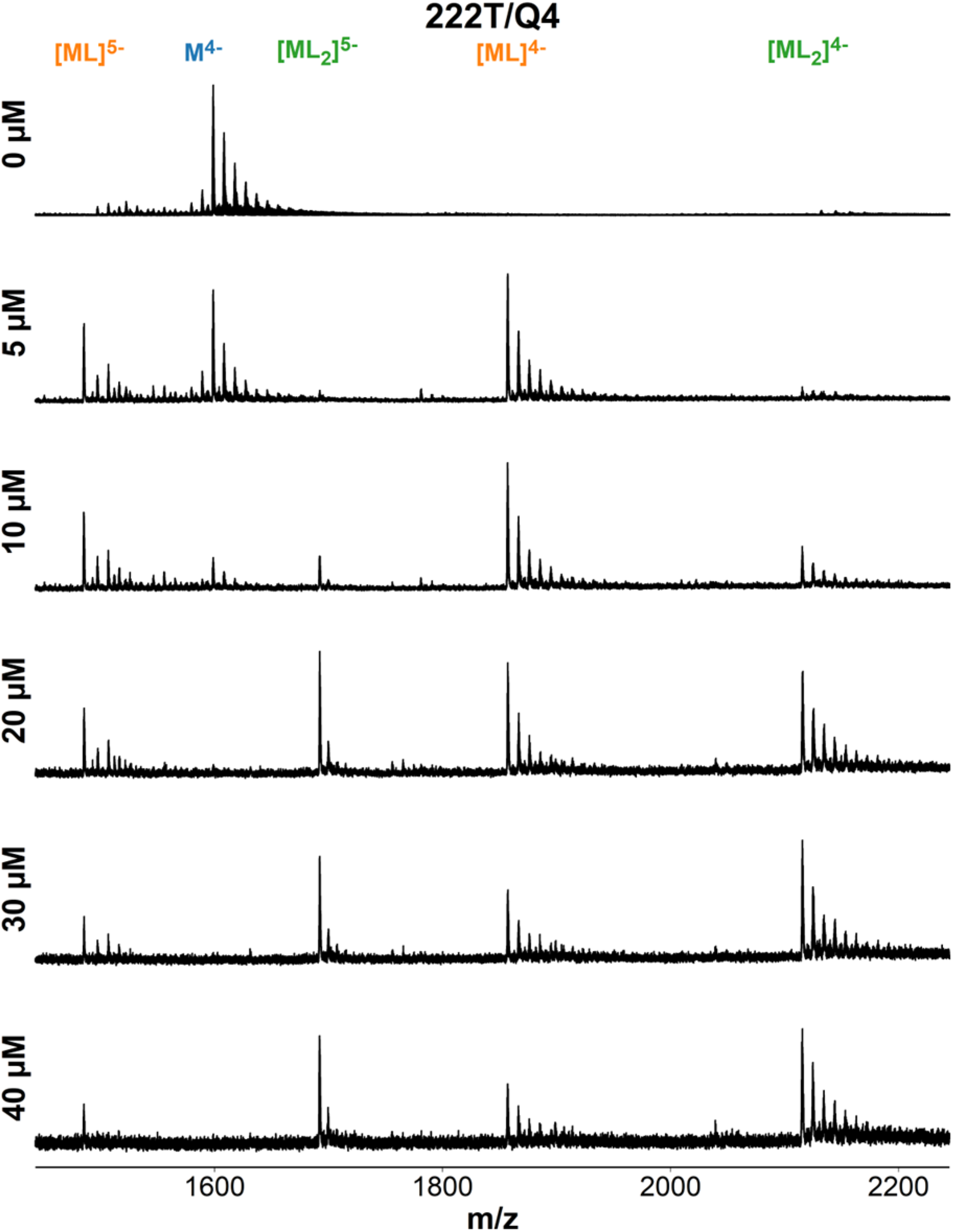
ESI-MS titration of 222T (dT**GGG**TT**GGG**TT**GGG**TT**GGG**T) with foldamer QQQQ. Samples contain 10 µM DNA, 0-40 µM ligand, 0.5 mM KCl, 100 mM TMAA (pH 6.8).

**Figure S80.**
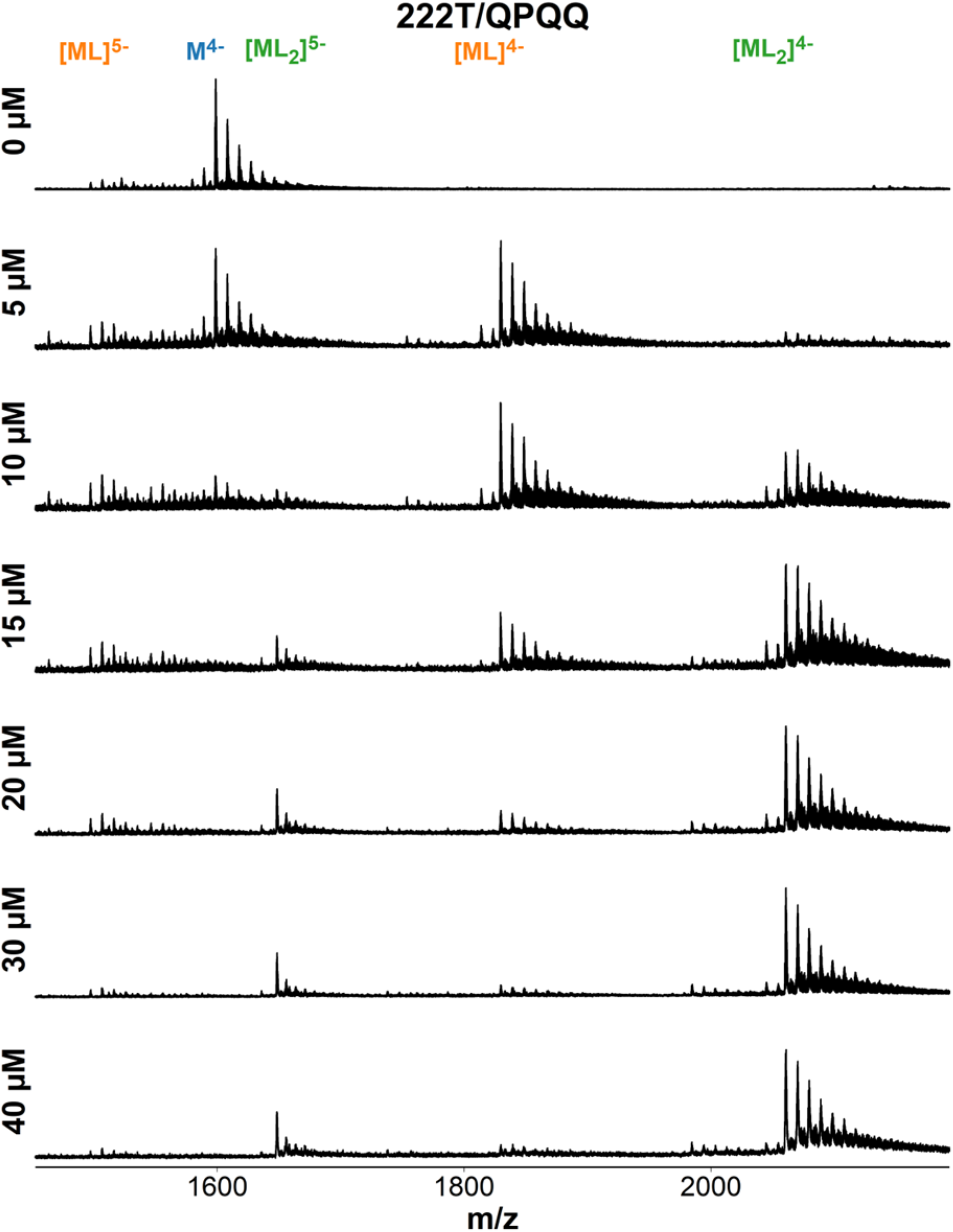
ESI-MS titration of 222T (dT**GGG**TT**GGG**TT**GGG**TT**GGG**T) with foldamer QQPQ. Samples contain 10 µM DNA, 0-40 µM ligand, 0.5 mM KCl, 100 mM TMAA (pH 6.8).

**Figure S81.**
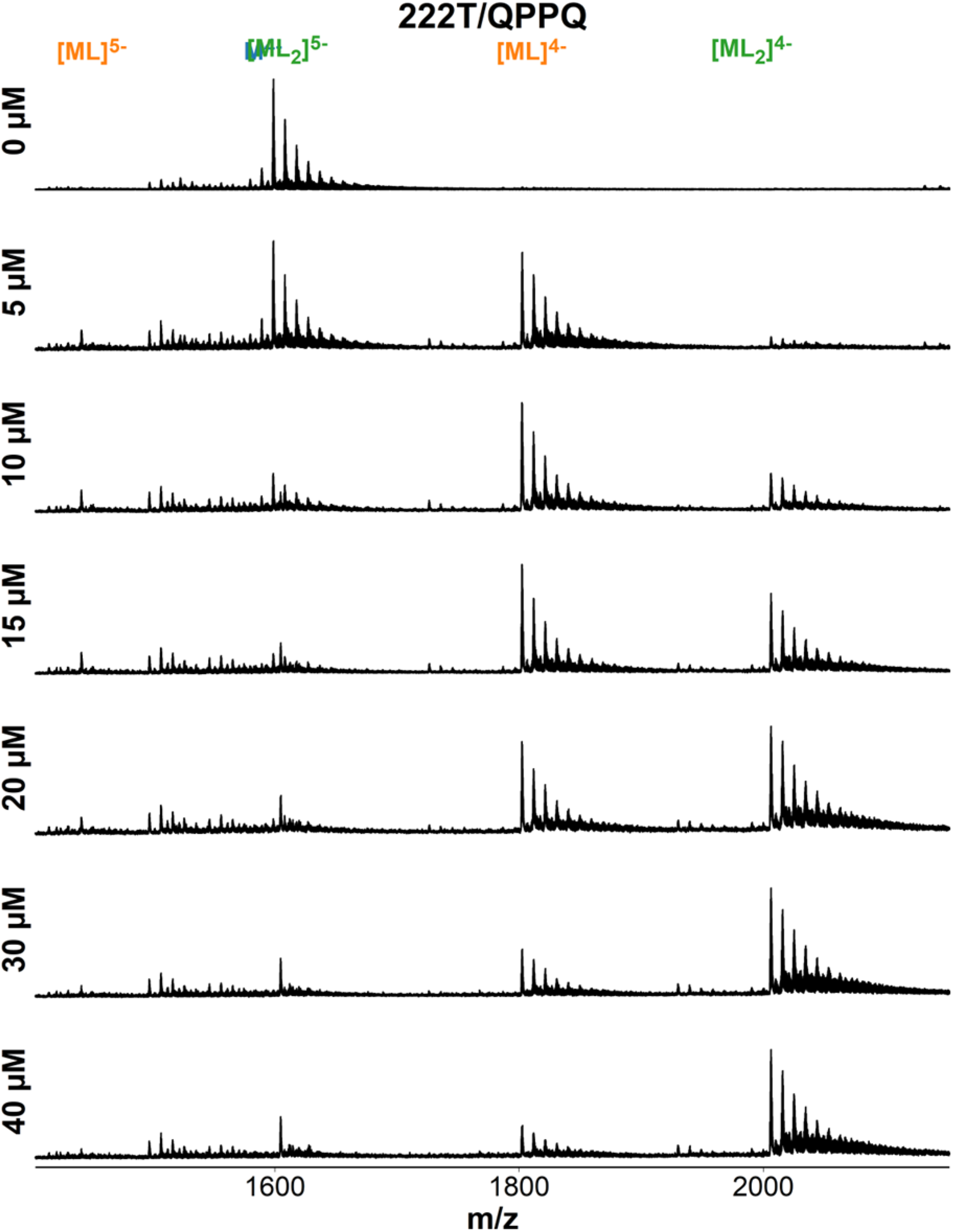
ESI-MS titration of 222T (dT**GGG**TT**GGG**TT**GGG**TT**GGG**T) with foldamer QPPQ. Samples contain 10 µM DNA, 0-40 µM ligand, 0.5 mM KCl, 100 mM TMAA (pH 6.8).

**Figure S82.**
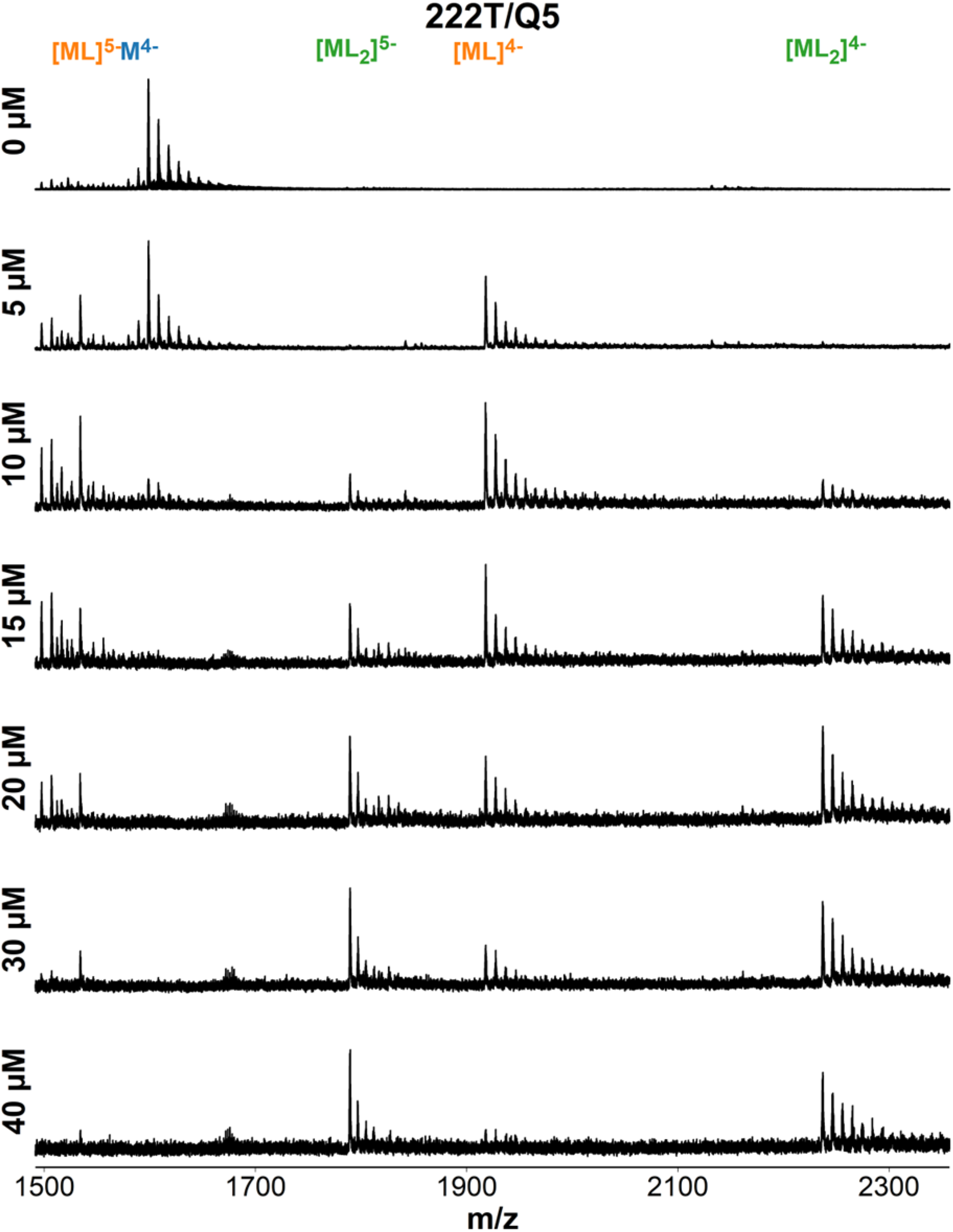
ESI-MS titration of 222T (dT**GGG**TT**GGG**TT**GGG**TT**GGG**T) with foldamer QQQQQ. Samples contain 10 µM DNA, 0-40 µM ligand, 0.5 mM KCl, 100 mM TMAA (pH 6.8).

**Figure S83.**
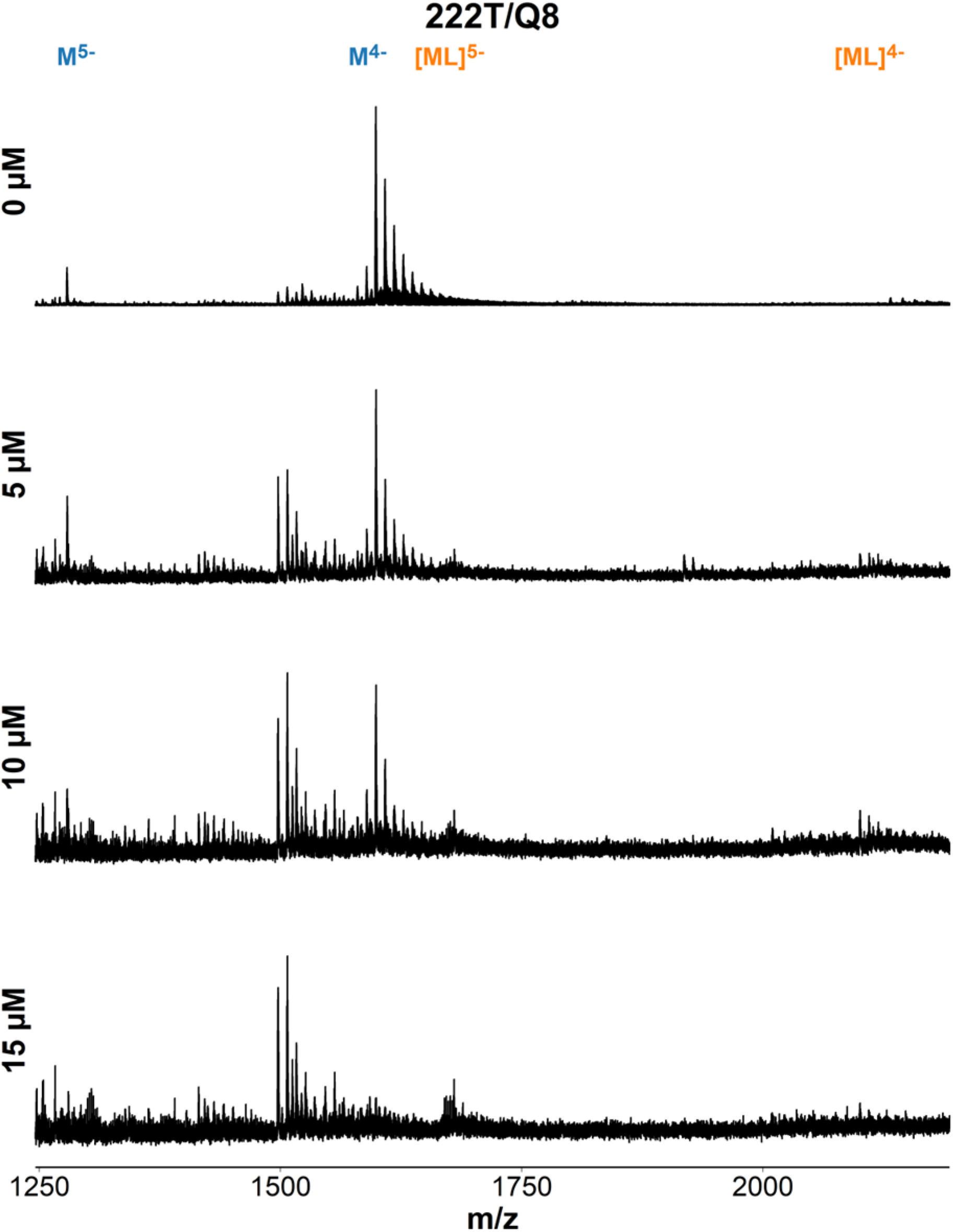
ESI-MS titration of 222T (dT**GGG**TT**GGG**TT**GGG**TT**GGG**T) with foldamer QQQQQQQQ. Samples contain 10 µM DNA, 0-15 µM ligand, 0.5 mM KCl, 100 mM TMAA (pH 6.8).

**Figure S84.**
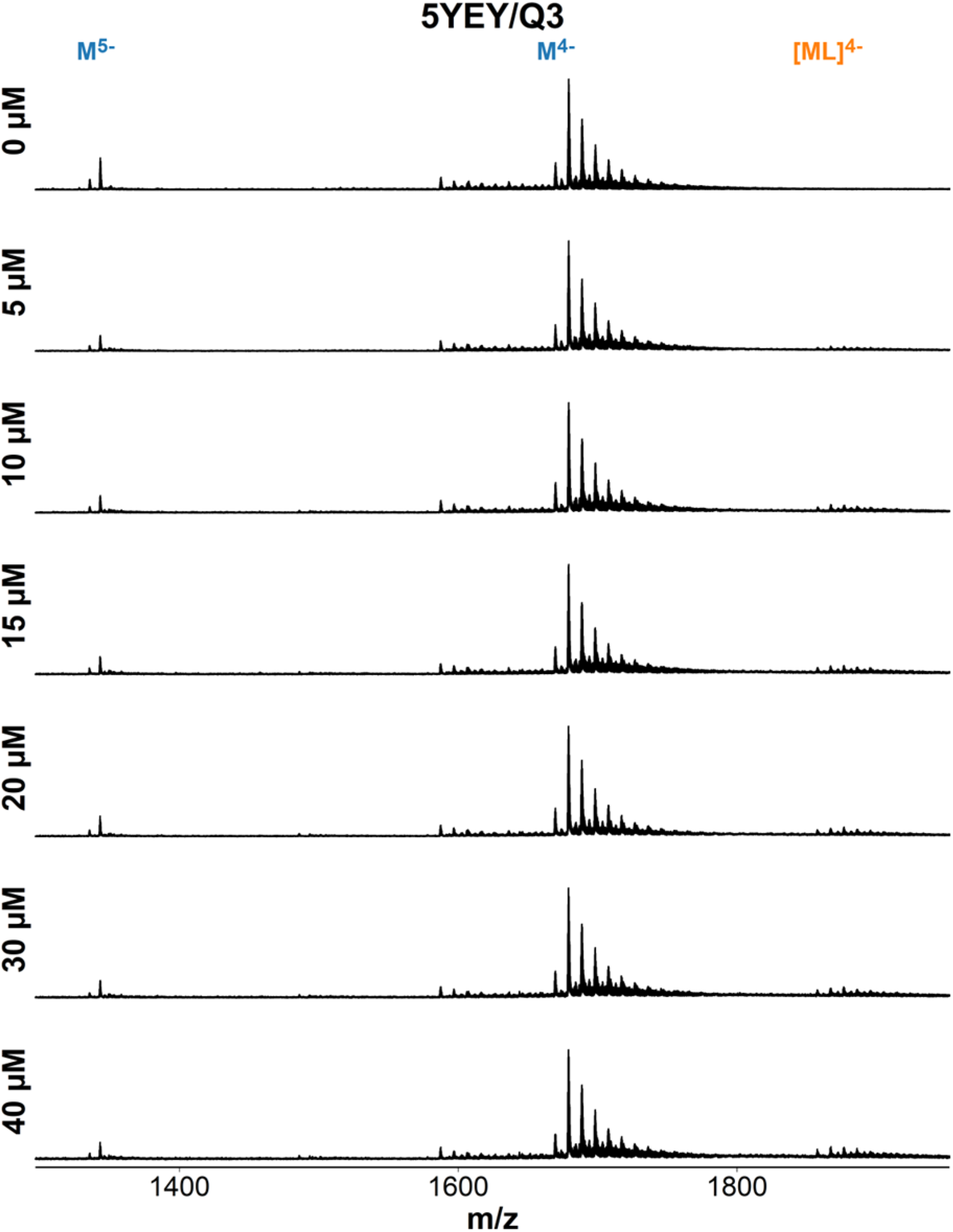
ESI-MS titration of 5YEY (d**GGG**TTA**GGG**TTA**GGG**TTT**GGG**) with foldamer QQQ. Samples contain 10 µM DNA, 0-40 µM ligand, 0.5 mM KCl, 100 mM TMAA (pH 6.8).

**Figure S85.**
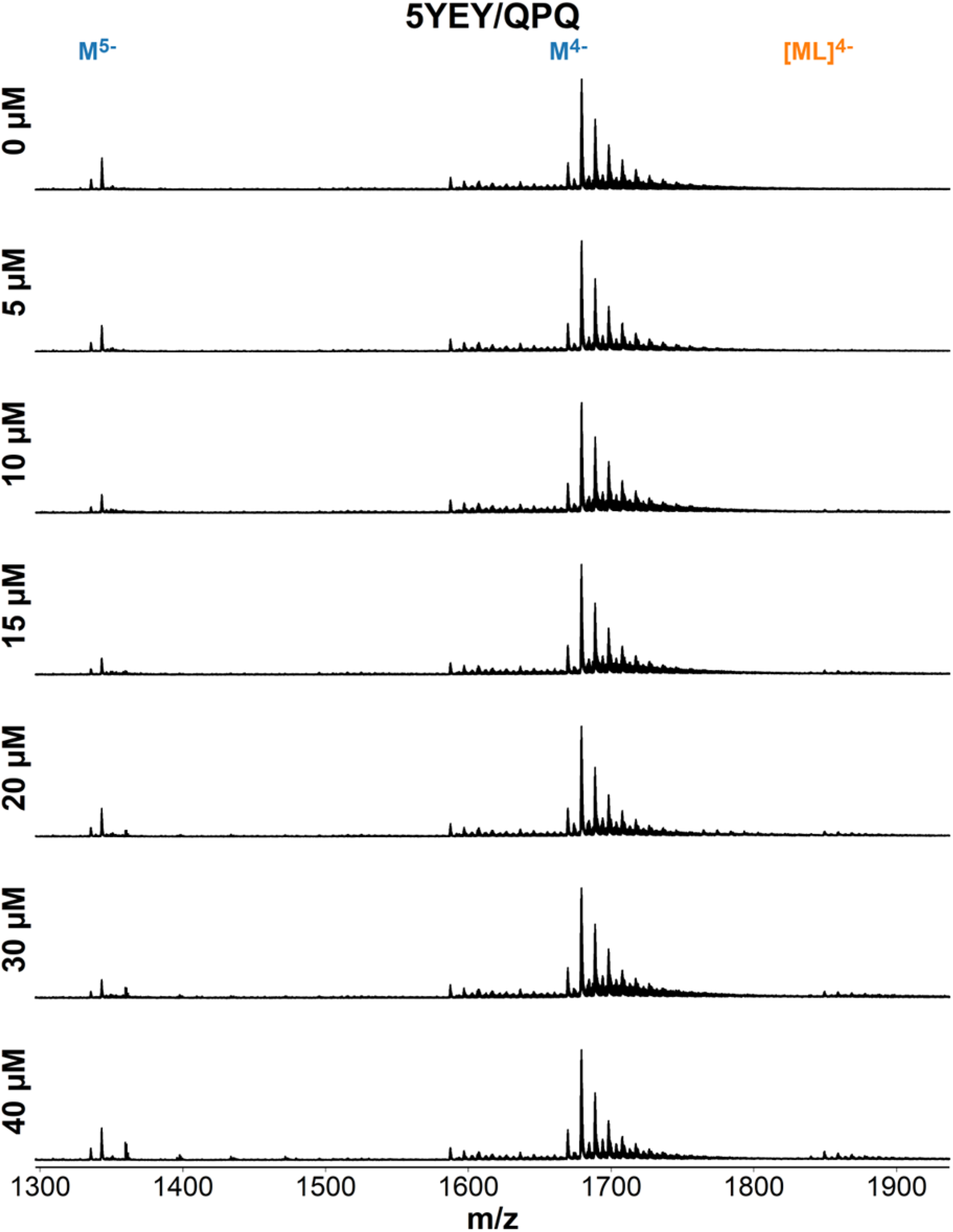
ESI-MS titration of 5YEY (d**GGG**TTA**GGG**TTA**GGG**TTT**GGG**) with foldamer QPQ. Samples contain 10 µM DNA, 0-40 µM ligand, 0.5 mM KCl, 100 mM TMAA (pH 6.8).

**Figure S86.**
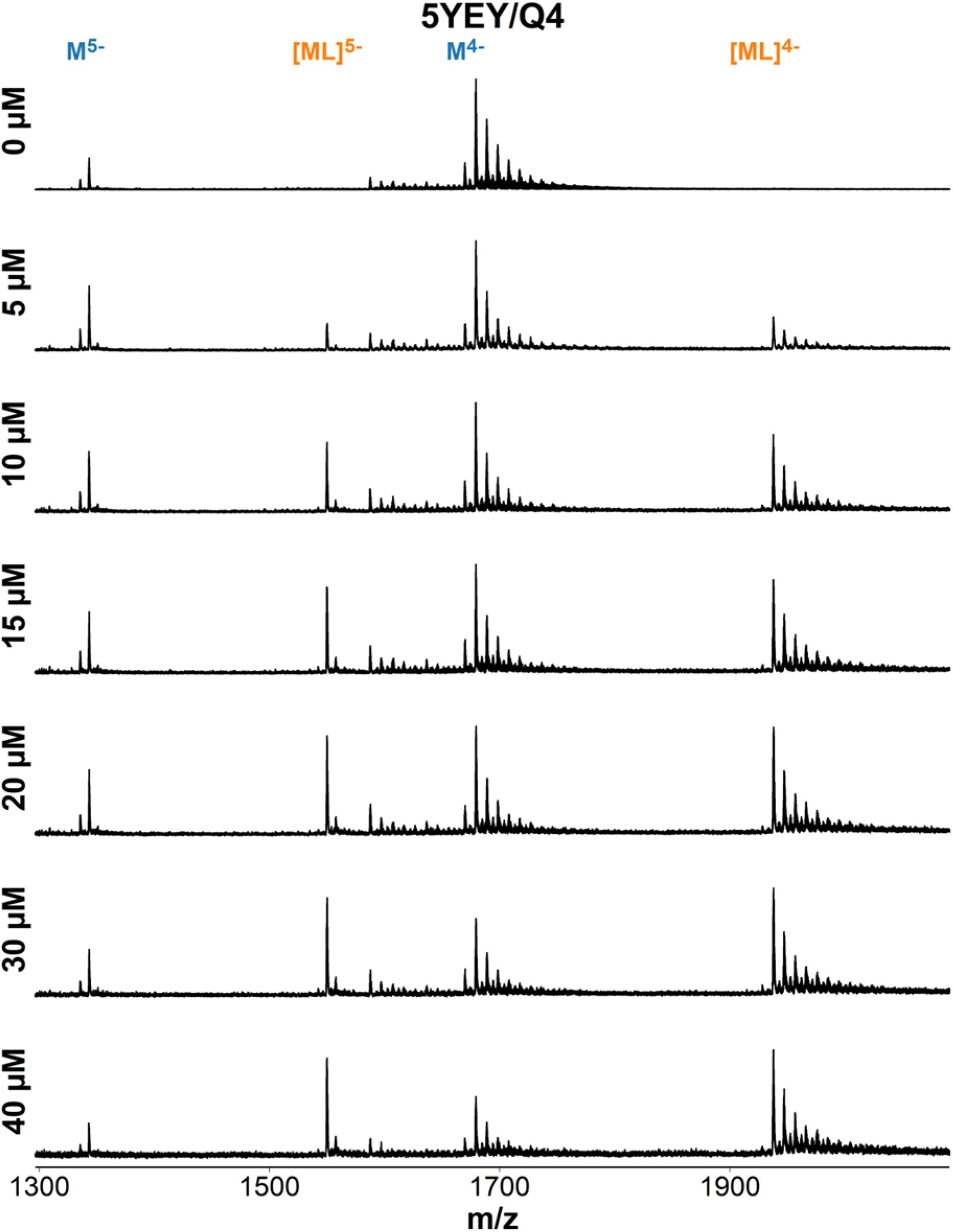
ESI-MS titration of 5YEY (d**GGG**TTA**GGG**TTA**GGG**TTT**GGG**) with foldamer QQQQ. Samples contain 10 µM DNA, 0-40 µM ligand, 0.5 mM KCl, 100 mM TMAA (pH 6.8).

**Figure S87.**
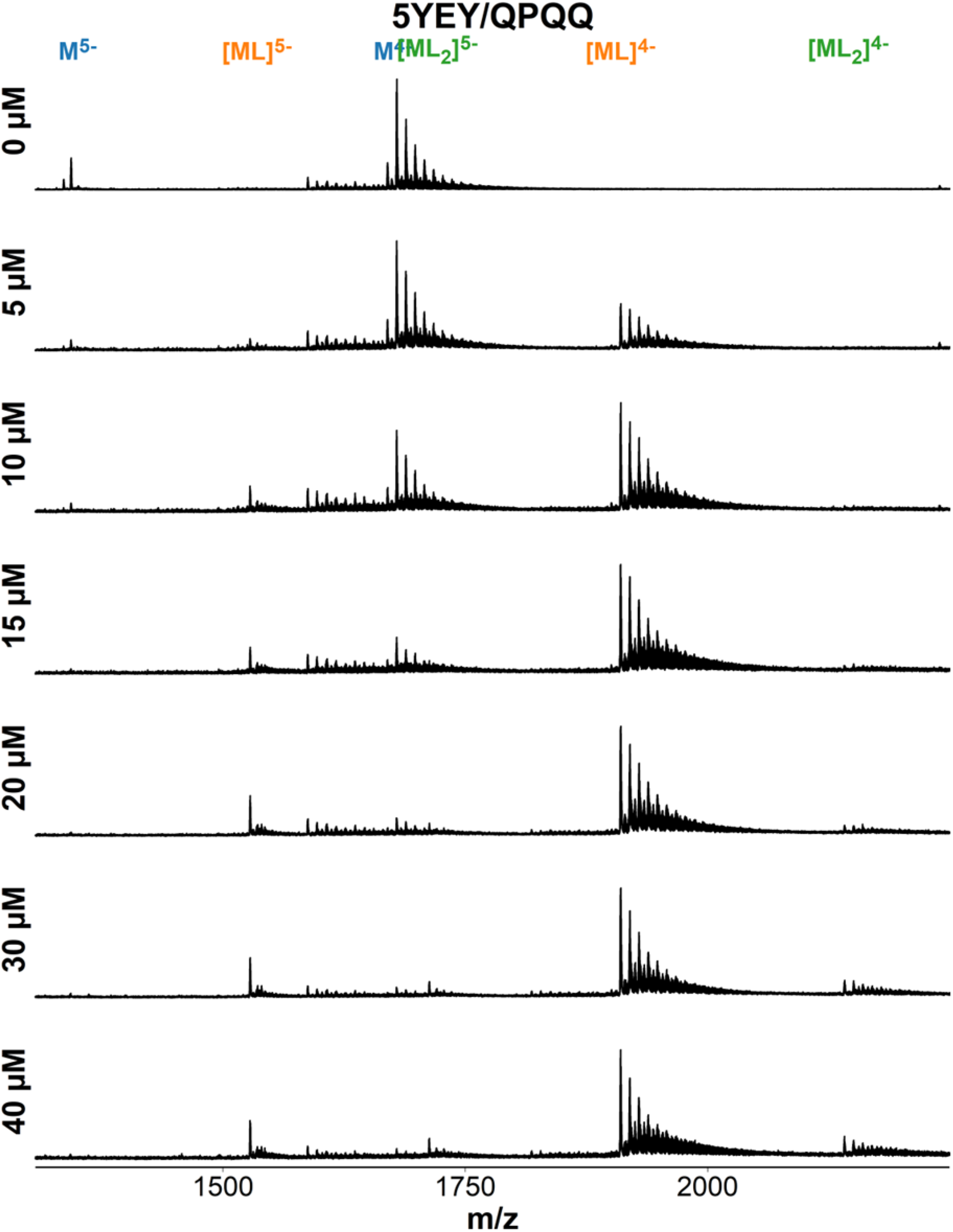
ESI-MS titration of 5YEY (d**GGG**TTA**GGG**TTA**GGG**TTT**GGG**) with foldamer QQPQ. Samples contain 10 µM DNA, 0-40 µM ligand, 0.5 mM KCl, 100 mM TMAA (pH 6.8).

**Figure S88.**
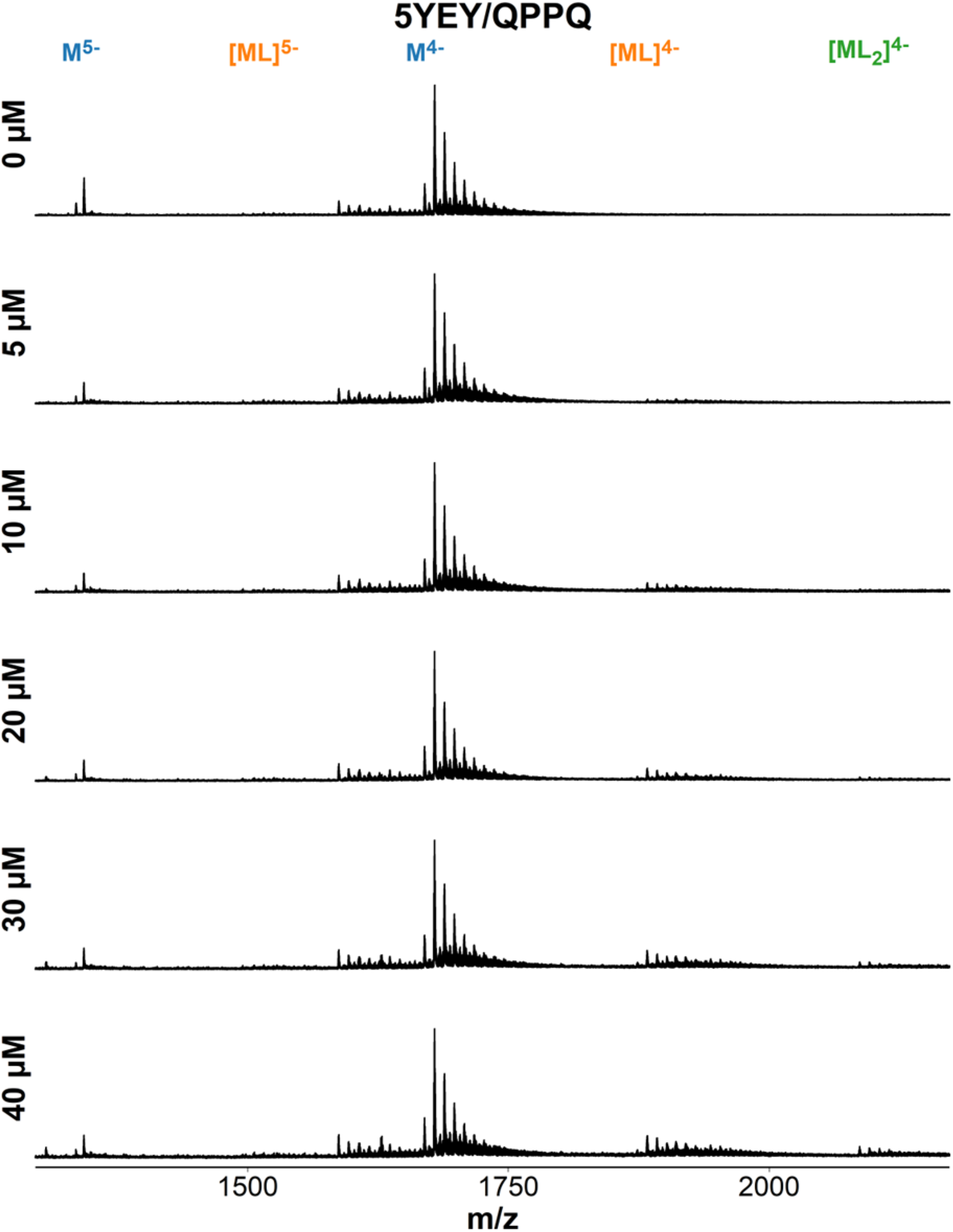
ESI-MS titration of 5YEY (d**GGG**TTA**GGG**TTA**GGG**TTT**GGG**) with foldamer QPPQ. Samples contain 10 µM DNA, 0-40 µM ligand, 0.5 mM KCl, 100 mM TMAA (pH 6.8).

**Figure S89.**
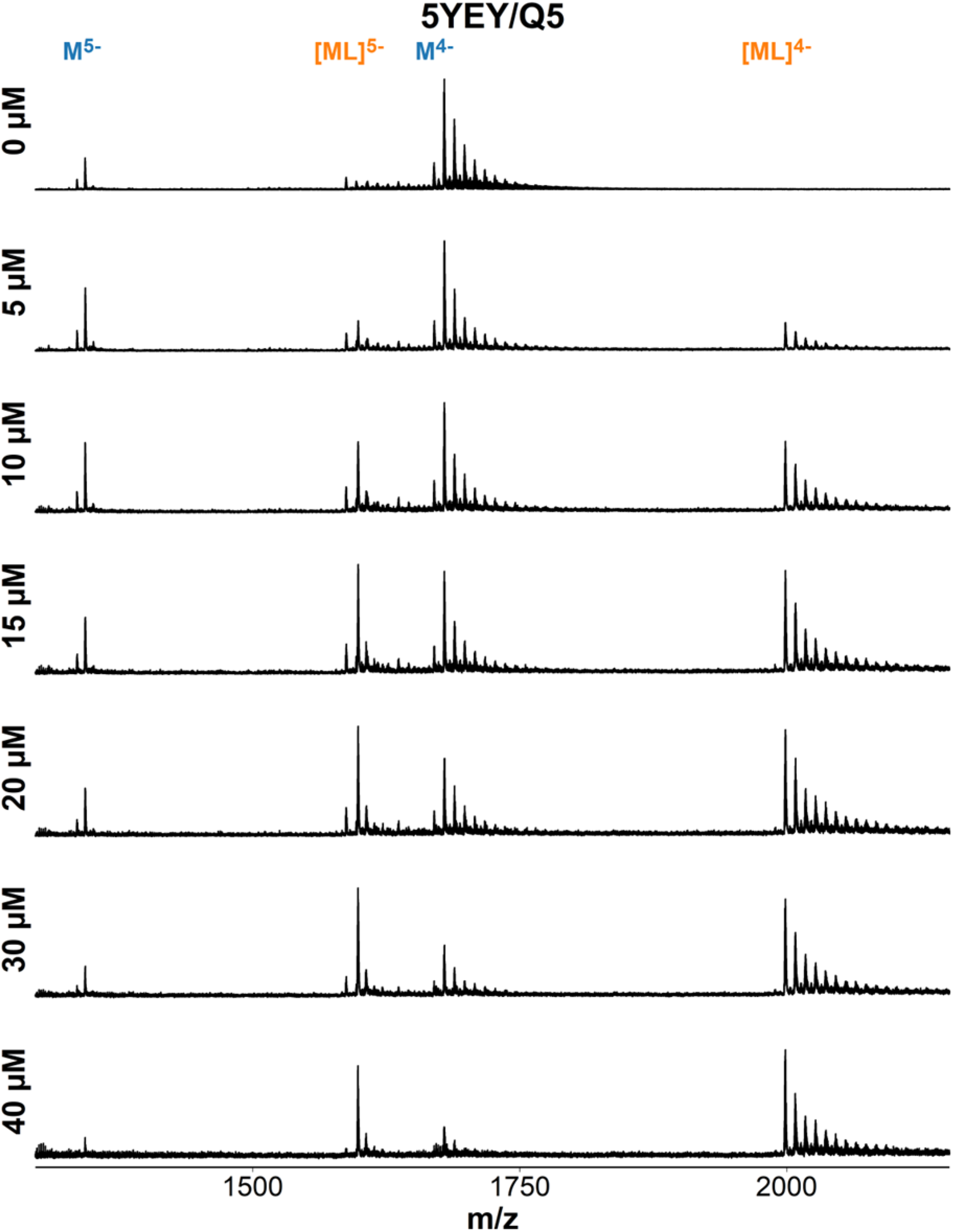
ESI-MS titration of 5YEY (d**GGG**TTA**GGG**TTA**GGG**TTT**GGG**) with foldamer QQQQQ. Samples contain 10 µM DNA, 0-40 µM ligand, 0.5 mM KCl, 100 mM TMAA (pH 6.8).

**Figure S90.**
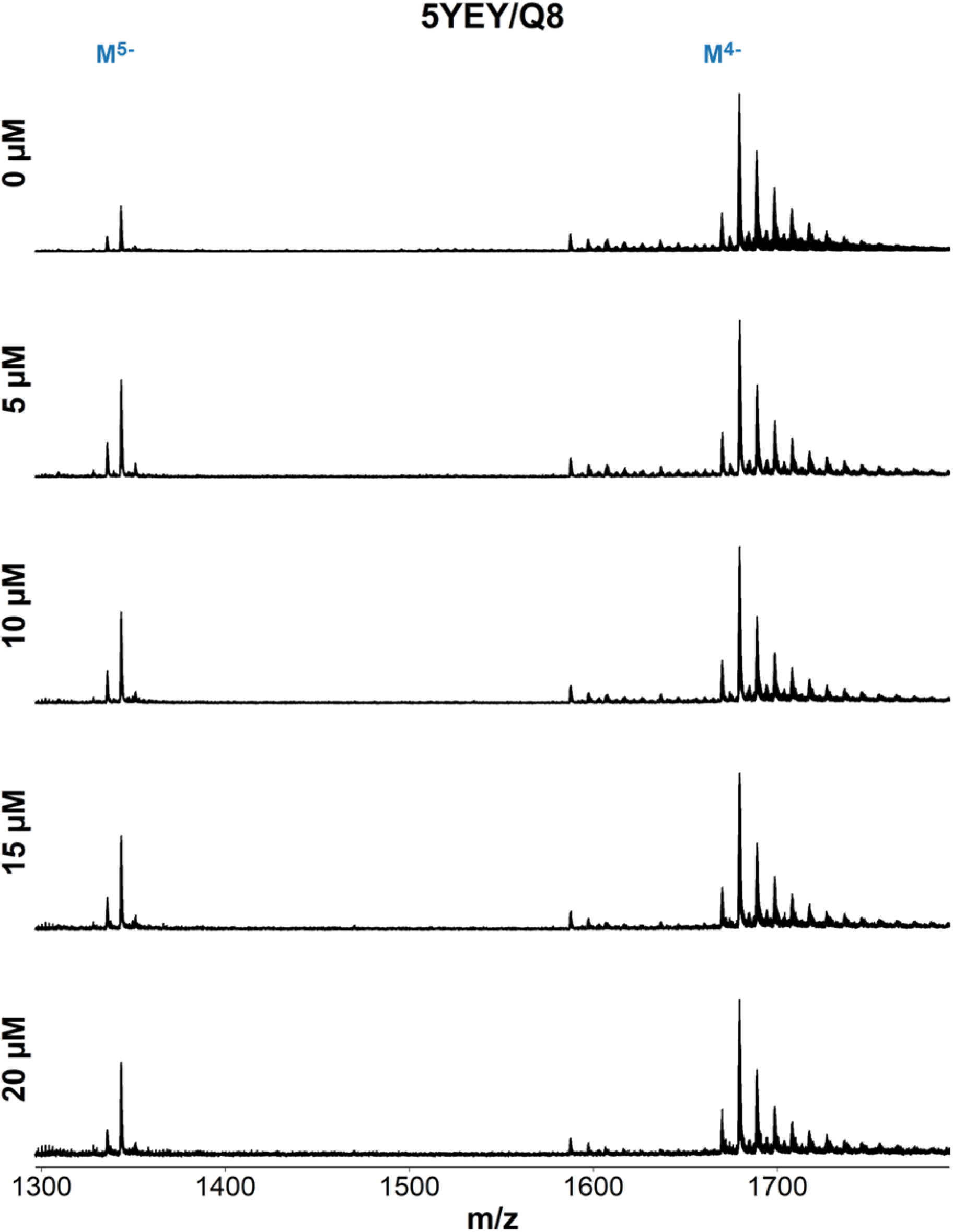
ESI-MS titration of 5YEY (d**GGG**TTA**GGG**TTA**GGG**TTT**GGG**) with foldamer QQQQQQQQ. Samples contain 10 µM DNA, 0-20 µM ligand, 0.5 mM KCl, 100 mM TMAA (pH 6.8).

**Figure S91.**
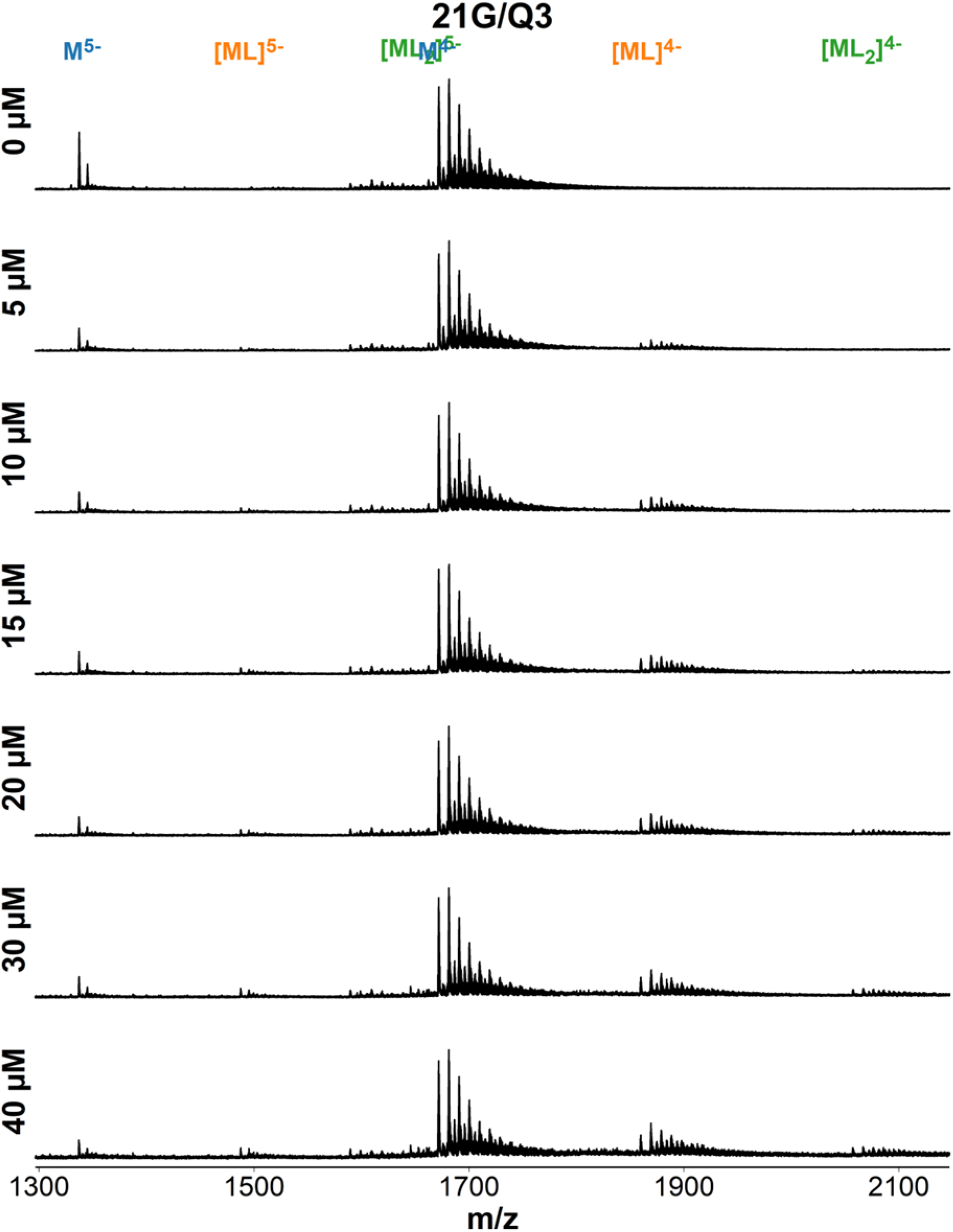
ESI-MS titration of 21G (d**GGG**TTA**GGG**TTA**GGG**TTA**GGG**) with foldamer QQQ. Samples contain 10 µM DNA, 0-40 µM ligand, 0.5 mM KCl, 100 mM TMAA (pH 6.8).

**Figure S92.**
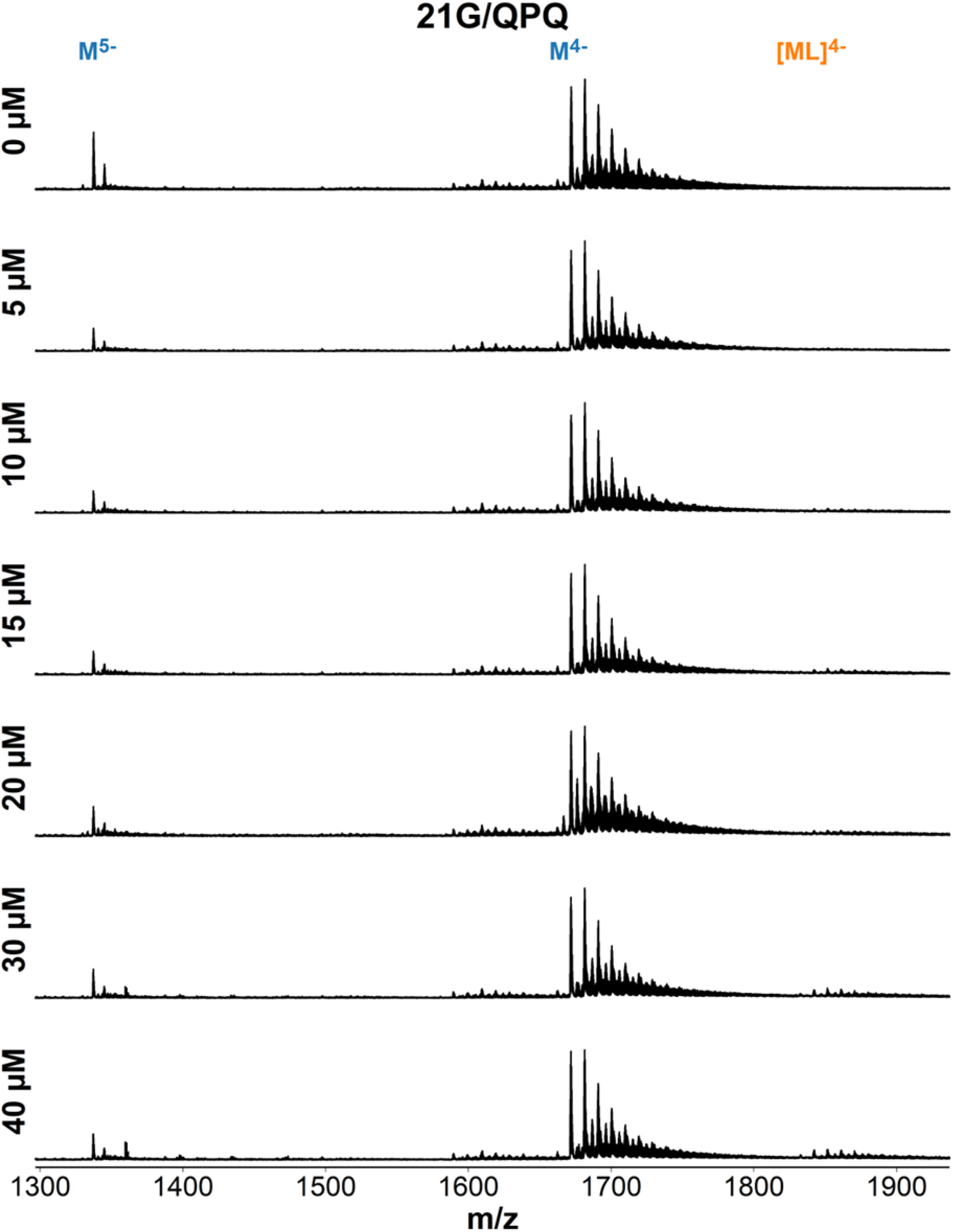
ESI-MS titration of 21G (d**GGG**TTA**GGG**TTA**GGG**TTA**GGG**) with foldamer QPQ. Samples contain 10 µM DNA, 0-40 µM ligand, 0.5 mM KCl, 100 mM TMAA (pH 6.8).

**Figure S93.**
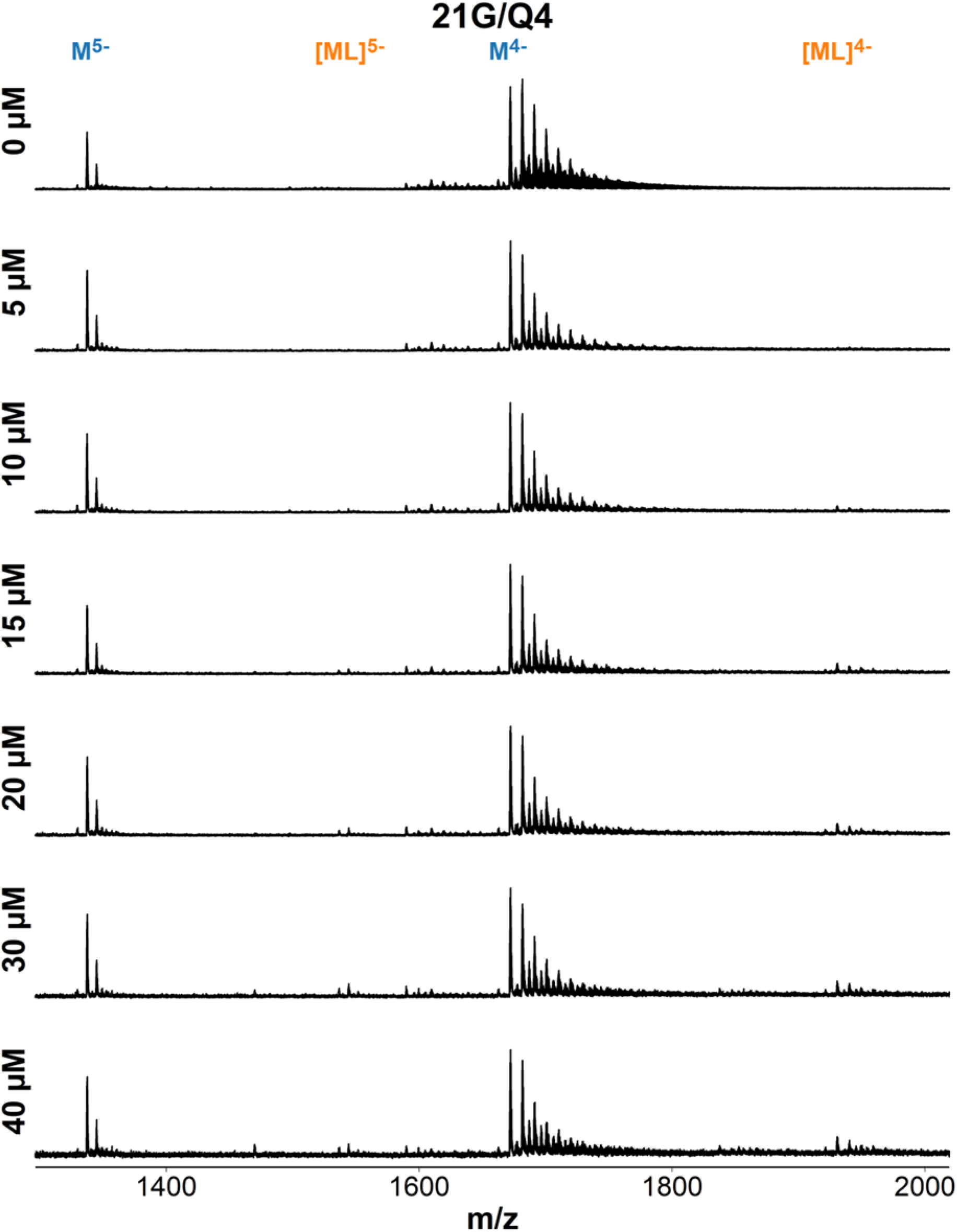
ESI-MS titration of 21G (d**GGG**TTA**GGG**TTA**GGG**TTA**GGG**) with foldamer QQQQ. Samples contain 10 µM DNA, 0-40 µM ligand, 0.5 mM KCl, 100 mM TMAA (pH 6.8).

**Figure S94.**
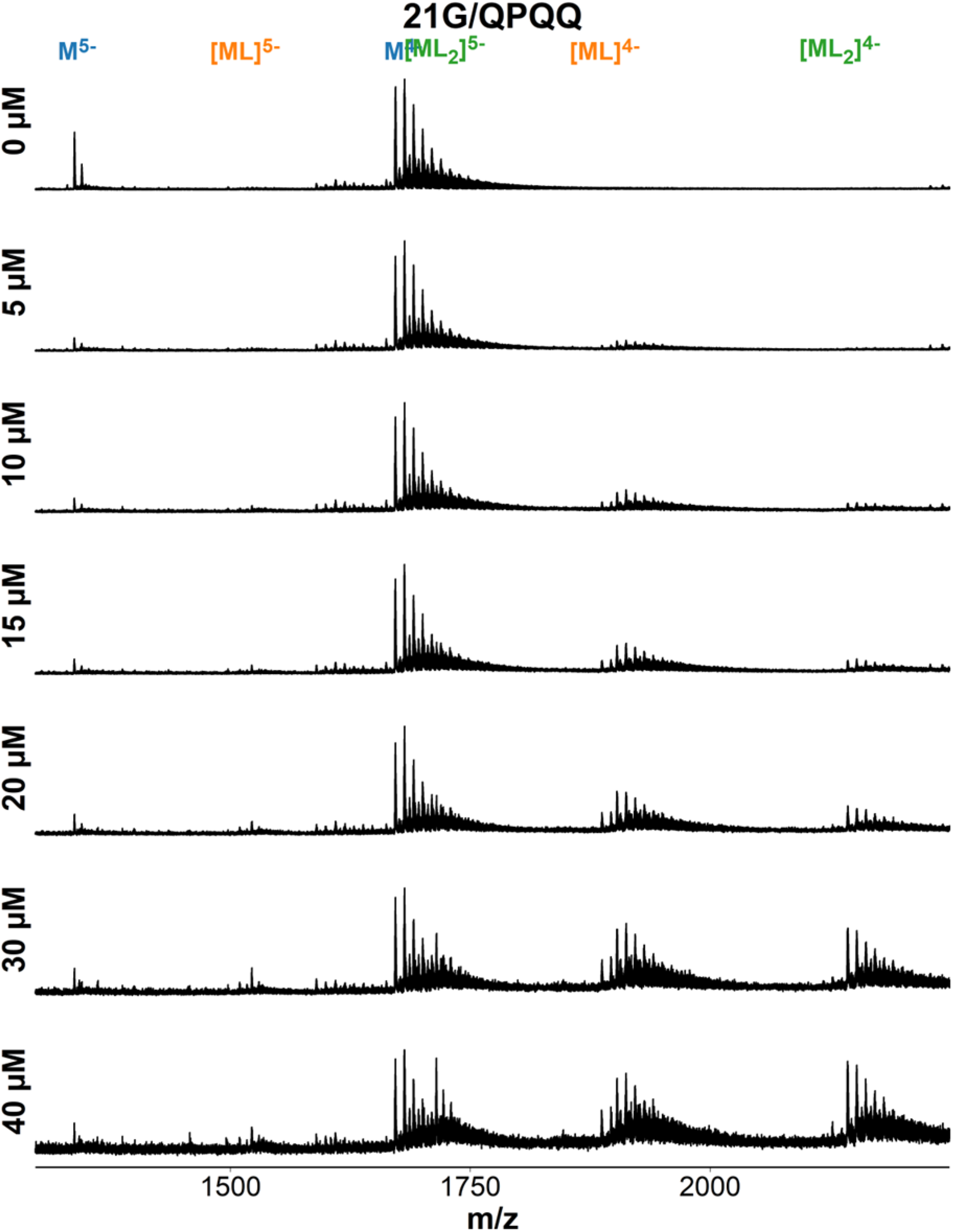
ESI-MS titration of 21G (d**GGG**TTA**GGG**TTA**GGG**TTA**GGG**) with foldamer QQPQ. Samples contain 10 µM DNA, 0-40 µM ligand, 0.5 mM KCl, 100 mM TMAA (pH 6.8).

**Figure S95.**
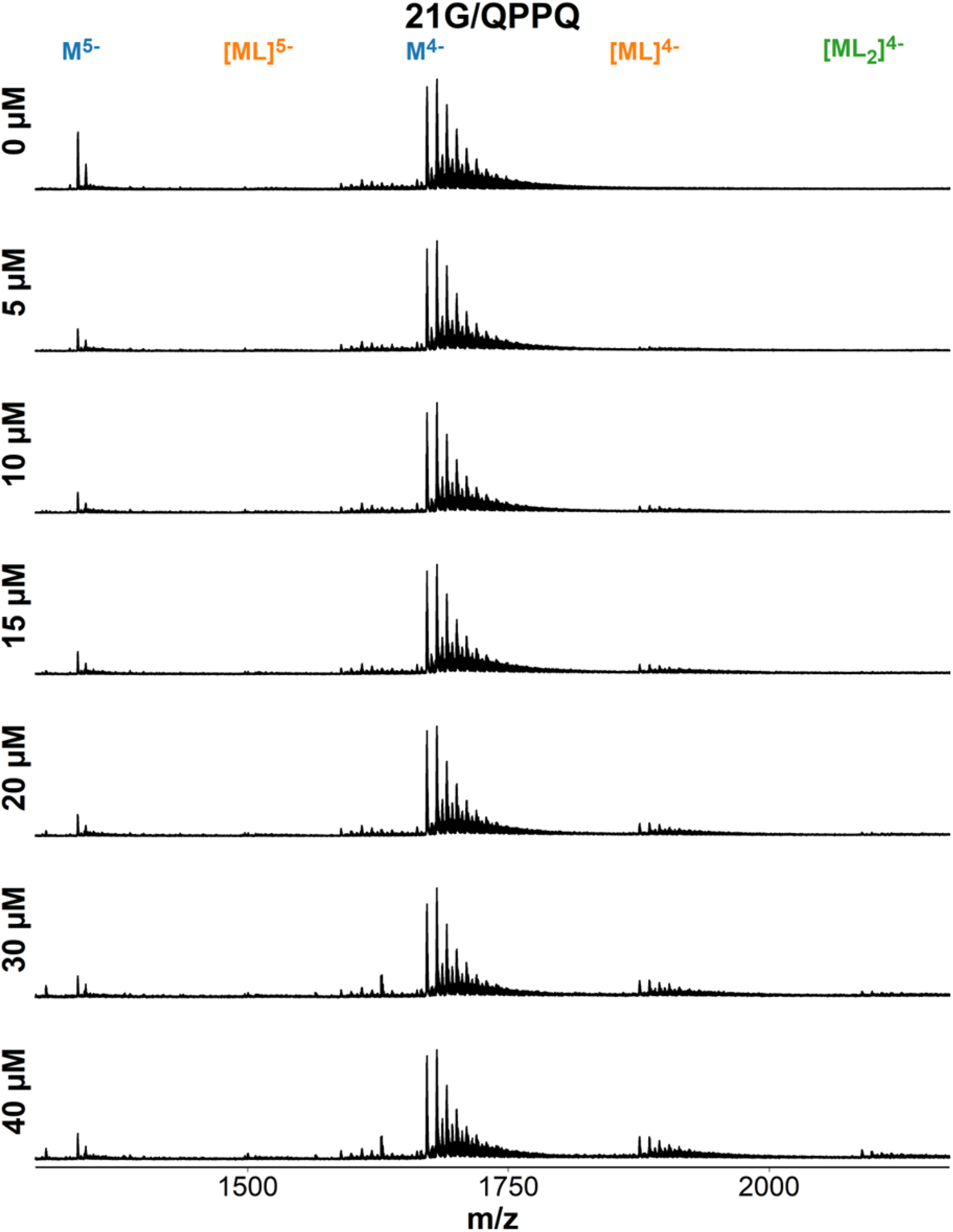
ESI-MS titration of 21G (d**GGG**TTA**GGG**TTA**GGG**TTA**GGG**) with foldamer QPPQ. Samples contain 10 µM DNA, 0-40 µM ligand, 0.5 mM KCl, 100 mM TMAA (pH 6.8).

**Figure S96.**
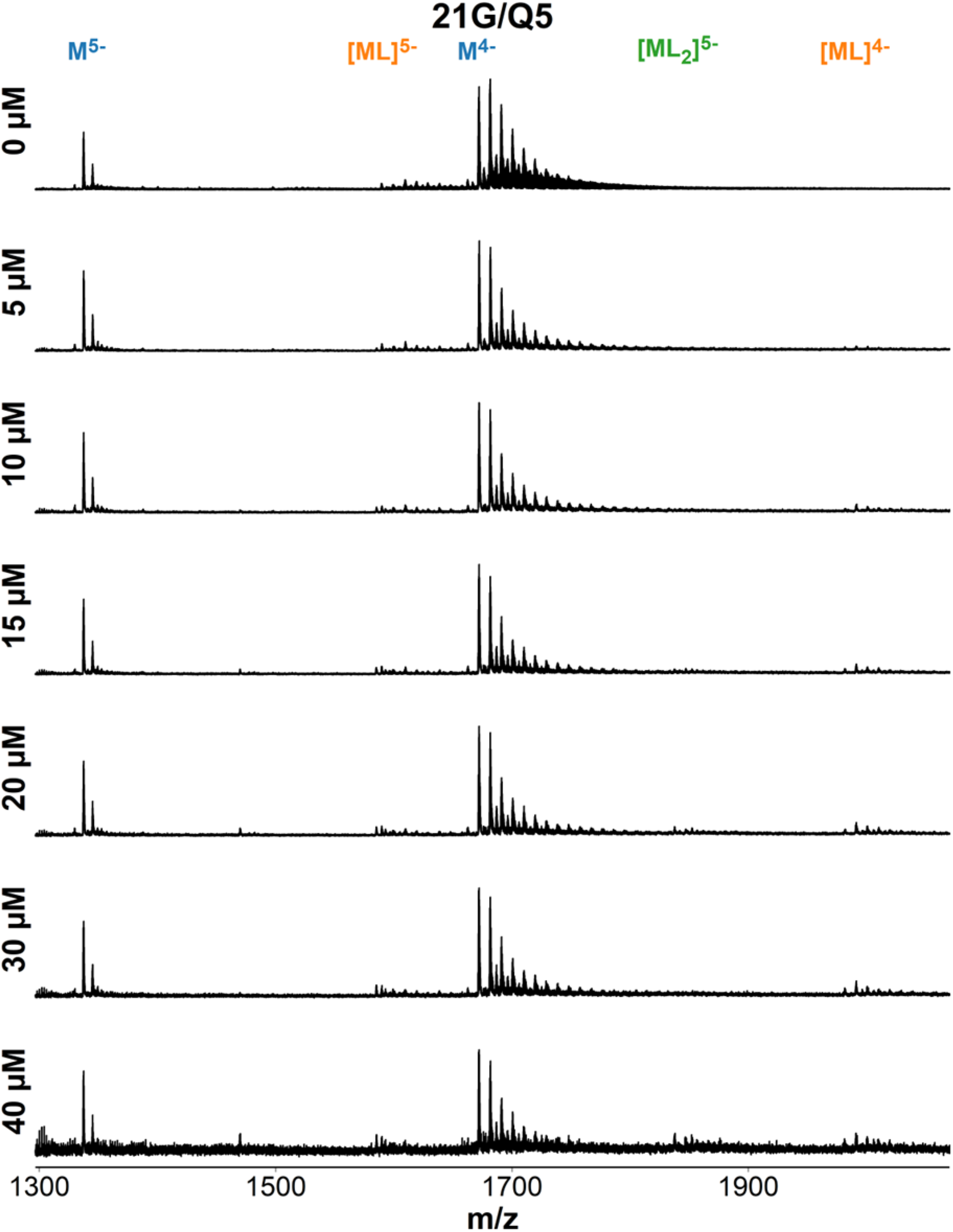
ESI-MS titration of 21G (d**GGG**TTA**GGG**TTA**GGG**TTA**GGG**) with foldamer QQQQQ. Samples contain 10 µM DNA, 0-40 µM ligand, 0.5 mM KCl, 100 mM TMAA (pH 6.8).

**Figure S97.**
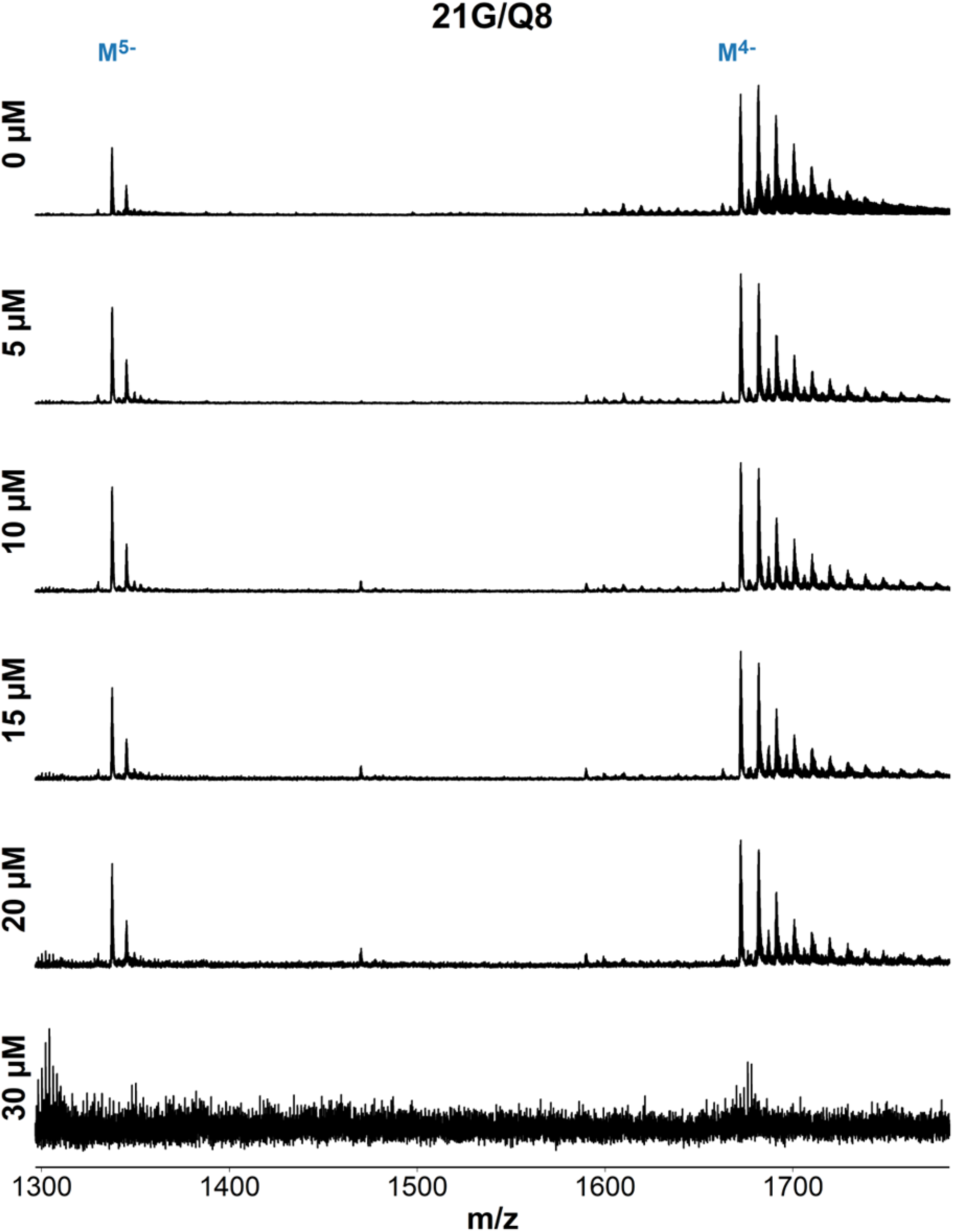
ESI-MS titration of 21G (d**GGG**TTA**GGG**TTA**GGG**TTA**GGG**) with foldamer QQQQQQQQ. Samples contain 10 µM DNA, 0-30 µM ligand, 0.5 mM KCl, 100 mM TMAA (pH 6.8).

**Figure S98.**
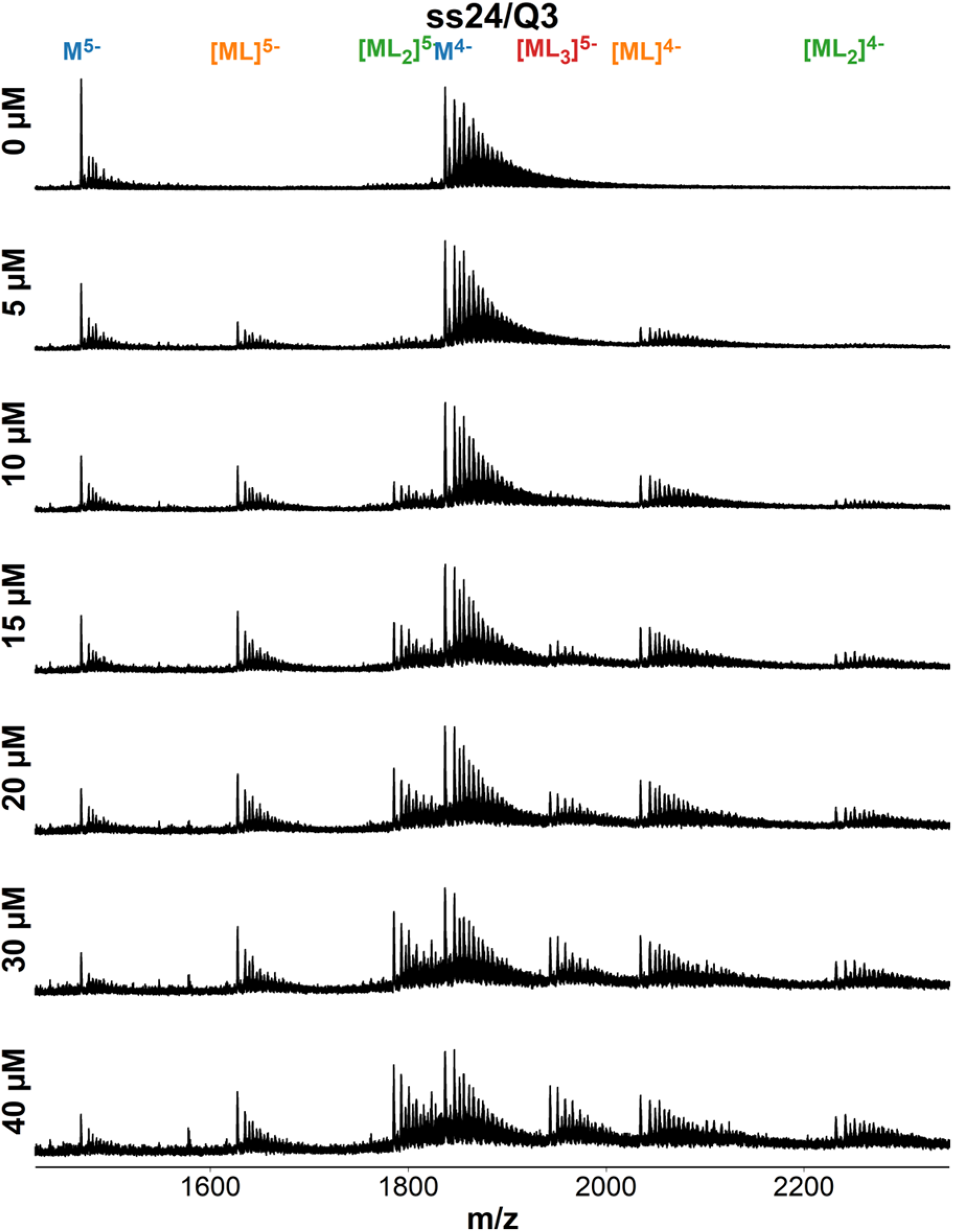
ESI-MS titration of ss24 (dTGCCATGCTACTGAGATGACGCTA) with foldamer QQQ. Samples contain 10 µM DNA, 0-40 µM ligand, 0.5 mM KCl, 100 mM TMAA (pH 6.8).

**Figure S99.**
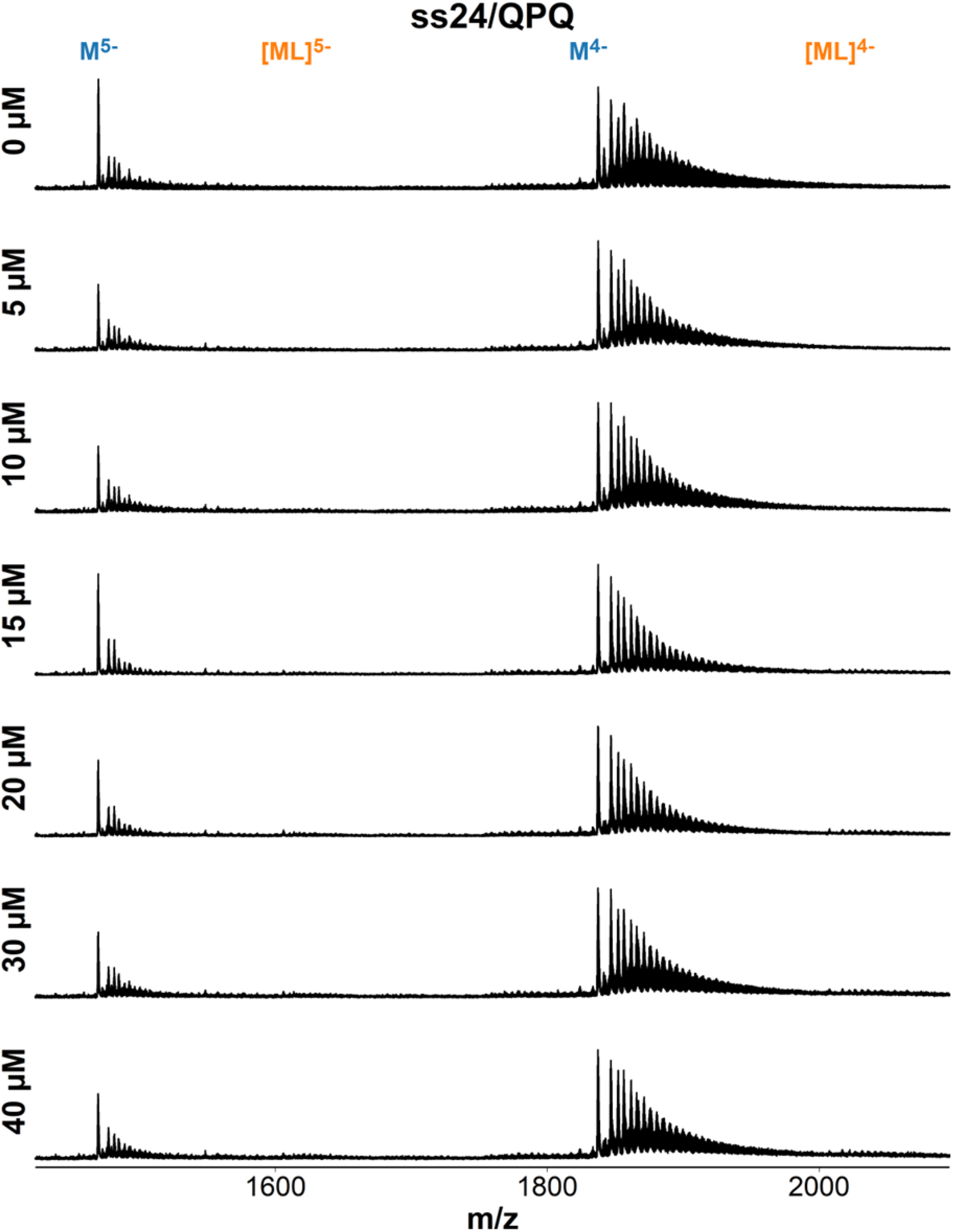
ESI-MS titration of ss24 (dTGCCATGCTACTGAGATGACGCTA) with foldamer QPQ. Samples contain 10 µM DNA, 0-40 µM ligand, 0.5 mM KCl, 100 mM TMAA (pH 6.8).

**Figure S100.**
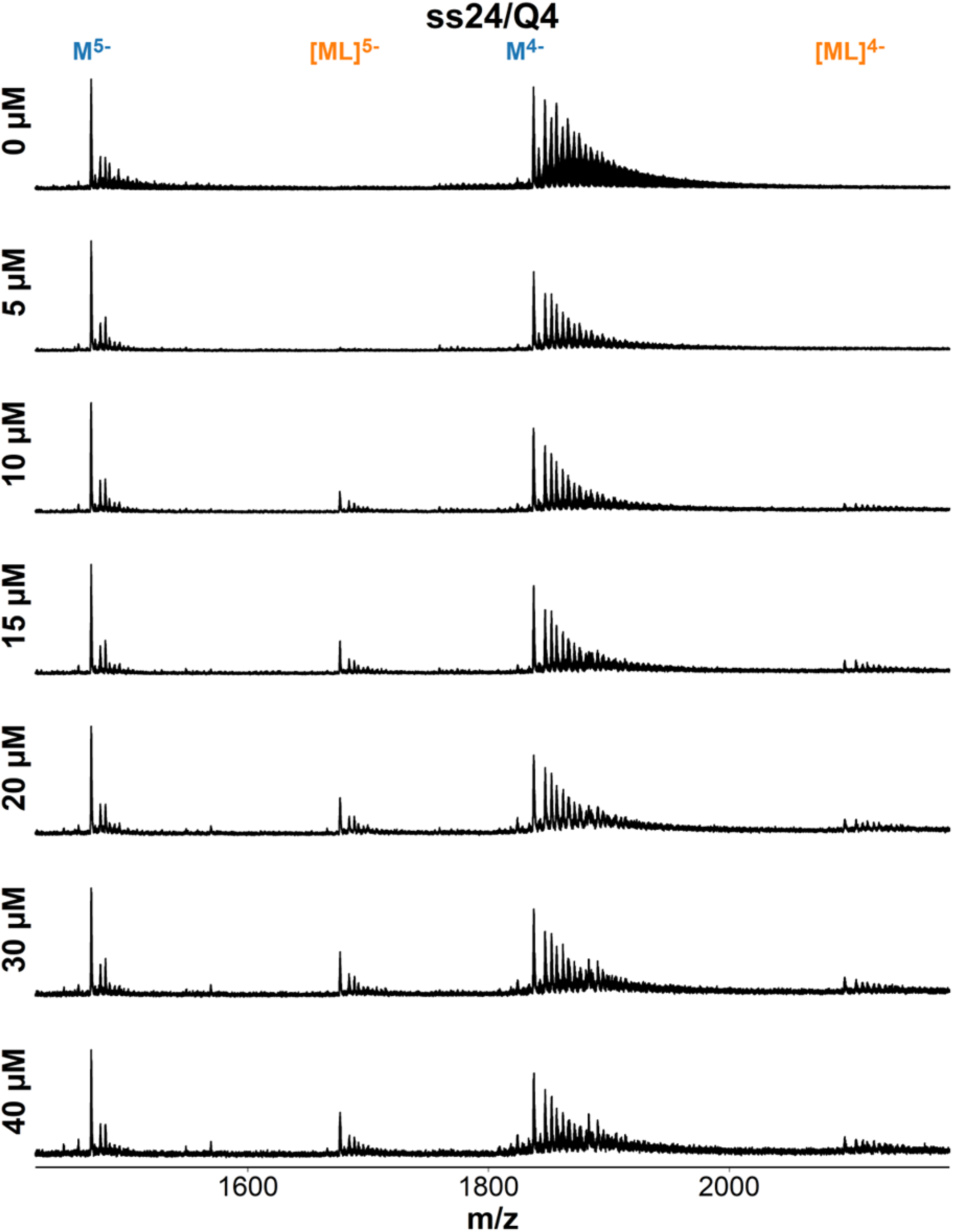
ESI-MS titration of ss24 (dTGCCATGCTACTGAGATGACGCTA) with foldamer QQQQ. Samples contain 10 µM DNA, 0-40 µM ligand, 0.5 mM KCl, 100 mM TMAA (pH 6.8).

**Figure S101.**
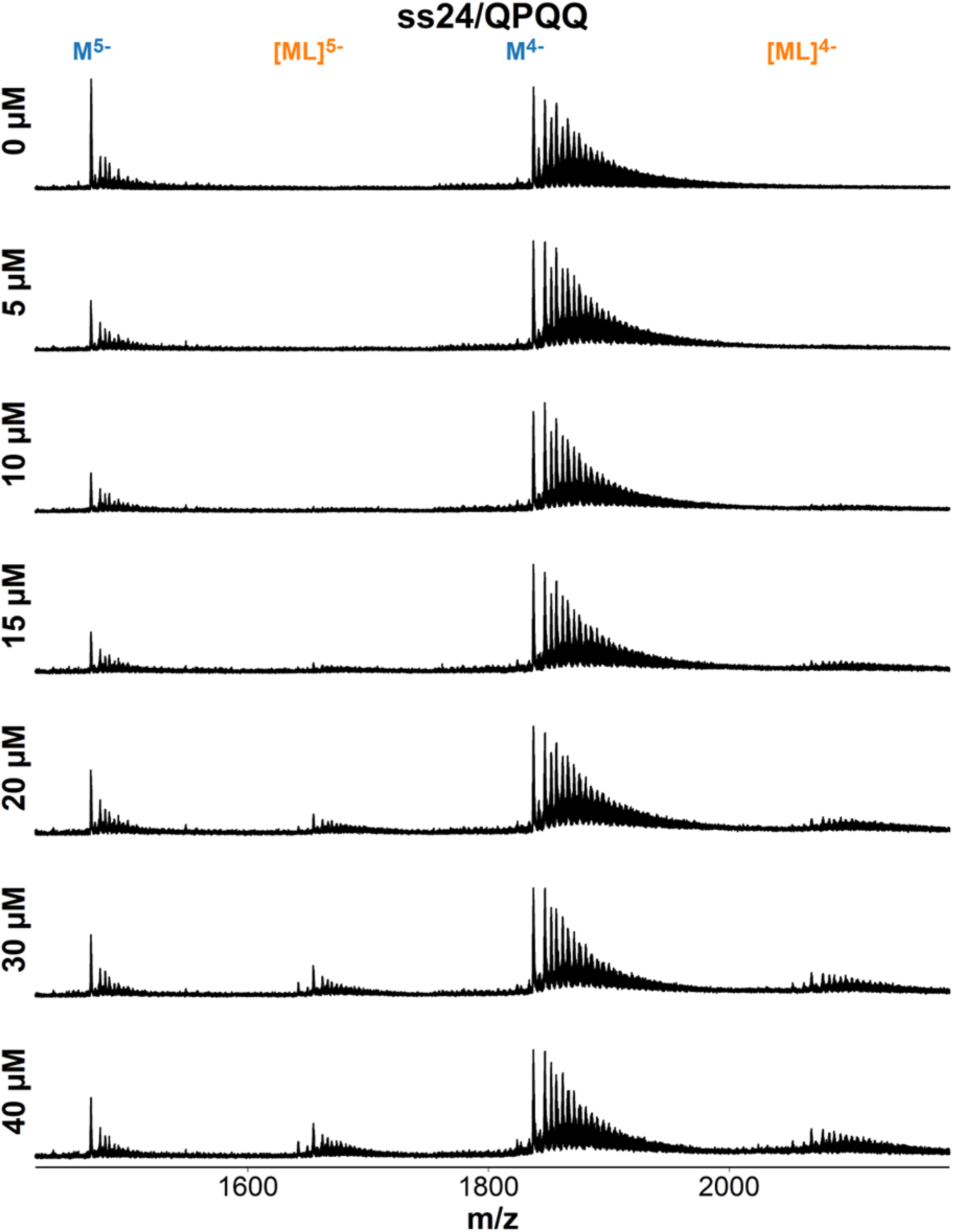
ESI-MS titration of ss24 (dTGCCATGCTACTGAGATGACGCTA) with foldamer QQPQ. Samples contain 10 µM DNA, 0-40 µM ligand, 0.5 mM KCl, 100 mM TMAA (pH 6.8).

**Figure S102.**
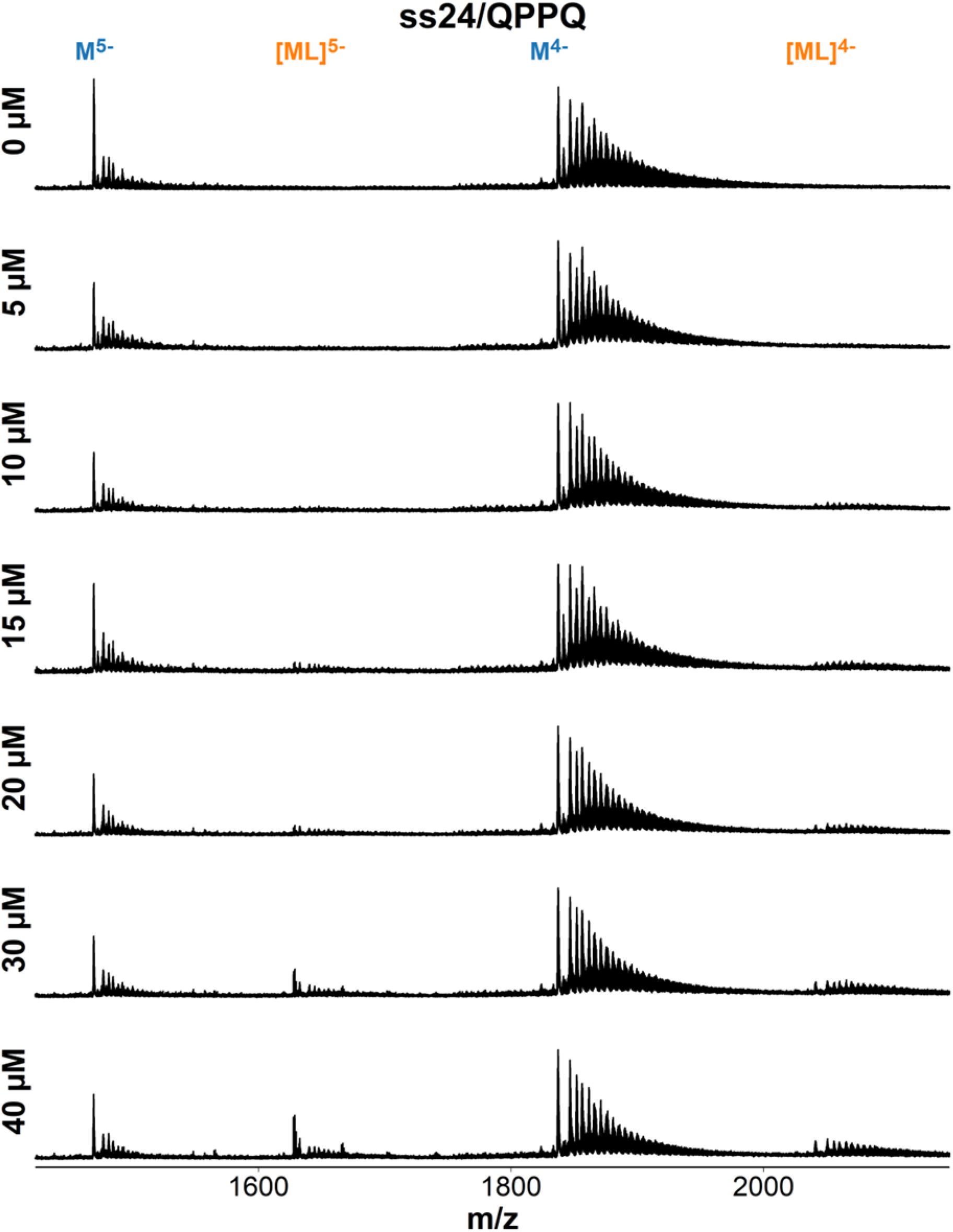
ESI-MS titration of ss24 (dTGCCATGCTACTGAGATGACGCTA) with foldamer QPPQ. Samples contain 10 µM DNA, 0-40 µM ligand, 0.5 mM KCl, 100 mM TMAA (pH 6.8).

**Figure S103.**
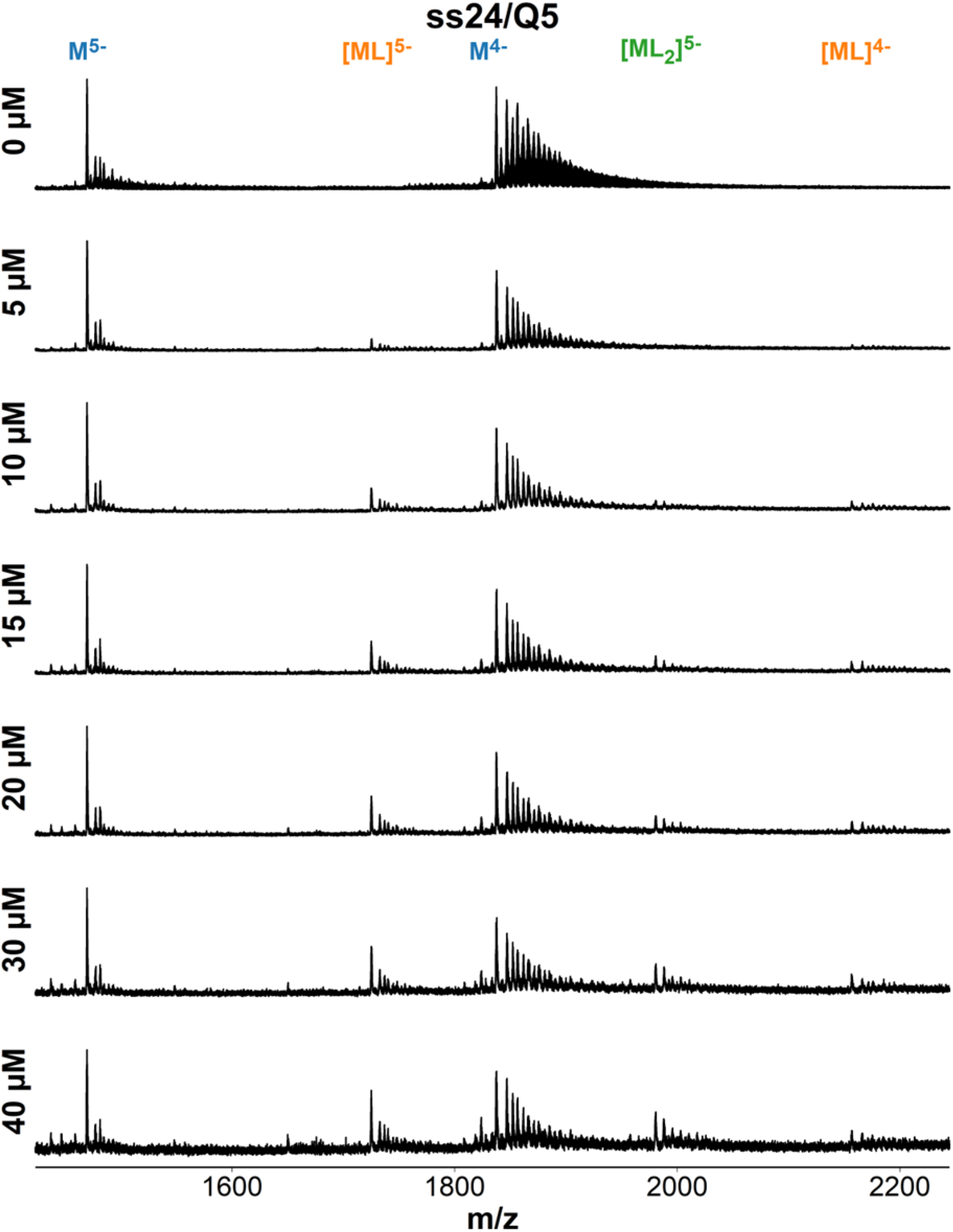
ESI-MS titration of ss24 (dTGCCATGCTACTGAGATGACGCTA) with foldamer QQQQQ. Samples contain 10 µM DNA, 0-40 µM ligand, 0.5 mM KCl, 100 mM TMAA (pH 6.8).

**Figure S104.**
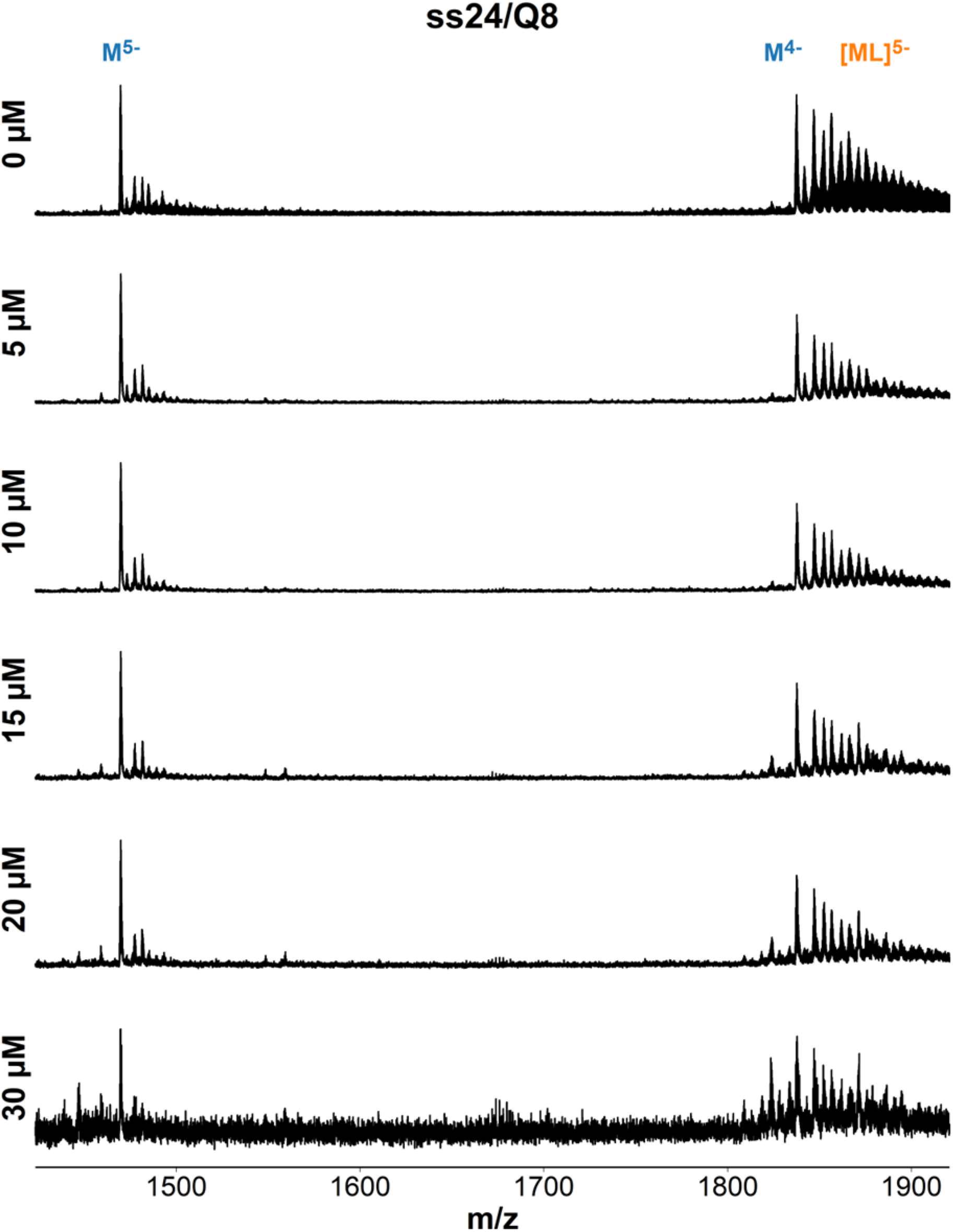
ESI-MS titration of ss24 (dTGCCATGCTACTGAGATGACGCTA) with foldamer QQQQQQQQ. Samples contain 10 µM DNA, 0-30 µM ligand, 0.5 mM KCl, 100 mM TMAA (pH 6.8).

**Figure S105.**
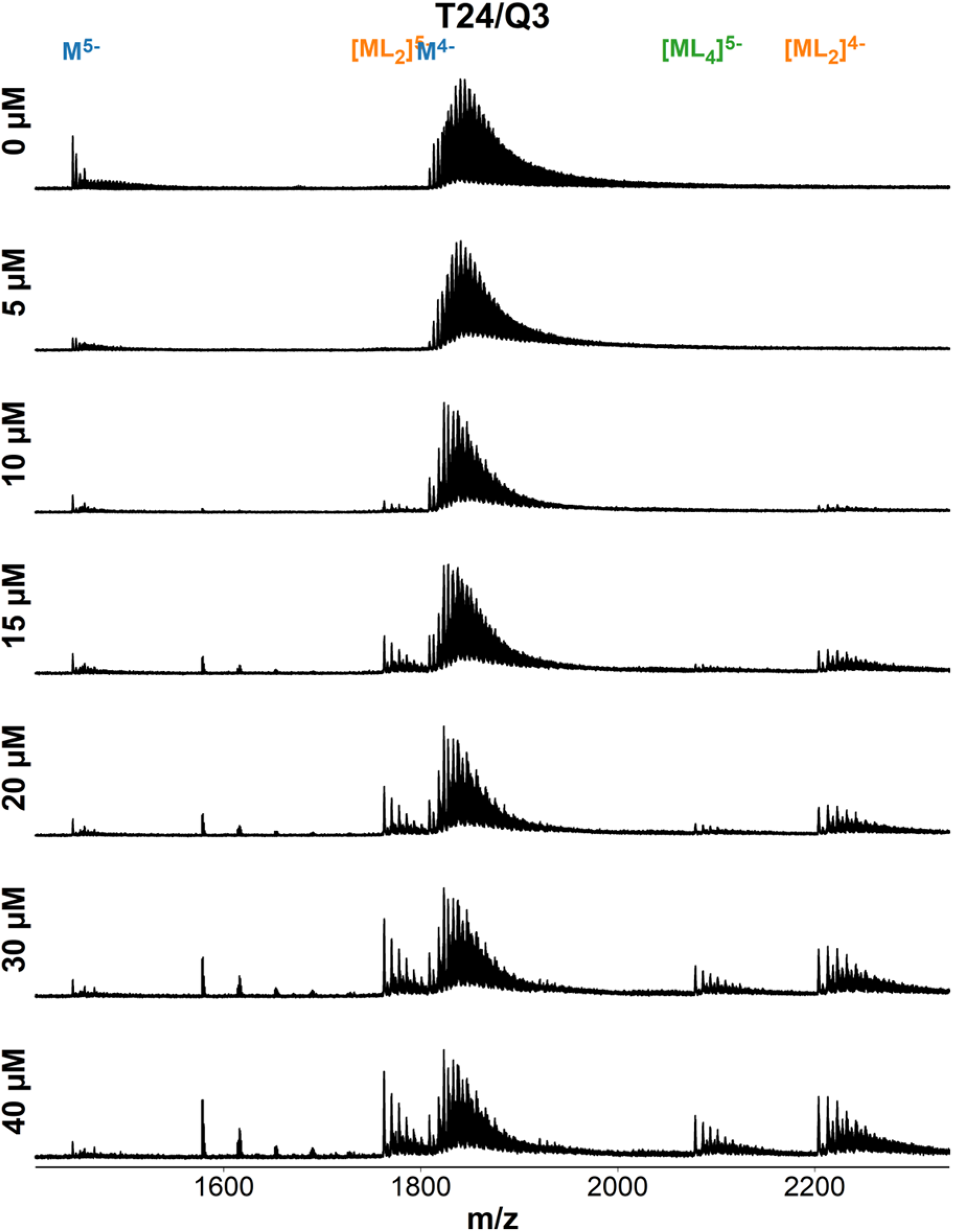
ESI-MS titration of T24 (dTTTTTTTTTTTTTTTTTTTTTTTT) with foldamer QQQ. Samples contain 10 µM DNA, 0-40 µM ligand, 0.5 mM KCl, 100 mM TMAA (pH 6.8).

**Figure S106.**
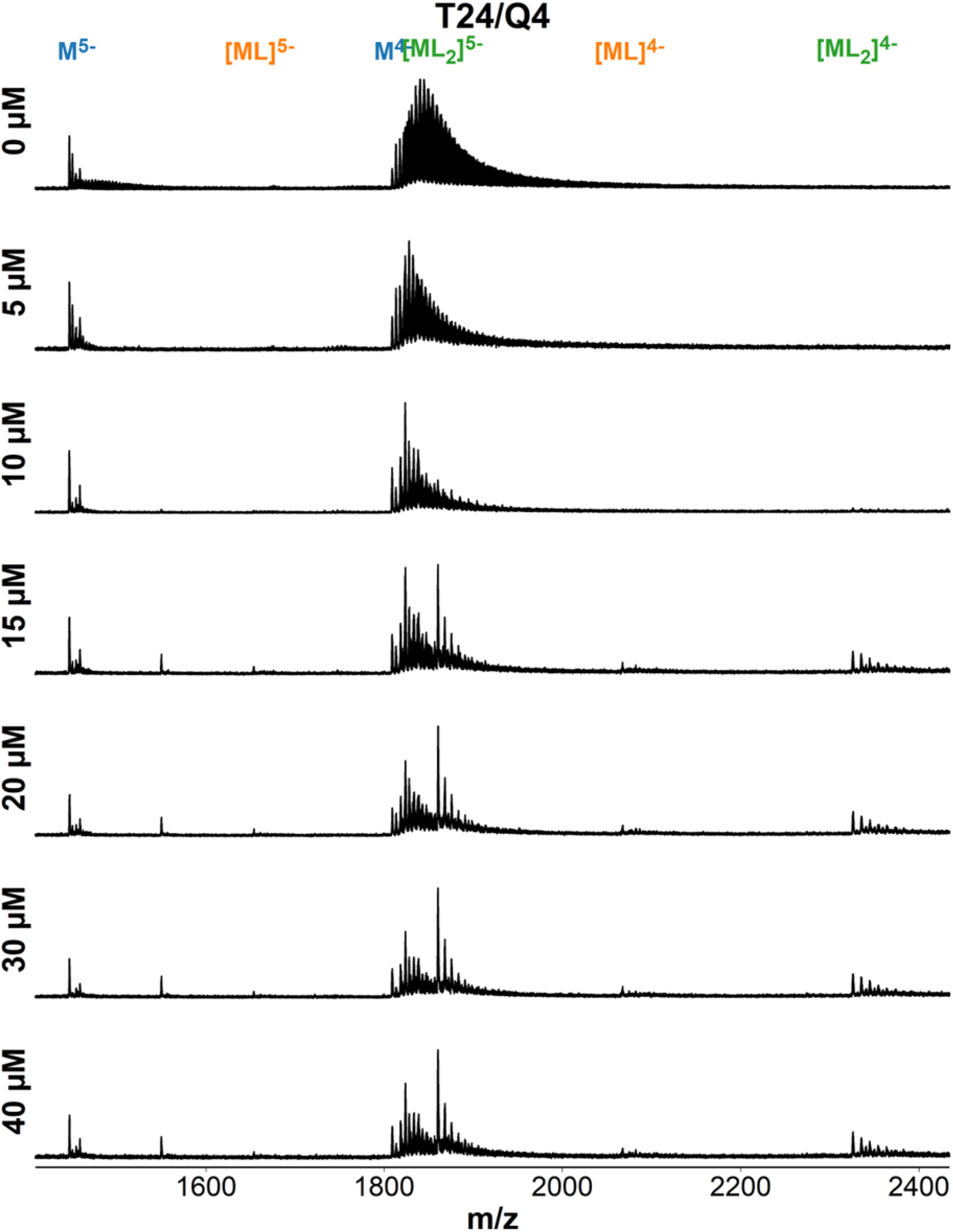
ESI-MS titration of T24 (dTTTTTTTTTTTTTTTTTTTTTTTT) with foldamer QQQQ. Samples contain 10 µM DNA, 0-40 µM ligand, 0.5 mM KCl, 100 mM TMAA (pH 6.8).

**Figure S107.**
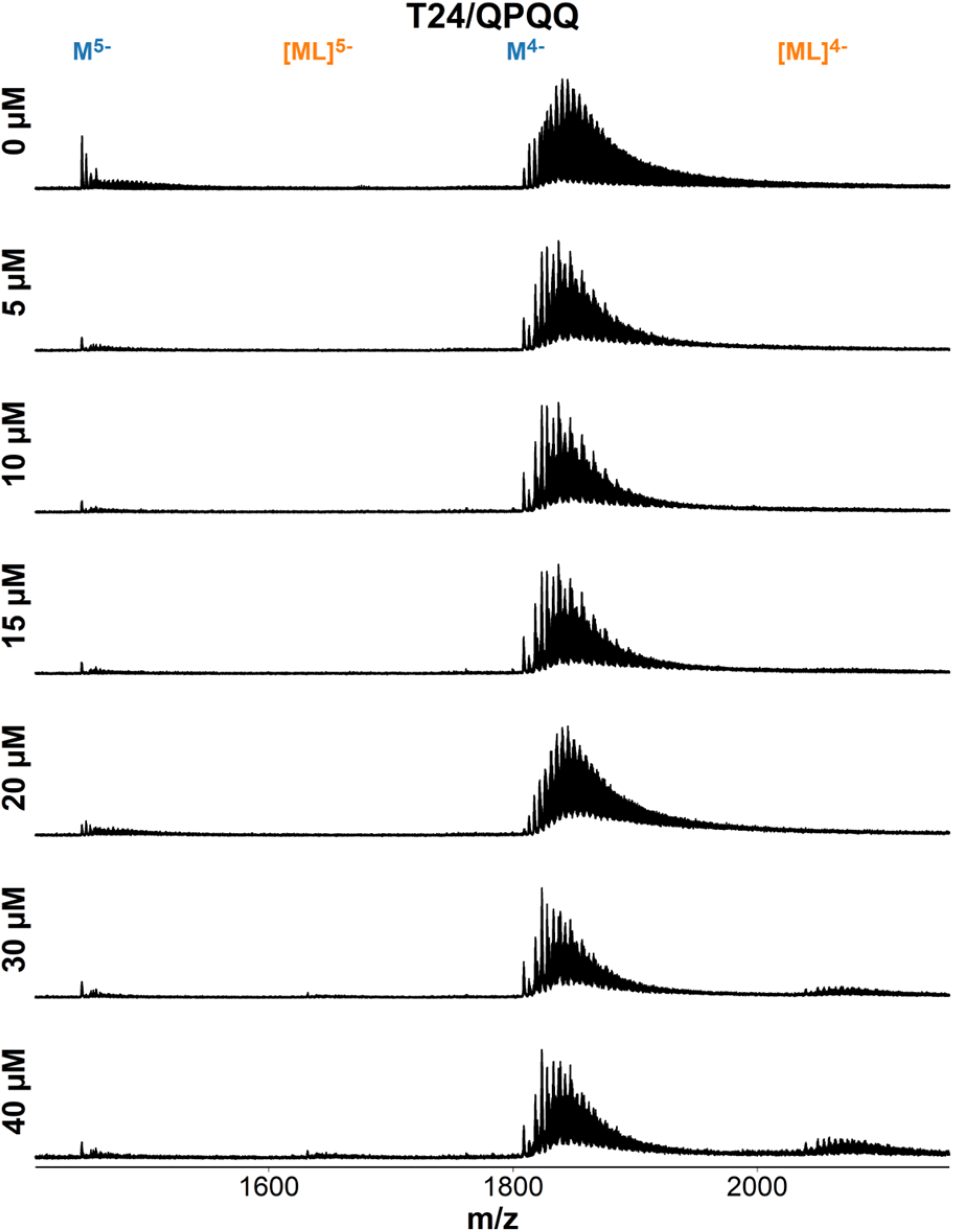
ESI-MS titration of T24 (dTTTTTTTTTTTTTTTTTTTTTTTT) with foldamer QQPQ. Samples contain 10 µM DNA, 0-40 µM ligand, 0.5 mM KCl, 100 mM TMAA (pH 6.8).

**Figure S108.**
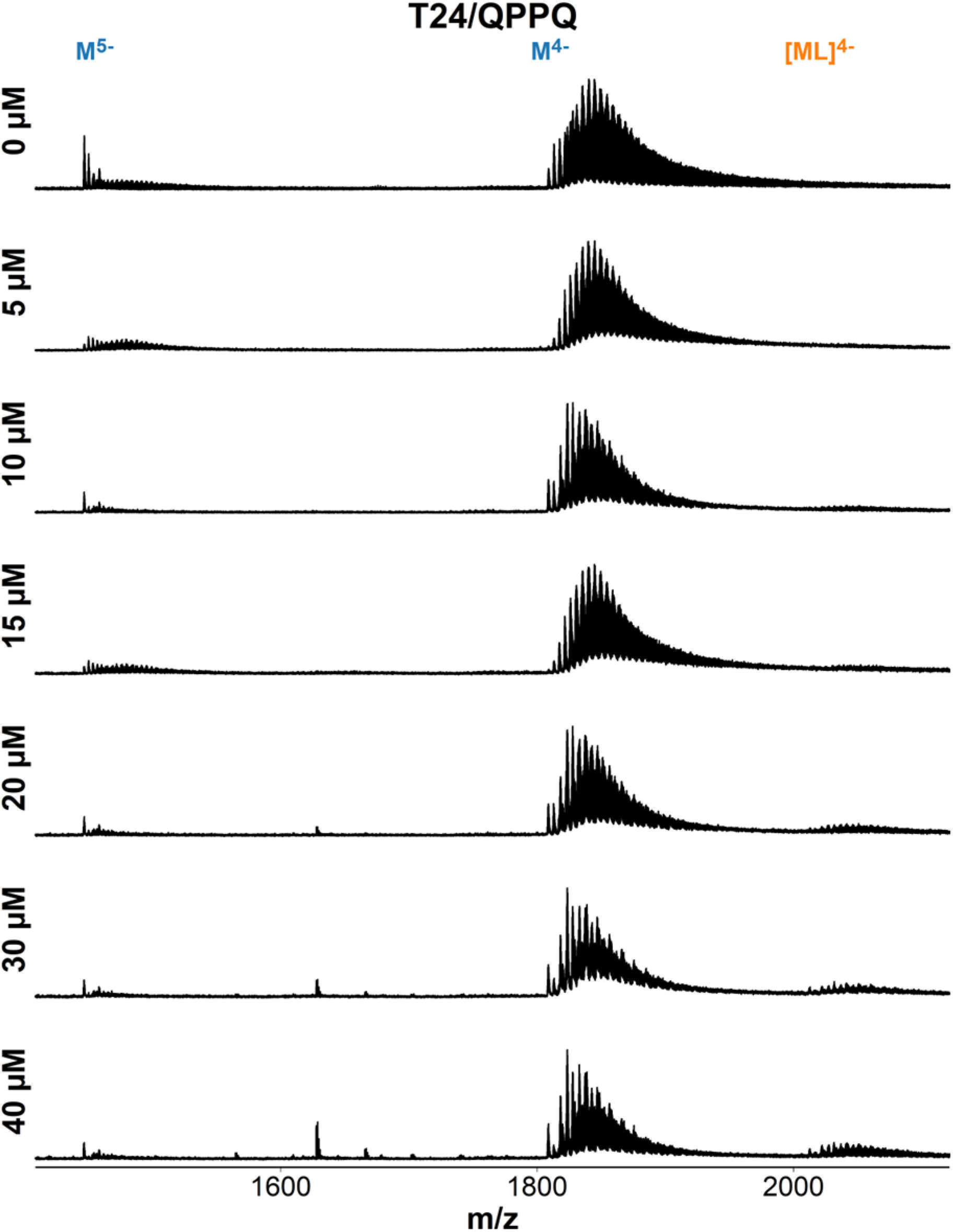
ESI-MS titration of T24 (dTTTTTTTTTTTTTTTTTTTTTTTT) with foldamer QPPQ. Samples contain 10 µM DNA, 0-40 µM ligand, 0.5 mM KCl, 100 mM TMAA (pH 6.8).

**Figure S109.**
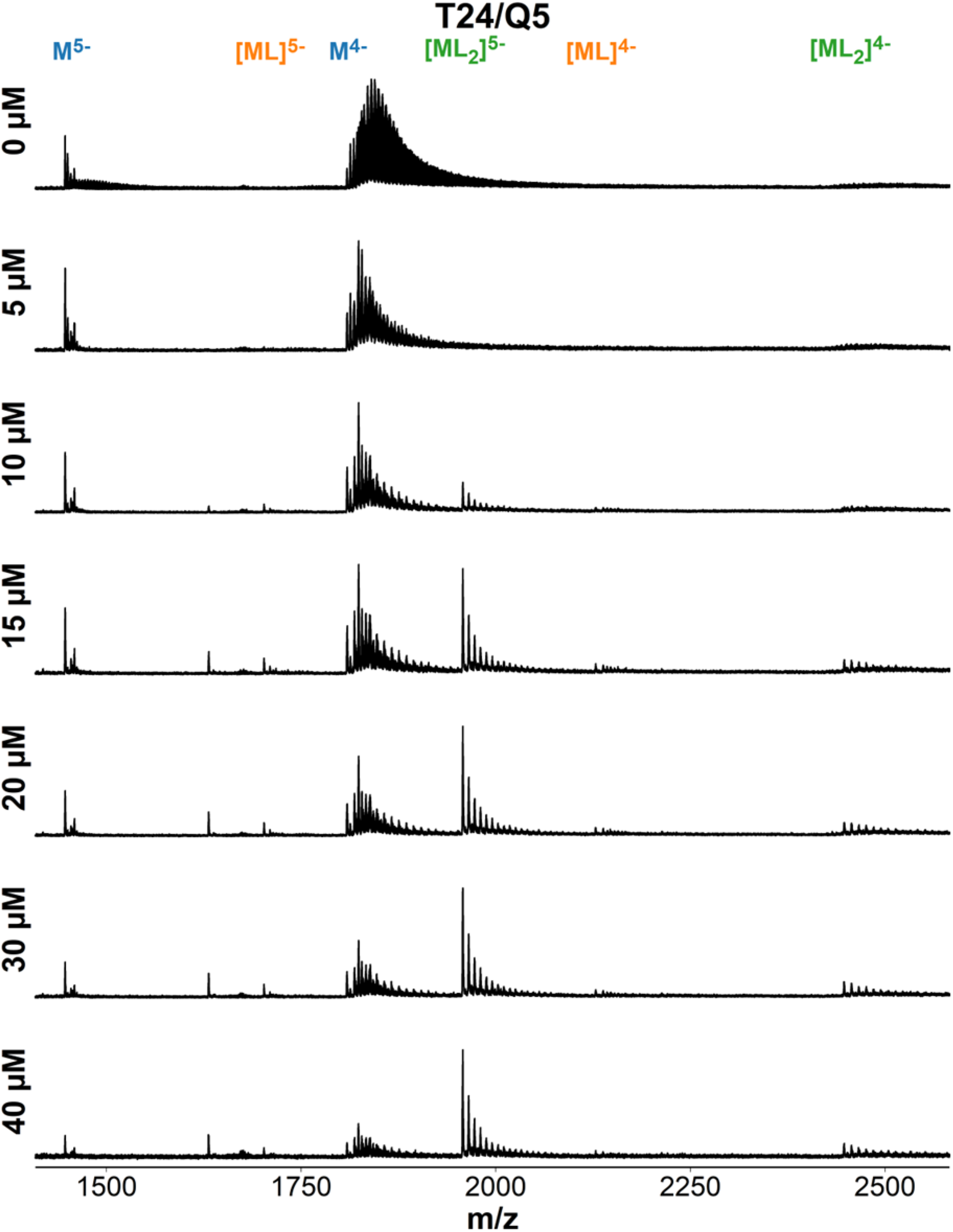
ESI-MS titration of T24 (dTTTTTTTTTTTTTTTTTTTTTTTT) with foldamer QQQQQ. Samples contain 10 µM DNA, 0-40 µM ligand, 0.5 mM KCl, 100 mM TMAA (pH 6.8).

**Figure S110.**
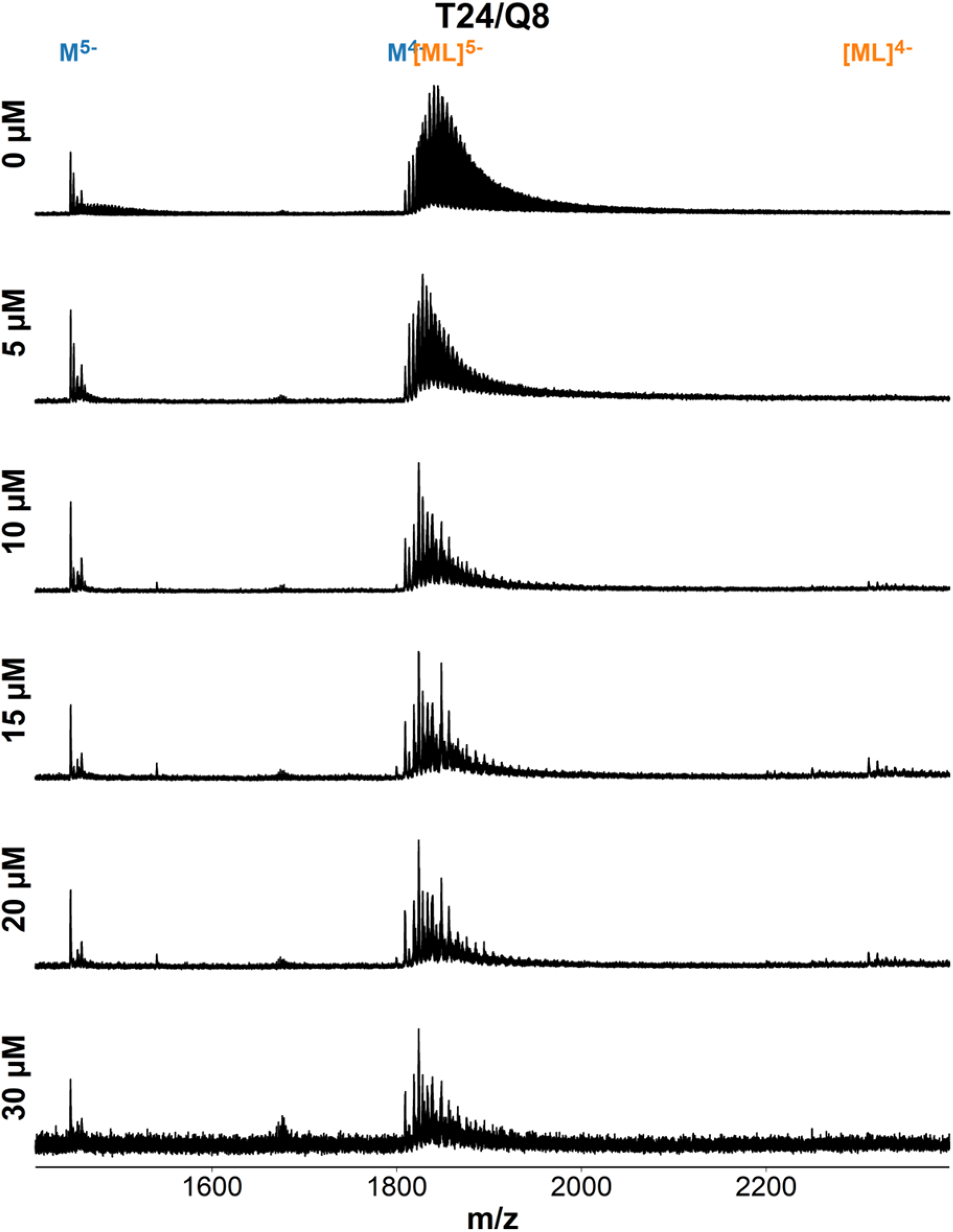
ESI-MS titration of T24 (dTTTTTTTTTTTTTTTTTTTTTTTT) with foldamer QQQQQQQQ. Samples contain 10 µM DNA, 0-30 µM ligand, 0.5 mM KCl, 100 mM TMAA (pH 6.8).

#### T24 induces CD on Q_n_-type foldamers

Among our ligand screening results, we noticed that Q-mers (i.e. Q3, Q4, Q5, Q8) form high-affinity 2:1 complexes (low µM KD) with single-stranded sequence T24. We investigated this phenomenon via CD (Figure S111).

**Figure S111.**
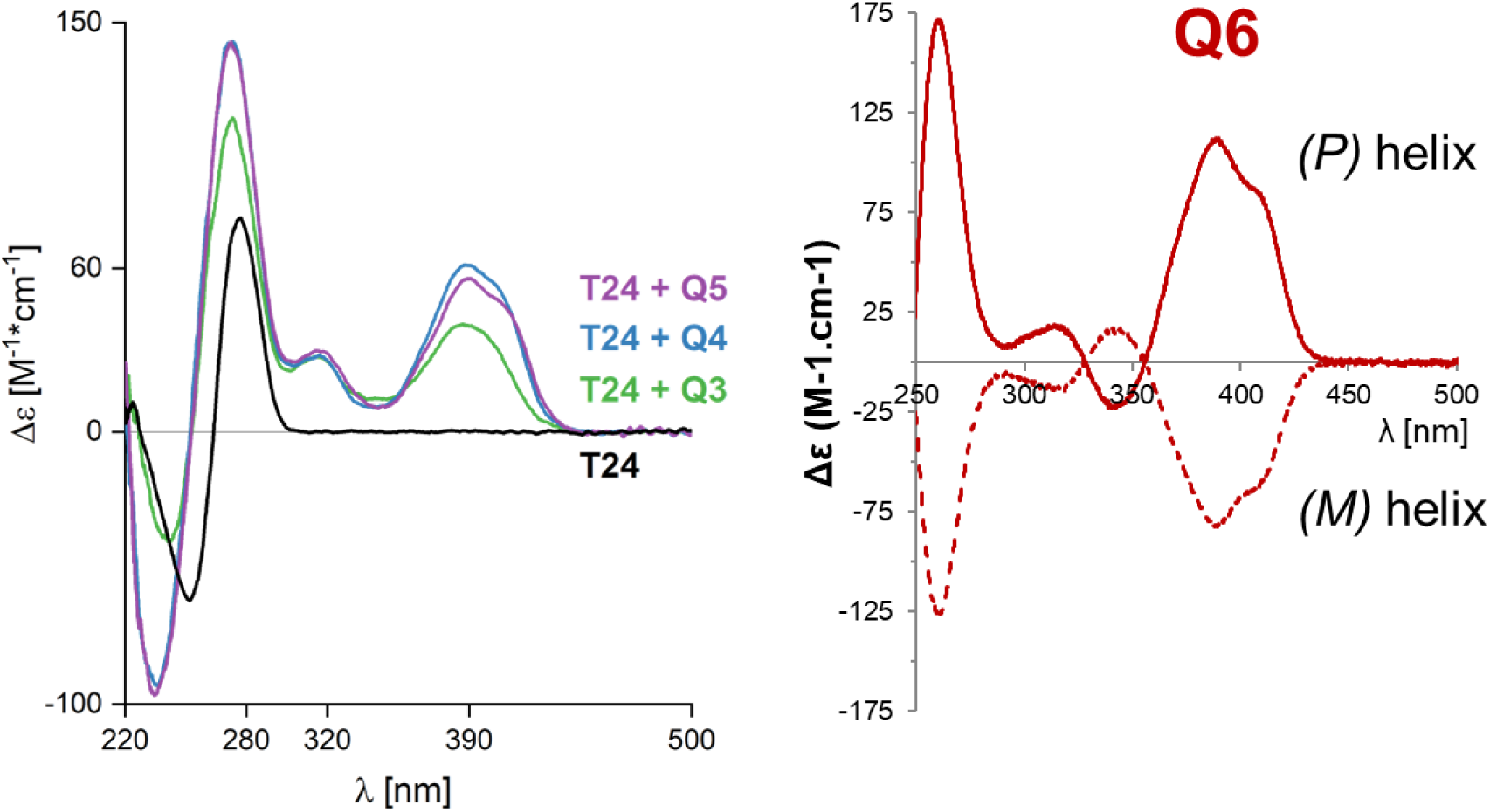
**Left**: CD signatures of dT_24_ with different Q-mer foldamers. Samples contain: 10 µM T24, 40 µM Q_3_/30 µM Q_4_/24 µM Q_5_ (normalized to 120 µM Q monomer) and 100 mM TMAA (pH 6.8). **Right**: Enantiomer-separated CD signatures of the (M) and (P) helix for Q6.

The Q3/Q4/Q5 we used in our studies has no CD signature on its own, since they are racemic mixtures. But with T24, we see CD bands that can be interpreted as enantiomeric excess of the *(P)*-helix. We do not know that kind of structure T24 and Q-mer foldamers form; our attempt of crystallizing a T24/Q4 complex was unsuccessful. Based on the high 2:1 cooperativity and the induced CD on the (right-handed) *(P)*-helix we speculate that T24 and Qn foldamers associate into some sort of double helix, with the foldamer dimerizing to accommodate the full length of the DNA sequence.

U-rich motifs play a role in RNA expression,^6,7^ which is why we attempted to reproduce our results on U24 (Figure S112). Nonetheless, there was to remarkable induced CD in U24 upon adding Qn foldamer.

**Figure S112.**
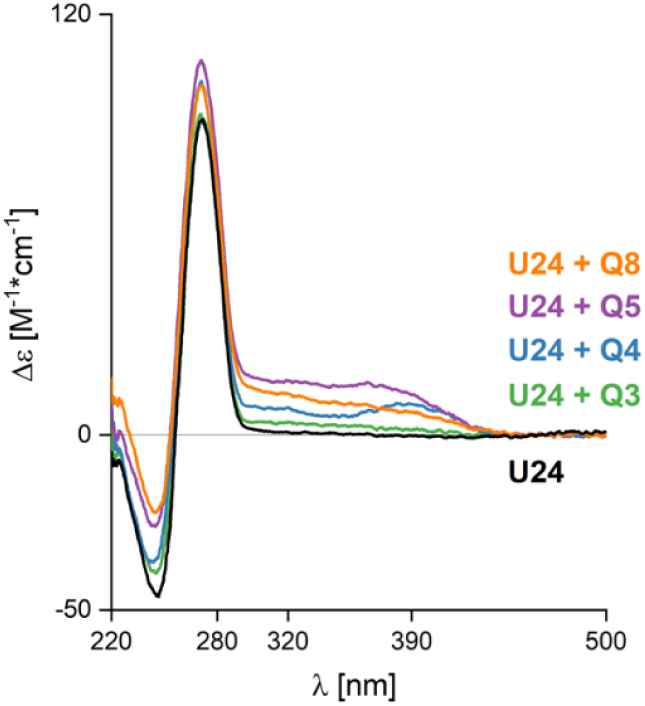
CD signatures of rU_24_ with different Q-mer foldamers. Samples contain: 10 µM U24, 40 µM Q_3_/30 µM Q_4_/24 µM Q_5_/15 µM Q_8_ (normalized to 120 µM Q monomer) and 100 mM TMAA (pH 6.8).

#### Foldamer-induced disruption of G-quadruplex

Figure S32 shows that the decline in DNA signal response is proportional to the number of Q units in the foldamer. Each Q unit carries a positive charge, so we suspect that the loss of signal is due to ligand-induced DNA aggregation, with the ligand acting as a sort of flocculant that helps overcome charge repulsions between DNA polyanions. To test this hypothesis, we picked T30177TT, a parallel G-quadruplex that is stable even in low K^+^ and Q8, the ligand with the fastest decrease in response factor. If the ligand causes DNA aggregation, the CD signature of T30177TT should decrease the more ligand is added.

**Figure S113.**
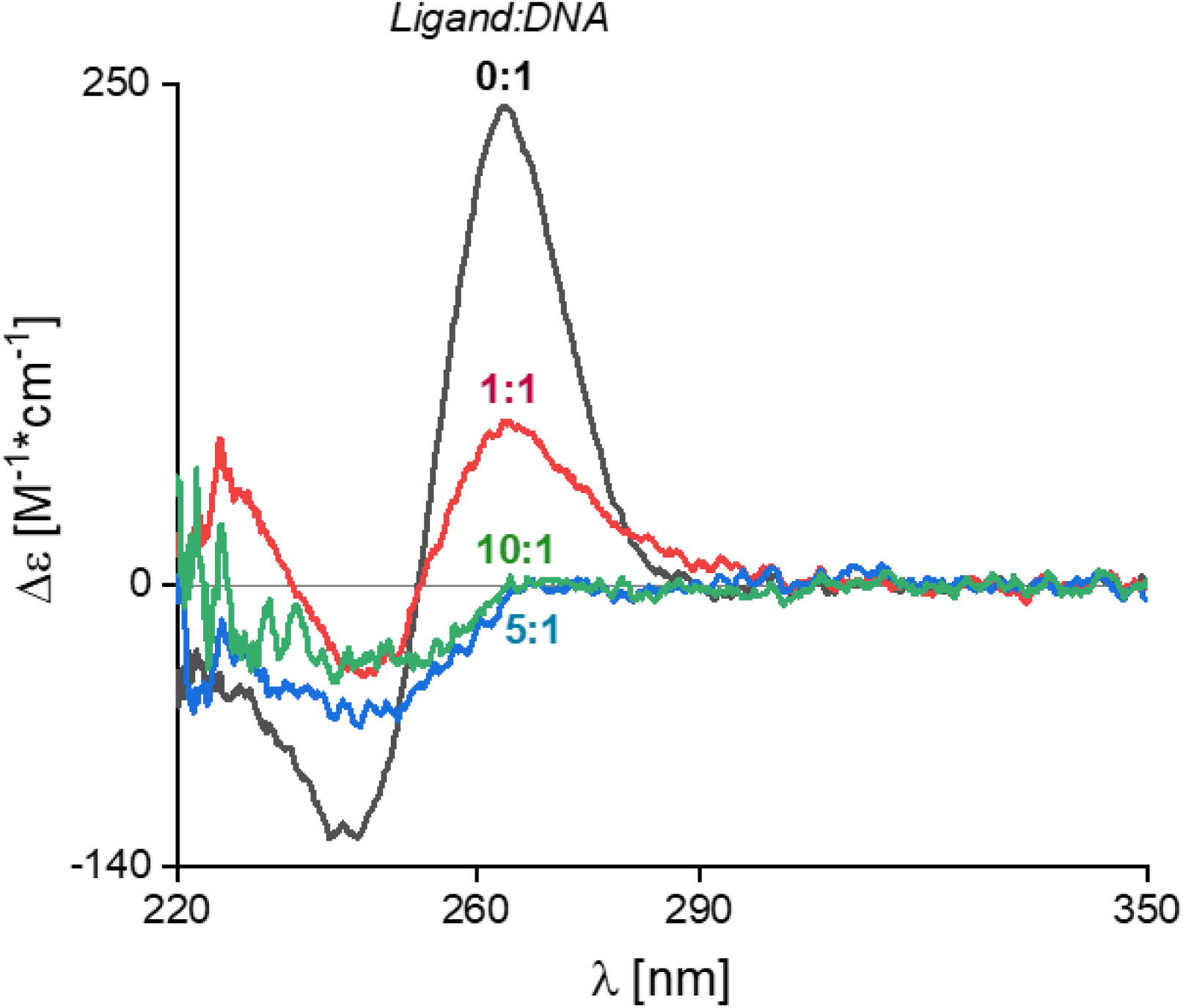
Tracking the ligand-induced aggregation of T30177TT (dTT**G**T**GG**T**GGG**T**GGG**T**GGG**T) by adding more and more Q8. Samples contain 1 µM DNA, 0/1/5/10 µM Q_8_, 1 mM KCl and 100 mM TMAA (pH 6.8).

The experimental results are consistent with our hypothesis, indicating that G-quadruplexes with too many positive charges run the risk of causing DNA aggregation, or even precipitation (aka ‘salting out’ a biomolecule by neutralizing all of its charges).

#### X-ray crystallography

**Table S5.** Data acquisition and refinement parameters for the 222T/QQPQ crystal. Parenthesis show statistics for the highest-resolution shell.

|  |  |
| --- | --- |
| Space group | P42 |
| Unit cell (Å) | a = b = 32.71, c = 61.01, $\alpha = \beta = \gamma = 90.000^\circ$ |
| Wavelength (Å) | 0.979 |
| Resolution range (Å) | 30.5-2.512 (2.602-2.512) |
| Completeness (%) | 99.10 (100.0) |
| Total reflections | 29748 (2985) |
| Unique reflections | 2226 (218) |
| Multiplicity | 13.4 (13.7) |
| Average I/$\sigma$ | 21.61 (4.09) |
| Wilson B-Factor | 58.79 |
| R-merge | 0.1469 (1.054) |
| R-means | 0.1531 (1.095) |
| R-pim | 0.0426 (0.2952) |
| CC $\frac{1}{2}$ | 0.999 (0.872) |
| CC* | 1 (0.965) |
| Reflections used in refinement | 2207 (218) |
| Reflections used for R-free | 221 (22) |
| R-work | 0.2563 (0.2933) |
| R-free | 0.3037 (0.4455) |
| CC (work) | 0.902 (0.773) |
| CC (free) | 0.990 (0.336) |
| r.m.s. bond deviation (Å) | 0.018 |
| r.m.s. angle deviation ($^\circ$) | 1.84 |
| Clashscore | 23.19 |
| Number of non-hydrogen atoms | 531 |
| DNA atoms | 388 |
| QQPQ atoms | 243 |
| Other atoms (K<sup>+</sup> and Mg<sup>2+</sup>) | 2 |
| Average B factors | 66.24 (DNA: 62, QQPQ: 79) |
| DNA atoms | 61.87 |
| QQPQ atoms | 78.21 |
| Other atoms (K<sup>+</sup> and Mg<sup>2+</sup>) | 69.63 |

**Figure S114.**
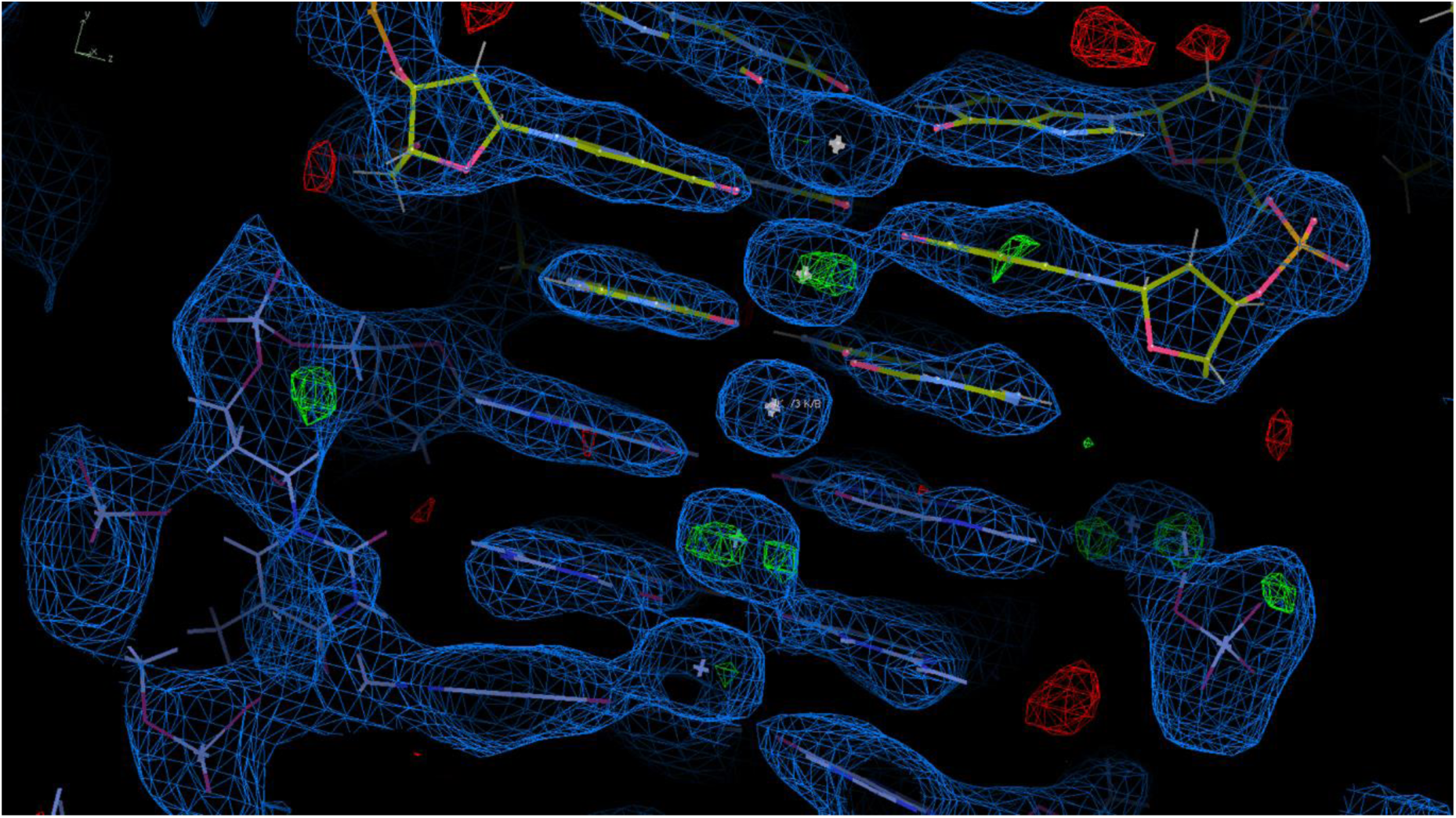
Electron density map of the G-quadruplex core structure

**Figure S115.**
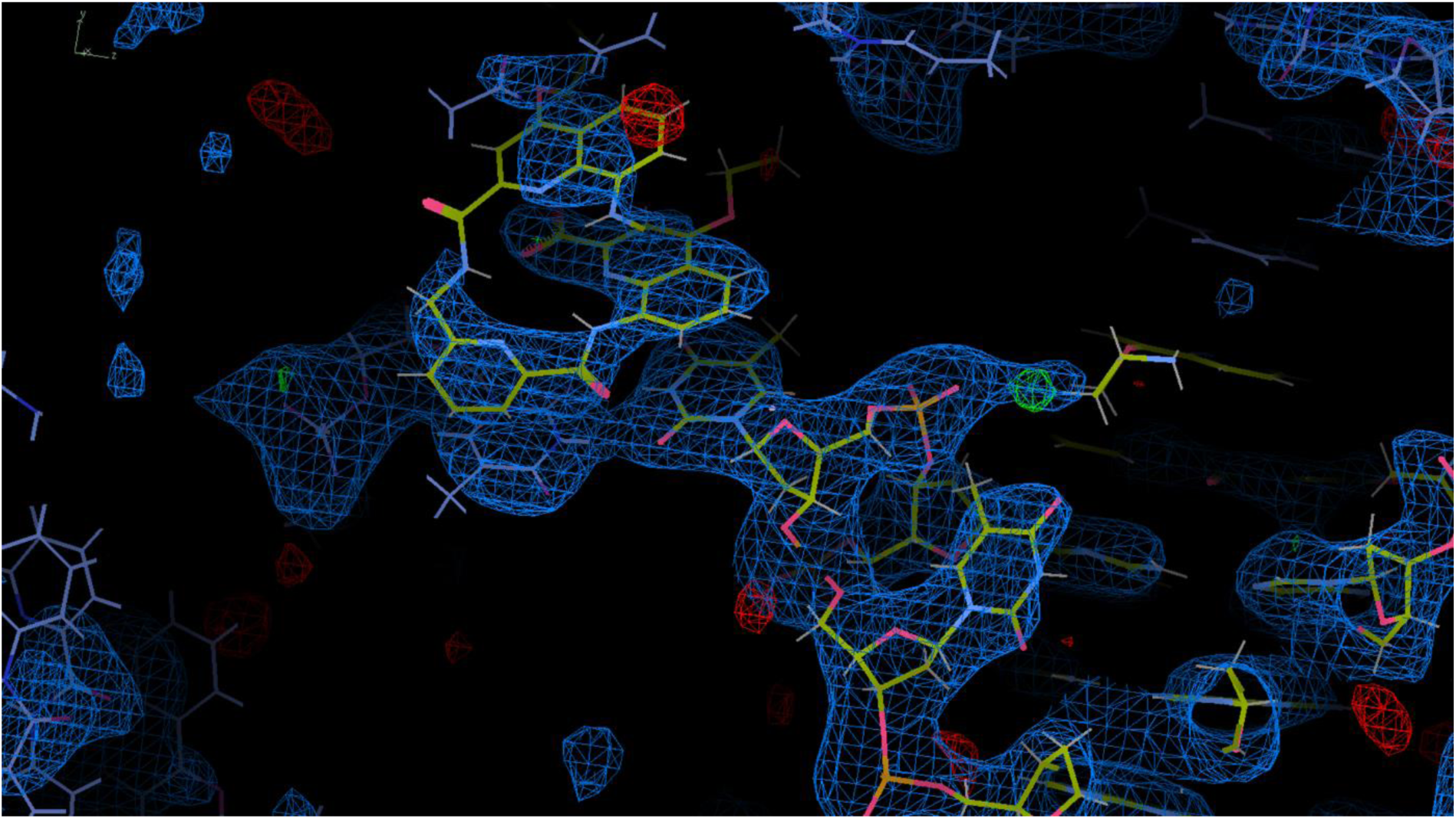
Electron density map of the QQPQ foldamer stacked on top of a thymine in the 222T loop region.

**Figure S116.**
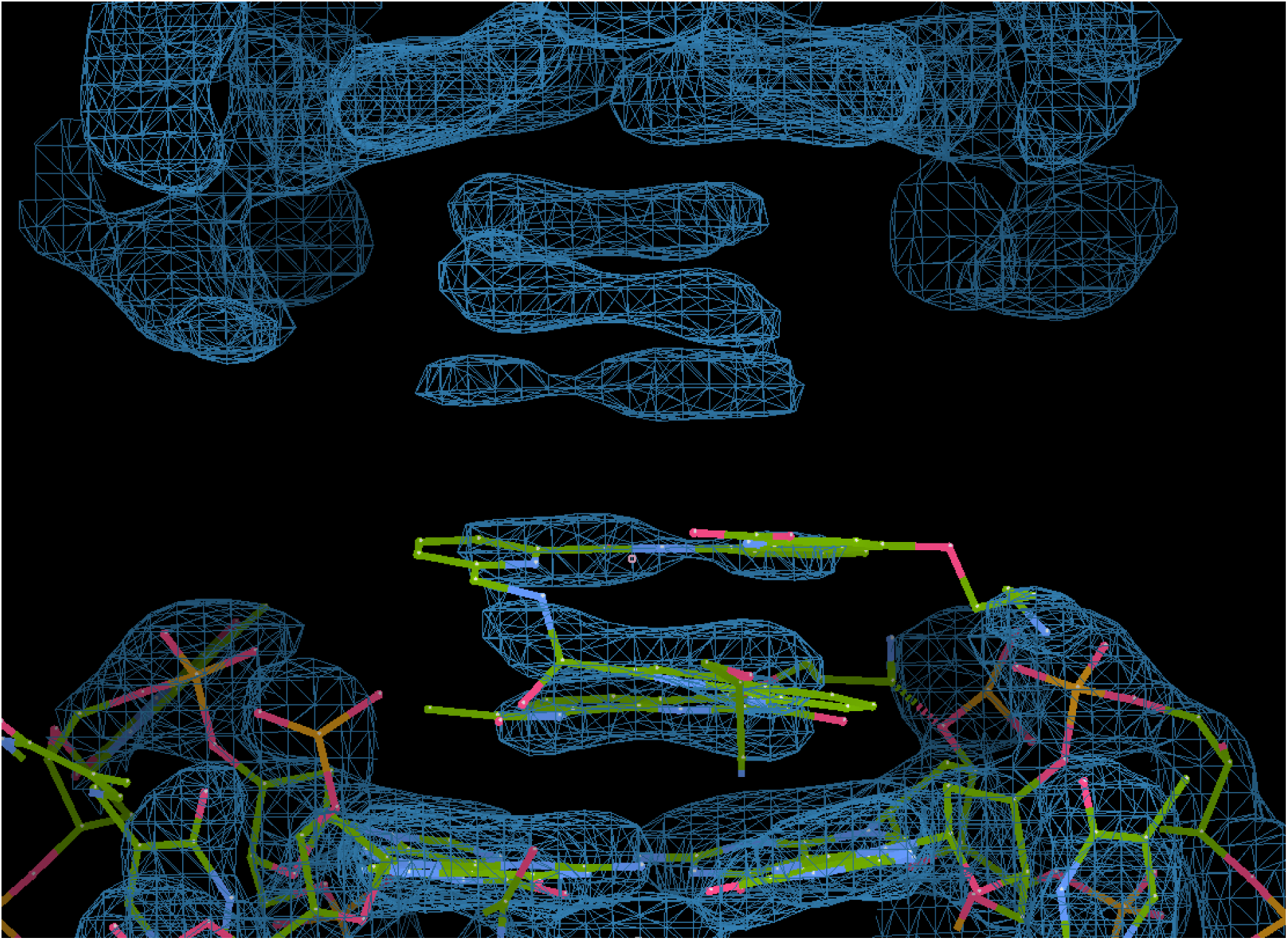
Electron density map of the QQPQ foldamer stacked onto the G-tetrad of 222T, showing two adjacent unit cells on top and on the bottom.

**Figure S117.**
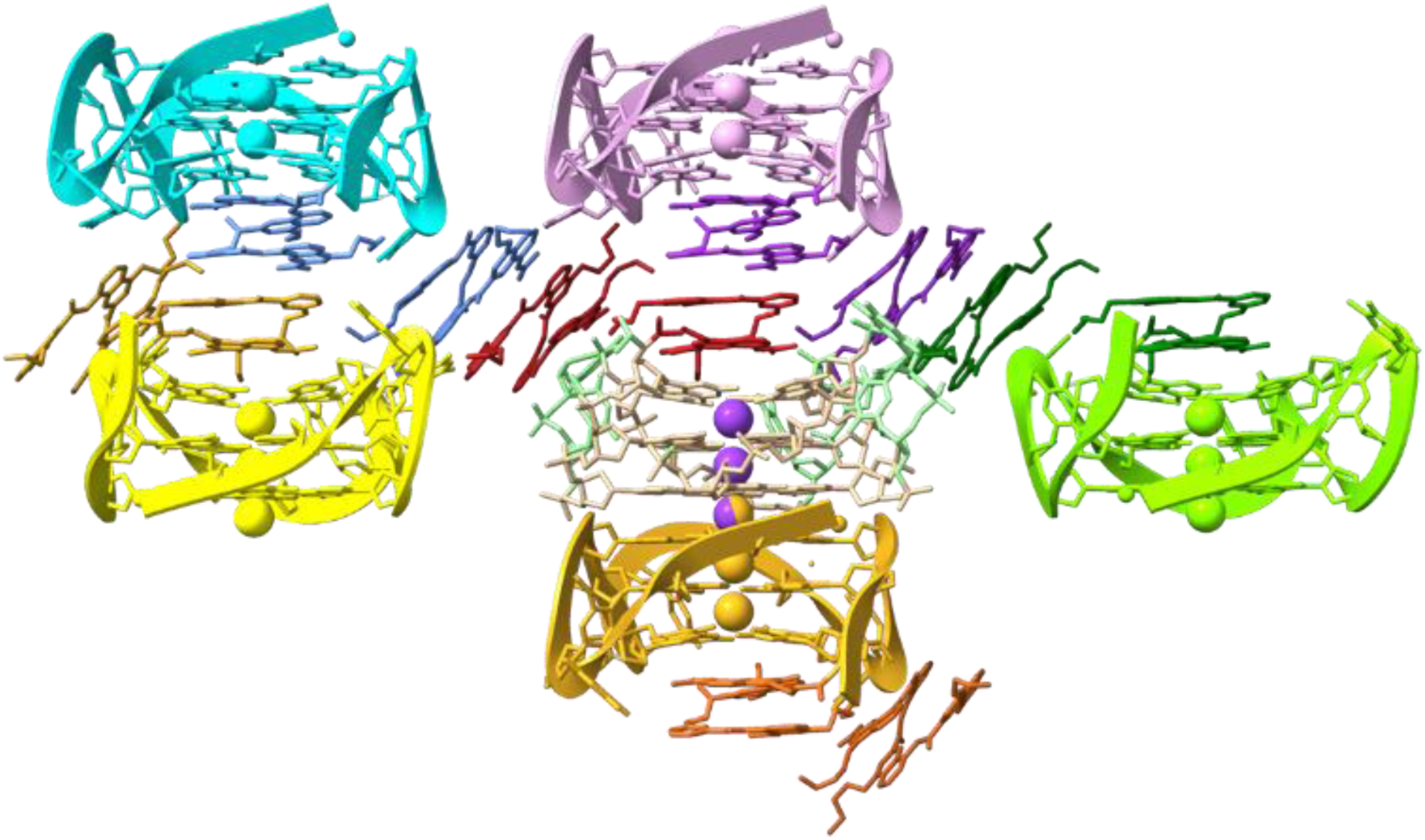
Crystal packing of the 222T/QQPQ crystal. Different colors show different unit cells.

**Figure S118.**
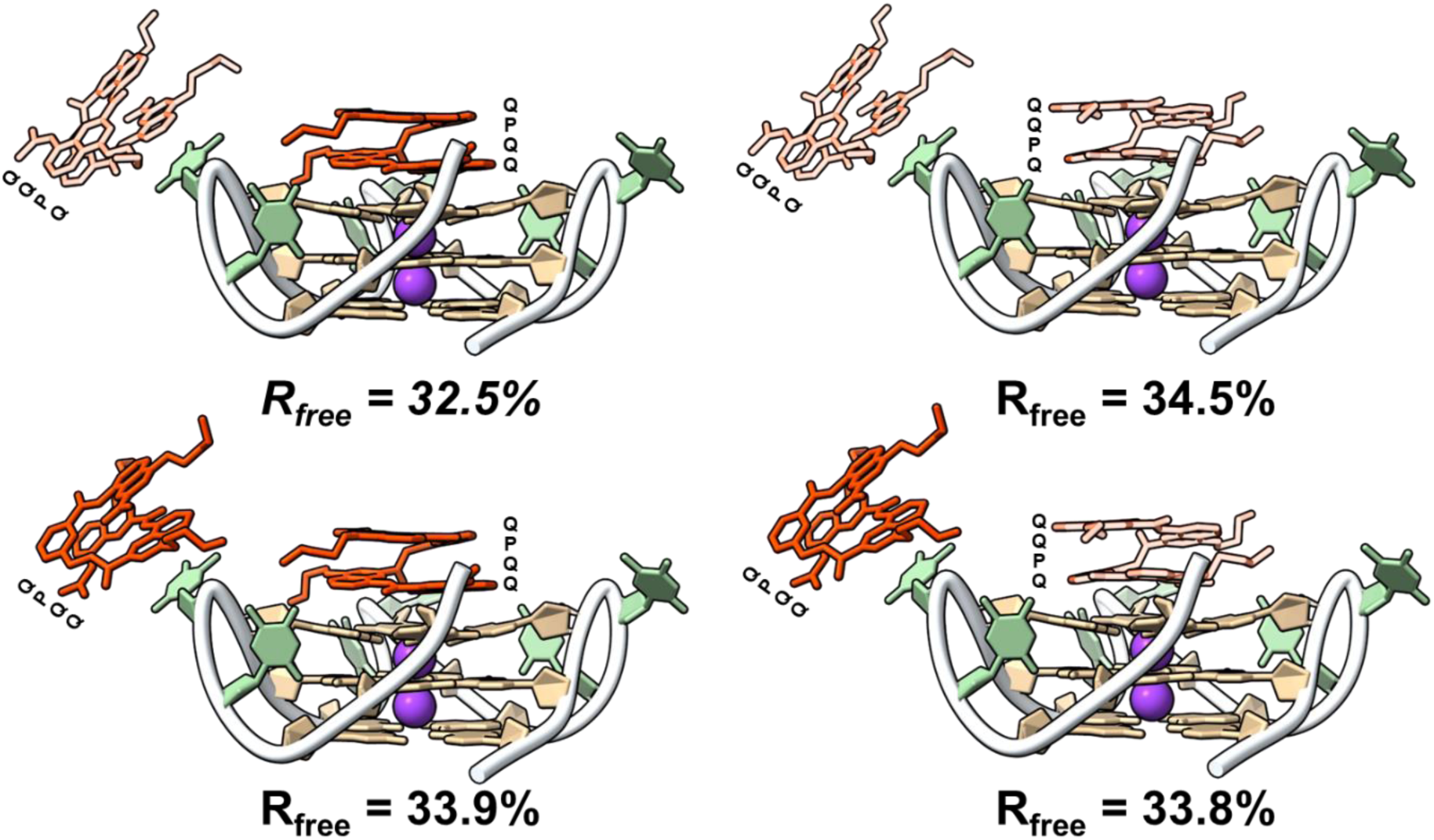
Flipping the foldamer between C- and N-terminal binding to test whether any conformation is statistically preferential. Schemes represent the four different models and their resulting R_free_ values.

#### 1D and 2D NMR spectra

**Figure S119.**
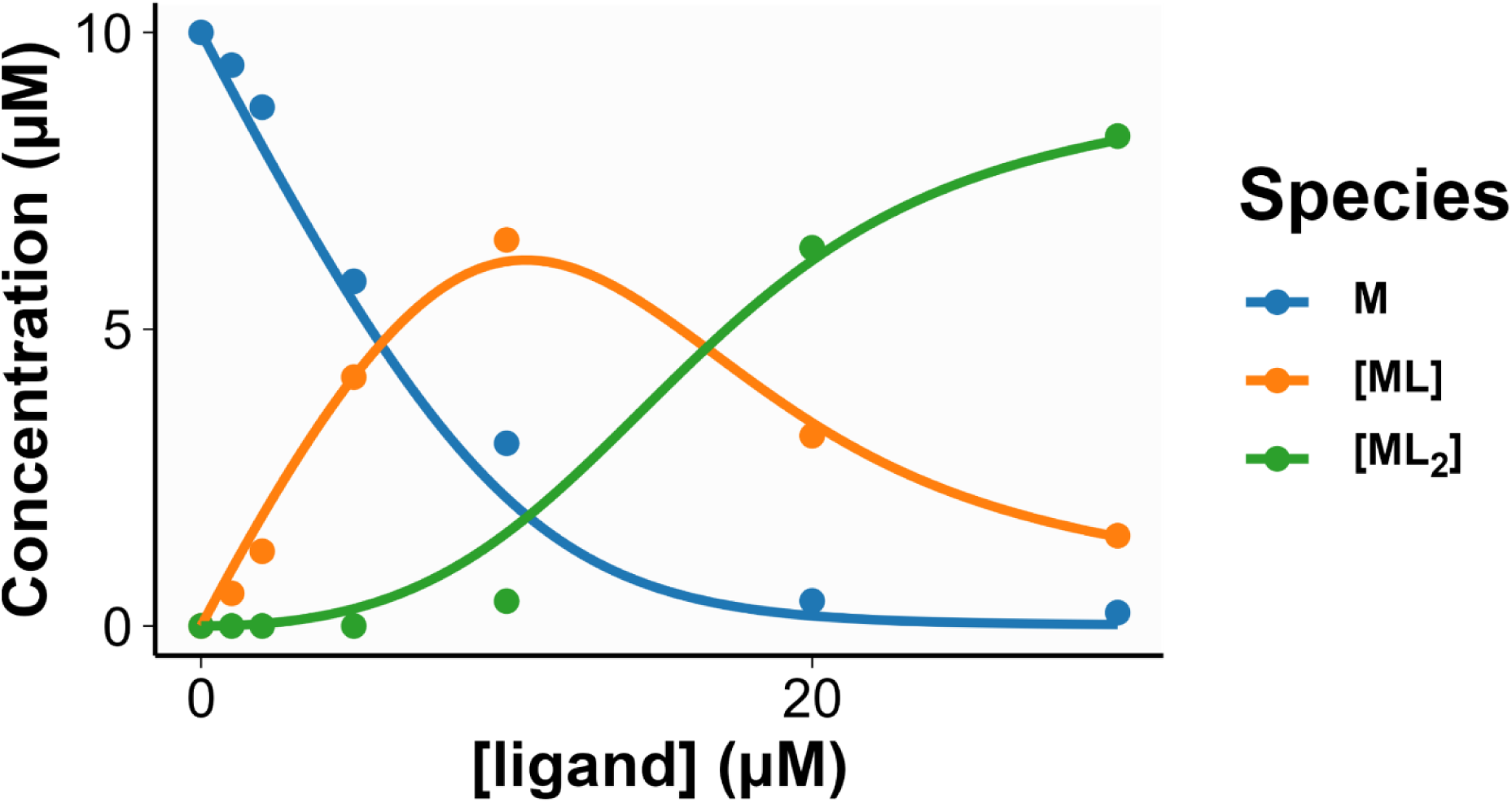
Dynamic fitting of MS titration data for 2LK7 (dTT**G_3_**T**G_3_**T**G_3_**T**G_3_**T) and foldamer QQPQ with K_D1_ = 0.18 ± 0.08 µM and K_D2_ = 2.1 ± 0.7 µM. Samples contain 10 µM DNA, 0/1/2/5/10/20/30 µM QQPQ, 0.5 mM KCl and 100 mM TMAA (pH 6.8).

**Figure S120.**
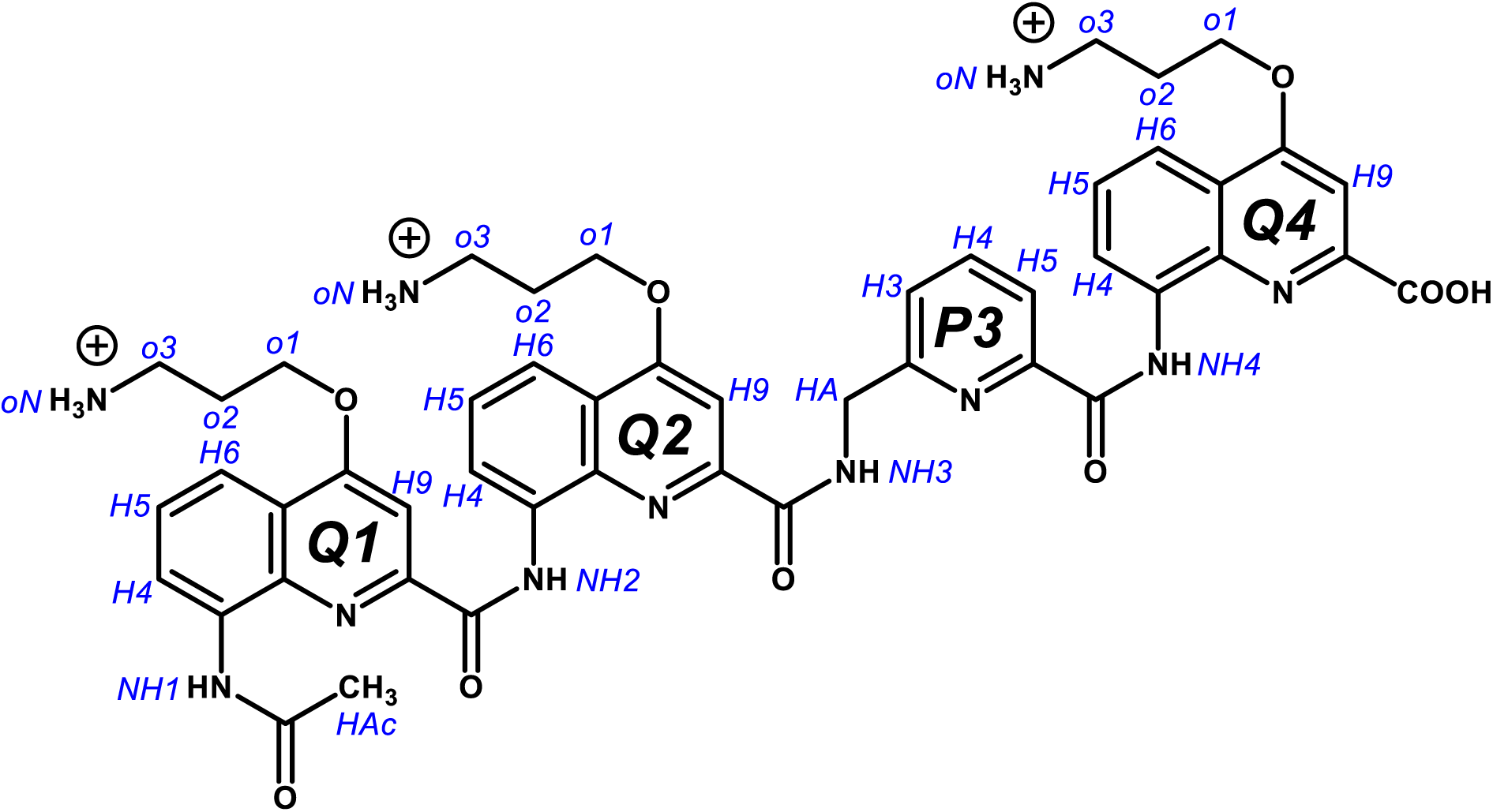
Proton labels in the QQPQ molecule.

**Figure S121.**
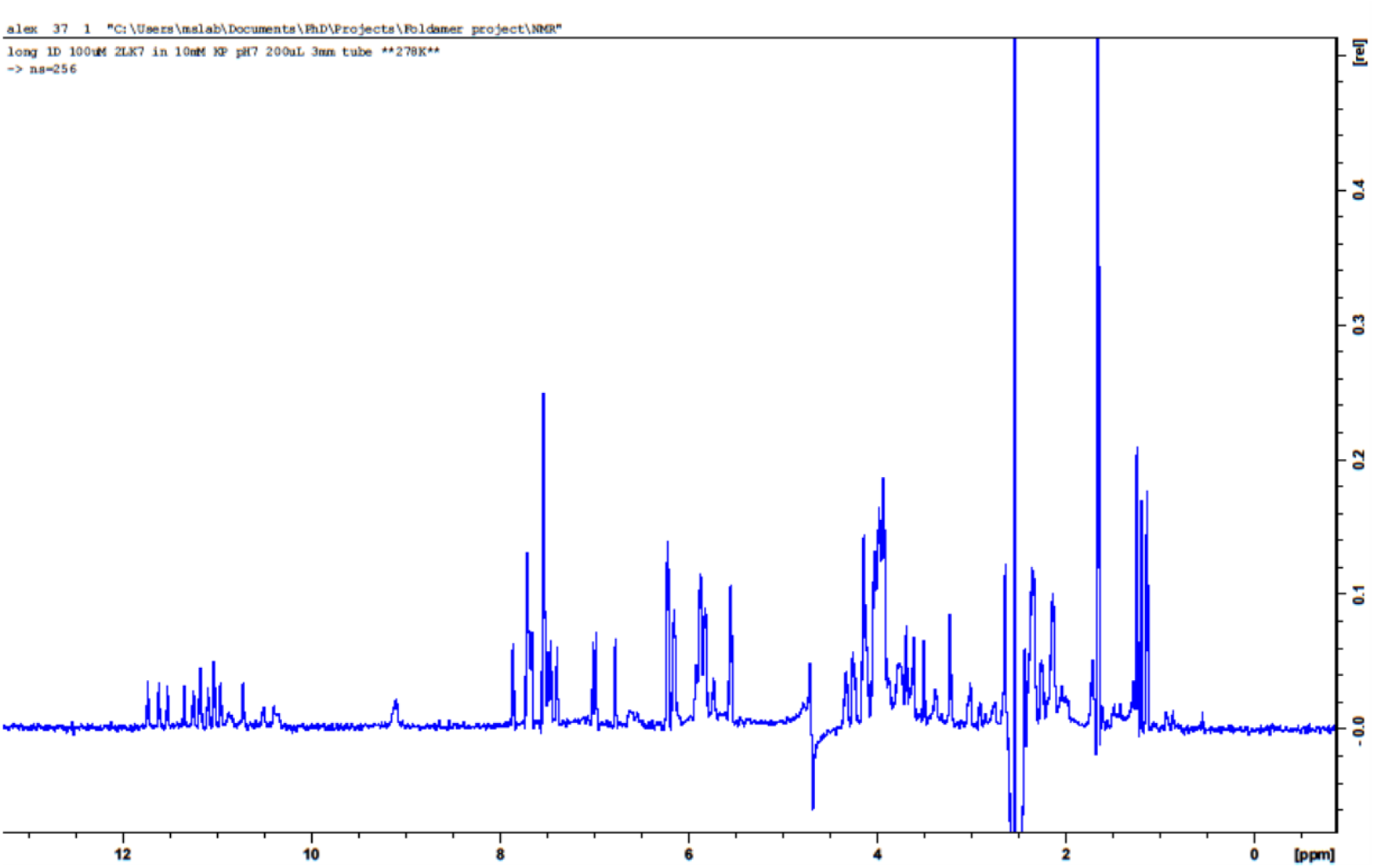
Full 1D NMR spectrum of 2LK7 (dTT**G_3_**T**G_3_**T**G_3_**T**G_3_**T) on a Bruker Avance 700 MHz at 278 K. Sample matrix: 100 µM DNA, 10 mM potassium phosphate buffer (pH 7), 90/10 H_2_O/D_2_O. Water signal was suppressed by excitation sculpting.

**Figure S122.**
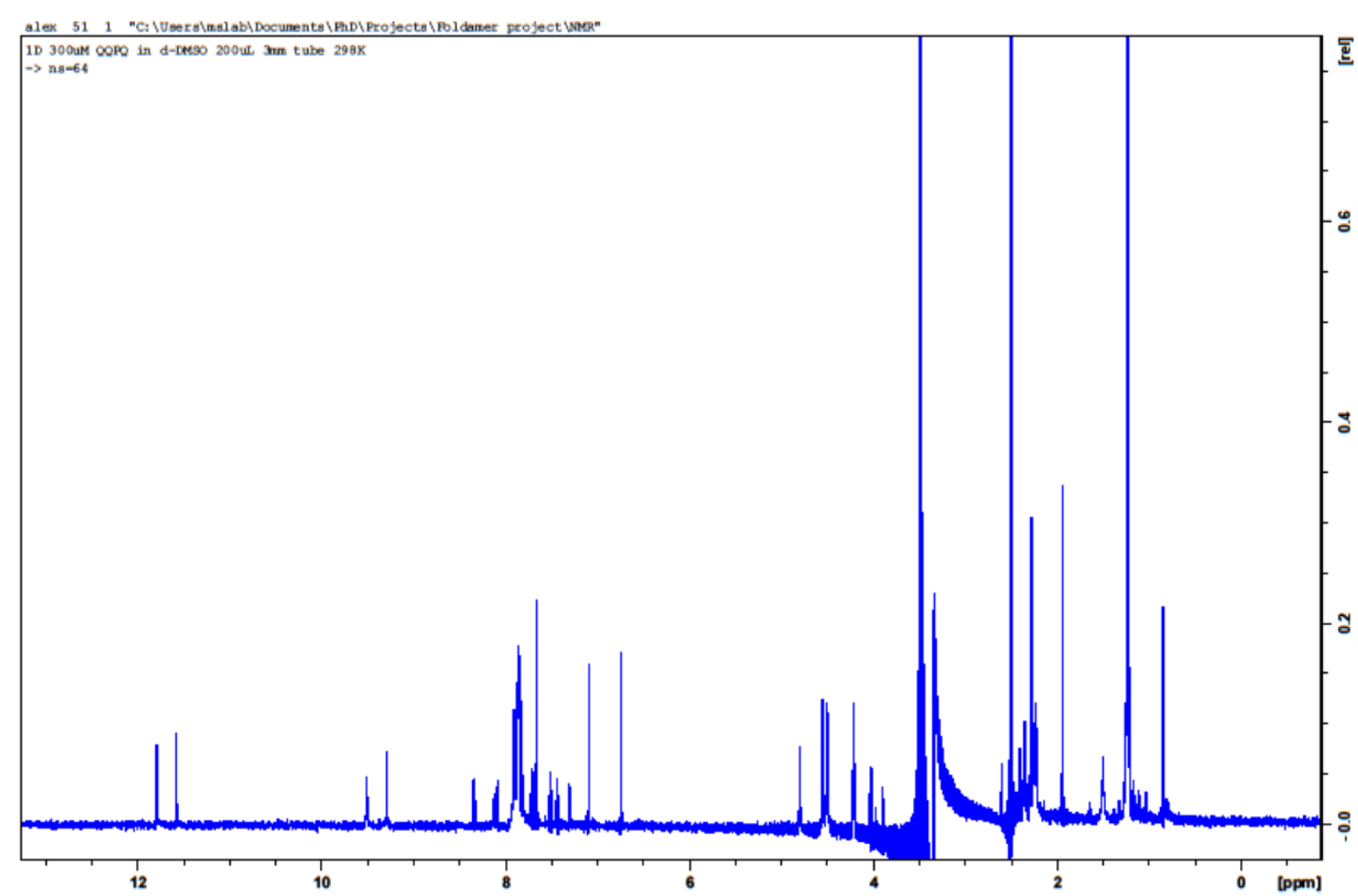
Full 1D NMR spectrum of Foldamer QQPQ on a Bruker Avance 700 MHz at 298 K. Sample matrix: 300 µM ligand in d6-DMSO.

**Figure S123.**
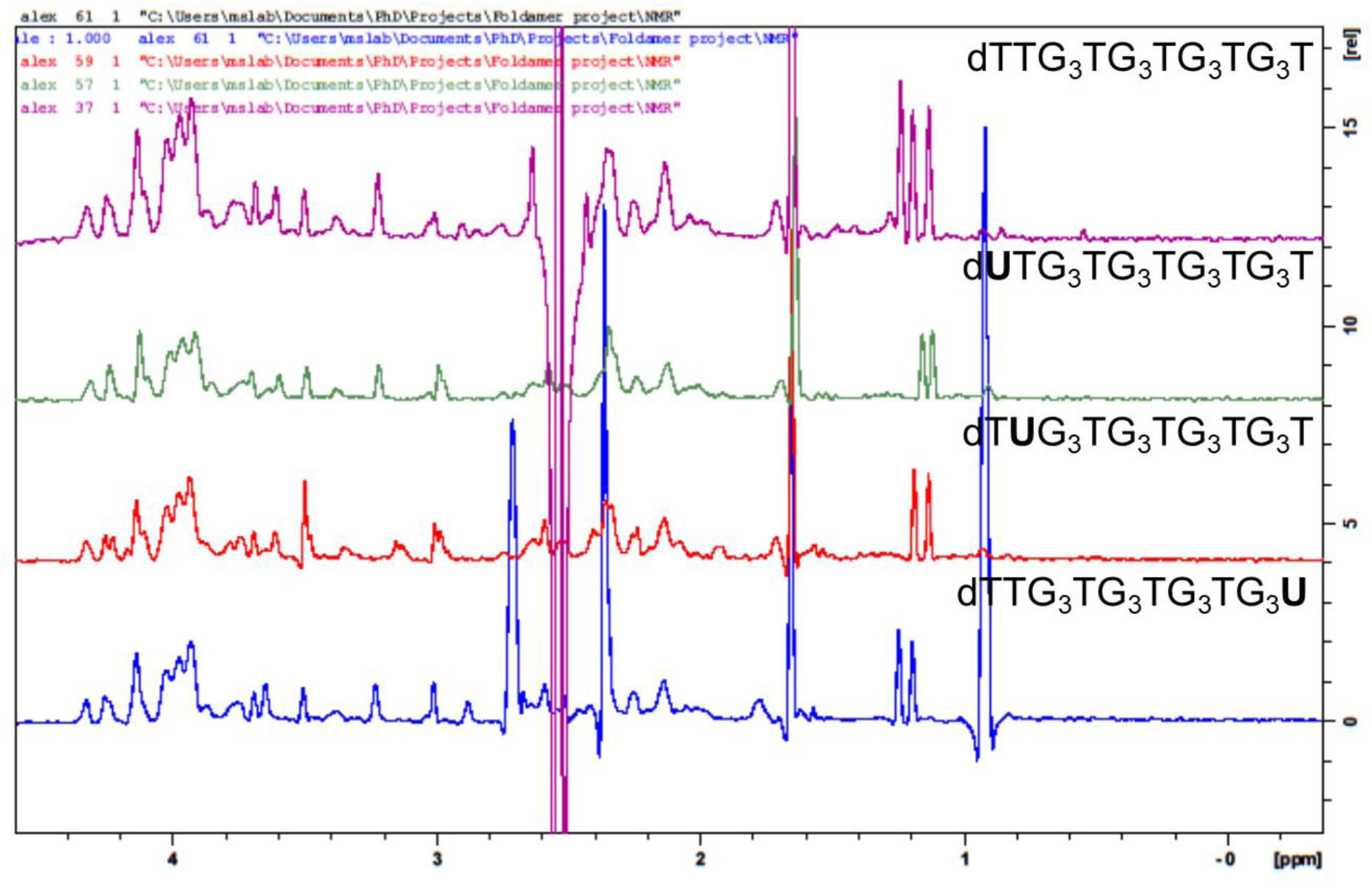
1D NMR spectra of 2LK7 and labeled derivatives on a Bruker Avance 700 MHz at 278 K, showing the low shift region. Sample matrix: 100 µM DNA, 10 mM potassium phosphate buffer (pH 7), 90/10 H_2_O/D_2_O. Water signal was suppressed by excitation sculpting.

**Figure S124.**
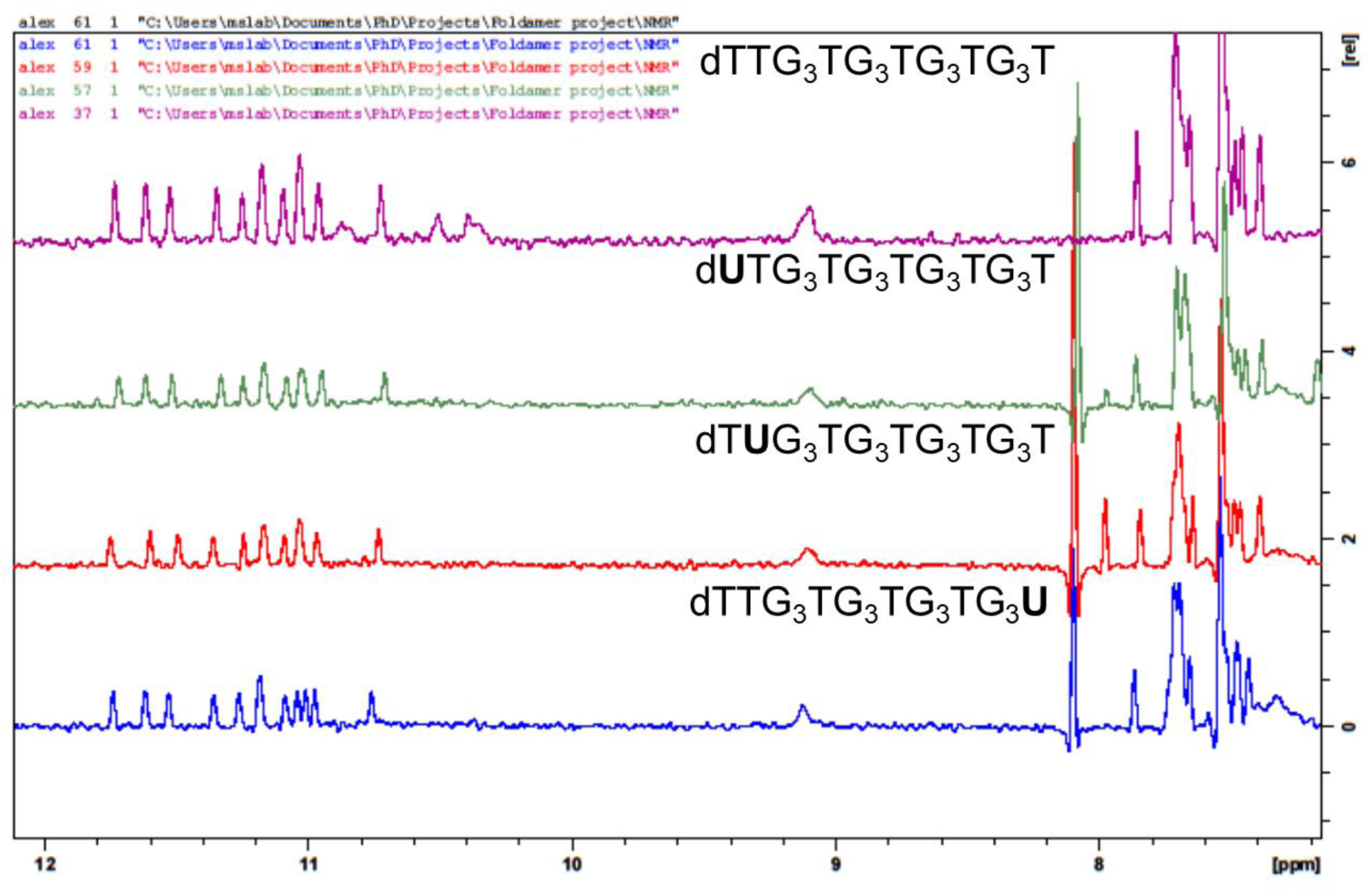
1D NMR spectra of 2LK7 and labeled derivatives on a Bruker Avance 700 MHz at 278 K, showing the high shift region. Sample matrix: 100 µM DNA, 10 mM potassium phosphate buffer (pH 7), 90/10 H_2_O/D_2_O. Water signal was suppressed by excitation sculpting.

**Figure S125.**
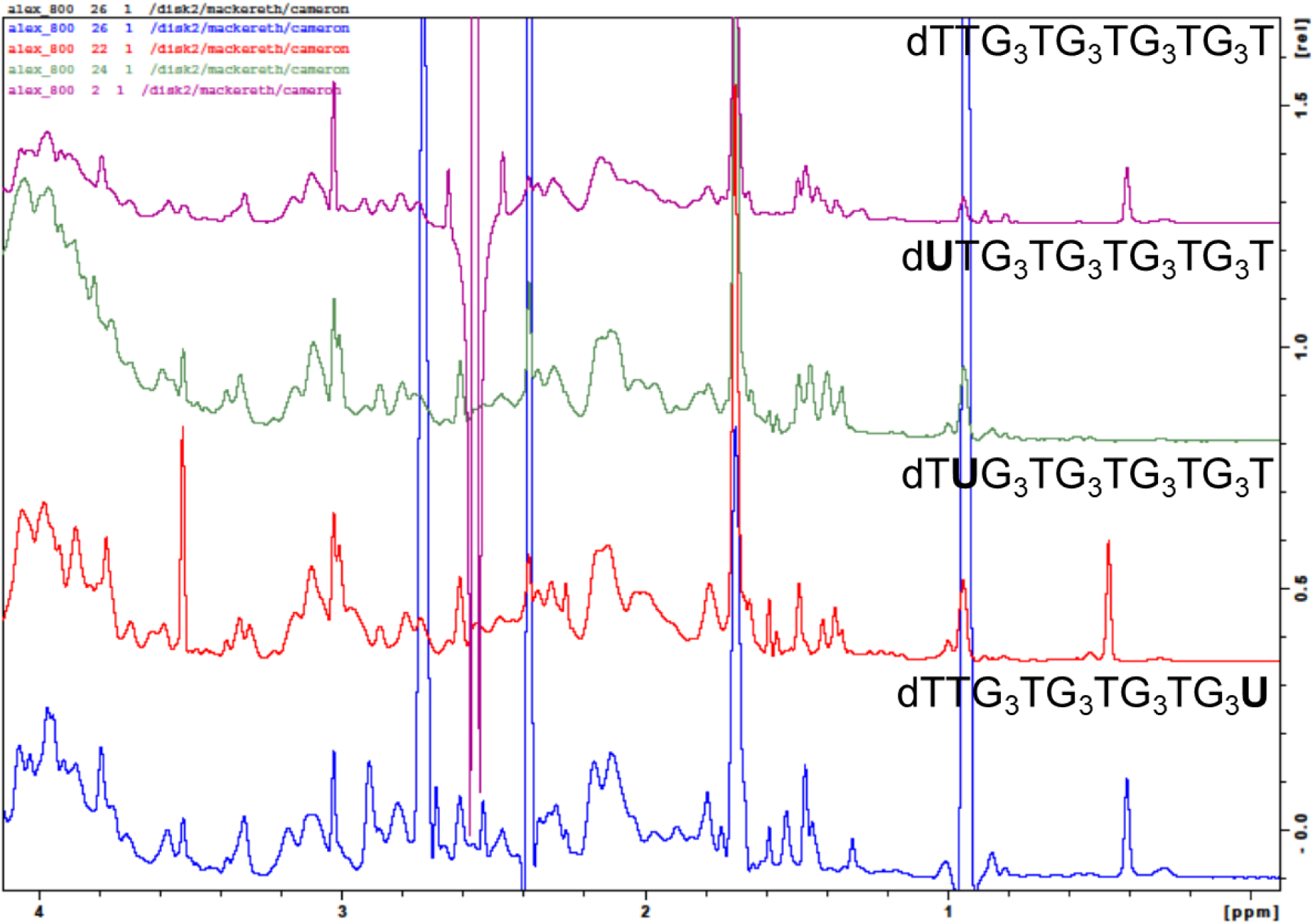
1D NMR spectra of 2LK7 and labeled derivatives with QQPQ on a Bruker Avance 700 MHz at 278 K, showing the low shift region. Sample matrix: 100 µM DNA,300 µM ligand. 10 mM potassium phosphate buffer (pH 7), 90/10 H_2_O/D_2_O.

**Figure S126.**
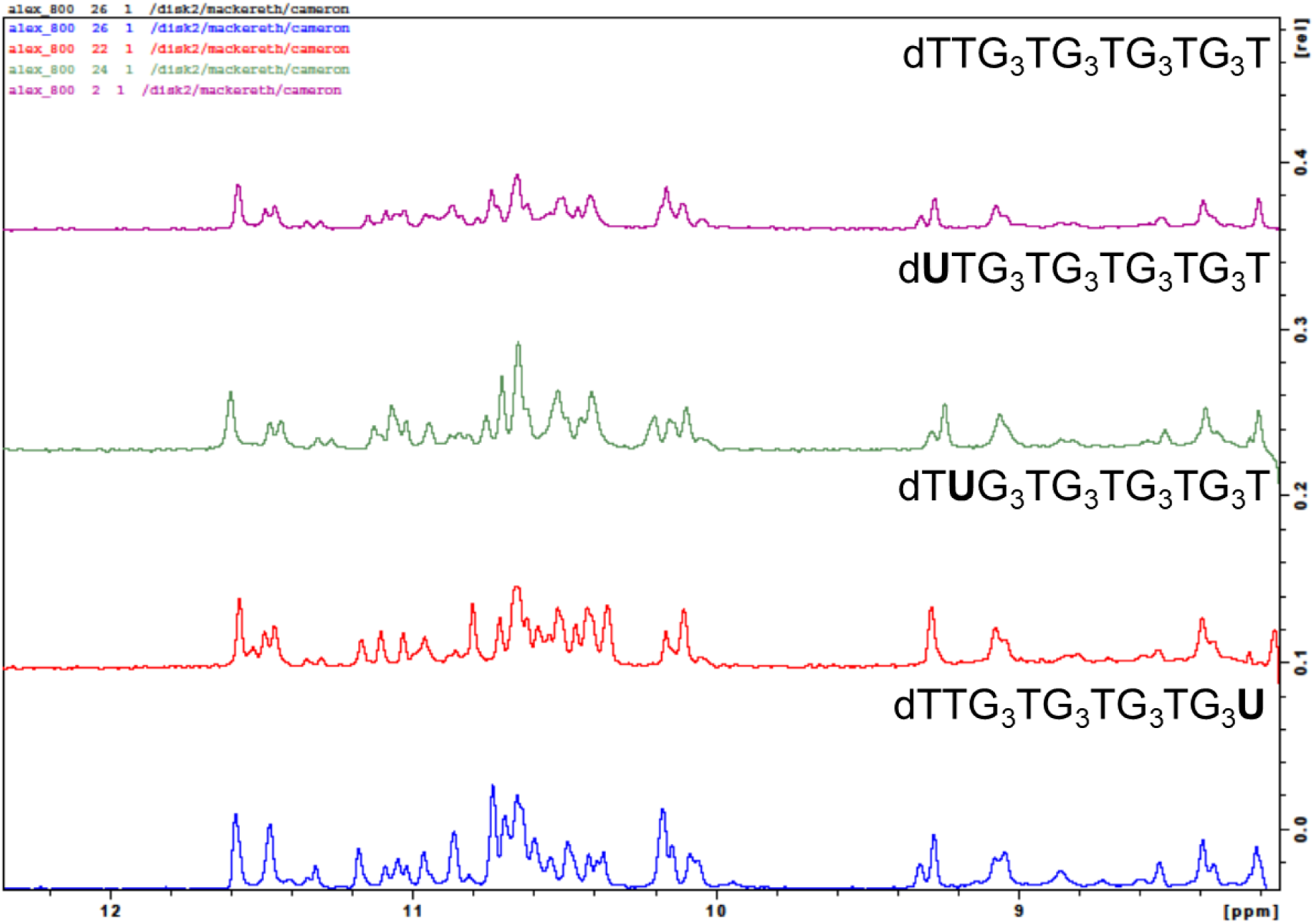
1D NMR spectra of 2LK7 and labeled derivatives with QQPQ on a Bruker Avance 700 MHz at 278 K, showing the high shift region. Sample matrix: 100 µM DNA,300 µM ligand. 10 mM potassium phosphate buffer (pH 7), 90/10 H_2_O/D_2_O.

**Figure S127.**
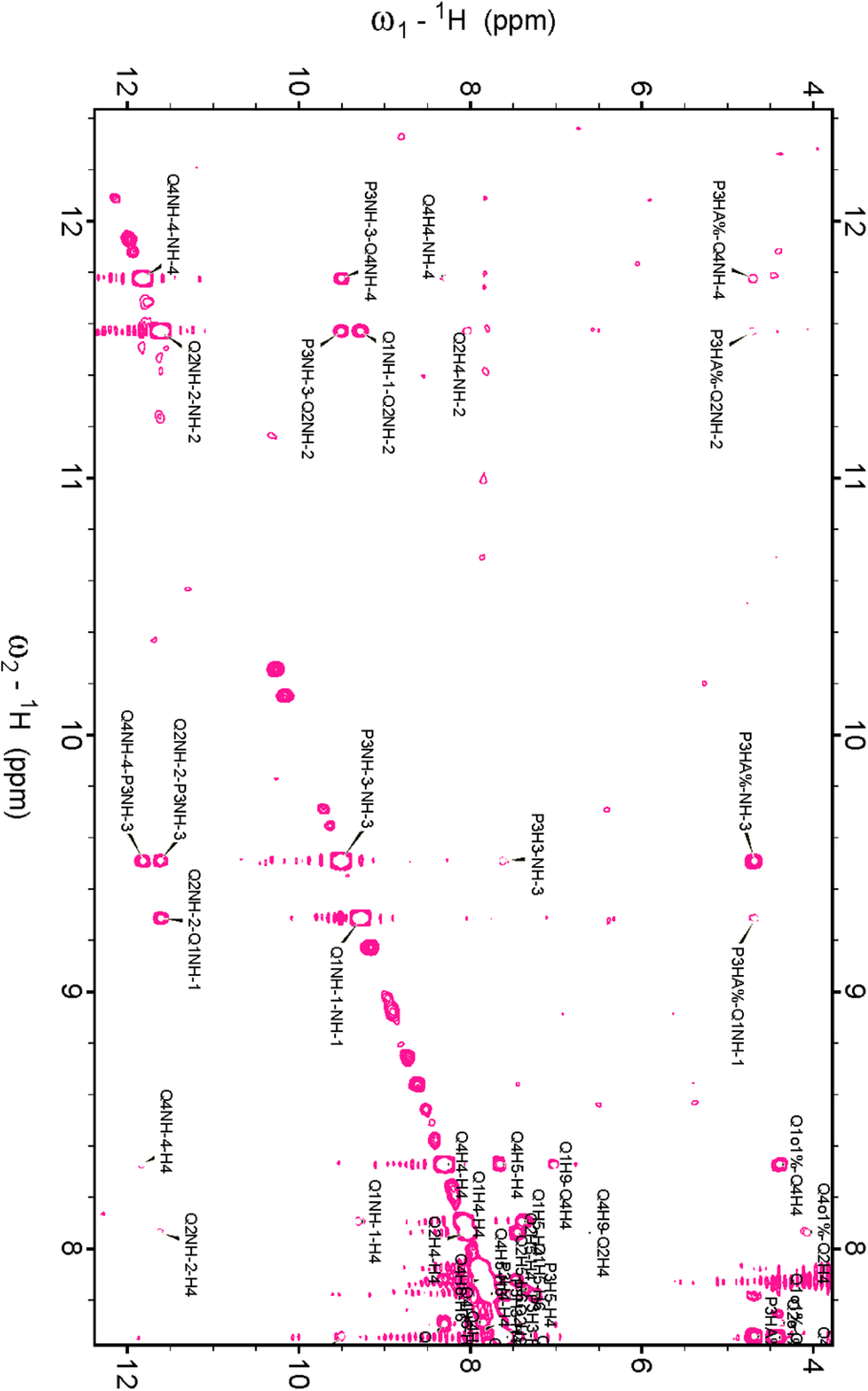
^1^H-^1^H NOESY of QQPQ on a Bruker Avance-III 800 MHz; high shift region containing amide protons (NH). Sample matrix: 300 µM QQPQ in d6-DMSO.

**Figure S128.**
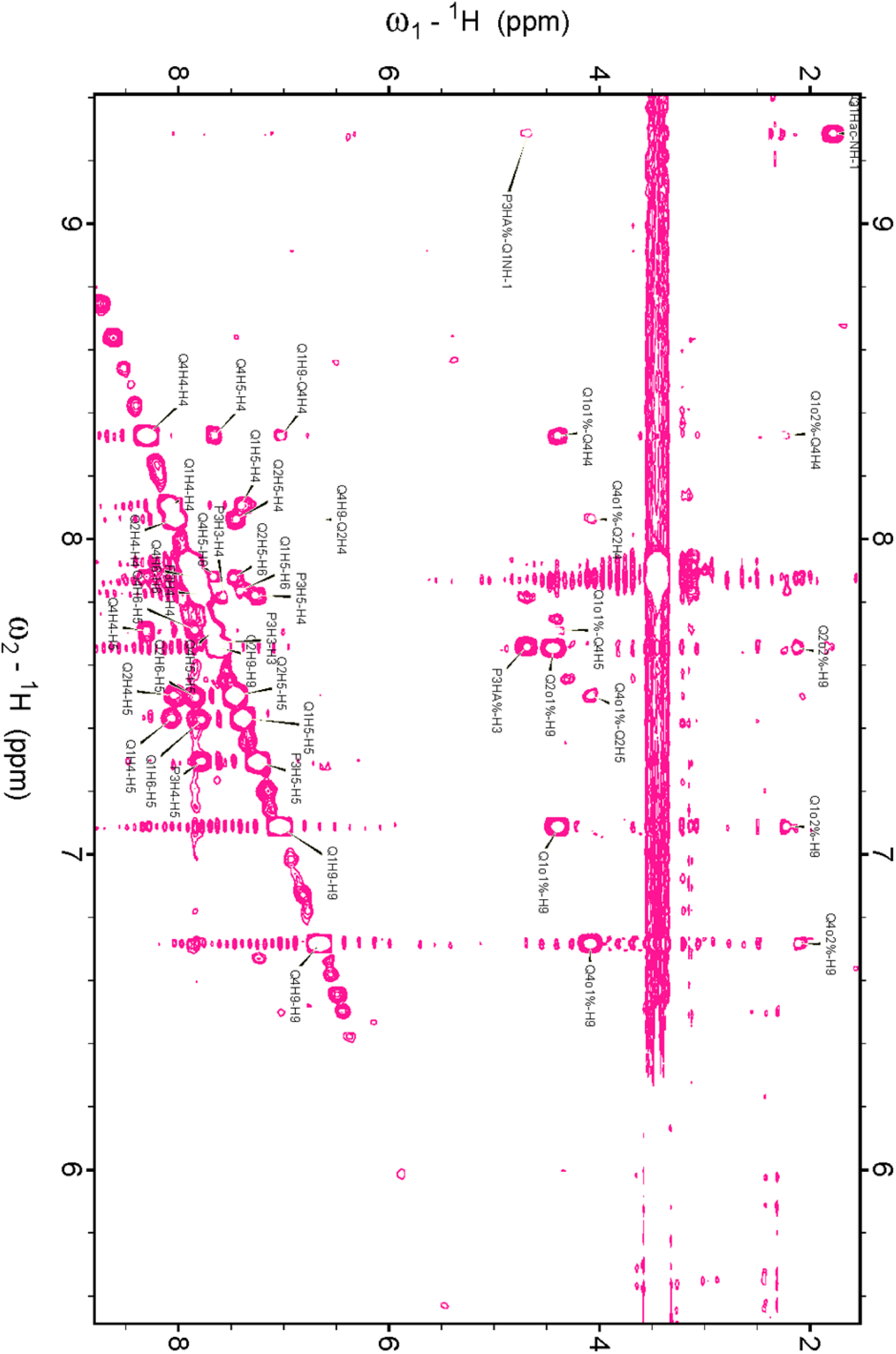
^1^H-^1^H NOESY of QQPQ on a Bruker Avance-III 800 MHz; middle region containing the aromatic protons in the foldamer core (H4, H5, H6, H9). Sample matrix: 300 µM QQPQ in d6-DMSO.

**Figure S129.**
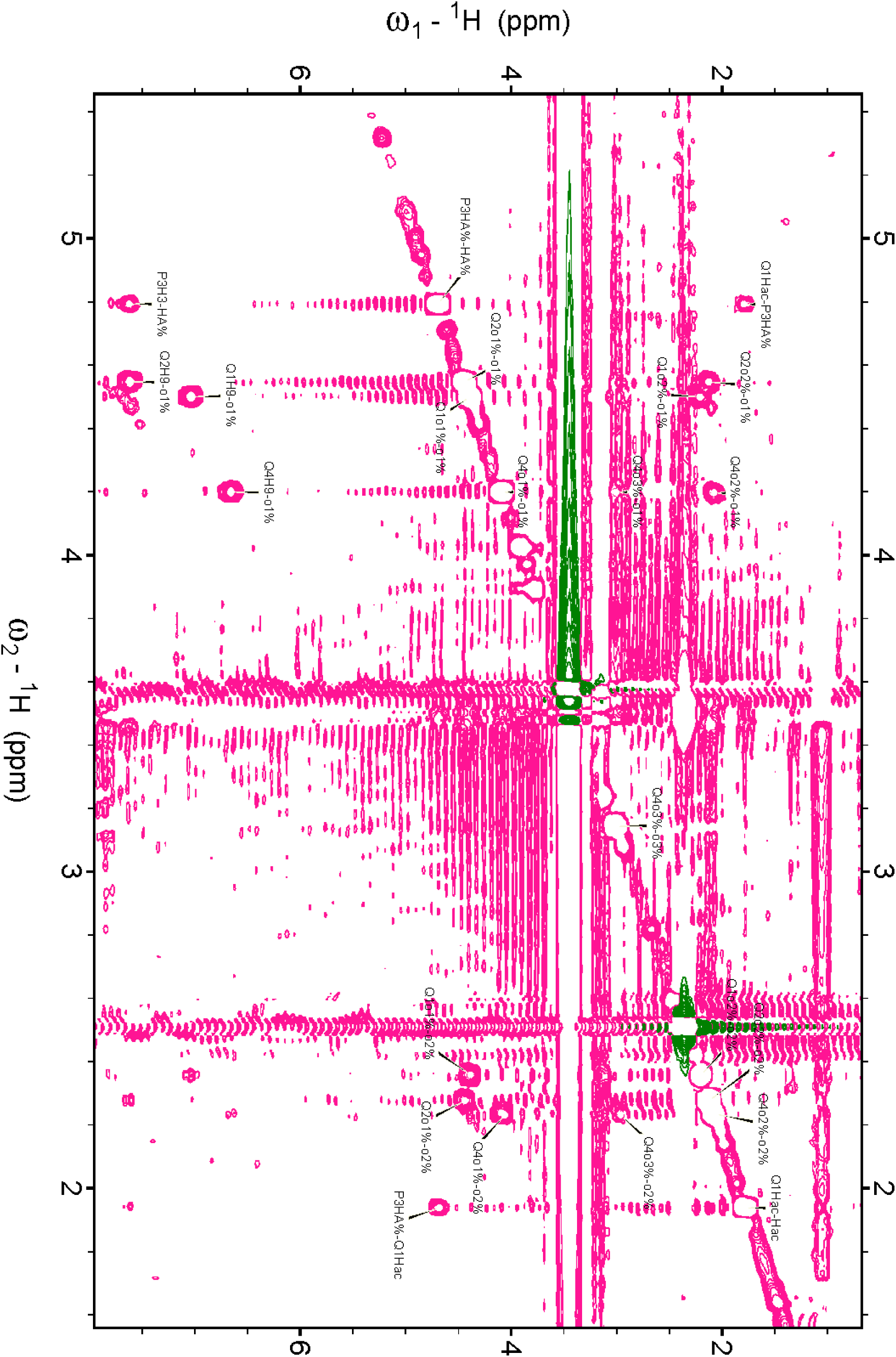
^1^H-^1^H NOESY of QQPQ on a Bruker Avance-III 800 MHz; low shift region containing the aliphatic protons, which are mostly in the side chains(o1, o2, o3, HA, HAc). Sample matrix: 300 µM QQPQ in d6-DMSO.

**Figure S130.**
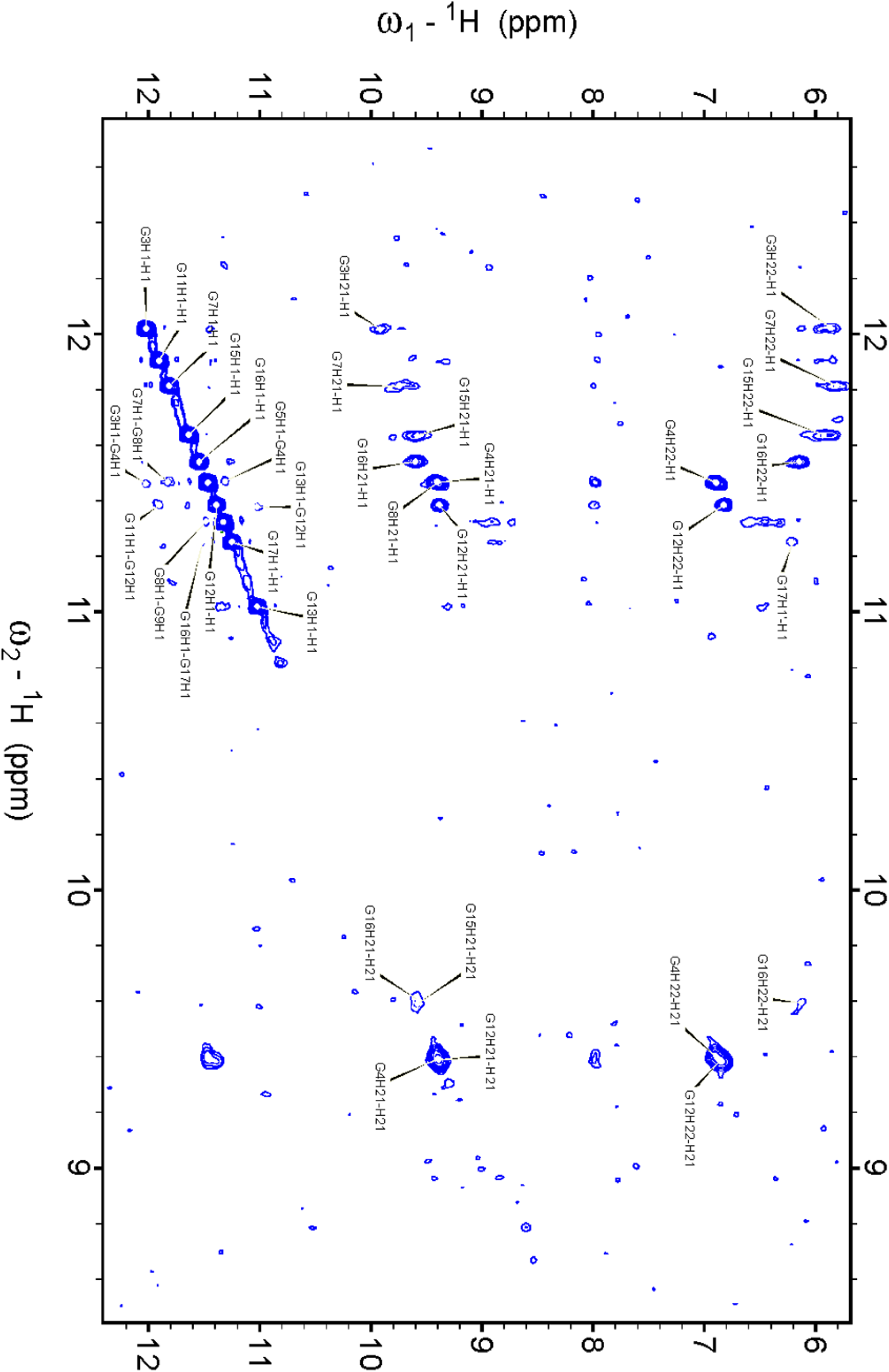
^1^H-^1^H NOESY of 2LK7 on a Bruker Avance-III 800 MHz; high shift region showing the guanine H1/H2 protons. Sample matrix: 100 µM 2LK7, 10 mM potassium phosphate buffer (pH 7), 90/10 H_2_O/D_2_O.

**Figure S131.**
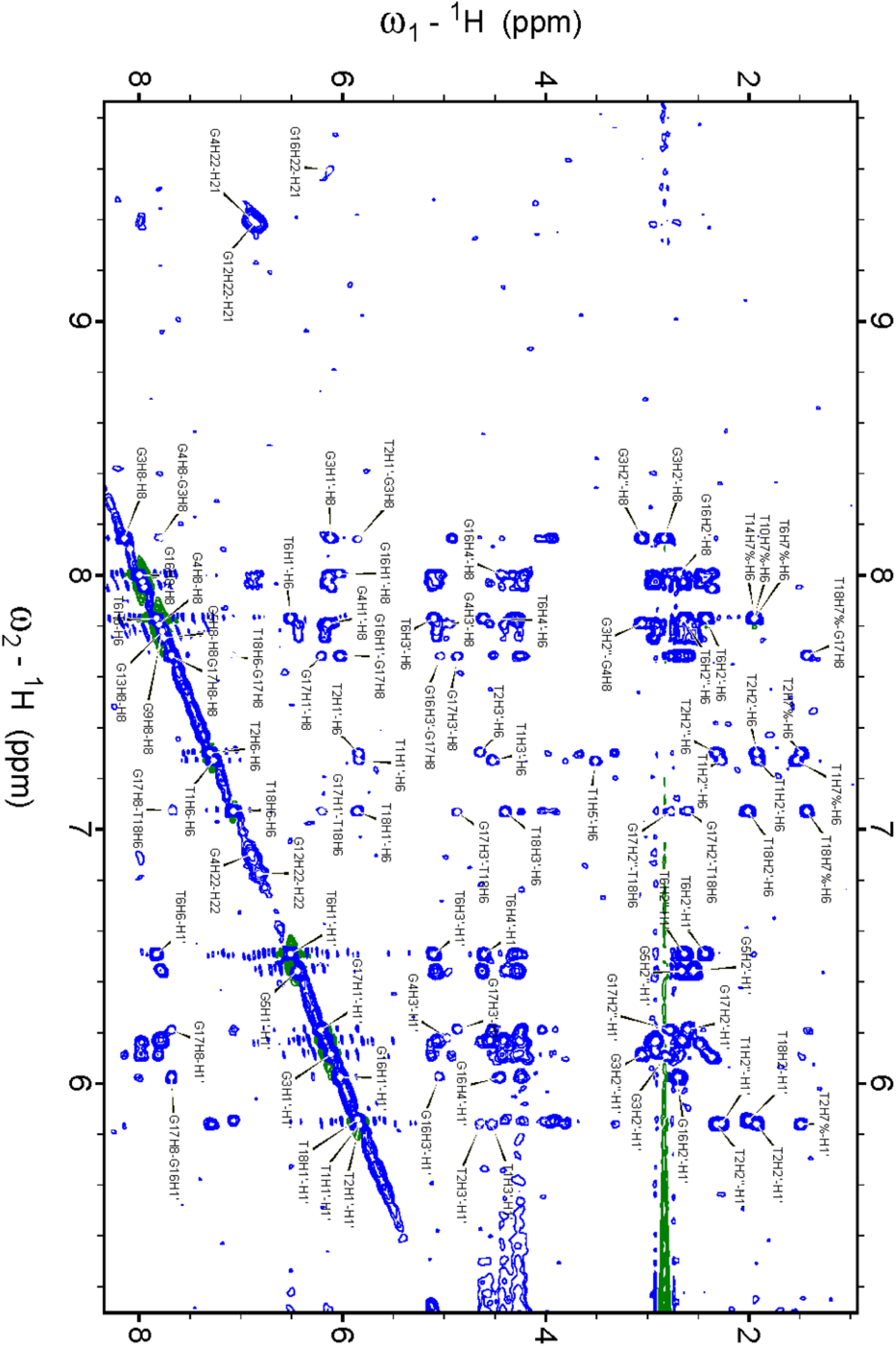
^1^H-^1^H NOESY of 2LK7 on a Bruker Avance-III 800 MHz; middle region containing the guanine H8, thymine H6 and sugar H1’ protons. Sample matrix: 100 µM 2LK7, 10 mM potassium phosphate buffer (pH 7), 90/10 H_2_O/D_2_O.

**Figure S132.**
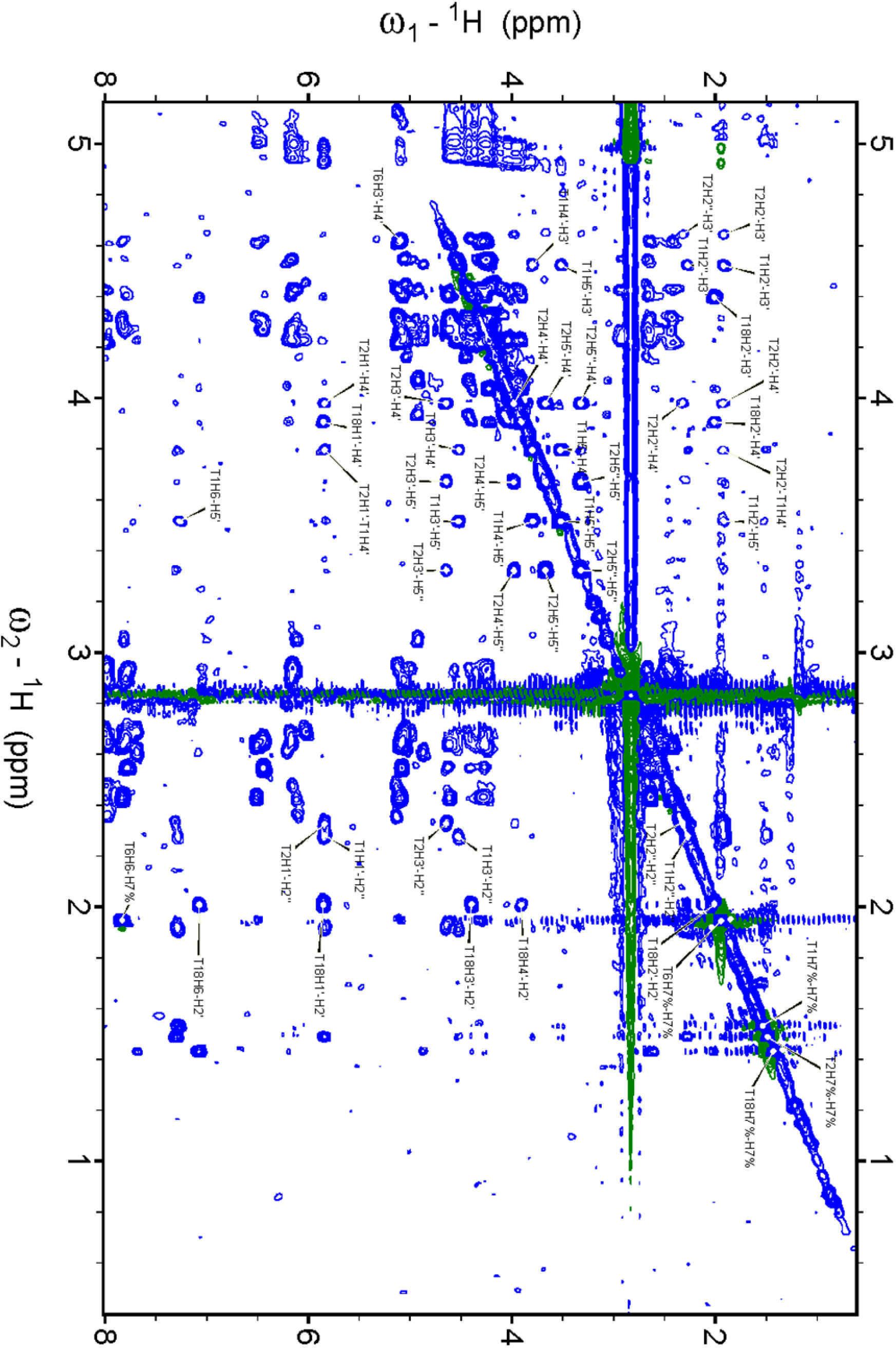
^1^H-^1^H NOESY of 2LK7 on a Bruker Avance-III 800 MHz; low shift region containing sugar protons (H2’ to H5’) and the thymine methyl group (H7). Sample matrix: 100 µM 2LK7, 10 mM potassium phosphate buffer (pH 7), 90/10 H_2_O/D_2_O.

**Figure S133.**
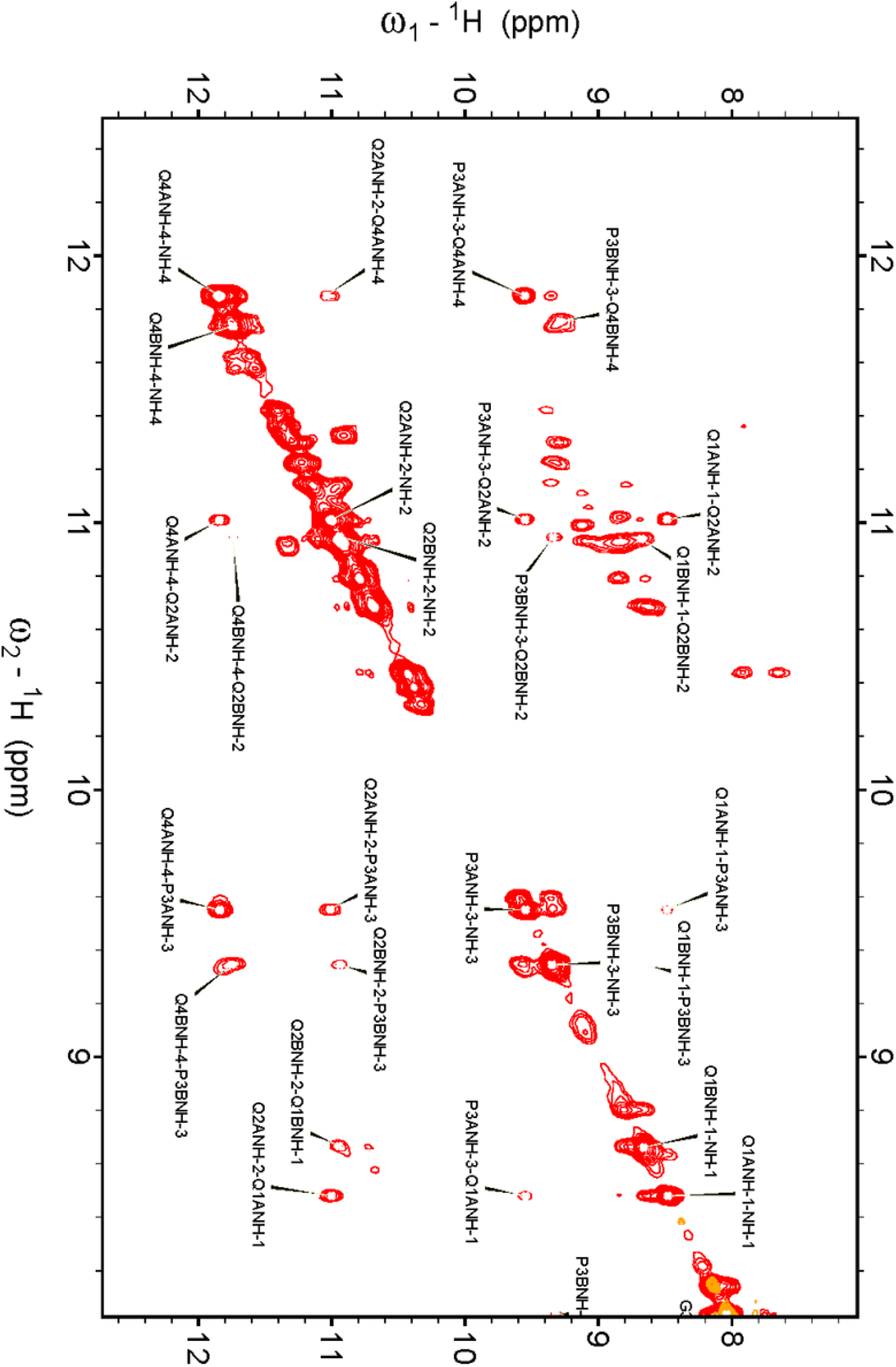
^1^H-^1^H NOESY of 2LK7/QQPQ on a Bruker Avance-III 800 MHz; high shift region containing guanine H1 protons and foldamer NH protons. Sample matrix: 100 µM 2LK7, 300 µM QQPQ, 10 mM potassium phosphate buffer (pH 7), 90/10 H_2_O/D_2_O.

**Figure S134.**
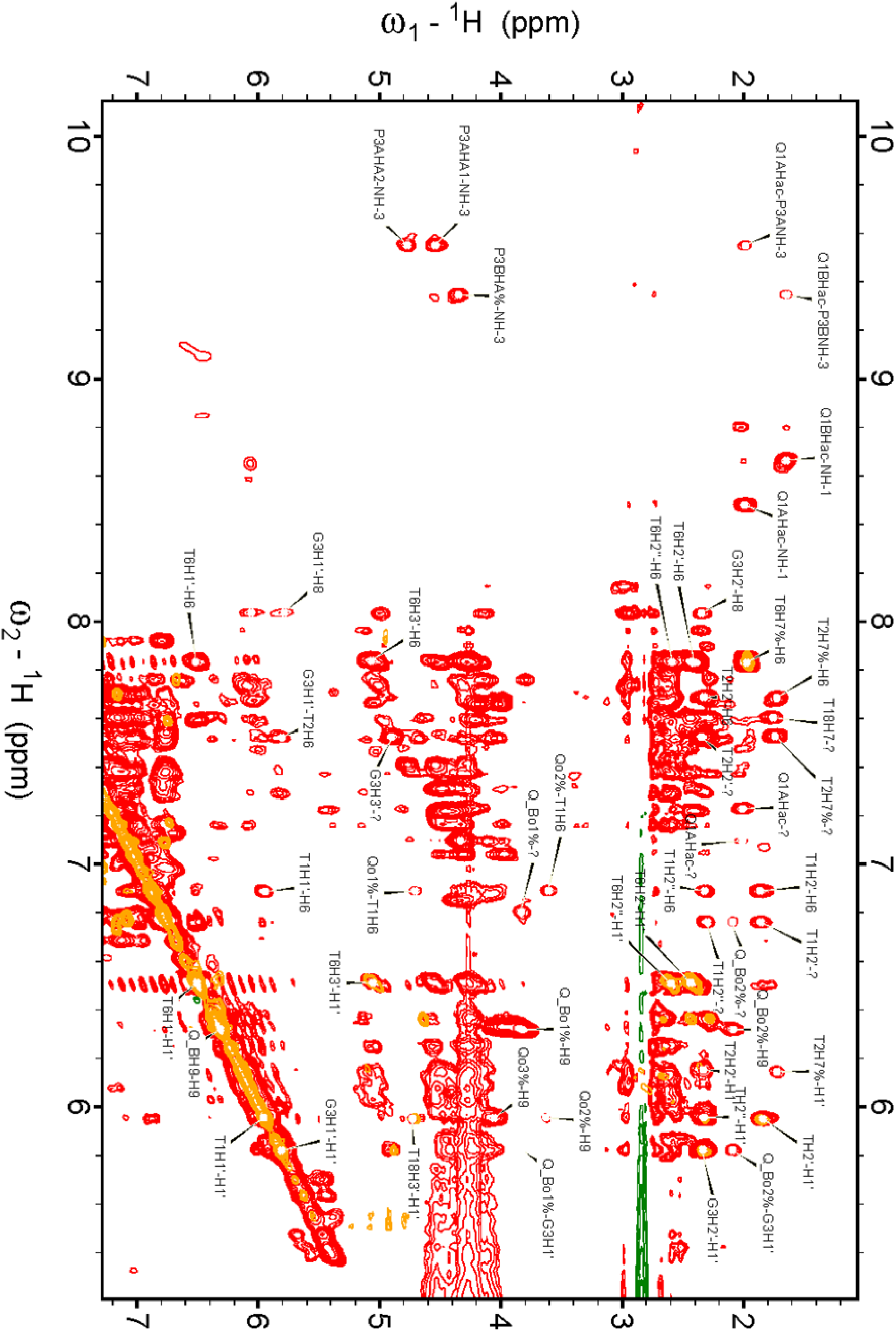
^1^H-^1^H NOESY of 2LK7/QQPQ on a Bruker Avance-III 800 MHz; middle region containing aromatic protons of nucleobases and foldamer (as well as H1’ sugar protons). Sample matrix: 100 µM 2LK7, 300 µM QQPQ, 10 mM potassium phosphate buffer (pH 7), 90/10 H_2_O/D_2_O. The orange overlay is a TOCSY spectrum of the same system.

#### Molecular dynamics

##### Methods

###### Force field modifications for the foldamer

The single-point energy of the foldamer structure was calculated using quantum mechanical (QM) density functional theory (DFT) with the B3LYP functional and 6-31G(d) basis set, employing the ORCA 6.0 software package, then converted to the molden file format.^8–18^ Two-stage Restrained Electrostatic Potential (RESP) atomic charge calculations were then performed in Multiwfn.^19,20^ An initial antechamber file was generated with Amber atom types, -2 formal charge (accounting for two phosphates), ^21,22^ wherein AM1-BCC partial charges were then replaced with those of the RESP calculation. A prepin file was generated using prepgen to define the topology and molecular parameters of QQPQ, and the force field parameters were subsequently derived using parmchk2.^23^

###### Complex preparation

The complexes were prepared in PyMOL (3.0.0, Schrödinger, NY, USA) from the crystal structure. The latter was first cleaned up: a single 222T unit was kept, extra magnesium and potassium ions were removed (only the 2 tetrad-bound K^+^ were conserved), as well as the second QQPQ bound on the loop at the interface with other units. Missing 3’ and 5’ thymine atoms were added at the system preparation step (*vide infra*).

Variant complexes were then generated by adding (on the 5’-tetrad) and/or flipping (binding through its C-instead of N-terminus) and/or mirroring (to its right-handed enantiomer) QQPQ, where necessary. A total of 8 complexes were prepared and named following the nomenclature:

- 3 or 5, to designate the 3’-tetrad or 5’-tetrad binding interface
- N or C, for N-terminus or C-terminus QQPQ binding interface
- L or R, for left-handed or right-handed enantiomer

Hence, the “original” crystal structure is 3NM, while e.g. 3NM/5CM refers to a 2:1 complex, with the L enantiomer bound through it N-terminus on the 3’-tetrad, and C-terminus on the 5’ end.

For the Hybrid-1 structural variant of 5YEY, the structure of 2JSM was used as a base. The extra 5’ residues were removed and the 5’O capped with a hydrogen. The residues were renumbered and A18 was mutated to a T18. Potassium cations were positioned at their expected coordination sites, in between tetrads. QQPQ was stacked on top of the 5’ tetrad with a random orientation.

##### MD preparation

The systems were then prepared for molecular dynamics using the Leap program from the Amber24 suite ^23^. The OL21 force field was used for DNA.^24,25^

The structure was explicitly solvated in a truncated octahedral box of water molecules, using the OPC model with the ad hoc Li/Merz ion parameters of atomic ions (12-6 set),^26,27^ with a minimum of 14 Å between the solute and the box edge. K^+^ and Cl^-^ ions were added to neutralize the system and adjust the ionic strength to 100 mM. The number of ions was determined SLTCAP method, using Equation S1 simplified by Machado and Pantano into Equation S2, where ν_*w*_ is the water volume of the simulation box in reduced units, *c*_0_ the salt concentration, *Q* the total charge of the complex, and 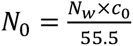 with *N*_*w*_ the number of water mole
   cules in the simulation box.^28^

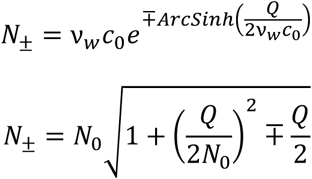

Note that the simpler SPLIT method described by Machado and Pantano cannot be applied as our system does not satisfy the *N*_0_ ≫ *Q* condition; however it yields identical values.

###### Molecular dynamics

All simulation steps were performed with pmemd.cuda (v. 18.0) from the CUDA version of AMBER,^29–31^ on an NVIDIA H100 PCIe Tensor core GPU (CUDA version: 12.4) from the DOREMI CALI v3 cluster of the *Mésocentre de Calcul Intensif Aquitain* (Université de Bordeaux, France). The system was minimized for 20000 cycles using the steepest descent algorithm for the first 4000 steps and the conjugate gradient for the next 16000 steps. The system was then heated at constant volume from 0 to 298 K over 18 ps then kept for 2 ps at the final temperature, using a time step of 2 fs, the Langevin thermostat with a 2.0 ps^-1^ collision frequency and a different seed for the pseudo-random number generation for every run to avoid synchronization artifacts,^32^ an 8 Å non-bonded cutoff, and the bonds involving hydrogen were constrained with the SHAKE algorithm. The system was further equilibrated five times at 298 K with the parameters above and the pressure kept at 1.0 bar with the Berendsen barostat,^33^ before production MD simulations were run for a microsecond.

###### Data analysis

Trajectory files cleanup, alignment, filtering and analysis was performed with R 4.4.2, using the bio3d package and custom scripts.^34^ The determination of RMSD was performed with the rmsd function. All atom-atom were measured with the dist.xyz function, complemented with the com function to determine center of masses for ring-ring distances. Molecular structure images were created in PyMOL 3.0.

##### QQPQ/222T

###### Structures mid-production

**Figure S135.**
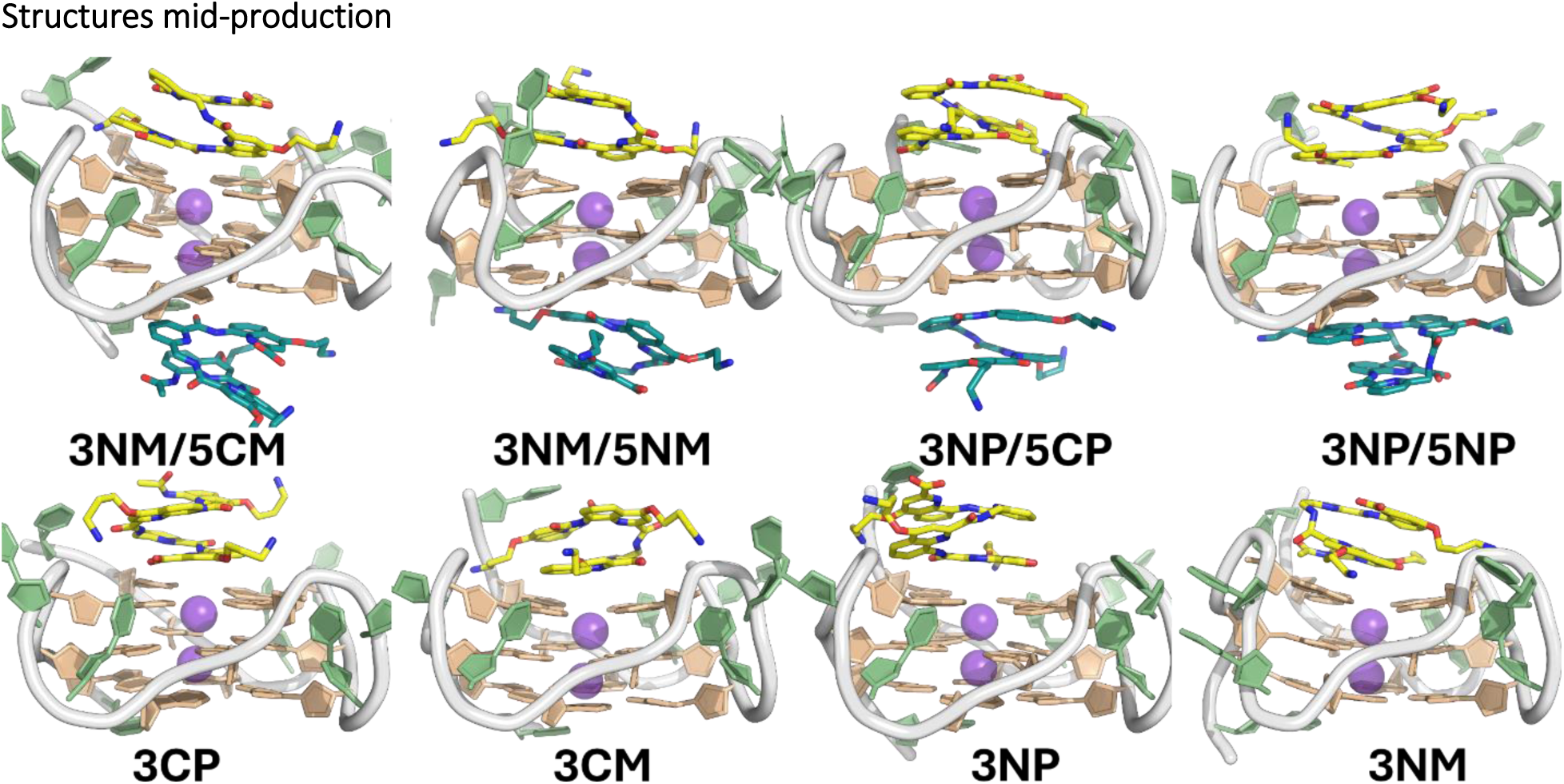
Minimized, heated and equilibrated structures of the 1:1 (bottom) and 2:1 (top) QQPQ:222T complexes submitted to 1 microsecond MD simulations, here shown after 500 ns.

###### RMSD

**Figure S136.**
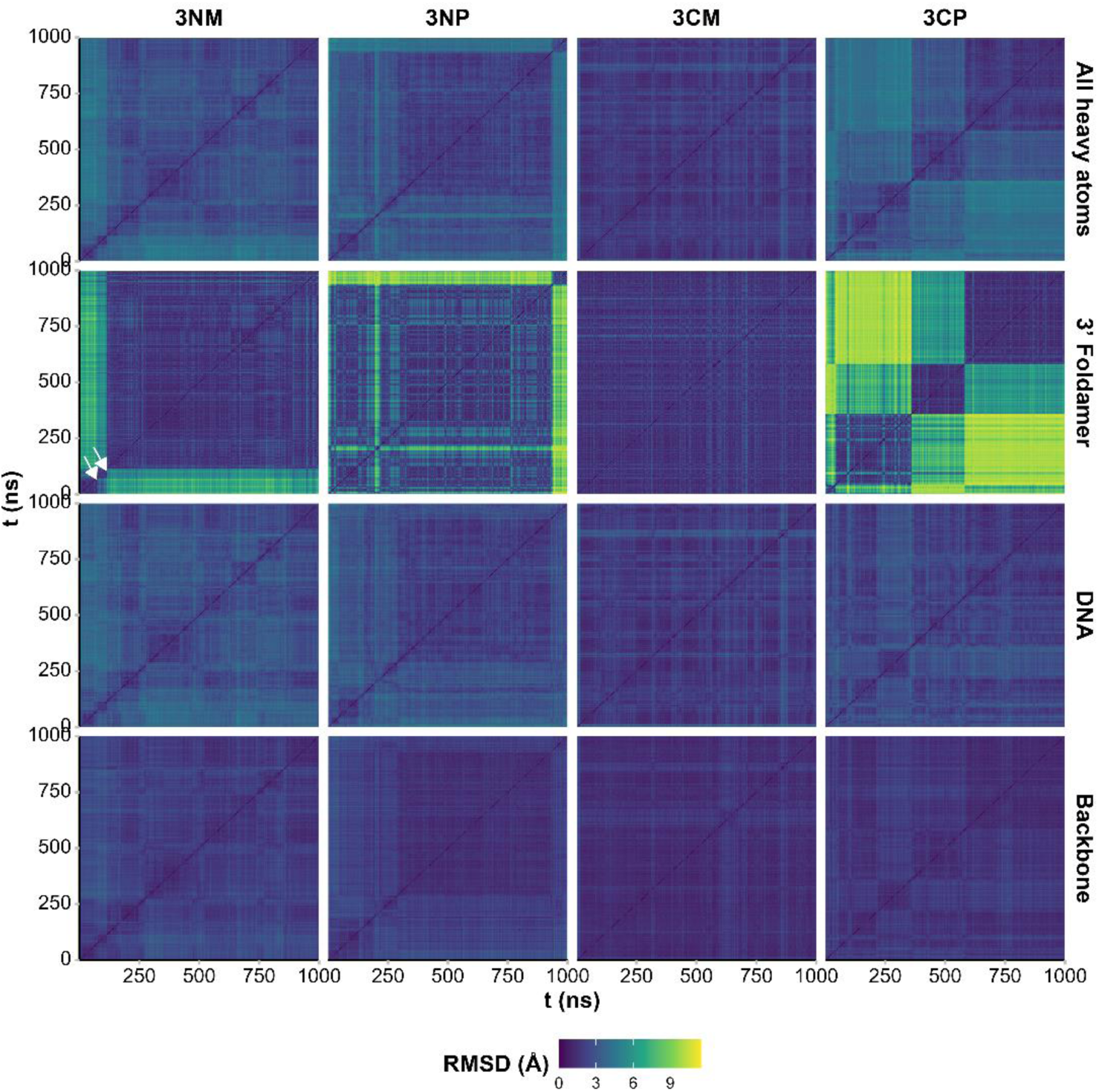
Pairwise RMSD of 1:1 QQPQ:222T complexes calculated on all heavy atoms, and only on the foldamer, the DNA, and the DNA backbone. Lighter colors points to structural changes, while similar-colored squares along the diagonal indicate a relative structural stability during the corresponding time range. An example of foldamer rotation events are indicated by white arrows for 3NM.

**Figure S137.**
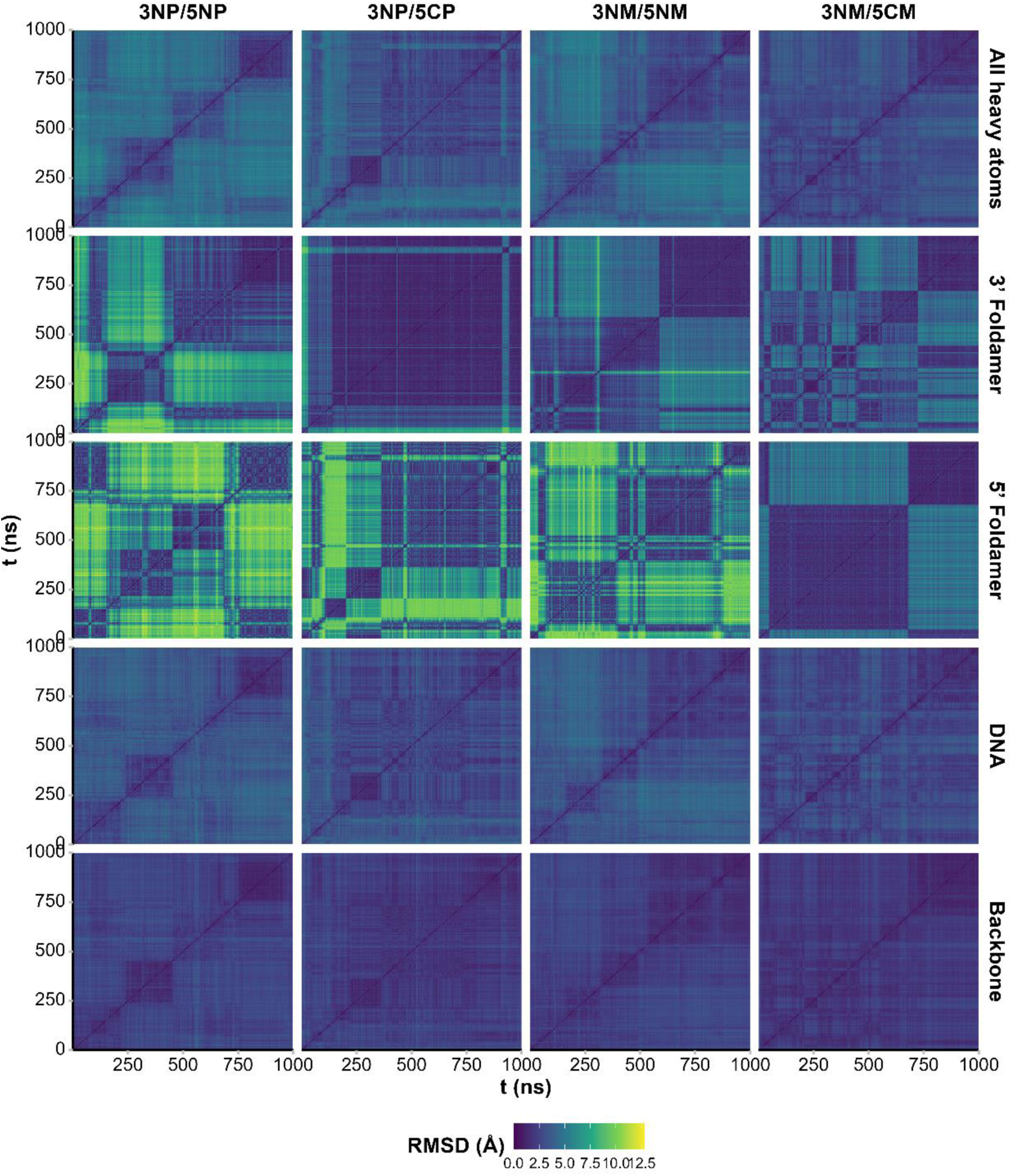
Pairwise RMSD of 2:1 QQPQ:222T complexes calculated on all heavy atoms, and only on either foldamer, the DNA, and the DNA backbone. Lighter colors points to structural changes, while similar-colored squares along the diagonal indicate a relative structural stability during the corresponding time range.

###### Tetrad stability

**Figure S138.**
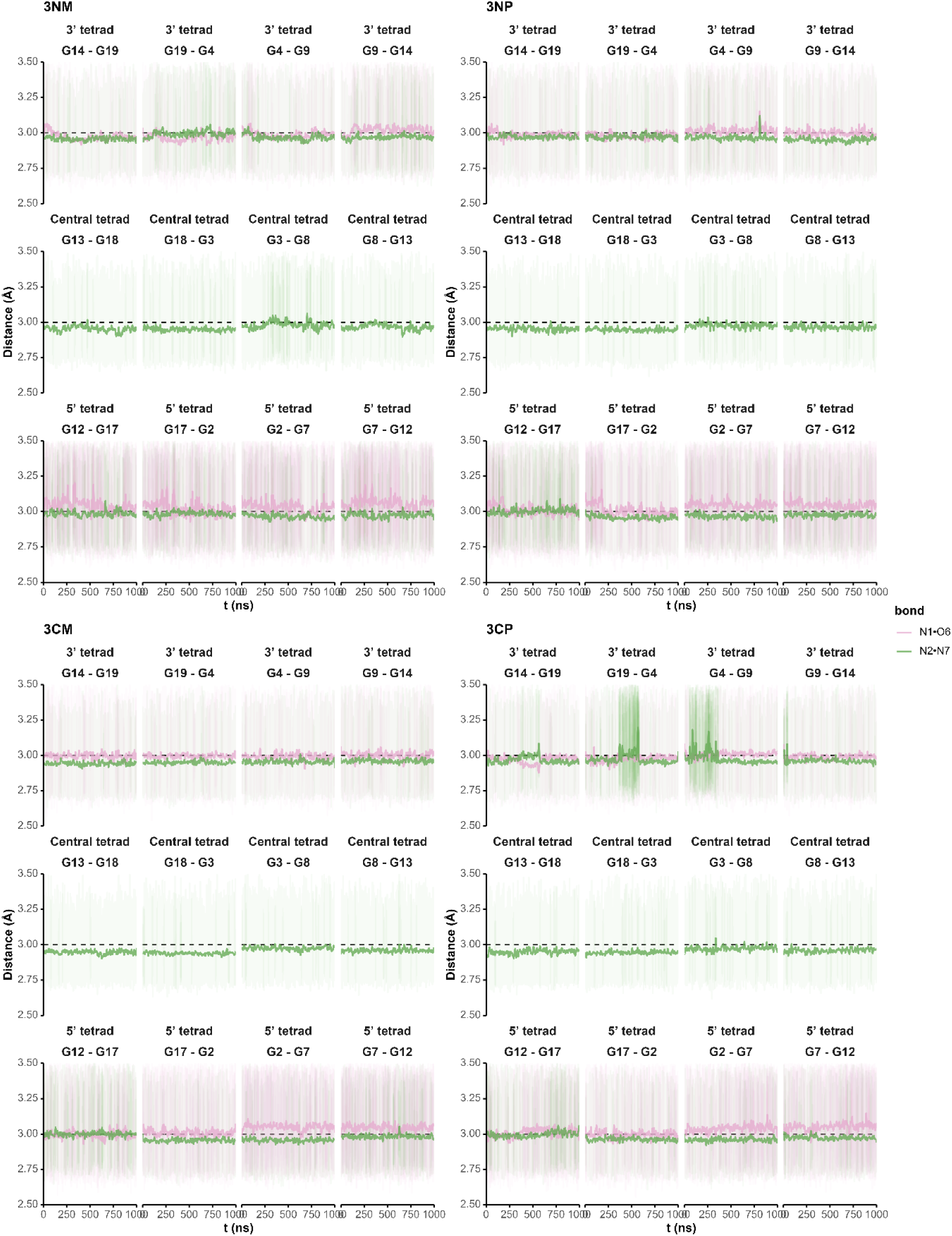
H-bonds within tetrads are stable for all 1:1 complexes, here shown through the donor-acceptor distance during the trajectory.

**Figure S139.**
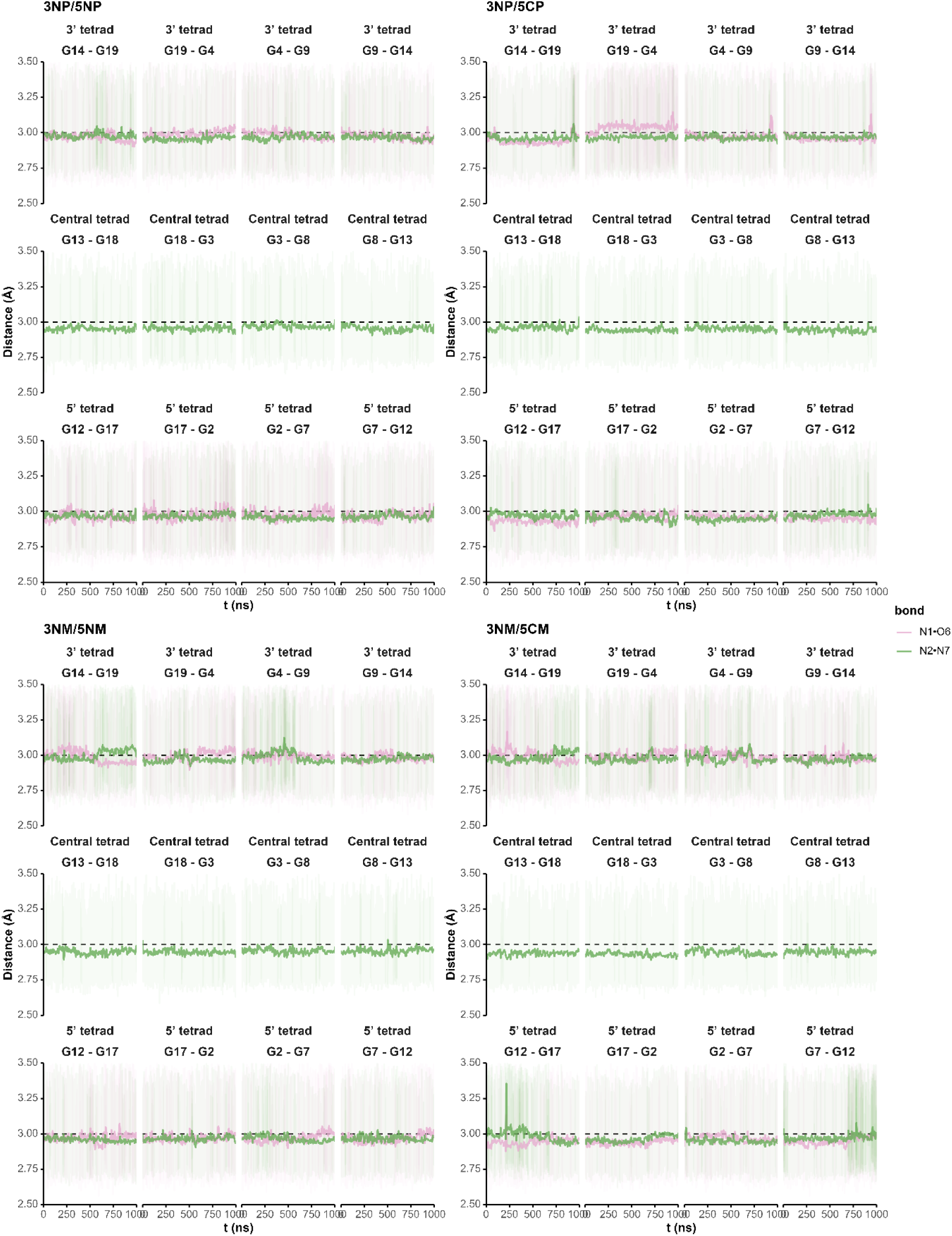
H-bonds within tetrads are stable for all 2:1 complexes, here shown through the donor-acceptor distance during the trajectory.

###### Side-chain H-bonding

**Figure S140.**
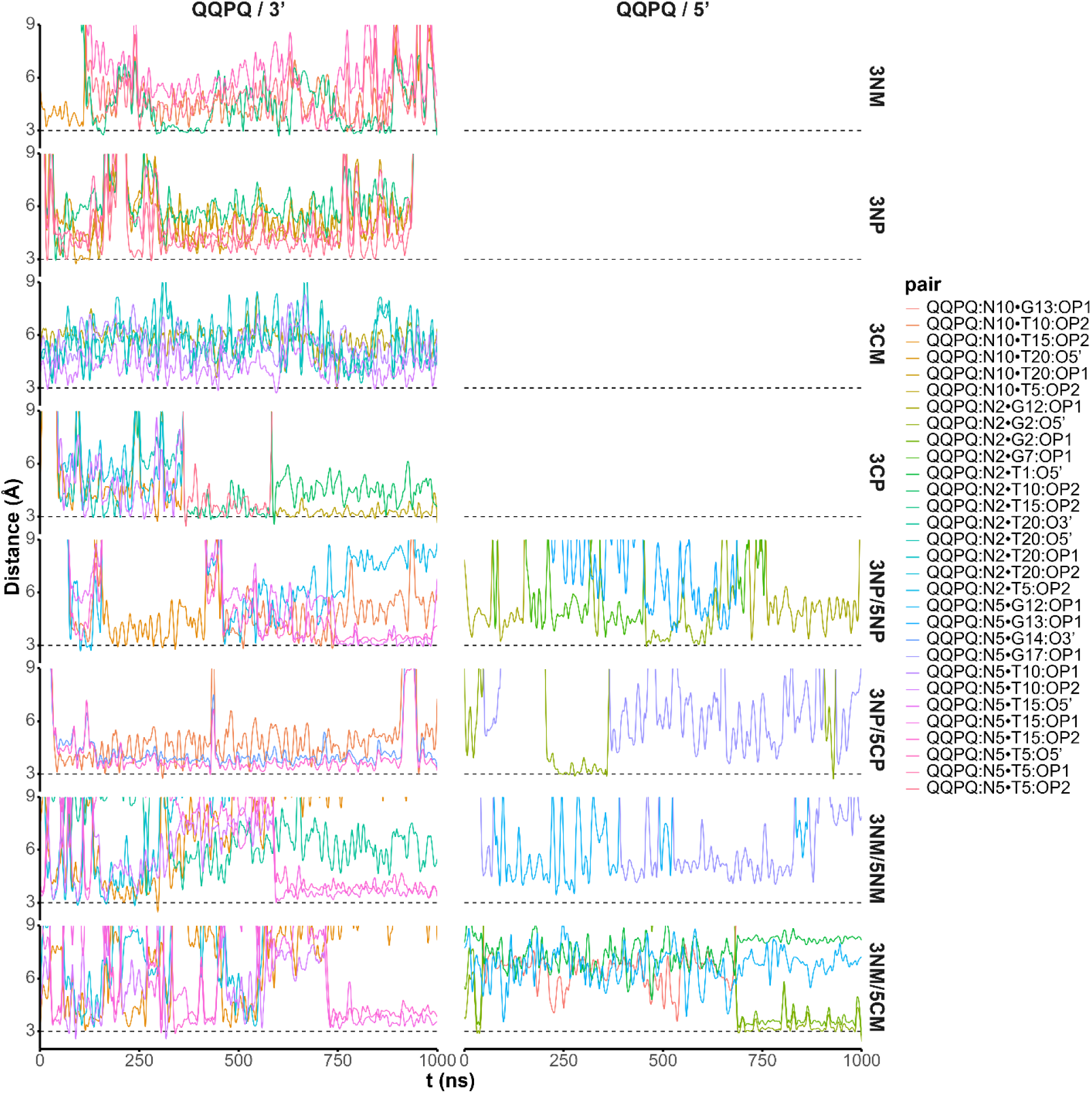
Donor-acceptor distances of H-bonds involving the ammonium groups of quinoline side chains and the DNA backbone. Only contacts for which the distance is below 3.1 Å with a frequency > 0.1 are shown for the sake of clarity.

**Figure S141.**
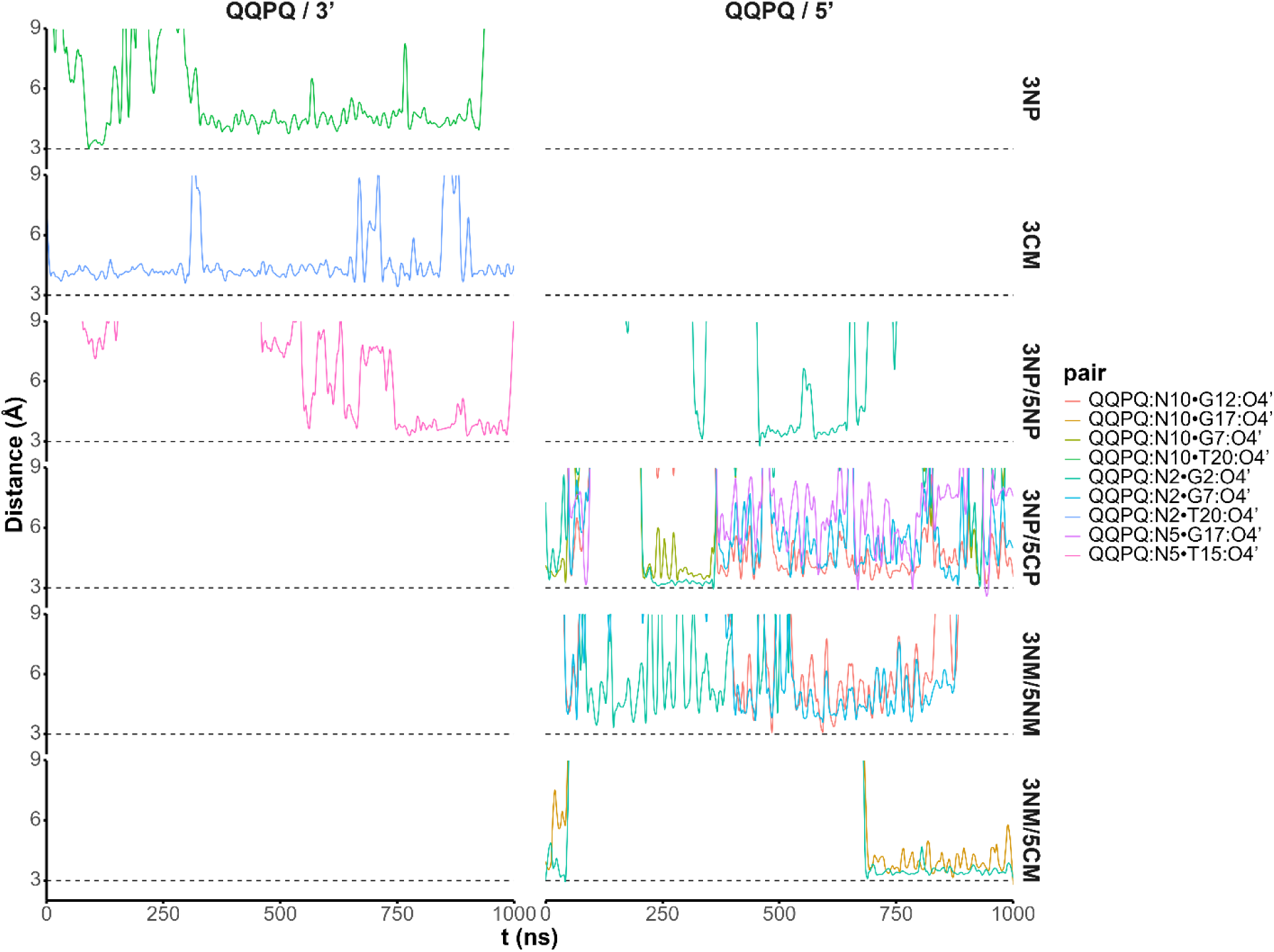
Donor-acceptor distances of H-bonds involving the ammonium groups of quinoline side chains and the DNA sugars. Only contacts for which the distance is below 3.1 Å with a frequency > 0.1 are shown for the sake of clarity.

**Figure S142.**
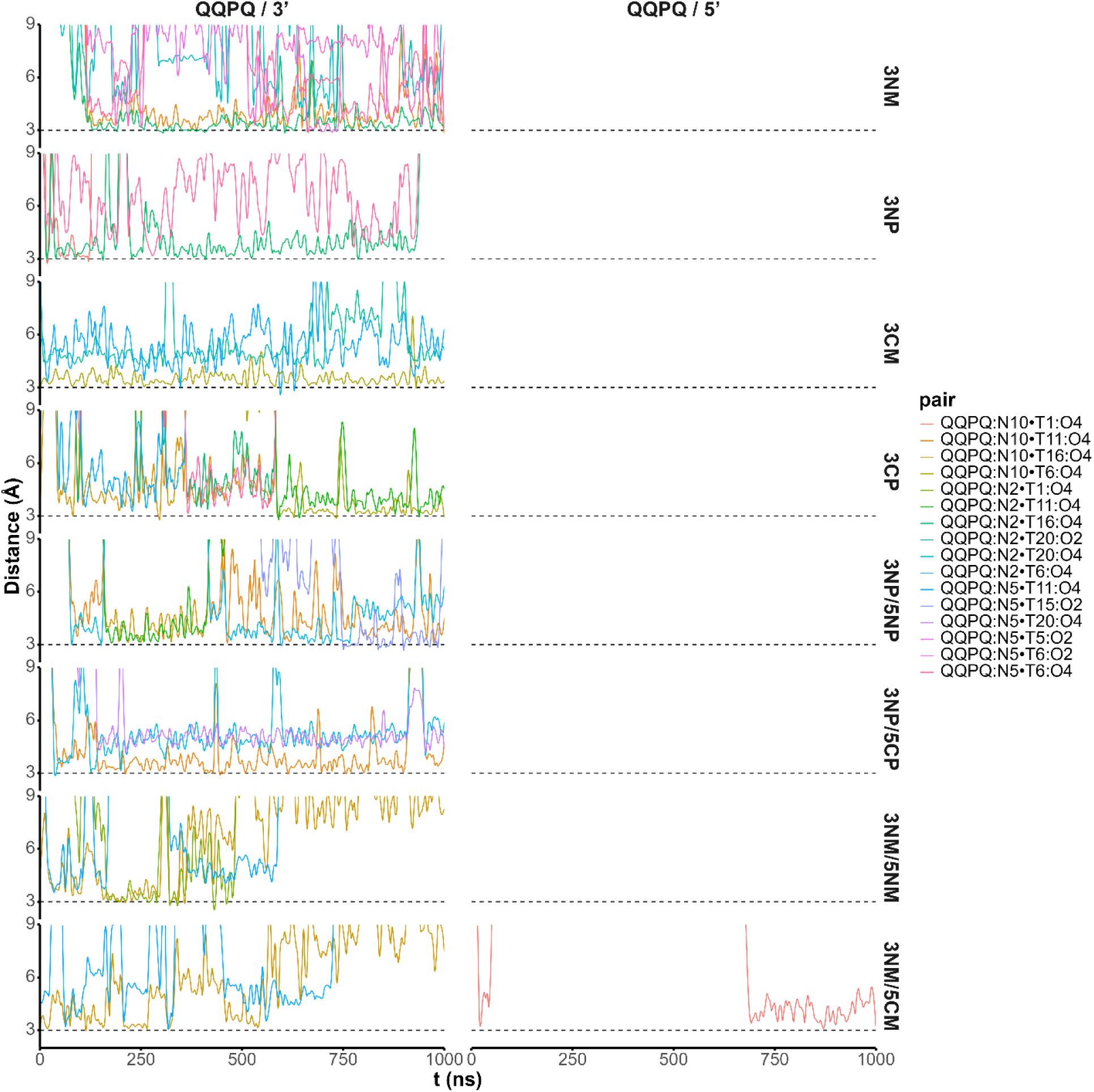
Donor-acceptor distances of H-bonds involving the ammonium groups of quinoline side chains and the DNA thymines from loop and termini (T1 and T20). Only contacts for which the distance is below 3.1 Å with a frequency > 0.1 are shown for the sake of clarity.

###### Stacking distances

**Figure S143.**
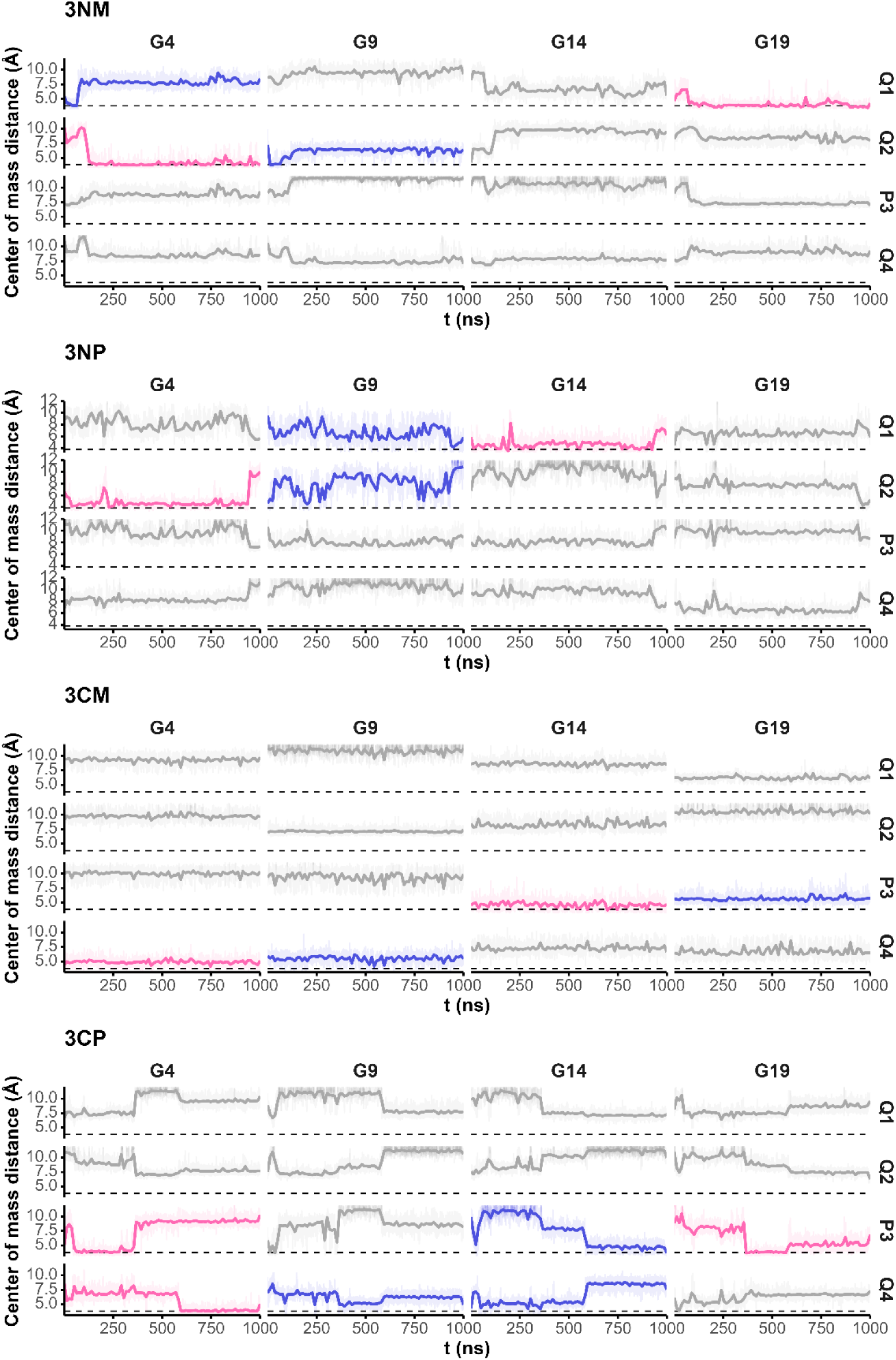
Distances between the aromatic cycles of QQPQ and the 3’ tetrad guanines, for the 1:1 complexes. Aromatic cycles are numbered from 1 to 4 from the N- to C-termini. The data is colored by frequency of stacking (pink: often stacked, purple: occasionally stacked, grey: generally not stacked), estimated with the distance between G18 and G19 (dashed line) used as reference of ‘proper’ π-stacking

**Figure S144.**
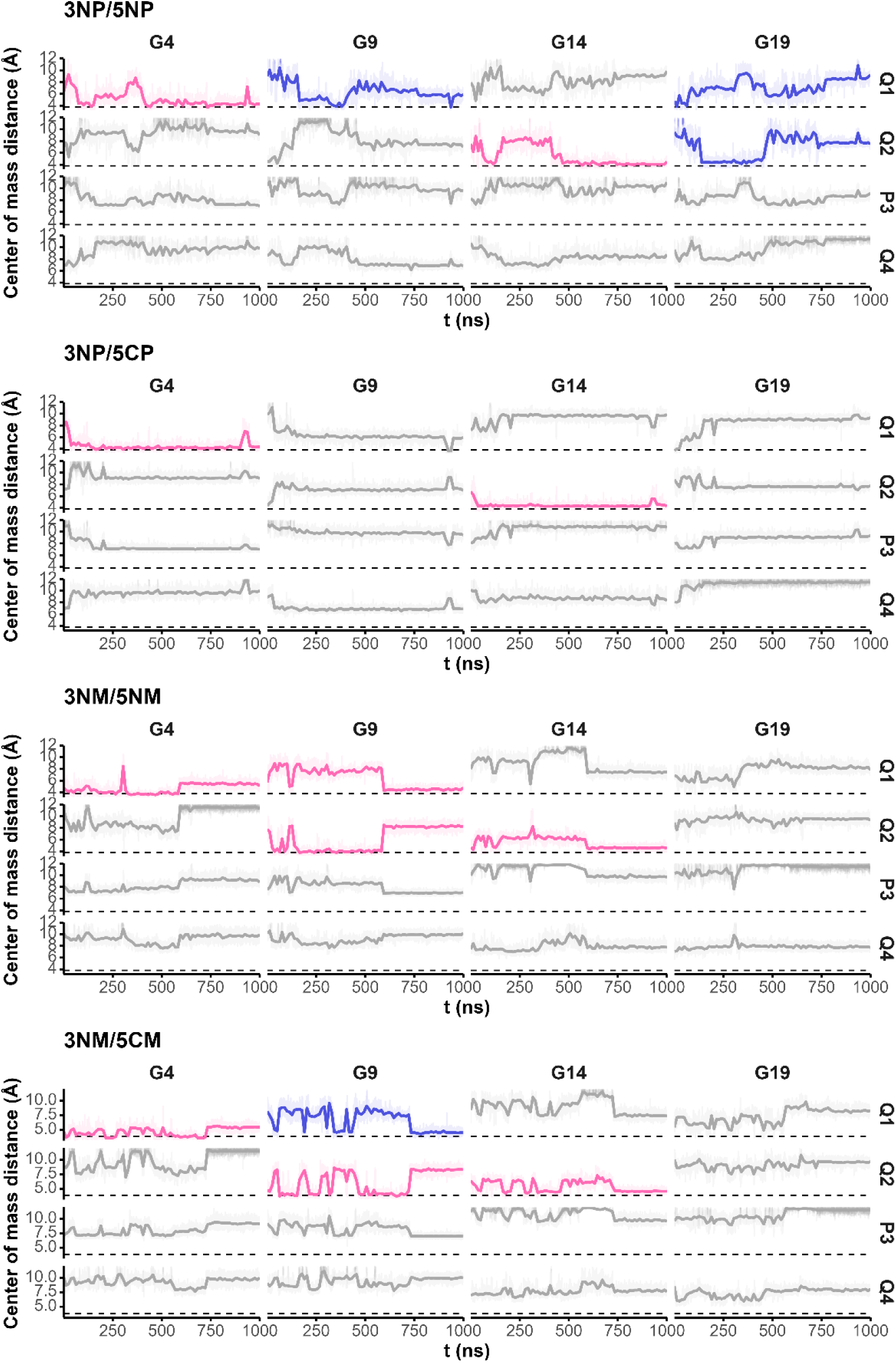
Distances between the aromatic cycles of QQPQ and the 3’ tetrad guanines, for the 2:1 complexes. Aromatic cycles are numbered from 1 to 4 from the N- to C-termini. The data is colored by frequency of stacking (pink: often stacked, purple: occasionally stacked, grey: generally not stacked), estimated with the distance between G18 and G19 (dashed line) used as reference of ‘proper’ π-stacking

**Figure S145.**
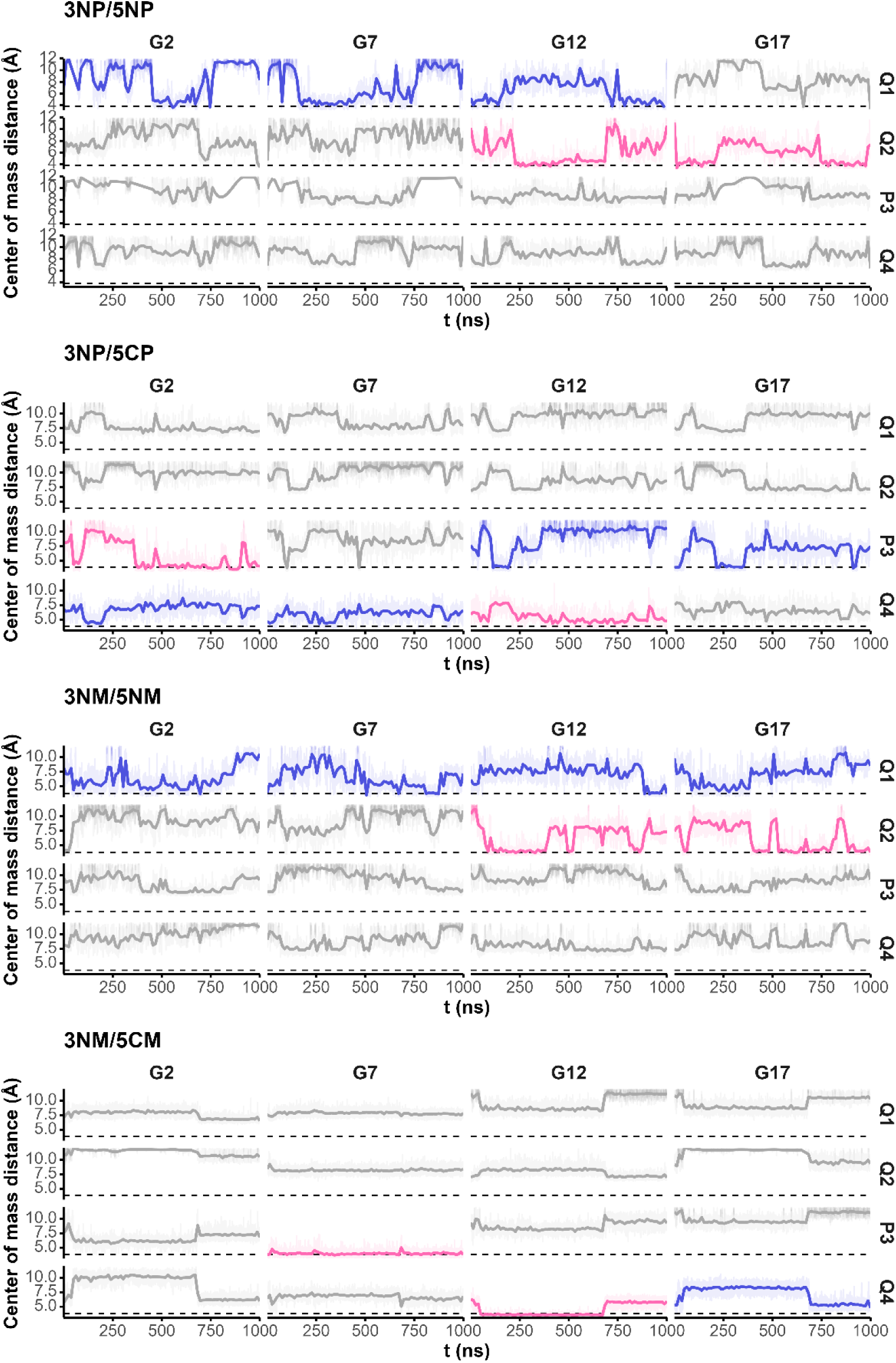
Distances between the aromatic cycles of QQPQ and the 5’ tetrad guanines, for the 2:1 complexes. Aromatic cycles are numbered from 1 to 4 from the N- to C-termini. The data is colored by frequency of stacking (pink: often stacked, purple: occasionally stacked, grey: generally not stacked), estimated with the distance between G18 and G19 (dashed line) used as reference of ‘proper’ π-stacking

###### QQPQ rotation

**Figure S146.**
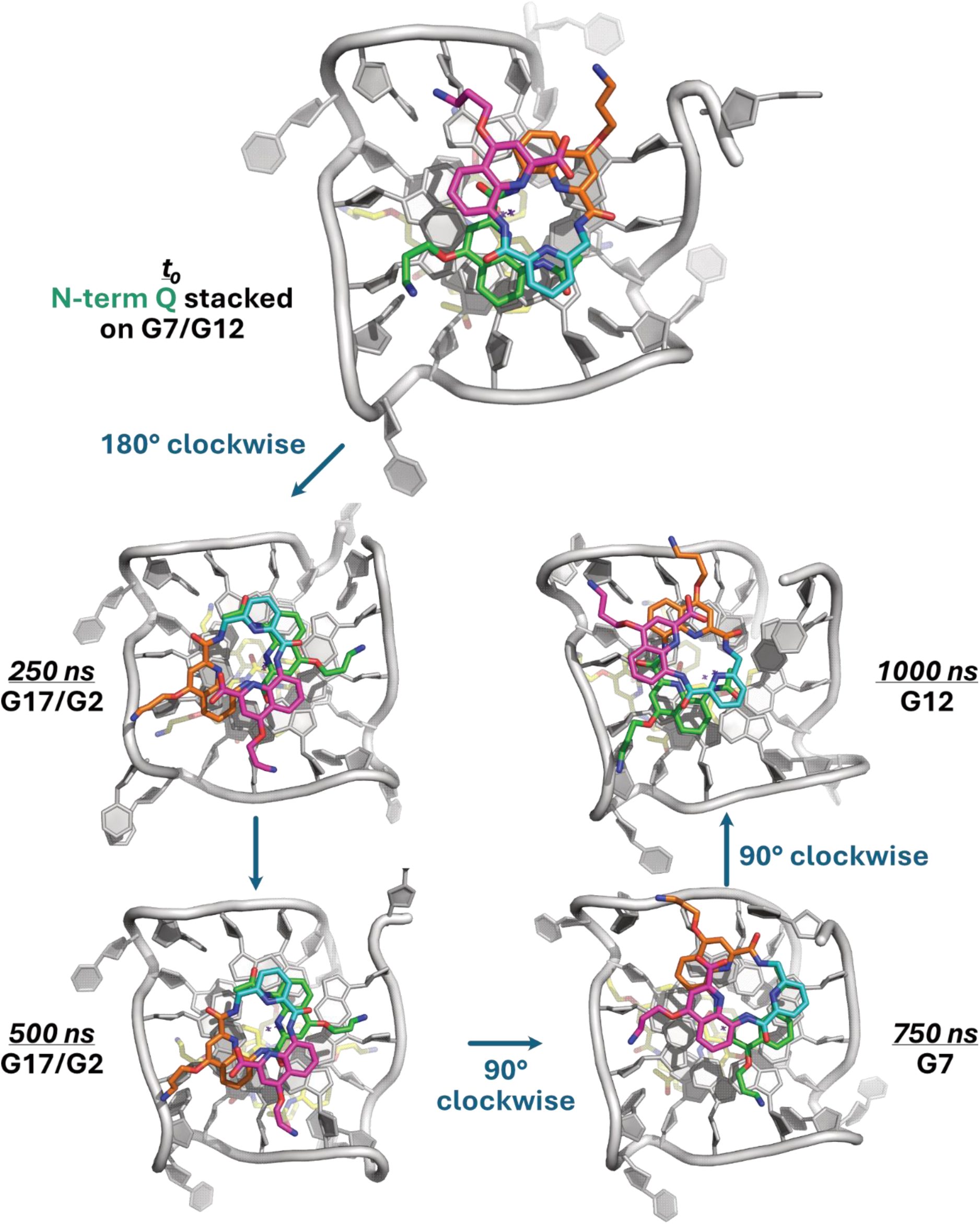
Example of a complete clockwise rotation of QQPQ (M enantiomer), here stacked by its N-terminal quinoline (green) on the 5’-end of 222T (complex 3NM/5NM). Note the very significant dynamics of 5’ and 3’ dTs during the simulation. Corresponding cycle-cycle distances are given in Figure S145 (see the sequential stacking for 3NM/5NM: Q1).

###### Thymine binding

**Figure S147.**
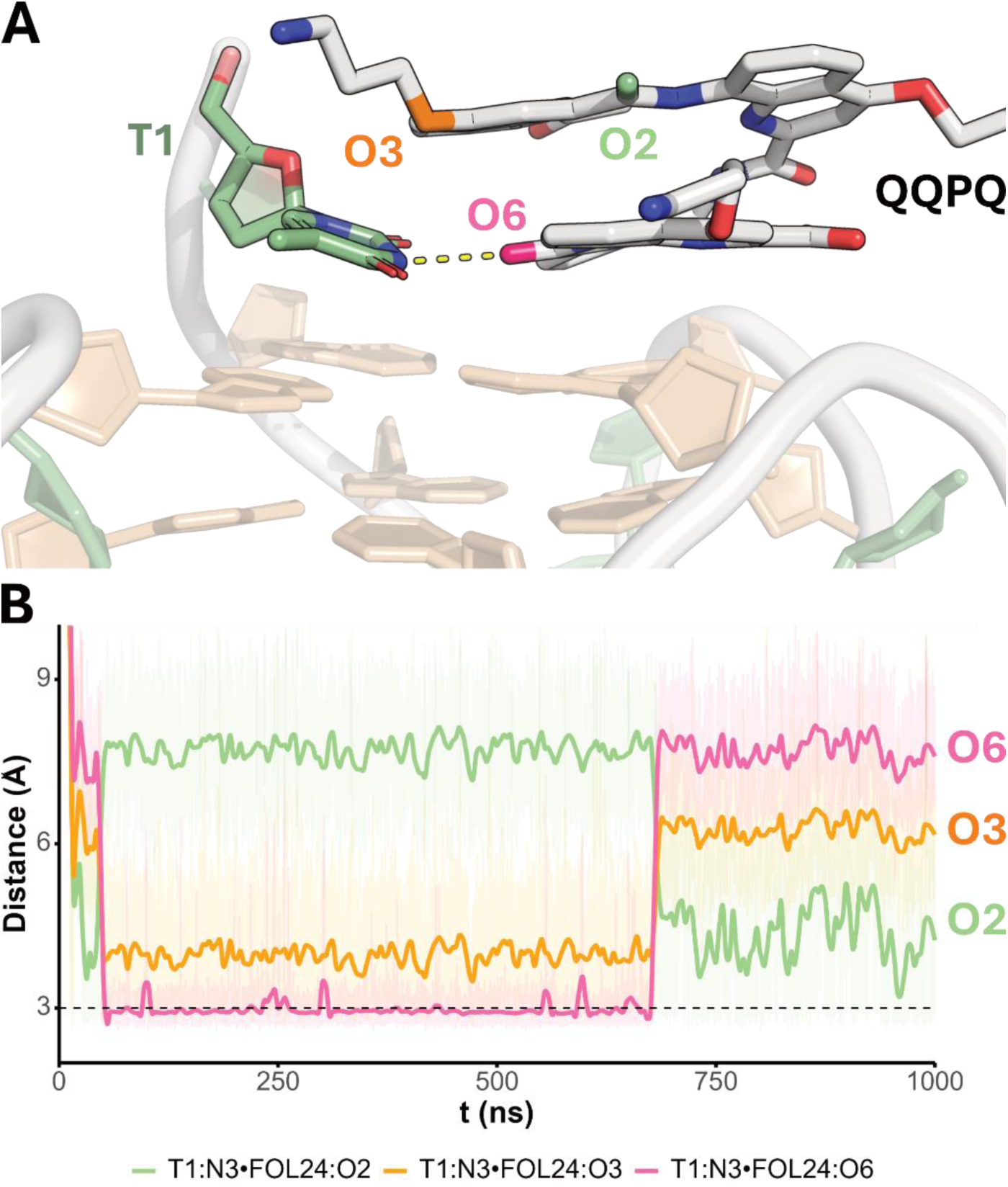
Example of discrete binding mode of QQPQ involving binding to a thymine. A. In 3NM5CM, QQPQ binds on the 5’-end by establishing an H-bond between its O6 (pink) and the N3 of dT1. B. This interaction has a significant lifetime, being maintained here for more than half a microsecond. This leads to an inhibition of the rotation of QQPQ, as visible in Figure S145 (see 3NM/5CM: Q4/G12)

###### RMSD

**Figure S148.**
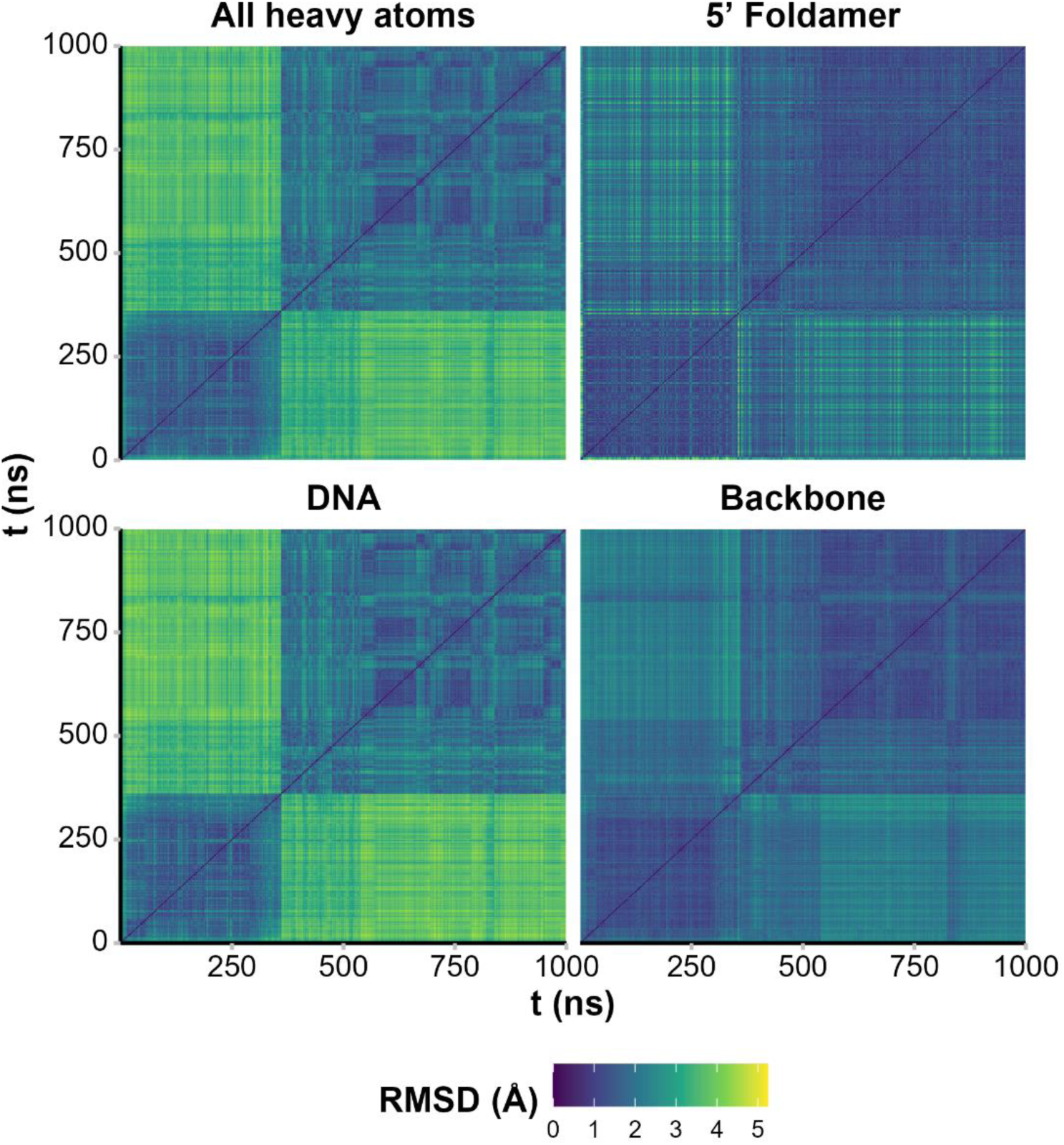
Pairwise RMSD of the 1:1 QQPQ:5YEY complexes calculated on all heavy atoms, and only on either foldamer, the DNA, and the DNA backbone. Lighter colors points to structural changes, while similar-colored squares along the diagonal indicate a relative structural stability during the corresponding time range.

###### QQPQ/5YEY T18 interactions

**Figure S149.**
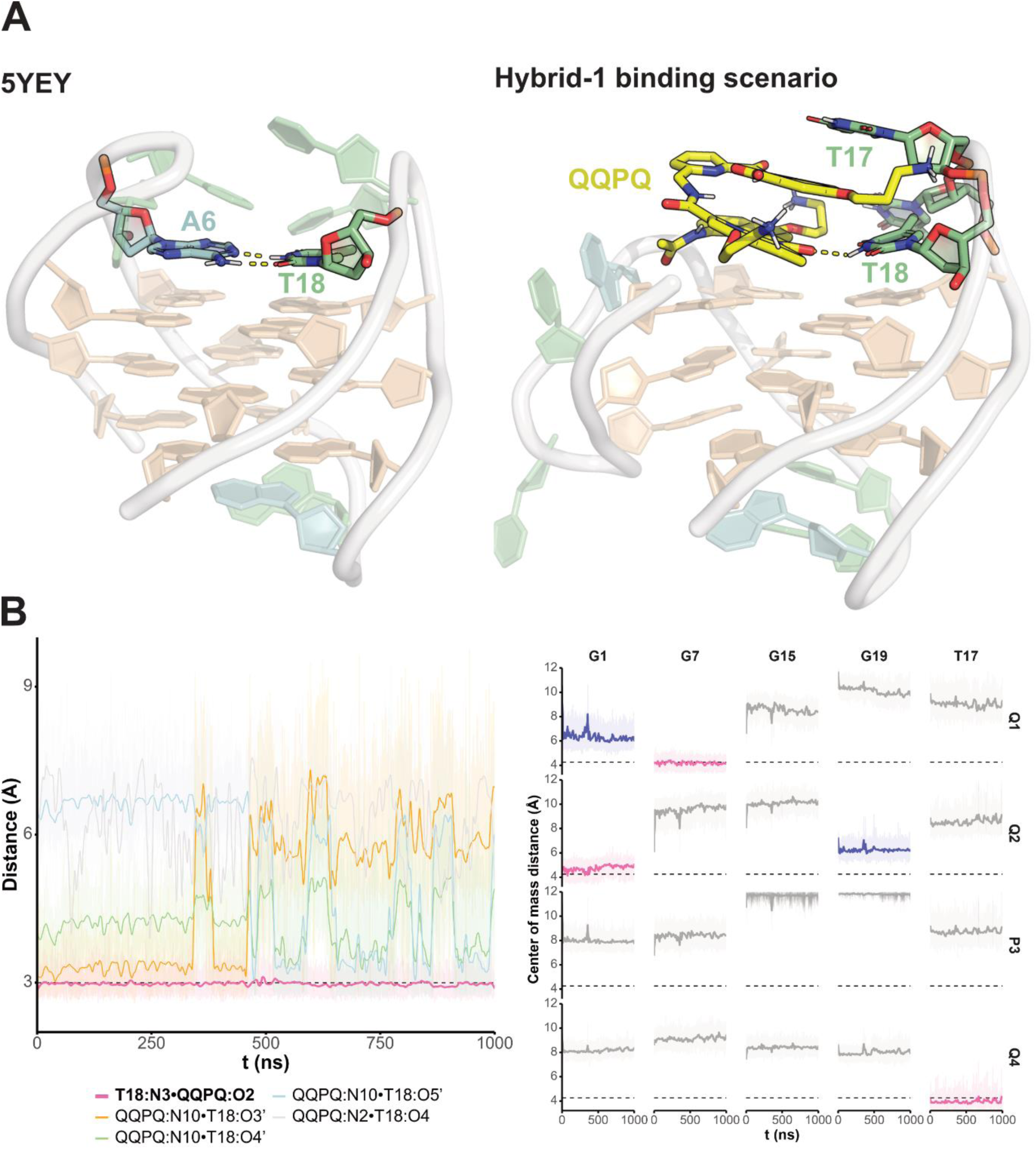
Exploring the binding of QQPQ to 5YEY converted to a hybrid-1 type loop arrangement. A. The deposited structure of 5YEY (left),^35^ and a random frame extracted from the MD simulation of its hybrid-1 topology complexed by QQPQ. In this scenario, the A6:T18 base pair is disrupted giving space above the 5’ tetrad for QQPQ to stack. Further, T18 is available to bind the amide oxygen at the N-terminus of QQPQ, reminiscent of the binding mode depicted in Figure S147. T17 stacks on top of the C-terminus of QQPQ. B. Distances between H-bond donor and acceptor groups of QQPQ and T18, showing the stability of the pairing between QQPQ and the N3 of T18. Only bonds formed (i.e. distance < 3.1 Å) more than 5% of the time are shown. C. Distances between the aromatic cycles of QQPQ and the 5’ tetrad guanines and T17 of 5YEY. Aromatic cycles are numbered from 1 to 4 from the N- to C-termini. The data is colored by frequency of stacking (pink: often stacked, purple: occasionally stacked, grey: generally not stacked), estimated with the distance between G20 and G21 (dashed line) used as reference of ‘proper’ π-stacking. The ligand is clearly locked in place, stacking with its first quinoline to G7, and being capped by T17, which stacks on Q4.

## Full wwPDB X-ray Structure Validation Report

### 1 Overall quality at a glance

The following experimental techniques were used to determine the structure:

*X-RAY DIFFRACTION*

The reported resolution of this entry is 2.51 Å.

Percentile scores (ranging between 0-100) for global validation metrics of the entry are shown in the following graphic. The table shows the number of entries on which the scores are based.

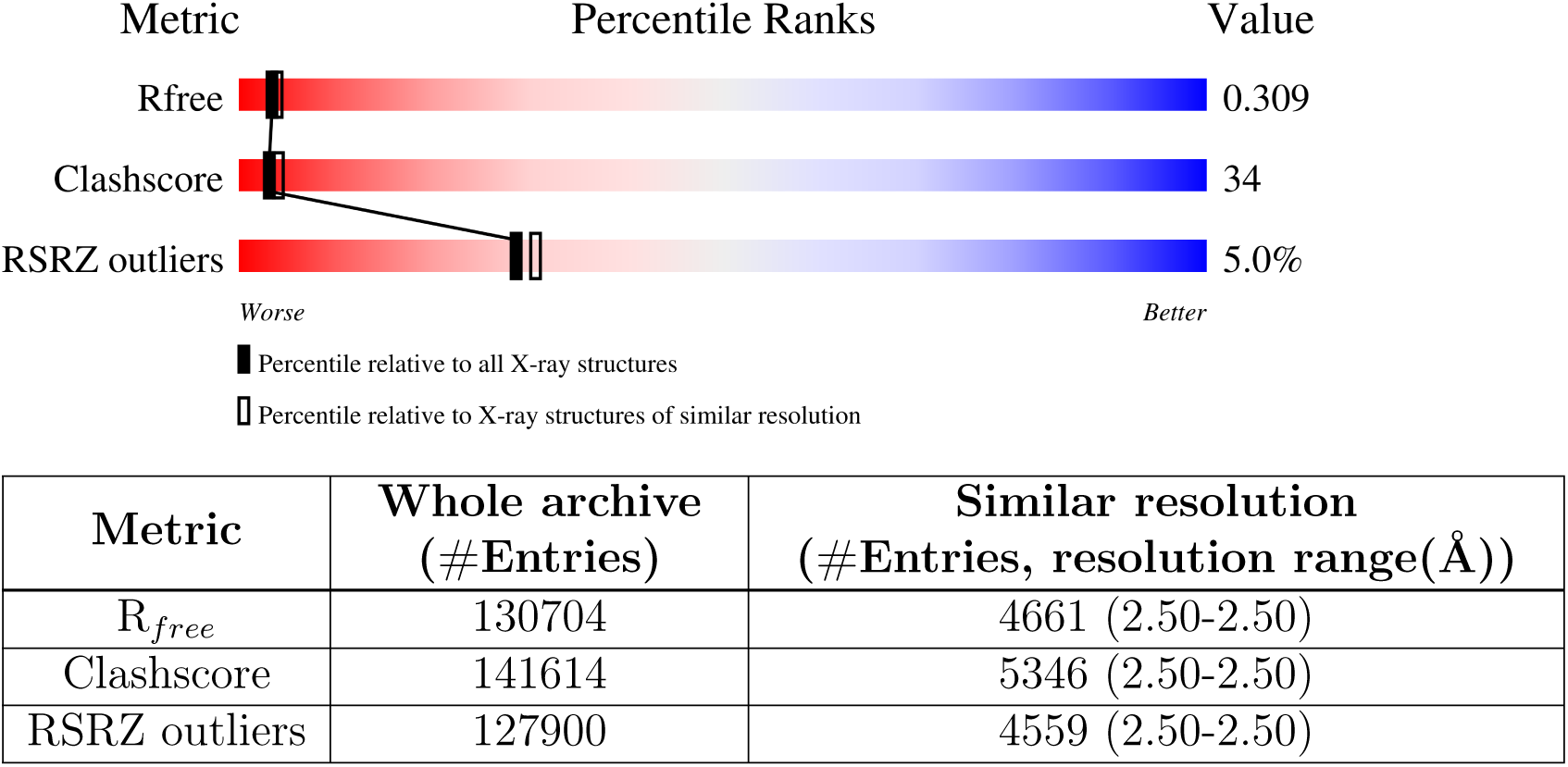

The table below summarises the geometric issues observed across the polymeric chains and their fit to the electron density. The red, orange, yellow and green segments of the lower bar indicate the fraction of residues that contain outliers for *>*=3, 2, 1 and 0 types of geometric quality criteria respectively. A grey segment represents the fraction of residues that are not modelled. The numeric value for each fraction is indicated below the corresponding segment, with a dot representing fractions *<*=5% The upper red bar (where present) indicates the fraction of residues that have poor fit to the electron density. The numeric value is given above the bar.

| Mol | Chain | Length | Quality of chain |
| --- | --- | --- | --- |
| 1 | A | 20 | 5%<br>65%<br>30%<br>5% |
| 2 | B | 5 | 40%<br>40%<br>20% |
| 2 | C | 5 | 80%<br>20% |

The following table lists non-polymeric compounds, carbohydrate monomers and non-standard residues in protein, DNA, RNA chains that are outliers for geometric or electron-density-fit criteria:

| Mol | Type | Chain | Res | Chirality | Geometry | Clashes | Electron density |
| --- | --- | --- | --- | --- | --- | --- | --- |
| 2 | QUK | B | 108 | - | - | X | X |
| 2 | QUK | C | 112 | - | - | - | X |
| 4 | MG | A | 105 | - | - | - | X |

### 2 Entry composition

There are 5 unique types of molecules in this entry. The entry contains 837 atoms, of which 306 are hydrogens and 0 are deuteriums.

In the tables below, the ZeroOcc column contains the number of atoms modelled with zero occupancy, the AltConf column contains the number of residues with at least one atom in alternate conformation and the Trace column contains the number of residues modelled with at most 2 atoms.

- Molecule 1 is a DNA chain called DNA (5’-D(*TP*GP*GP*GP*TP*TP*GP*GP*GP*TP* TP*GP*GP*GP*TP*TP*GP*GP*GP*T)-3’).

| Mol | Chain | Residues | Atoms |  |  |  |  |  | ZeroOcc | AltConf | Trace |
| --- | --- | --- | --- | --- | --- | --- | --- | --- | --- | --- | --- |
| 1 | A | 20 | Total | C | H | N | O | P | 0 | 0 | 1 |
|  |  |  | 592 | 180 | 204 | 72 | 117 | 19 |  |  |  |

- Molecule 2 is a protein called ACE-QUK-QUK-ZY9-QUK.

| Mol | Chain | Residues | Atoms |  |  |  |  | ZeroOcc | AltConf | Trace |
| --- | --- | --- | --- | --- | --- | --- | --- | --- | --- | --- |
| 2 | B | 5 | Total | C | H | N | O | 0 | 0 | 0 |
|  |  |  | 119 | 48 | 51 | 11 | 9 |  |  |  |
| 2 | C | 5 | Total | C | H | N | O | 0 | 0 | 0 |
|  |  |  | 119 | 48 | 51 | 11 | 9 |  |  |  |

- Molecule 3 is POTASSIUM ION (three-letter code: K) (formula: K).

| Mol | Chain | Residues | Atoms |  | ZeroOcc | AltConf |
| --- | --- | --- | --- | --- | --- | --- |
| 3 | A | 3 | Total | K | 0 | 0 |
|  |  |  | 3 | 3 |  |  |

- Molecule 4 is MAGNESIUM ION (three-letter code: MG) (formula: Mg).

| Mol | Chain | Residues | Atoms |  | ZeroOcc | AltConf |
| --- | --- | --- | --- | --- | --- | --- |
| 4 | A | 2 | Total | Mg | 0 | 0 |
|  |  |  | 2 | 2 |  |  |

- Molecule 5 is water.

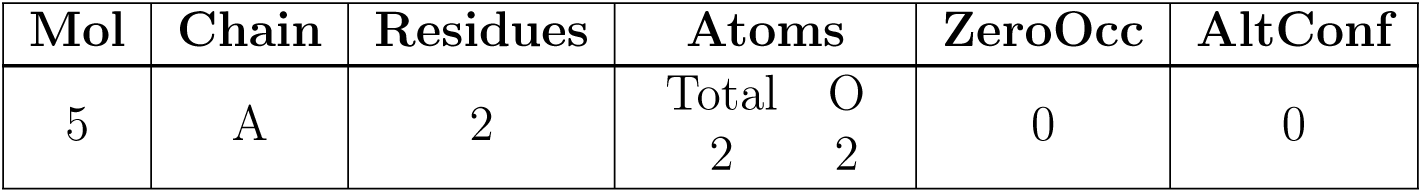

| Mol | Chain | Residues | Atoms |  | ZeroOcc | AltConf |
| --- | --- | --- | --- | --- | --- | --- |
| 5 | A | 2 | Total | O | 0 | 0 |
|  |  |  | 2 | 2 |  |  |

### 3 Residue-property plots

These plots are drawn for all protein, RNA, DNA and oligosaccharide chains in the entry. The first graphic for a chain summarises the proportions of the various outlier classes displayed in the second graphic. The second graphic shows the sequence view annotated by issues in geometry and electron density. Residues are color-coded according to the number of geometric quality criteria for which they contain at least one outlier: green = 0, yellow = 1, orange = 2 and red = 3 or more. A red dot above a residue indicates a poor fit to the electron density (RSRZ *>* 2). Stretches of 2 or more consecutive residues without any outlier are shown as a green connector. Residues present in the sample, but not in the model, are shown in grey.

- Molecule 1: DNA (5’-D(*TP*GP*GP*GP*TP*TP*GP*GP*GP*TP*TP*GP*GP*GP*TP*TP *GP*GP*GP*T)-3’)

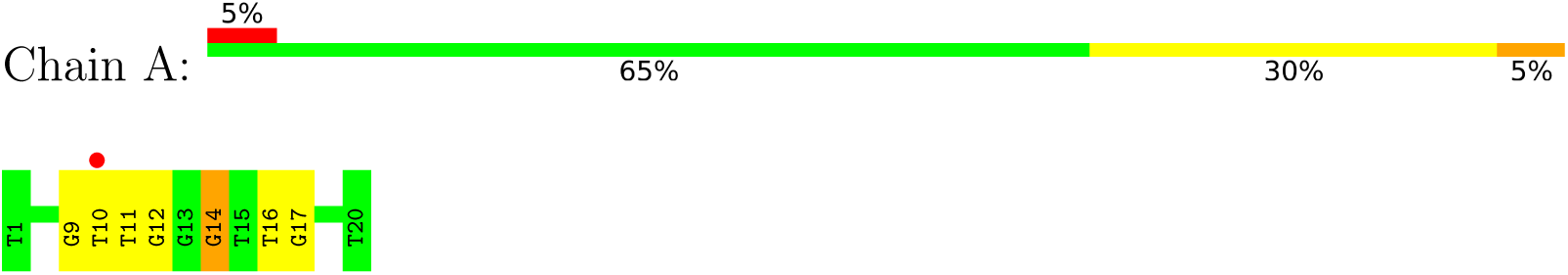

- Molecule 2: ACE-QUK-QUK-ZY9-QUK

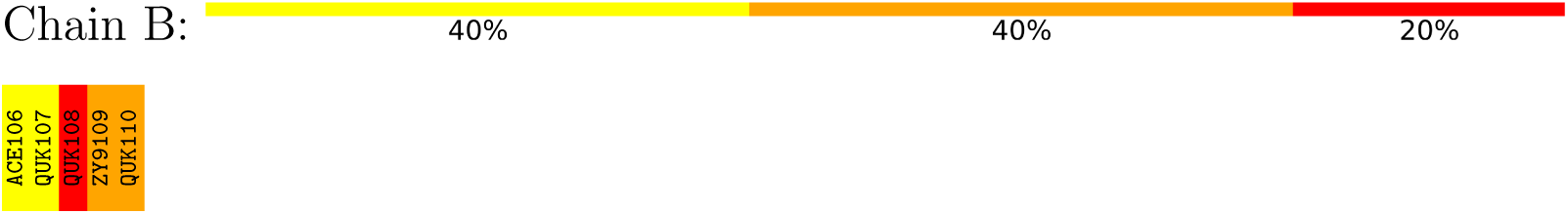

- Molecule 2: ACE-QUK-QUK-ZY9-QUK

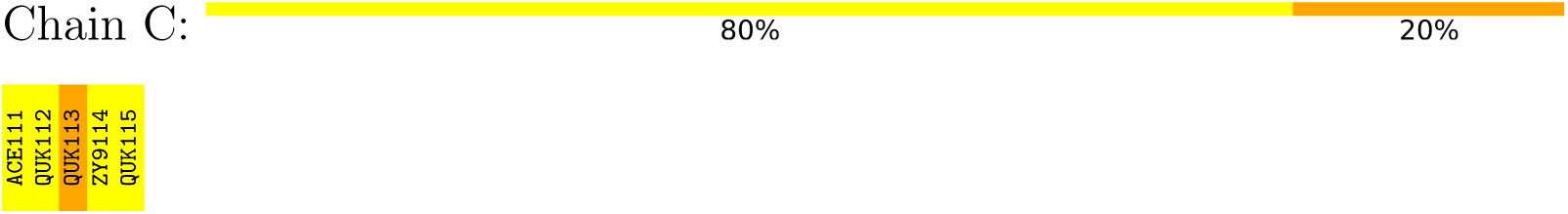

### 4 Data and refinement statistics

| Property | Value | Source |
| --- | --- | --- |
| Space group | P 42 | Depositor |
| Cell constants<br>a, b, c, $\alpha$, $\beta$, $\gamma$ | 32.71Å 32.71Å 61.01Å<br>90.00° 90.00° 90.00° | Depositor |
| Resolution (Å) | 30.50 – 2.51<br>32.71 – 2.51 | Depositor<br>EDS |
| % Data completeness<br>(in resolution range) | 99.1 (30.50-2.51)<br>99.9 (32.71-2.51) | Depositor<br>EDS |
| $R_{merge}$ | 0.15 | Depositor |
| $R_{sym}$ | (Not available) | Depositor |
| $\langle I/\sigma(I) \rangle$ <sup>1</sup> | 3.29 (at 2.51Å) | Xtriage |
| Refinement program | PHENIX 1.21rc1_5084 | Depositor |
| R, $R_{free}$ | 0.272 , 0.308<br>0.270 , 0.309 | Depositor<br>DCC |
| $R_{free}$ test set | 224 reflections (10.01%) | wwPDB-VP |
| Wilson B-factor (Å<sup>2</sup>) | 59.9 | Xtriage |
| Anisotropy | 0.454 | Xtriage |
| Bulk solvent $k_{sol}$(e/Å<sup>3</sup>), $B_{sol}$(Å<sup>2</sup>) | 0.28 , 40.8 | EDS |
| L-test for twinning<sup>2</sup> | $\langle \vert L\vert \rangle = 0.50$, $\langle L^2 \rangle = 0.35$ | Xtriage |
| Estimated twinning fraction | 0.479 for h,-k,-l | Xtriage |
| $F_o, F_c$ correlation | 0.93 | EDS |
| Total number of atoms | 837 | wwPDB-VP |
| Average B, all atoms (Å<sup>2</sup>) | 74.0 | wwPDB-VP |

<sup>2</sup>Theoretical values of $\langle |L| \rangle$ , $\langle L^2 \rangle$ for acentric reflections are 0.5, 0.333 respectively for untwinned datasets, and 0.375, 0.2 for perfectly twinned datasets.

Xtriage’s analysis on translational NCS is as follows: *The largest off-origin peak in the Patterson function is 10.48% of the height of the origin peak. No significant pseudotranslation is detected*.

^1^Intensities estimated from amplitudes.

^2^Theoretical values of *< |L| >*, *< L*^2^ *>* for acentric reflections are 0.5, 0.333 respectively for untwinned datasets, and 0.375, 0.2 for perfectly twinned datasets.

### 5 Model quality

#### 5.1 Standard geometry

Bond lengths and bond angles in the following residue types are not validated in this section: ZY9, K, MG, QUK, ACE

The Z score for a bond length (or angle) is the number of standard deviations the observed value is removed from the expected value. A bond length (or angle) with *|Z| >* 5 is considered an outlier worth inspection. RMSZ is the root-mean-square of all Z scores of the bond lengths (or angles).

| Mol | Chain | Bond lengths |  | Bond angles |  |
| --- | --- | --- | --- | --- | --- |
|  |  | RMSZ | #\|Z\| >5 | RMSZ | #\|Z\| >5 |
| 1 | A | 1.71 | 0/435 | 1.46 | 1/675 (0.1%) |

Chiral center outliers are detected by calculating the chiral volume of a chiral center and verifying if the center is modelled as a planar moiety or with the opposite hand.A planarity outlier is detected by checking planarity of atoms in a peptide group, atoms in a mainchain group or atoms of a sidechain that are expected to be planar.

| Mol | Chain | #Chirality outliers | #Planarity outliers |
| --- | --- | --- | --- |
| 2 | B | 0 | 1 |

There are no bond length outliers.

All (1) bond angle outliers are listed below:

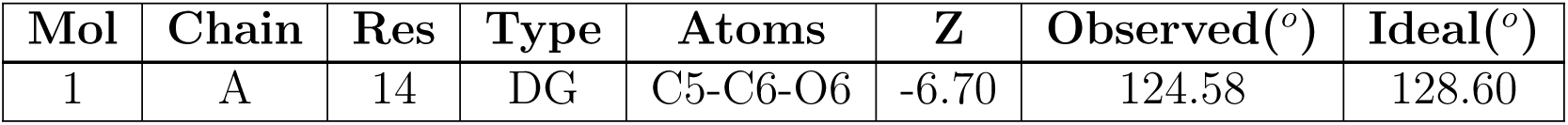

There are no bond length outliers. All (1) bond angle outliers are listed below:
| Mol | Chain | Res | Type | Atoms | Z | Observed(°) | Ideal(°) |
| --- | --- | --- | --- | --- | --- | --- | --- |
| 1 | A | 14 | DG | C5-C6-O6 | -6.70 | 124.58 | 128.60 |

There are no chirality outliers.

All (1) planarity outliers are listed below:

There are no chirality outliers. All (1) planarity outliers are listed below:
| Mol | Chain | Res | Type | Group |
| --- | --- | --- | --- | --- |
| 2 | B | 108 | QUK | Peptide |

#### 5.2 Too-close contacts

In the following table, the Non-H and H(model) columns list the number of non-hydrogen atoms and hydrogen atoms in the chain respectively. The H(added) column lists the number of hydrogen atoms added and optimized by MolProbity. The Clashes column lists the number of clashes within the asymmetric unit, whereas Symm-Clashes lists symmetry-related clashes.

| Mol | Chain | Non-H | H(model) | H(added) | Clashes | Symm-Clashes |
| --- | --- | --- | --- | --- | --- | --- |
| 1 | A | 388 | 204 | 204 | 13 | 0 |
| 2 | B | 68 | 51 | 3 | 18 | 0 |
| 2 | C | 68 | 51 | 3 | 1 | 0 |
| 3 | A | 3 | 0 | 0 | 0 | 0 |
| 4 | A | 2 | 0 | 0 | 0 | 0 |
| 5 | A | 2 | 0 | 0 | 0 | 0 |
| All | All | 531 | 306 | 210 | 24 | 0 |

The all-atom clashscore is defined as the number of clashes found per 1000 atoms (including hydrogen atoms). The all-atom clashscore for this structure is 34.

All (24) close contacts within the same asymmetric unit are listed below, sorted by their clash magnitude.

| Atom-1 | Atom-2 | Interatomic distance (Å) | Clash overlap (Å) |
| --- | --- | --- | --- |
| 2:B:106:ACE:H2 | 2:B:108:QUK:C | 1.87 | 1.05 |
| 2:B:109:ZY9:N | 2:B:110:QUK:OXT | 2.10 | 0.82 |
| 2:B:108:QUK:C | 2:B:110:QUK:OXT | 2.33 | 0.76 |
| 2:B:106:ACE:CH3 | 2:B:108:QUK:C | 2.64 | 0.75 |
| 2:B:106:ACE:H2 | 2:B:109:ZY9:N | 2.08 | 0.67 |
| 1:A:9:DG:H4' | 1:A:11:DT:C6 | 2.33 | 0.63 |
| 1:A:9:DG:C8 | 2:B:106:ACE:O | 2.52 | 0.63 |
| 2:B:106:ACE:H2 | 2:B:109:ZY9:C2 | 2.29 | 0.62 |
| 1:A:14:DG:C4 | 2:B:108:QUK:C4 | 2.83 | 0.61 |
| 2:B:106:ACE:H2 | 2:B:108:QUK:O | 2.01 | 0.60 |
| 1:A:9:DG:N7 | 2:B:106:ACE:O | 2.35 | 0.60 |
| 2:B:108:QUK:C10 | 2:B:110:QUK:OXT | 2.51 | 0.59 |
| 1:A:9:DG:C5 | 2:B:106:ACE:O | 2.63 | 0.51 |
| 2:B:108:QUK:C10 | 2:B:110:QUK:C | 2.92 | 0.48 |
| 2:C:111:ACE:H2 | 2:C:113:QUK:C | 2.43 | 0.47 |
| 1:A:9:DG:C5 | 2:B:106:ACE:C | 2.88 | 0.46 |
| 1:A:11:DT:H2'' | 1:A:12:DG:O5' | 2.16 | 0.46 |
| 1:A:14:DG:H4' | 1:A:16:DT:C6 | 2.52 | 0.45 |
| 1:A:14:DG:C5 | 2:B:108:QUK:C3 | 3.02 | 0.43 |
| 1:A:14:DG:C8 | 2:B:108:QUK:C5 | 3.02 | 0.43 |
| 2:B:106:ACE:CH3 | 2:B:108:QUK:O | 2.65 | 0.42 |
| 1:A:10:DT:C5' | 1:A:11:DT:OP2 | 2.68 | 0.41 |
| 1:A:16:DT:O2 | 1:A:17:DG:H2' | 2.21 | 0.40 |
| 1:A:14:DG:C4 | 2:B:108:QUK:C3 | 3.05 | 0.40 |

There are no symmetry-related clashes.

#### 5.3 Torsion angles

##### 5.3.1 Protein backbone

There are no protein backbone outliers to report in this entry.

##### 5.3.2 Protein sidechains

There are no protein residues with a non-rotameric sidechain to report in this entry.

##### 5.3.3 RNA

There are no RNA molecules in this entry.

#### 5.4 Non-standard residues in protein, DNA, RNA chains

8 non-standard protein/DNA/RNA residues are modelled in this entry.

In the following table, the Counts columns list the number of bonds (or angles) for which Mogul statistics could be retrieved, the number of bonds (or angles) that are observed in the model and the number of bonds (or angles) that are defined in the Chemical Component Dictionary. The Link column lists molecule types, if any, to which the group is linked. The Z score for a bond length (or angle) is the number of standard deviations the observed value is removed from the expected value. A bond length (or angle) with *|Z| >* 2 is considered an outlier worth inspection. RMSZ is the root-mean-square of all Z scores of the bond lengths (or angles).

| Mol | Type | Chain | Res | Link | Bond lengths |  |  | Bond angles |  |  |
| --- | --- | --- | --- | --- | --- | --- | --- | --- | --- | --- |
| | | | | | Counts | RMSZ | $\#\vert Z\vert > 2$ | Counts | RMSZ | $\#\vert Z\vert > 2$ |
| 2 | ZY9 | C | 114 | 2 | 10,10,11 | 0.72 | 1 (10%) | 11,12,14 | 0.91 | 1 (9%) |
| 2 | ZY9 | B | 109 | 2 | 10,10,11 | 0.83 | 1 (10%) | 11,12,14 | 1.10 | 0 |
| 2 | QUK | B | 107 | 2 | 19,19,20 | 0.50 | 0 | 22,25,27 | 0.81 | 1 (4%) |
| 2 | QUK | B | 110 | 2 | 20,20,20 | 0.66 | 0 | 26,27,27 | 0.95 | 2 (7%) |
| 2 | QUK | B | 108 | 2 | 19,19,20 | 1.02 | 2 (10%) | 22,25,27 | 2.66 | 3 (13%) |
| 2 | QUK | C | 112 | 2 | 19,19,20 | 0.44 | 0 | 22,25,27 | 0.92 | 2 (9%) |
| 2 | QUK | C | 113 | 2 | 19,19,20 | 0.66 | 1 (5%) | 22,25,27 | 1.63 | 4 (18%) |
| 2 | QUK | C | 115 | 2 | 20,20,20 | 0.45 | 0 | 26,27,27 | 0.95 | 2 (7%) |

In the following table, the Chirals column lists the number of chiral outliers, the number of chiral centers analysed, the number of these observed in the model and the number defined in the Chemical Component Dictionary. Similar counts are reported in the Torsion and Rings columns. ‘-’ means no outliers of that kind were identified.

| Mol | Type | Chain | Res | Link | Chirals | Torsions | Rings |
| --- | --- | --- | --- | --- | --- | --- | --- |
| 2 | ZY9 | C | 114 | 2 | - | 0/4/4/6 | 0/1/1/1 |
| 2 | ZY9 | B | 109 | 2 | - | 0/4/4/6 | 0/1/1/1 |
| 2 | QUK | B | 107 | 2 | - | 3/7/7/9 | 0/2/2/2 |
| 2 | QUK | B | 110 | 2 | - | 2/9/9/9 | 0/2/2/2 |
| 2 | QUK | B | 108 | 2 | - | 6/7/7/9 | 0/2/2/2 |
| 2 | QUK | C | 112 | 2 | - | 4/7/7/9 | 0/2/2/2 |
| 2 | QUK | C | 113 | 2 | - | 2/7/7/9 | 0/2/2/2 |
| 2 | QUK | C | 115 | 2 | - | 1/9/9/9 | 0/2/2/2 |

All (5) bond length outliers are listed below:

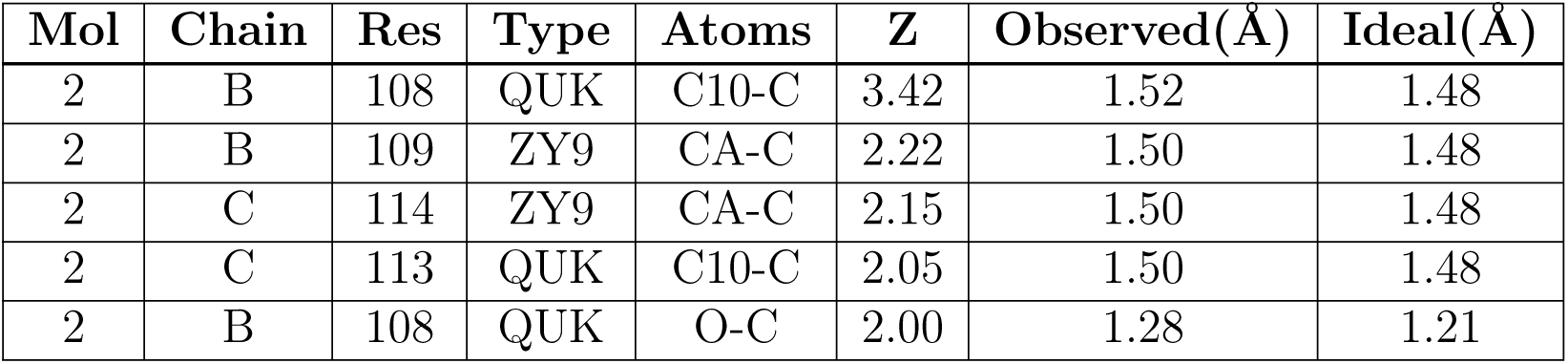

All (15) bond angle outliers are listed below:

| Mol | Chain | Res | Type | Atoms | Z | Observed(°) | Ideal(°) |
| --- | --- | --- | --- | --- | --- | --- | --- |
| 2 | B | 108 | QUK | C9-C10-C | -8.70 | 113.70 | 121.23 |
| 2 | B | 108 | QUK | O-C-C10 | -6.34 | 118.22 | 124.22 |
| 2 | B | 108 | QUK | C-C10-N11 | 5.60 | 120.15 | 114.66 |
| 2 | C | 113 | QUK | C9-C10-C | -4.11 | 117.67 | 121.23 |
| 2 | C | 113 | QUK | O-C-C10 | -3.39 | 121.01 | 124.22 |
| 2 | B | 110 | QUK | C3-CA-N | 3.00 | 126.35 | 120.36 |
| 2 | C | 113 | QUK | C-C10-N11 | 2.99 | 117.60 | 114.66 |
| 2 | C | 113 | QUK | CE-CD-CG | 2.83 | 123.73 | 113.61 |
| 2 | C | 115 | QUK | C3-CA-N | 2.69 | 125.74 | 120.36 |
| 2 | C | 112 | QUK | O-C-C10 | -2.51 | 121.85 | 124.22 |
| 2 | C | 115 | QUK | O-C-C10 | -2.47 | 116.23 | 121.24 |
| 2 | C | 114 | ZY9 | O-C-CA | -2.30 | 122.05 | 124.22 |
| 2 | B | 110 | QUK | C7-CA-N | -2.14 | 113.98 | 118.07 |
| 2 | B | 107 | QUK | O-C-C10 | -2.08 | 122.25 | 124.22 |
| 2 | C | 112 | QUK | C9-C10-C | -2.02 | 119.48 | 121.23 |

There are no chirality outliers.

All (18) torsion outliers are listed below:

There are no chirality outliers. All (18) torsion outliers are listed below:
| Mol | Chain | Res | Type | Atoms |
| --- | --- | --- | --- | --- |
| 2 | B | 108 | QUK | O-C-C10-N11 |
| 2 | B | 108 | QUK | O-C-C10-C9 |
| 2 | C | 113 | QUK | O-C-C10-C9 |
| 2 | C | 115 | QUK | CE-CD-CG-OB |
| 2 | C | 112 | QUK | C6-C8-OB-CG |
| 2 | C | 112 | QUK | CE-CD-CG-OB |
| 2 | B | 110 | QUK | CE-CD-CG-OB |
| 2 | C | 112 | QUK | C9-C8-OB-CG |
| 2 | B | 108 | QUK | C6-C8-OB-CG |
| 2 | C | 112 | QUK | CG-CD-CE-N1 |
| 2 | B | 108 | QUK | CG-CD-CE-N1 |
| 2 | B | 107 | QUK | CD-CG-OB-C8 |
| 2 | B | 108 | QUK | C9-C8-OB-CG |
| 2 | C | 113 | QUK | CE-CD-CG-OB |
| 2 | B | 108 | QUK | CD-CG-OB-C8 |
| 2 | B | 107 | QUK | CE-CD-CG-OB |
| 2 | B | 107 | QUK | CG-CD-CE-N1 |
| 2 | B | 110 | QUK | CG-CD-CE-N1 |

There are no ring outliers.

4 monomers are involved in 15 short contacts:

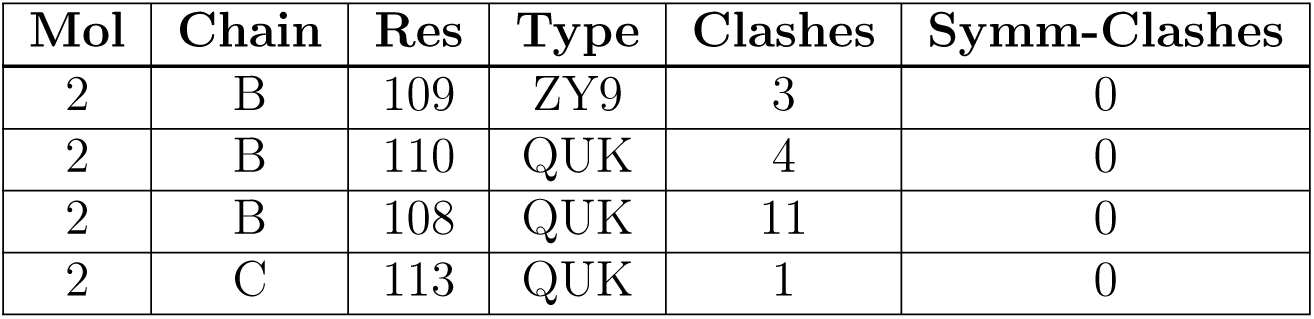

#### 5.5 Carbohydrates

There are no monosaccharides in this entry.

#### 5.6 Ligand geometry

Of 5 ligands modelled in this entry, 5 are monoatomic - leaving 0 for Mogul analysis.

There are no bond length outliers.

There are no bond angle outliers.

There are no chirality outliers.

There are no torsion outliers.

There are no ring outliers.

No monomer is involved in short contacts.

#### 5.7 Other polymers

There are no such residues in this entry.

#### 5.8 Polymer linkage issues

There are no chain breaks in this entry.

### 6 Fit of model and data

#### 6.1 Protein, DNA and RNA chains

In the following table, the column labelled ‘#RSRZ*>* 2’ contains the number (and percentage) of RSRZ outliers, followed by percent RSRZ outliers for the chain as percentile scores relative to all X-ray entries and entries of similar resolution. The OWAB column contains the minimum, median, 95*^th^* percentile and maximum values of the occupancy-weighted average B-factor per residue. The column labelled ‘Q*<* 0.9’ lists the number of (and percentage) of residues with an average occupancy less than 0.9.

| Mol | Chain | Analysed | <RSRZ> | #RSRZ > 2 | OWAB(Å<sup>2</sup>) | Q < 0.9 |
| --- | --- | --- | --- | --- | --- | --- |
| 1 | A | 20/20 (100%) | 0.33 | 1 (5%) 28 30 | 47, 59, 90, 99 | 0 |
| 2 | B | 0/5 | - | - | - | - |
| 2 | C | 0/5 | - | - | - | - |
| All | All | 20/30 (66%) | 0.33 | 1 (5%) 28 30 | 47, 59, 90, 99 | 0 |

All (1) RSRZ outliers are listed below:

| Mol | Chain | Res | Type | RSRZ |
| --- | --- | --- | --- | --- |
| 1 | A | 10 | DT | 2.6 |

#### 6.2 Non-standard residues in protein, DNA, RNA chains

In the following table, the Atoms column lists the number of modelled atoms in the group and the number defined in the chemical component dictionary. The B-factors column lists the minimum, median, 95*^th^* percentile and maximum values of B factors of atoms in the group. The column labelled ‘Q*<* 0.9’ lists the number of atoms with occupancy less than 0.9.

| Mol | Type | Chain | Res | Atoms | RSCC | RSR | B-factors(Å<sup>2</sup>) | Q < 0.9 |
| --- | --- | --- | --- | --- | --- | --- | --- | --- |
| 2 | ZY9 | C | 114 | 10/11 | 0.56 | 0.35 | 67,83,102,108 | 0 |
| 2 | QUK | B | 108 | 18/19 | 0.66 | 0.67 | 61,94,133,141 | 0 |
| 2 | QUK | B | 110 | 19/19 | 0.71 | 0.35 | 63,81,124,124 | 0 |
| 2 | QUK | C | 113 | 18/19 | 0.73 | 0.40 | 53,87,126,134 | 0 |
| 2 | QUK | C | 112 | 18/19 | 0.76 | 0.46 | 61,100,161,161 | 0 |
| 2 | ZY9 | B | 109 | 10/11 | 0.76 | 0.27 | 60,74,95,101 | 0 |
| 2 | QUK | C | 115 | 19/19 | 0.76 | 0.37 | 54,82,118,118 | 0 |
| 2 | QUK | B | 107 | 18/19 | 0.78 | 0.36 | 48,77,169,169 | 0 |

#### 6.3 Carbohydrates

There are no monosaccharides in this entry.

#### 6.4 Ligands

In the following table, the Atoms column lists the number of modelled atoms in the group and the number defined in the chemical component dictionary. The B-factors column lists the minimum, median, 95*^th^* percentile and maximum values of B factors of atoms in the group. The column labelled ‘Q*<* 0.9’ lists the number of atoms with occupancy less than 0.9.

| Mol | Type | Chain | Res | Atoms | RSCC | RSR | B-factors(Å<sup>2</sup>) | Q<0.9 |
| --- | --- | --- | --- | --- | --- | --- | --- | --- |
| 4 | MG | A | 105 | 1/1 | 0.55 | 0.48 | 78,78,78,78 | 0 |
| 3 | K | A | 103 | 1/1 | 0.65 | 0.15 | 28,28,28,28 | 1 |
| 3 | K | A | 102 | 1/1 | 0.69 | 0.13 | 41,41,41,41 | 0 |
| 3 | K | A | 101 | 1/1 | 0.71 | 0.12 | 42,42,42,42 | 0 |
| 4 | MG | A | 104 | 1/1 | 0.88 | 0.48 | 79,79,79,79 | 0 |

#### 6.5 Other polymers

There are no such residues in this entry.

## Footnotes

1 Vallade M., Sai Reddy P., Fischer L., Huc I., Enhancing aromatic foldamer helix dynamics to probe interactions with protein surfaces, *Eur. J. Org. Chem*., **2018**, 5489.

